# Systematic structural and functional analysis of metal-binding sites in native proteomes

**DOI:** 10.1101/2025.11.10.687516

**Authors:** JP. Quast, A. Pagotto, RJ. Separovich, M. Dünnebacke, L. Gillet, X. Luo, E. Züger, G. Magnusson, MSP. Correia, C. Ciofalo, C. Andreini, A. Rosato, P. Beltrao, N. Zamboni, N. de Souza, P. Picotti

## Abstract

Metal-binding proteins (MBPs) are widespread and critical for many biological processes. Proteomics-based approaches have identified MBPs in various systems but typically cannot determine the location of metal-binding sites or shed light on the functional roles of metals. Thus, a systems-wide map of metal binding sites and an understanding of the consequences of metal binding for protein function are still lacking. Here, we used limited proteolysis-coupled mass spectrometry (LiP-MS) combined with chelator treatment to globally map metal-protein interactions and binding sites in native protein lysates. We applied the pipeline, named Metal-LiP, to *Escherichia coli,* identified 981 peptides on 297 proteins that exhibited dose responsiveness to chelator treatment, and quantified the relative metal-binding affinity. These proteins are strongly enriched for known or predicted MBPs, based on our novel compiled ground truth dataset of previously annotated (n=1013) and predicted (n=850) MBPs. Further, chelator-responsive peptides mapped at or near annotated metal-binding sites in 3D protein structures, showing that Metal-LiP can identify metal-binding sites, and provided novel experimental evidence for previously predicted sites. Notably, Metal-LiP identified 225 novel candidate MBPs and we validated five of these novel MBPs (GatZ, AhpC, GrxB, MsrC, TrhO) using inductively coupled plasma-mass spectrometry to detect metal binding in purified proteins. Finally, we used orthogonal approaches to probe the broad functional consequences of metal binding. Using a combination of Metal-LiP, thermal proteome profiling, size-exclusion chromatography coupled to mass spectrometry, and structural analyses, we identified cases where metal binding affects protein stability, modulates the assembly states of protein complexes, or is likely to affect enzymatic activity. We thereby propose functional roles for metal binding in 98 of our Metal-LiP hits and in 225 proteins in total. In summary, we describe an approach to identify *bona fide* MBPs with binding site-level resolution across the proteome. We have combined it with other methods to identify 424 candidate MBPs in the *E coli* proteome, to map known and novel candidate metal-binding sites for 297 of these proteins, and to suggest functional roles of metal binding.

## Introduction

Metal-binding proteins (MBPs) are widespread in cellular systems and make up ∼ 40% of proteins with known structure^1,2^. Metals and MBPs are involved in almost all (∼ 80%) metabolic pathways^5^, and their dysregulation causes human disease^6^.

A wide range of experimental approaches have been used to identify MBPs. Structure-based analyses (*e.g.,* X-ray crystallography, cryo-electron microscopy) can locate metal ions and their binding sites within single proteins^7^. However, these techniques study one protein at a time and can identify artifactual binding sites due to non-native buffer conditions during protein purification^4,7^. More recently, proteomics-based approaches have been developed to systematically profile MBPs in a single experiment^8–11^. For example, thermal proteome profiling (TPP) exploits the characteristic shift in a protein’s melting temperature upon ligand binding to quantify its binding affinity^12^. Thermal profiling studies using chelator treatments on protein lysates have identified hundreds of MBPs and their metal-binding specificities in different model systems^10,11^. However, TPP cannot determine the precise metal-binding site, thus precluding insights into the structural features regulating metal-protein interactions. Further, while selective binding of metal ions to MBPs is known to affect catalysis^2^, structural stability^3^, and protein interactions^4^, previous work has not systematically investigated the functional effects but is based on studies of a few individual proteins. Knowledge of the metal-protein interactome would therefore be greatly expanded by the development of approaches to globally map metal-protein interactions and binding sites, and to study the functional consequences of these interactions, under physiologically relevant conditions.

Limited proteolysis-coupled mass spectrometry (LiP-MS) is a structural proteomics technique that captures changes in protease accessibility of protein targets with peptide-level resolution and on a proteome-wide scale^13^. Since ligand binding alters the protease accessibility of proteins, we have previously applied LiP-MS to systematically map binding sites for protein-protein^14^, protein-drug^15^, and protein-metabolite^16^ interactions in thousands of proteins in complex lysates. Here, we hypothesized that binding or removal of metal ions is sufficient to alter the protease accessibility of metal-binding sites. We assessed the application of LiP-MS to the large-scale study of metal-binding proteins and sites, leveraging changes in proteolytic accessibility caused by chelator-induced metal removal. Our modified pipeline, termed Metal-LiP, involves size-exclusion filtration to remove free metal ions from native lysates, followed by chelator treatment in twelve concentrations to extract protein-bound metals in a dose-responsive manner. Quantitative LiP-MS was then performed to derive peptide-level chelator-response curves, which were used to identify candidate metal-binding proteins and sites across the proteome.

We applied Metal-LiP to generate a proteome-wide map of MBPs and metal-binding sites in *Escherichia coli*. In total, we identified 981 peptides in 297 proteins that exhibited dose responsiveness for at least one of six chelator (EDTA, EGTA, DTPA, TETA, TPEN, DiP) treatments. To benchmark our Metal-LiP hits, we compiled a ground-truth dataset of around 1000 previously annotated MBPs; we further compiled a set of 850 predicted MBPs based on multiple predictors. We found that the Metal-LiP hits are strongly enriched for annotated or predicted MBPs and are involved in metal-associated molecular functions (*e.g.,* metal-binding, ribosome, RNA binding). Importantly, chelator-responsive peptides are spatially proximal to known or predicted metal-binding sites in 3D protein structures, thus confirming that our approach identified MBPs with site-level resolution. We then assessed the molecular consequences of metal removal. Quantitative thermal proteome profilings of EDTA-treated bacterial lysates revealed a role of metals in protein stabilization or destabilization for a subset of our candidate MBPs Further, SEC-MS analyses conducted in parallel highlighted cases where metal binding modulates the assembly state of protein complexes and structural analyses revealed cases where metal binding is likely to modulate catalytic processes. We identified 424 candidate MBPs overall, of which 225 were novel, detected known and novel candidate metal-binding sites for 297 proteins,, and orthogonally validated metal binding for five out of six tested candidates (GatZ, AhpC, GrxB, MsrC, TrhO) in experiments with purified proteins *in vitro*. Thus, we show that Metal-LiP can globally profile metal-induced structural changes in proteomes and will be useful for MBP and binding-site mapping in other cellular systems.

## Results

### Metal-LiP, a LiP-MS-based workflow to investigate the metal-binding proteome

To profile protein-metal interactions in native proteomes, we developed a novel workflow termed Metal-LiP that combines a dose-response treatment of native lysates with a chelator (Fig. 1A), size-exclusion filtration and LiP-MS (Fig 1B). Briefly, *E. coli* grown in modified M9 medium were lysed, followed by filtration to remove free metal ions. We adjusted the protein concentration of the lysate to 1 µg/µL and treated it with a twelve-dosage series of six different chelators (Fig. 1A) to remove metals from proteins in a dose-dependent manner, followed by a second filtration step to remove the metal-chelator complexes. Samples were then processed using LiP-MS, generating differential cleavage patterns depending on metal occupancy. We then identified candidate metal-binding proteins as those that showed dose-response behavior upon chelator treatment and extracted information such as half maximal effective concentration (EC_50_) values from the dose-response curves.

**Fig. 1:**
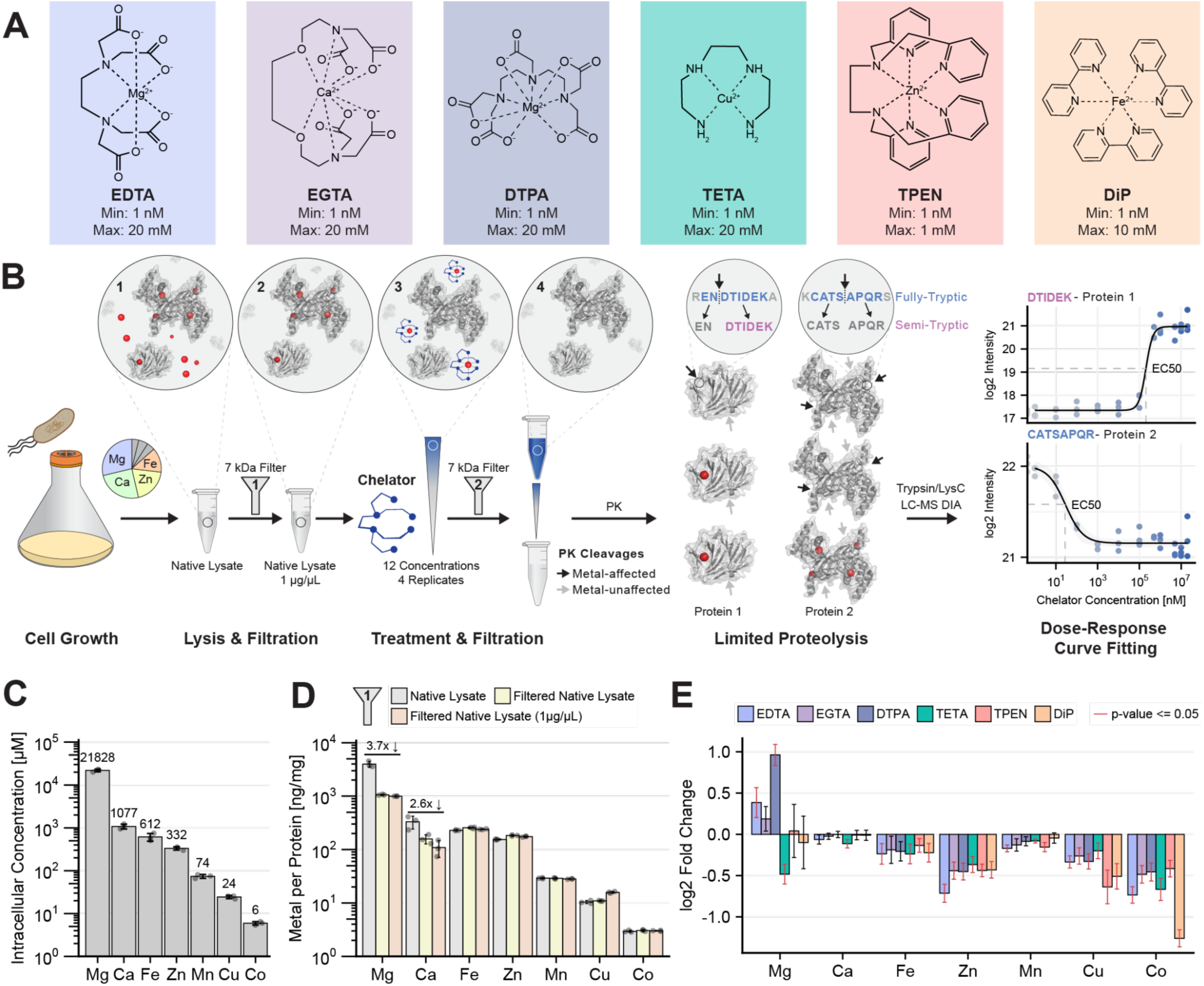
Development of Metal-LiP to study metal-binding proteomes. **A** The panel of chelators used in the LiP-MS workflow. Each chelator was applied in a dosage series; the lowest and highest concentrations are indicated. **B** LiP-MS workflow for the identification and characterisation of metal-binding proteins. Inserts 1-4 depict the likely molecular events at each step; red spheres represent metal ions and the chelator is schematised. In insert 1, metal ions are present in both free and protein-bound form. In insert 2, free metal ions have been removed by filtration. In insert 3, the chelator has stripped metal ions off of proteins. In insert 4, the chelator-metal complexes have been removed by a second filtration step. PK: Proteinase K **C** Intracellular metal concentrations of *E. coli* grown in modified M9 medium determined by ICP-MS. **D** Size filtration removes some metal ions from protein lysates. Plotted is the amount of metal by protein in the native lysate, upon the first filtration step, and in the protein concentration-adjusted samples. Buffer metal concentrations were subtracted from all samples. **E** Chelator effect on metal abundance in samples after the second Metal-LiP filtration step. The bar plot shows the log2 fold change for the indicated metals after chelator treatment (max concentration, see Fig. 1A) compared to vehicle control. Error bars show the 95% confidence interval. Red error bars indicate a significant change (*p*-value ≤ 0.05) as determined by a Welch’s t-test. All metal abundances in panels C-E were determined by ICP-MS in triplicate.

We developed Metal-LiP by optimising the standard LiP workflow^17^ to be compatible with the use of chelators, and applied it to the *E. coli* proteome. First, we modified the composition of the M9 growth medium to more accurately reflect physiological metal concentrations (Supplementary Fig. 1 and Methods). We confirmed that bacteria grown in this modified M9 medium had intracellular metal concentrations in the expected range^18,19^ (Fig. 1C). Second, we included an initial filtration step of the native lysate to remove any free metals that could reduce the effectiveness of the chelator to disrupt protein-metal interactions, due to competition (Fig. 1B and Supplementary Fig. 2A). This filtration step reduced the concentration of magnesium and calcium ions by 3.7 and 2.9 fold, respectively (Fig. 1D and Supplementary Fig. 2B). The effect on these metals is consistent with the fact that they form the weakest interactions according to the Irving-Williams series^20^ and have the highest intracellular concentrations, while metals such as zinc and copper tightly bind proteins and are not present as free hydrated ions inside cells^21^.

We introduced a second size-exclusion filtration step immediately after the chelator treatment and prior to protease addition to remove metal-chelator complexes from the sample (Fig. 1B and Supplementary Fig. 2A) and to avoid effects of the chelator treatment on either protease activity or the MS measurement in the LiP-MS step (Supplementary Fig. 2C, D and Methods). We confirmed that filtration reduced chelator concentrations in the lysate using LC-MS measurements (Supplementary Fig. 2E). The hydrophobic chelators TPEN and DiP were more efficiently removed (>99%) compared to the hydrophilic chelators EDTA, EGTA and DTPA (∼80%). Unfortunately, we could not consistently quantify TETA. Based on these results, all five quantified chelators were reduced below the level at which we saw effects on Proteinase K (PK) activity or ion suppression (Supplementary Fig. 2C, D). To further reduce any potential effect on trypsin activity, we added 3 mM CaCl_2_ to all samples at the trypsin digestion step.

We confirmed that chelator treatment removed metals from proteins by using ICP-MS to measure metal ion concentrations after the second Metal-LiP filtration step. We saw a clear decrease of cobalt, copper and zinc concentrations in samples treated with all 6 chelators, compared to vehicle-treated controls (Fig. 1E). Iron and manganese showed a small reduction, indicating that these metals are less accessible to chelators. The largest decrease in absolute concentration was observed for zinc due to higher absolute levels (Supplementary Fig. 2B). Counterintuitively, we noted an increase in magnesium concentration after chelator treatment relative to the control, especially for EDTA and DTPA. This may be due to the large amount of residual free magnesium in the samples at the time of chelator treatment (i.e., after the first filtration) (Supplementary Fig. 2B), such that less efficient removal of magnesium-chelator complexes relative to free magnesium would lead to an apparent increase in magnesium in treated samples compared to vehicle controls (Supplementary Fig. 2E). While some chelators had stronger effects on metal concentration than others, and this further differed slightly between metals, the metal-depletion effect was generally consistent across chelators (see Discussion).

An alternative approach to the use of chelators to identify protein-metal interactions is to directly treat lysates with metal ions. We decided against this approach for multiple reasons. First, many metal salts have a low solubility at neutral pH (e.g. FeCl_3_), which reduces the usable metal concentration range. Second, we observed a strong inhibition of PK activity by several metals (Supplementary Fig. 3A). Third, binding of free metal ions to an unoccupied metal-binding site might not be favoured since associative metal transfer is preferred *in vivo*^22^. Further, for some MBPs, metal ions might need to be incorporated into the structure during the protein folding process^23^. Fourth, a LiP-MS experiment to identify structurally-altered proteins upon adding a magnesium dosage series (Supplementary Fig. 3B) identified about 7000 peptides (15%) and 1000 proteins (54%) as hits (Supplementary Fig. 3C), and no enrichment for metal-binding proteins. This indicates an unspecific effect (Supplementary Fig. 3D). We therefore chose the chelator-based workflow for Metal-LiP.

In summary, we have developed a limited proteolysis-based method, termed Metal-LiP, to identify metal-binding proteins in any biological system of interest.

### Metal-LiP identifies known and novel MBPs in *E. coli*

To identify metal-binding proteins in the *E. coli* proteome, we treated cell lysates with a 12-dosage series of each of the six chelators (Fig. 1A), followed by LiP-MS. To identify hits, we used a computational workflow modified from our previously described approach^15^. In every experiment, we fitted a 4-parameter log logistic regression curve to peptide precursor intensities, after filtering for peptide precursors (hereafter referred to as precursors) found in at least 2 replicates in at least 4 chelator concentrations. Significant precursors were then defined as those with a Pearson correlation between the dose-response curve and the measured data points of r ≥ 0.8 and an adjusted p-value ≤ 0.05 (ANOVA) (Supplementary Fig. 4A, B, C; Methods). To minimise false positives, curves were further assessed qualitatively; cases with unrealistic fitting parameters or patterns of missing data, or poor curve fits, were excluded from further analysis. Finally, we further refined the hit list by checking for hits that were supported by multiple lines of evidence (Supplementary Fig. 4D, E), and by recovering precursors that were close to significance (r ≥ 0.5) in a given condition (i.e., chelator treatment) if they were significantly changing in other conditions (Supplementary Fig. 4F).

We identified between 252 (DiP) and 882 (TETA) precursors that showed good dose responsive behavior, depending on the chelator (Fig. 2A), with most chelators (EDTA, EGTA, DTPA and TPEN) yielding roughly 500-600 hits each. Our various hit refinement steps (Methods) added around 200-300 significant precursors for each chelator (Supplementary Figure 5A-C) and allowed some low-confidence precursors to be orthogonally verified (Supplementary Fig. 5D). In total, we observed 297 proteins with significant changes in proteolytic accessibility across all chelators (Supplementary Table 1),

**Fig. 2:**
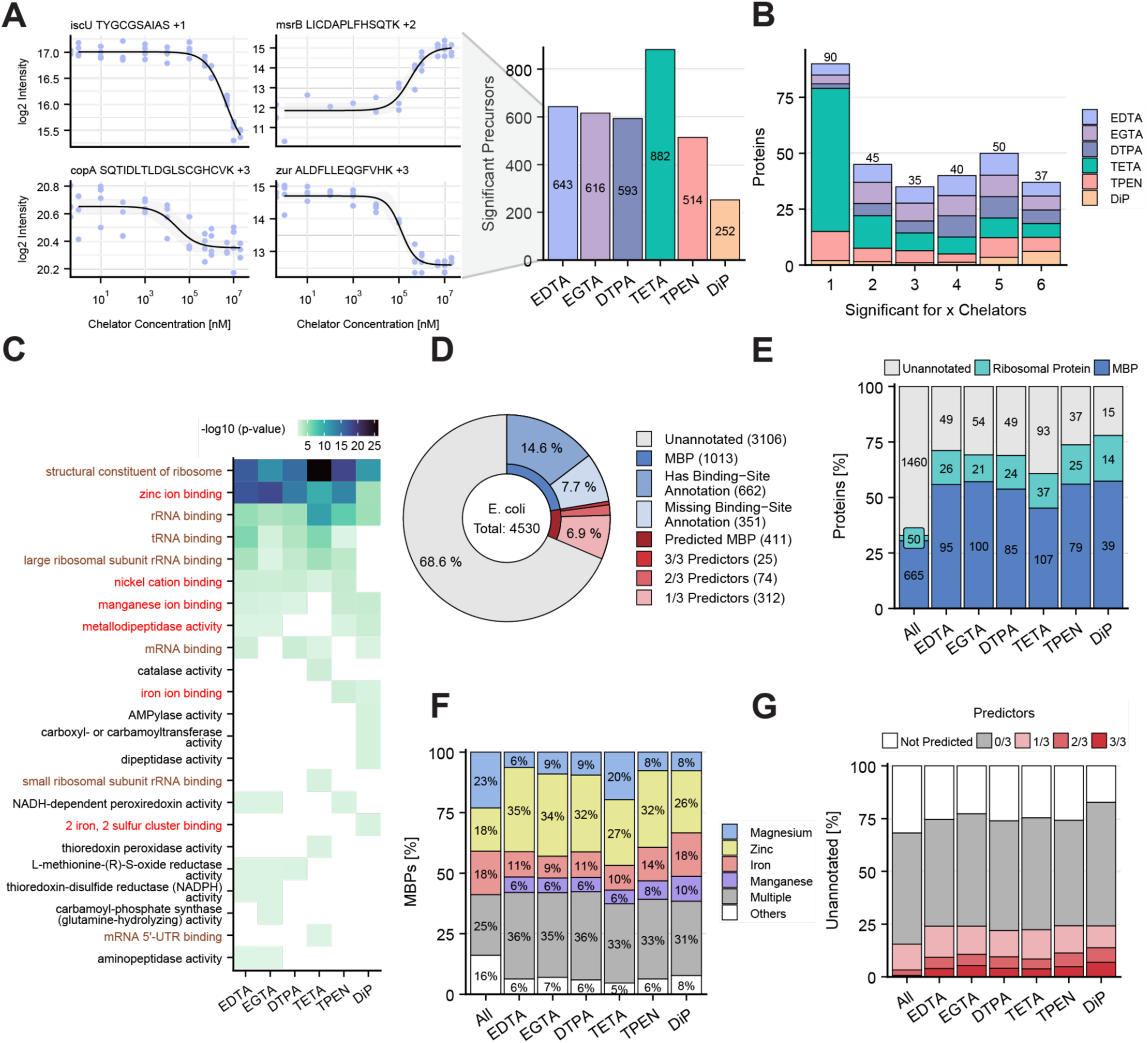
Metal-LiP-MS systematically identifies metal-binding proteins in the *E. coli* proteome. **A** The bar plot (right) shows the final number of significant hits (i.e., precursors that showed good dose-response curves and passed our filtration and validation criteria) for each chelator. The dose-response curves (left) are four examples of responses to EDTA for the indicated precursors, which map to the known MBPs IscU, msrB, CopA and Zur. **B** The stacked bar plot shows the overlap of hit proteins for the six chelators. Numbers above each bar indicate the number of proteins with the corresponding overlap. **C** Molecular function gene ontology enrichment for hit proteins from all chelators. The significance cutoff for the Fisher’s exact test is a p-value of 0.01. Only terms with at least two significant proteins are plotted. Red terms are related to metal-binding, brown terms to RNA-binding. **D** Metal-binding proteins (MBPs) and metal-binding protein predictions for the *E. coli* proteome. MBPs were extracted from UniProt and QuickGO. The category is further divided into proteins with and without an annotated binding site. Metal-binding predictions (Metal3D, Master of Metals (MOM) and GASS-Metal) are shown only for proteins that are not known MBPs. **E** Enrichment of annotated MBPs among Metal-LiP hits for each chelator. “All” indicates all detected proteins. Ribosomal proteins that are not annotated MBPs are shown in addition, since the ribosome is known to be sensitive to metal depletion. The number of significant proteins in each category is indicated. For each chelator the increase in the MBP fraction is statistically significant (p-value < 1e-5) as assessed by Fisher’s exact test. **F** Metal-binding specificity for annotated MBPs (from panel D) among Metal-LiP hits for each chelator. The “Multiple” category contains proteins for which multiple metals are annotated. The “Others” category includes all metal annotations with a frequency below 5%. **G** MBP prediction for unannotated MBPs (from panel D) among Metal-LiP hits for each chelator. The “Not Predicted” category means that no prediction was performed for the given protein, typically because the AlphaFold prediction did not pass our confidence cutoff, while the “0/3” category means that none of the predictors predicted a metal-binding site in the given protein.

For each hit protein, we calculated the chelator concentration corresponding to half-maximal proteolytic accessibility change (i.e., the EC_50_ value). This provided an estimate of the apparent binding strength of the metal ions to proteins in the cell extract and allowed direct comparison of chelator efficacies. Ranked EC_50_ values of precursors correlated well especially between EDTA, EGTA, DTPA and TETA (Supplementary Fig. 6A), suggesting that we are able to capture a relative affinity relationship of metal-protein interactions in a complex proteome. Among hits with annotated metal-binding sites, zinc-binding proteins tended to display higher EC_50_ values than magnesium-binding proteins (Supplementary Fig. 6B).

Of the 297 hit proteins in our Metal-LiP experiment (Supplementary Table 1), 207 proteins (70%) were significant for at least two chelators (Fig. 2B). Most of the proteins affected by a single chelator correspond to hits upon treatment with TETA. Reassuringly, our hit list includes several known MBPs, such as IscU, CopA, MsrB and Zur, which exhibited dose responsiveness to EDTA (Fig. 2A). Also, ribosome-related and RNA-binding terms were strongly enriched in gene ontology (GO) enrichment analysis for molecular function (Fig. 2C), as expected since ribosome structure depends strongly on metal ions like magnesium^24^. Further, there was a strong enrichment for zinc-binding, indicating that zinc-binding proteins might be preferentially identified by our workflow. Consistent with this, chelator treatment resulted in the largest absolute change in zinc concentrations, of all the measured metals (Supplementary Fig. 2B). Other metal-related terms such as nickel-, manganese-, iron- and metal-binding were also enriched, in some cases for only a subset of chelators. Interestingly, “iron ion binding” and “2 iron, 2 sulfur cluster binding” were enriched specifically for the proteins changing upon addition of DiP, consistent with the selectivity of this chelator for iron^25^.

In order to more systematically assess the performance of our method in capturing known MBPs, we compiled a ground truth MBP dataset for the *E. coli* proteome based on the UniProt and QuickGO databases, consisting of around 1000 proteins previously annotated as binding metals (Fig. 2D, Supplementary Fig. 7A, Supplementary Table 2, and Methods). Further, we compiled a second set of 850 predicted MBPs using the metal-binding site predictors Metal3D, GASS-Metal and Master of Metals (MOM)^26–28^(Supplementary Table 3), with the aim of prioritising candidates for follow-up analysis and validation.

We asked whether MBPs, as defined by this ground truth dataset, were enriched in our LiP-MS hits. Roughly 50% of our hits were previously annotated MBPs, significantly higher (p-value ≤ 0.05) than the fraction of all detected proteins annotated as MBPs (30%) (Fig. 2E). Our identified MBPs comprise considerably more proteins annotated as binding to zinc and manganese, and fewer magnesium- and iron-binding proteins, with DiP and TETA again showing some differences to other chelators (Fig 2F).

To assess whether proteins with LiP hits were biased towards particular functions compared to all annotated metal-binding proteins, we performed enrichment analysis across six major functional categories (catalytic, structural, regulator, transport, electron transfer, biosynthesis) that we annotated manually based on information in UniProt (Methods). We saw no significant differences in the overall distribution of hits versus all detected MBPs (Supplementary Fig. 5E), indicating that our method samples all types of functional MBPs equally well.

Metal-LiP also identified candidate novel MBPs (i.e., those not previously annotated as binding metals and therefore not present in our ground truth dataset). We identified around 50 such proteins for most individual chelator treatments and 116 proteins in total as potentially new MBPs (Fig. 2E). Of these, 72 proteins were hits for at least two chelators (Supplementary Fig. 5F), indicating a large number of potentially novel metal-binding proteins with multiple lines of evidence. Around 25% of hit proteins without previous metal-binding annotation were predicted by at least one predictor to contain a metal-binding site, a higher fraction than for all detected unannotated proteins (Fig. 2G), further supporting that these hits represent novel metal-binding proteins. Overall, we have shown that Metal-LiP identifies MBPs across the proteome, including several proteins with no prior links to metal biology.

### Proteolytic accessibility changes upon metal chelation are near metal-binding sites

Changes detected by LiP-MS can be mapped onto protein structures, enabling identification of the affected protein region. We exploited this feature to ask whether the 981 peptides identified as Metal-LiP hits were located at known metal-binding sites or if they were distributed more diffusely across proteins. A domain-level GO enrichment analysis revealed that hit Metal-LiP peptides mapping to InterPro^29^ domains were enriched for “metal ion binding” and “zinc ion binding” domain terms (Supplementary Fig. 8A), consistent with our protein-level enrichment analysis and highlighting the binding site-level resolution of Metal-LiP.

To investigate the structural context of Metal-LiP hits at higher resolution, we calculated the 3D distances between hit peptides and annotated metal-binding sites in AlphaFold predicted structures, excluding ribosomal proteins from this analysis (Methods). In proteins with one annotated metal-binding site that contained at least one hit peptide (74 proteins across chelators), hit peptides were significantly closer to metal-binding sites than detected but unchanging peptides (Wilcoxon rank sum test p-value < 0.05), as measured by the minimum distance between the peptide and the metal-coordinating amino acids within the binding site (Fig. 3A). This supports that our chelator-based approach detects proteolytic accessibility changes in proteins due to removal of metals from their binding sites. There were no major differences in these distances for the different chelators. For proteins with one or multiple annotated metal-binding sites (111 proteins across all chelators), the median minimum distance of any hit peptide to the closest annotated metal-binding site was 4.6 Å on average over all chelators (Supplementary Fig. 8B), in comparison to 12.7 Å for non-hits, again showing that hit peptides map nearer to known metal-binding sites. These data also indicate that chelator-induced structural rearrangements do not only affect the amino acids directly involved in metal coordination (for which we would expect a distance of 0 Å), but also amino acids distal to the binding site. This is likely due to conformational changes upon metal binding as we previously showed for metabolite-binding proteins captured by LiP-MS^16^ .

**Fig. 3:**
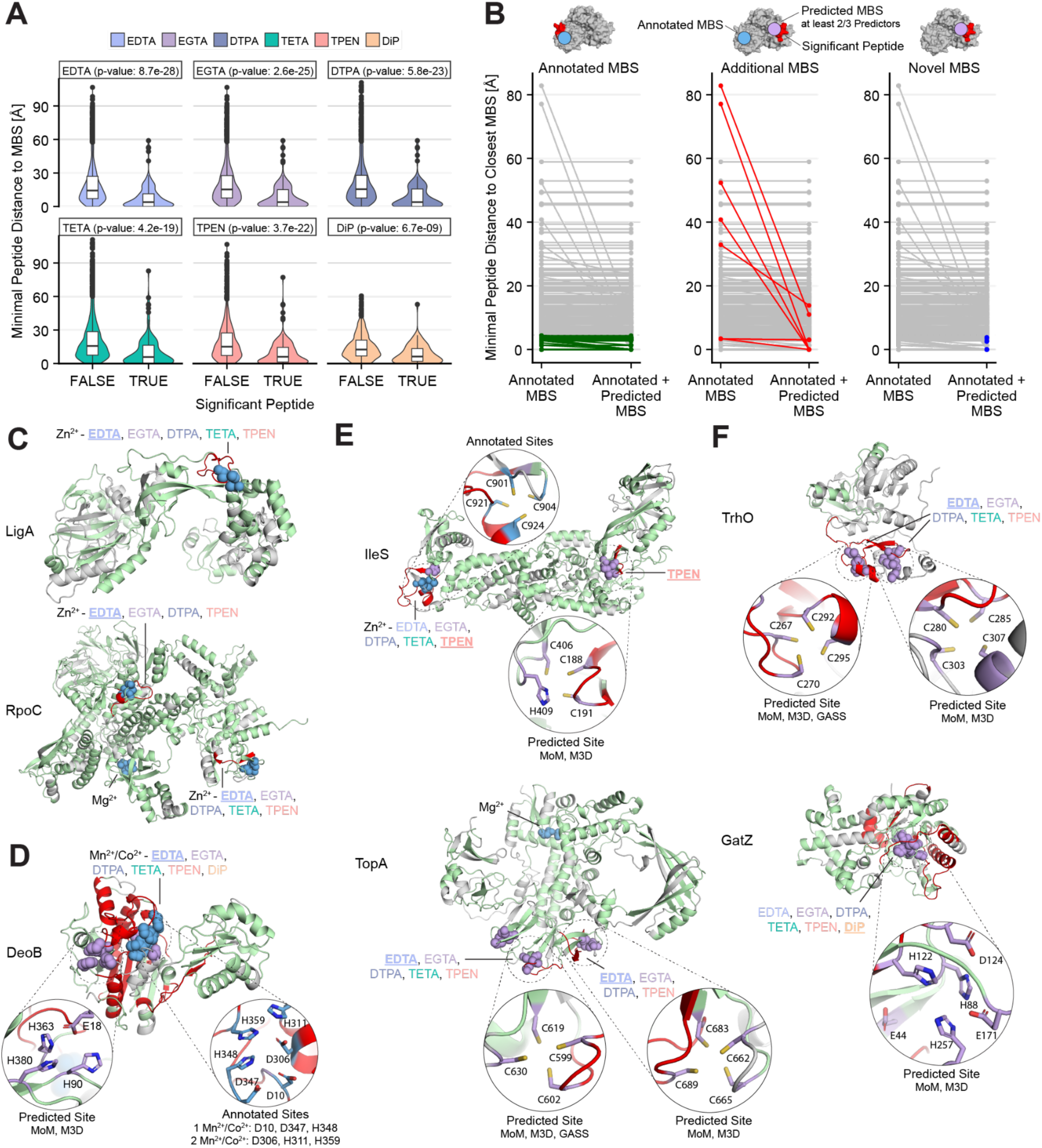
Chelator-induced structural changes are proximal to known metal-binding sites. **A** The violin plot shows the minimal distance distribution of peptides to the annotated metal-binding site of MBPs with one annotated site. For each chelator, significantly changing (TRUE) and non-changing (FALSE) peptides are plotted separately. p-values comparing the ranks of changing and non-changing peptides are shown (Wilcoxon rank sum test). **B** Each plot shows the minimal distance of hit peptides to the closest metal-binding site (MBS) for all proteins with annotated metal-binding sites and significant Metal-LiP peptides. Minimal distances to annotated MBSs are shown on the left, whereas distances to annotated or predicted (minimum 2/3 predictors) MBSs are shown on the right. If a peptide has distances for both “annotated” and “annotated + predicted” MBSs, its points are connected by a line. Coloured are peptides within 4.6 Å of an annotated MBS (left, green), peptides that have a shorter distance to a predicted than to an annotated site (middle, red), and peptides that are within 4.6 Å of only a predicted site (right, blue). **C-F** Significant Metal-LiP peptides are mapped onto AlphaFold predicted structures of the indicated proteins. Grey: Regions not detected by MS, Green: Detected, but non-changing regions, Red: Significantly changing peptides, Blue: Annotated metal-binding sites, Violet: metal-binding sites predicted by at least 2/3 predictors. We indicate which chelators have a significant peptide in the metal-binding site and underline the chelator for which the illustrated peptides have been mapped to the AlphaFold structure.

Next, we compared MBPs from the six major functional groups (catalytic, structural, regulator, transport, electron transfer, biosynthesis) in terms of their minimal peptide distances from annotated metal-binding sites, for both hit and non-hit peptides (Supplementary Fig. 8C). For proteins in the catalytic, structural and regulator groups, but not in the other functional groups, hit peptides were significantly closer to metal-binding sites than non-hit peptides, suggesting a bias for identifying known metal-binding sites depending on the functional group. However, due to the low number of proteins with regulator, transport, electron transfer, biosynthesis annotations, it is generally difficult to draw conclusions for these categories. Counterintuitively, hit-peptides of structural MBPs compared to catalytic MBPs were closer to known metal-binding sites (Mann-Whitney U p-value: 1.46e-06), suggesting a larger conformational effect upon metal removal in catalytic MBPs.

To further assess the potential of Metal-LiP for the discovery of new metal-binding sites, we then calculated the distance of hit peptides to both annotated and predicted sites, considering sites predicted by at least two of the three aforementioned methods. We defined each peptide as proximal to the binding site (<4.6 Å, as defined above for annotated sites) or distant (>4.6 Å). Of all hit proteins with annotated metal-binding sites, 71 proteins (64%) showed significant changes proximal to these sites (“Annotated MBS”, Fig. 3B). Among these proteins were the DNA Ligase (LigA), for which the changing peptide maps to the single annotated zinc-binding site for 5 out of 6 chelators (Fig. 3C). In another example, the DNA-directed RNA polymerase subunit beta’ (RpoC) contains three distinct annotated metal-binding sites (2 Zn^2+^, 1 Mg^2+^) of which two (2 Zn^2+^) are hits for at least 4 chelators, showing that our approach detects distinct metal-binding sites within the same protein (Fig. 3C). We also observed proteins where a change at the known metal-binding site was detected only in response to a single chelator, such as AlaS (EDTA) or UbiD (TPEN), showing that even though there is strong overlap between chelators, single-chelator hits are still informative (Supplementary Fig. 8D).

In some proteins, we also observed hit peptides mapping to regions distant (> 4.6 Å) from the annotated metal-binding site. For instance, DeoB, MetE and HisD showed significant peptides in the annotated site as well as the surrounding domain, indicating a broader structural effect upon metal removal (Fig. 3D, Supplementary Fig. 8E). Interestingly, in DeoB (Phosphopentomutase), the hit peptides map to an additional predicted metal-binding site in close proximity to, but distinct from, the annotated site. Our data thus provide experimental support for this predicted site and suggest that DeoB binds three rather than two metal ions (manganese or cobalt) as currently annotated. In total we identified four proteins (DeoB, TopA, HisB and IleS) where our hit peptides are closer to a predicted than an annotated metal-binding site (“Additional MBS”, Fig. 3B, E), and thus provide experimental support for such predicted sites.

In principle, a total of 490 Metal-LiP hit peptides represent candidate novel metal-binding sites, since they do not map at or near regions previously annotated to bind metals. However, many of these hits are likely to reflect allosteric or other large scale structural effects. To discover high confidence novel putative metal-binding sites, we focused on sites that were also predicted to bind metals. We used the cutoff of 4.6 Å to identify proteins without annotated metal-binding sites, where significantly changing peptides mapped at or near a predicted site (“Novel MBS”, Fig. 3B), and identified 10 such proteins. For instance, tRNA uridine(34) hydroxylase (TrhO) contains two predicted metal-binding sites, both of which were a hit with all chelators except for DiP (Fig. 3F). Similarly, the D-tagatose-1,6-bisphosphate aldolase subunit GatZ has a predicted site containing multiple possible ligands, and was a hit for all chelators (Fig. 3F).

Taken together, peptides that exhibit altered protease accessibility upon chelator treatment mapped near annotated metal-binding sites and were enriched in metal-binding domains. Importantly, we have shown that our approach identifies and characterises unannotated metal-binding sites in both previously known and newly discovered MBPs.

### Validation of potential novel MBPs

We went on to orthogonally validate a subset (n=7) of candidate novel MBPs from our Metal-LiP hits. For this, we selected proteins with predicted metal-binding sites (TrhO = two sites, GatZ = one site), with sites predicted only by Metal3D (TabA, Supplementary Fig. 8F), and with no predicted sites (FabI, AhpC, GrxB, and MsrC). HisD was also included as a positive control since it is known to bind 1 zinc ion^30^, and green fluorescent protein (GFP) as a negative control. We overexpressed N-terminally FLAG-tagged versions of each protein in *E. coli* (Supplementary Fig. 9A), purified the proteins using a FLAG bead-based purification workflow (Supplementary Fig. 9B) in combination with size exclusion chromatography (SEC) (Supplementary Fig. 9C). The metal concentrations in the purified protein preparations were then quantified by ICP-MS, and the metal occupancy was calculated as the molar ratio of metal to protein.

ICP-MS confirmed metal binding for almost all of the tested candidate MBPs. The positive control HisD showed a metal-binding site occupancy of close to 1 when we summed all the detected metals, indicating that each HisD protein molecule has one metal-binding site, as expected (Fig. 4A, Supplementary Fig. 10A). Consistent with its known specificity, HisD was found to bind mainly zinc, but also other metal ions to a lesser degree. By contrast, the negative control GFP did not bind any metal (Fig. 4A). Importantly, we observed metal binding for five of our seven selected Metal-LiP hits - GatZ, AhpC, GrxB, MsrC and TrhO, thus furthering evidence that they are novel MBPs that mainly bind zinc. We did not observe metal binding for FabI in this setup, and were unable to test TabA as it could not be purified (Fig. 4A, Supplementary Fig. 10A). We also compared different complex assembly states of GatZ (dimer and tetramer) and AhpC (dimer and decamer) that were obtained from separate SEC fractions (Supplementary Fig. 9C). While there was no large difference in total metal occupancy (calculated per monomer) for the GatZ tetramer and dimer, the tetramer bound a larger amount of copper than the dimer. Furthermore, the AhpC decamer bound considerably more total metal than the AhpC dimer (Supplementary Fig. 10B).

**Fig. 4:**
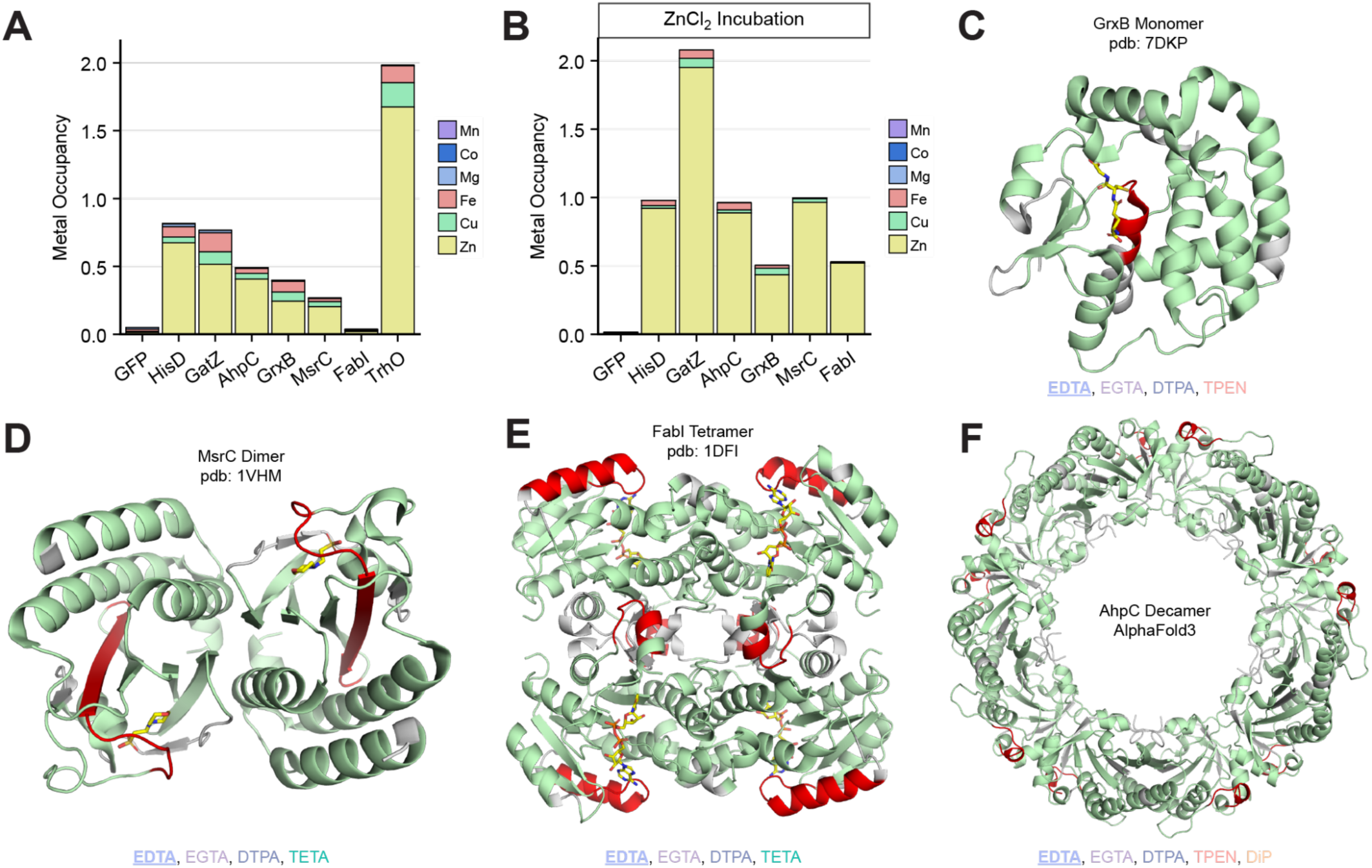
Candidate novel MBPs identified by Metal-LiP exhibit metal binding *in vivo* and *in vitro.* **A** Stacked bar plot of metal abundance normalised to protein abundance (metal occupancy) for the indicated purified proteins. **B** Metal occupancies of indicated purified proteins after incubation with a 2-fold molar excess of ZnCl_2_. All metal abundances in panels A-B were determined by ICP-MS in triplicate. For GatZ and AhpC, the molar ratios for the dimeric and decameric forms are shown, respectively. **C-F** Structures of GrxB, MsrC (dimer) and FabI (tetramer) and AlphaFold3 prediction of AhpC (decamer). Grey: Regions not detected by MS, Green: Detected, but non-changing regions, Red: Significantly changing peptides, Yellow: Ligands in active sites of proteins, glutathione (GrxB), MES (MsrC), NAD (FabI). We indicate which chelators have a significant peptide for the proteins and underline the chelator of which the labeled structure is displayed.

To quantitatively determine the maximum metal occupancy of each of the purified proteins under saturating conditions, we incubated the proteins with a 2-fold excess of metal (ZnCl_2_) and determined metal concentrations by ICP-MS after filtration. As above, positive control HisD showed an exact metal occupancy of 1, while negative control GFP did not exhibit metal binding (Fig. 4B, Supplementary Fig. 10C). For our candidate MBPs, we observed a metal occupancy of 2 for GatZ, 1 for AhpC and MsrC, and 0.5 for GrxB and FabI (Fig. 4B, Supplementary Fig. 10C). We note that TrhO was not included in this experiment due to low purification yield (Supplementary Fig. 10A). Incubation of proteins with CoCl_2_, MgCl_2_ and MnCl_2_ only showed additional binding of cobalt but not of magnesium and manganese, suggesting that these sites are indeed most specific for zinc (Supplementary Fig. 10C,D). With respect to metal occupancy for different protein assembly states in the presence of excess zinc, we found that both GatZ forms (dimer and tetramer) exhibited an occupancy of 2, whereas both AhpC forms (dimer and decamer) had an occupancy of 1 (Supplementary Fig. 10E).

We then examined metal-binding stoichiometries of our validated new MBPs in the context of their 3D structures. Both GatZ and TrhO, which have predicted metal-binding sites, were validated to bind zinc. For TrhO, two metal-binding sites are predicted (Fig. 3F), consistent with its observed metal occupancy of 2 as purified from the bacterial cell (Fig. 4A). For GatZ, however, we found two zinc ions bound per protein molecule, even though only a single site is predicted (Fig. 3F). Interestingly, this predicted site is highly similar to the dinuclear site in DeoB in terms of the type and number of coordinating amino acids (Fig. 3D), which may suggest that, like DeoB, GatZ is also bound by two zinc ions at a dinuclear site. Alternatively, GatZ may have an additional distinct metal-binding site, as evidenced by multiple GatZ Metal-LiP hits in a distant region of the protein (Fig. 3F).

For GrxB, we observed a metal occupancy of 0.5 for both *in vivo* and *in vitro* experiments (Fig. 3A, B), indicating that zinc coordination might involve a shared binding site across two subunits. Although GrxB is predominantly annotated as a monomer, we found that it elutes mostly as a dimer when analysed by size exclusion chromatography (Supplementary Fig. 9B). Mapping the significantly changing peptides to the protein structure revealed that these changes are located in the active site (Fig. 4C), although the precise amino acids that act as the coordinating ligands are not known. By contrast, for MsrC, we showed a metal occupancy of 1 (Fig. 4B), suggesting that each subunit of the dimer binds one zinc ion. The MsrC metal-binding site is likely to be located in or near the protein’s active site, due to the proximity of the Metal-LiP hit peptide to the active site (Fig. 4D).

FabI showed a metal occupancy of 0.5, again suggesting a binding site at the interface of two subunits. Hit peptides map to two distinct regions of the protein, connected by the active site, suggesting allosteric coupling of these two regions in a manner that is sensitive to the presence of a zinc ion (Fig. 4E). Since we did not observe metal bound to the purified protein, but only in the presence of excess added Zn, binding to FabI is likely to be weak. Interestingly, however, the FabI tetramer binds its substrate carrier protein ACP in the same region as the central structural change, and furthermore in a 2:4 ratio and thus in the same stoichiometry as zinc^31^. This could suggest a potential role of zinc in substrate binding. Lastly, for both decameric and dimeric AhpC we observed a metal occupancy of 1 suggesting that each protein binds one zinc ion. The observed structural change maps to the C-terminus of the protein, close to the active site that contains the reactive cysteines (Fig. 4F). While also here we do not observe a classical metal-binding site, we note that significantly changing peptides for the GrxB, MsrC, FabI and AhpC either contain or are in close proximity to cysteine residues, which could play a role in metal binding.

In summary, we have validated five novel zinc-binding proteins with metal occupancies of 0.5 to 2 and identified low-affinity or slower binding of metal to a sixth protein (FabI). While for GatZ and TrhO, we identified a clear binding site based on predictions, the binding site of the other four proteins is potentially located in protein-protein interfaces. This experiment confirms that Metal-LiP is able to identify novel metal-binding sites in proteins that were previously not annotated as MBPs.

### Functional roles of metal binding across the proteome

Next, we systematically analysed the potential functional effects of metal binding in the MBPs we identified. First, as an indication of which metal-binding events may modulate enzyme activity, we asked how many metal-binding events detected by MetalLiP structurally affect known enzyme active sites, using the active site annotations from UniProt and a distance cutoff of 10 Å (Methods). Out of the 297 hit proteins, 193 proteins are annotated as enzymes and 82 of those proteins have a known active site. Analysis of this set showed that in 38 proteins (13%) the Metal-LiP hit peptides mapped close to enzyme active sites (Supplementary Fig. 11A), suggesting that metal binding in these proteins may affect enzyme activity.

Our data show that a fraction of Metal-LiP hits mapped to protein regions distant from known or predicted metal-binding sites. While some of these hits may pinpoint novel metal-binding sites, others may reflect global structural effects arising from altered protein stability or changes in protein assembly state. We therefore next asked which of our metal-binding events could lead to differences in protein stability, and which could modulate protein complex assembly.

To identify metal-binding events that are important for protein stability, we employed a thermal protein profiling (TPP) workflow to measure the thermal stability of the *E. coli* proteome upon chelator treatment. In brief, we employed the same filtration steps as for the Metal-LiP workflow in combination with single dose EDTA treatment (20 mM), followed by a 9-temperature thermal denaturation series (37-76°C). We chose EDTA due to its broad selectivity (i.e., the EDTA hits overlap well with those of other chelators) and performance (i.e., good enrichment of known MBPs) in the Metal-LiP experiment. Using an 18-plex TMT labeling strategy in combination with high-pH fractionation and MS^3^-level acquisition to obtain melting curves for proteins (Fig. 5A), we quantified 1873 proteins consistently across three replicates (Supplementary Fig. 11B) and fitted melting curves for 1576 proteins. Of these, 81 proteins (5%) were significantly destabilised and 14 (1%) were significantly stabilised upon EDTA treatment (adjusted p-value ≤ 0.05, F-test) (Fig. 5B, Supplementary Table 4). As expected, both the destabilised and stabilised sets of proteins are enriched for MBPs (75% and 70% of proteins, respectively), as compared to 30% for all detected proteins (Fig. 5C). The metal composition of the MBPs with stability changes was similar to the Metal-LiP hits, with mainly zinc-binding proteins being destabilised and a trend towards a larger fraction of magnesium-binding proteins among the stabilised MBPs (Fig. 5D).

**Fig. 5:**
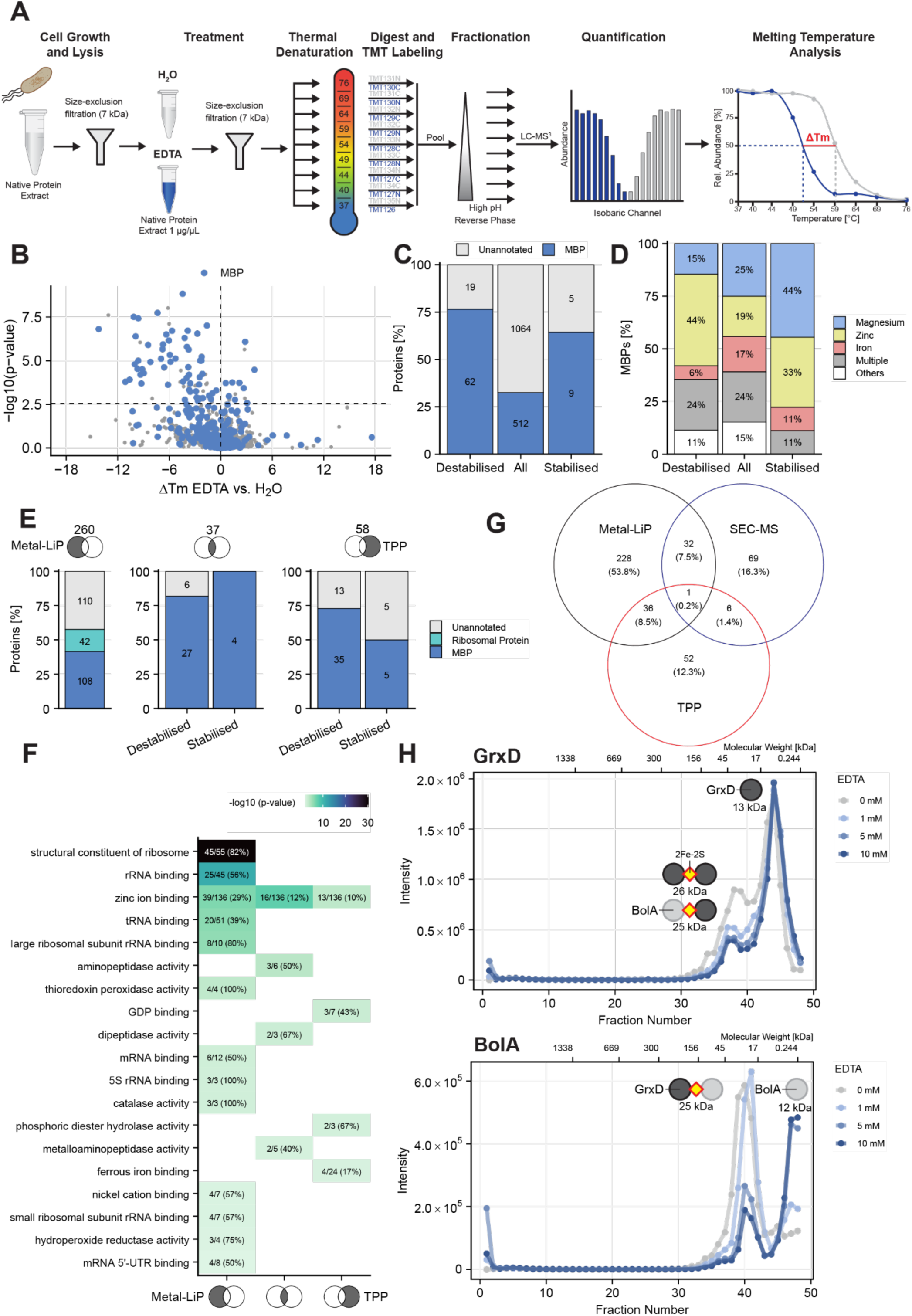
Chelator-induced metal removal affects protein stability and higher-order complex formation. **A** TPP workflow for the identification and characterisation of metal-binding proteins.Lysates were filtered before and after treatment with EDTA in line with the LiP-MS based workflow. A 9 temperature thermal denaturation series (37-76°C) was applied followed by filtration, 18-plex TMT labelling, high pH reverse phase separation and quantification by LC-MS^3^ measurement. **B** Volcano-plot of proteins with quantified melting temperatures. In blue are annotated MBPs. The significance cutoff is an adjusted p-value of 0.05. **C** Enrichment of the hit list of thermally destabilised and stabilised proteins for annotated MBPs. All indicates all detected proteins. The number of significant proteins in each category is indicated. For destabilised (p-value = 7.8e-17) and stabilised proteins (p-value = 0.02) the increase in the MBP fraction is statistically significant (p-value < 0.05) as assessed by a Fisher’s exact test. **D** The composition of the “MBP” category from panel C in terms of annotations of which metal is bound by each protein. The “Multiple” category contains proteins for which multiple metals are annotated. The “Others” category includes all metal annotations with a frequency below 5%. **E** Enrichment analysis for MBPs and ribosomal proteins of different overlap subsets between the LiP-MS and TPP experiment. The number indicates proteins in each subset. **F** Molecular function gene ontology enrichment for hit proteins from each LiP-MS and TPP overlap subset. The significance cutoff for the fisher’s exact test is a p-value of 0.01. Only terms with at least two significant proteins are plotted. **G** Overlap of significant proteins between the Metal-LiP, SEC-MS and TPP experiment. **H** SEC-MS protein abundance traces for GrxD (upper) and BolA (lower) with increasing concentrations of EDTA.

To understand the extent to which proteolytic accessibility changes observed by Metal-LiP reflect changes in protein stability, we assessed the overlap between hits in the TPP and Metal-LiP datasets (Fig. 5G). We observed that 37 proteins (about 39% of TPP hits for EDTA and 12% of LiP hits for all chelators) showed changes upon treatment with both readouts (Supplementary Table 5). This set of proteins consists largely of known (i.e., previously annotated) MBPs (83%) (Fig. 5E). Many of these proteins have catalytic metal sites, such as the binuclear and mononuclear zinc-binding sites in PepT and ArgE. However, there are also proteins with structural or regulator sites, such as Zur or bacterioferritin, suggesting that thermal stability is altered for both stabilising and active sites if the metal is removed. Further, of the 6 proteins not previously known to bind metals, 3 proteins (GatZ, HisI, AroG) are predicted to bind a metal ion by 2 predictors and one protein (YbeL) does not have a prediction since it does not meet the AlphaFold quality cutoff but contains a clear cluster of 4 cysteines that are likely to form a metal-binding site. Two proteins (WrbA and TyrB) do not have any metal-binding site predictions and would be interesting additional candidates for follow up as potentially novel MBPs. For the set of overlapping hits, there was no strong correlation between the extent of the structural changes observed in Metal-LiP and the thermal stability (ΔTm) of the protein as determined by TPP.

For the set of proteins with changes in LiP but not TPP, 42% are annotated MBPs. In addition, this set includes all significantly changing ribosomal proteins from the LiP experiment, suggesting that these proteins were mainly affected in structure but not thermal stability, upon chelator treatment. For about half of these LiP-exclusive proteins, we could measure good melting curves and can thus conclude that their thermal stability was not influenced by metal chelation, based on the sensitivity of the TPP approach. The remaining LiP-exclusive proteins were either non-melters or had non-classical melting behaviour suggestive of aggregation or multi-step thermal destabilisation.

The TPP exclusive set contains around 68% of annotated MBPs and includes most of the proteins that were stabilised upon EDTA treatment. Proteins in this set had much lower sequence coverage (median 58%) than proteins in the other two sets (median 75%) (Supplementary Fig. 11C), suggesting that this was in part why these proteins were not identified by Metal-LiP. We note that TPP requires identification of just one peptide per protein to detect a change in protein stability while LiP requires identification of the specific protein region involved in the corresponding structural change. Further, when we manually checked the 13 TPP exclusive proteins with a sequence coverage above 75%, we saw either non-classical melting behaviour or proteins with peptides just below the cutoff in LiP (data not shown). This indicates that many more TPP-exclusive hits upon metal chelation may be detectable by Metal-LiP given high enough sequence coverage.

Ribosomal terms were enriched in a GO analysis in the Metal-LiP exclusive set (Fig. 5F). Proteins of all sets were enriched for “Zinc ion binding”, with the overlapping set also enriched for peptidases, which are often metal-dependent. The “Nickel ion binding” and “ferrous iron binding” terms were enriched in the LiP set and the TPP set respectively. In summary, orthogonal analysis with TPP shows that 37 of our Metal-LiP hits (12%) and 95 candidate MBPs in total showed changes in protein stability upon metal removal, which likely represents a lower bound of this effect since several hits showed non-classical melting behavior in TPP. Furthermore, proteins with thermal stability alterations bind metals in both catalytic and structural sites, suggesting that also catalytic metals are involved in protein stabilisation.

Next, we asked whether some of the metal-binding events we detected are involved in the modulation of protein-protein interactions. To assess this, we performed size-exclusion chromatography mass spectrometry (SEC-MS) on *E. coli* lysates treated with EDTA (Supplementary Fig. 11D). Our analysis identified 2,126 *E. coli* proteins, of which 108 exhibited a significant change in assembly state upon EDTA treatment (Fig. 5G, Supplementary Table 6). These are candidate MBPs for which metal binding modulates protein complex assembly. Of these SEC-MS hits, 33 were also Metal-LiP hits, indicating that 11% of Metal-LiP hits reflect assembly changes. For example, we detected a clear decrease in the BolA-GrxD heterodimer (a known FeS cluster complex ^32,33^) and a corresponding increase in their respective monomers upon EDTA treatment (Fig. 5H). This observation suggests that the BolA-GrxD complex is sensitive to EDTA treatment and shows more broadly that Metal-LiP can detect metal-mediated assembly changes in protein complexes. Interestingly, our SEC-MS analysis also identified changes for 75 proteins (69.4%) that were not detected by Metal-LiP. For example, the homodimeric complex of the annotated MBP PhnO was destabilised by EDTA in both the SEC-MS (Supplementary Fig. 11E) and TPP (Fig. 5C) experiments. This may be explained by a shared metal-binding site at the interface of the dimeric complex, as predicted by AlphaFold3 (Supplementary Fig. 11F).

Taken together, orthogonal analyses suggest functional roles for metal binding in modulating either enzyme activity, stability or protein-protein interactions in hundreds of proteins in the *E. coli* proteome. These orthogonally-validated Metal-LiP hits will be excellent targets for future targeted studies into the functional classification of metal-binding proteins in *E. coli.* Combining results from Metal-LiP, TPP and SEC-MS we identified in total 424 known or candidate novel MBPs and mapped known or candidate novel metal-binding sites for 297 of these proteins.

## Discussion

We have developed Metal-LiP for the systematic identification of MBPs across the proteome. In brief, we combined the LiP-MS workflow with multi-dose chelator treatments and then identified proteins that showed altered protease accessibility in response to the chelator. This served as a readout for potential changes in metal occupancy upon chelator treatment and therefore for proteins that are likely to bind metals under the conditions tested. In the *E. coli* proteome, Metal-LiP preferentially identified known and predicted MBPs, but also 116 novel candidate MBPs, and precisely identified metal-binding sites.

Current systematic understanding of MBPs across proteomes is mainly based on computational analyses, with few proteome-wide experimental datasets available. Metal-LiP offers substantial advantages for profiling this important set of proteins since it identifies them without relying on known binding motifs, and avoids the cysteine bias inherent in methods such as ZnCPT^9^. Indeed, in orthogonal validation studies of 6 of the novel MBPs identified by Metal-LiP, we validated 5 out of 6 as bonafide MBPs. FabI could only be validated after incubation with metals and has therefore potentially low-affinity or slower kinetics. Only 2 of these proteins had previously predicted metal binding sites, underlying the substantial potential of Metal-LiP for MBP discovery. Notably, Metal-LiP offers peptide-level resolution and therefore can precisely localize metal-binding sites. Our systematic 3D structural analysis of Metal-LiP hit proteins with previously annotated metal binding sites showed that chelator-responsive peptides are on average 4.6 Å away from known metal-binding sites, indicating a close spatial relationship. Thus Metal-LiP can capture regions influenced by metal coordination, which includes regions that are near, rather than exclusively at, the metal binding site.

Interestingly, the six chelators we used had broadly similar metal-depletion effects. This lack of selectivity may be because the chelators are at drastically higher concentrations than metals in our experiment. Chelators should display metal selectivity when present in equimolar or substoichiometric amounts relative to metals, as selectivity is determined by the equilibrium constants for different metals. However, when chelators are in large excess, it is possible that even metals with lower affinity will be bound. We conclude that, since the 6 tested chelators were present in large excess, this led to a similar degree of metal removal from proteins in *E. coli* lysates. Furthermore, this suggests that all the metal that is accessible to chelation is removed from proteins in our setup. We note that chelation removes a relatively small proportion of the bound metal, providing support for previous studies that have suggested that there is a limited chelator-accessible pool of metal in MBPs^34^.

Metal co-ordination is likely to be a functionally important feature of most MBPs. This is particularly true for the subset of Metal-LiP hits that are enzymes, and where metal-binding occurs at or near the enzyme active site. Nevertheless, chelator-responsive peptides are not expected to map only to the metal-binding site since, as we have previously shown for LiP-MS, Metal-LiP can identify several types of molecular events. Indeed, integration with orthogonal datasets showed that the approach also identified proteins with altered thermal stability and protein complexes with changed assembly states upon metal chelation.

Thermal proteome profiling showed changes in stability in 6% of all measured proteins upon EDTA treatment, 38% of which were also identified as showing altered protease accessibility in Metal-LiP. Most proteins showed destabilisation upon metal removal, consistent with the stabilising role of metal ions in protein structure. We also note that, manual inspection of TPP melting curves revealed non-standard melting behaviours upon EDTA treatment, suggesting additional complexity in their response to metal removal that will require further studies. In addition, there were shifts in protein complex assembly states upon metal chelation, indicating that metal binding both stabilises and destabilises protein complexes. Notably, of the roughly 100 proteins that changed their assembly state based on SEC-MS after EDTA treatment, one third were also detected by Metal-LiP. These experiments emphasize the complementarity of Metal-LiP to these orthogonal approaches, but importantly also provide insight into proteome-wide changes upon metal chelation. Notably, our data identify metal binding as a potentially widespread and underexplored mechanism for regulating protein-protein interactions, with exciting implications for selectively tuning complex assembly.

Our approach has some limitations. First, membrane proteins are underrepresented in the dataset and the method requires high sequence coverage thus making it most likely to identify chelation-dependent changes in relatively abundant proteins. It may also miss genuine MBPs with inherently high proteolytic resistance or without significant conformational changes upon metal removal. Finally, weakly bound metals such as magnesium may be already removed during preparatory filtration steps and their binding therefore difficult to detect with this chelator-based approach.

In summary, we have developed the Metal-LiP approach to identify MBPs proteome-wide and applied it to discover novel bacterial MBPs and to advance our understanding of the structural consequences of metal removal. Understanding metal-dependent stabilisation and catalytic roles could inform the design of proteins with enhanced stability or new enzymatic functions. Further, our approach should be applicable to any proteome and to various physiological or environmental conditions, thus offering the potential to broadly study the regulation of MBPs and metal-dependent cellular processes.

## Methods

### Strains and growth conditions

#### Escherichia coli

The *E. coli* K-12 strain BW25113^35^ was grown in a modified M9 medium^36^ with adjusted metal concentrations (see detailed description below and detailed composition in Supplementary Table 7). A starter culture was inoculated in 50 mL of modified M9 medium from a single colony on an LB agar plate and grown overnight at 37°C (OD_600_ ∼ 2.5). The culture was spun down at 2500 g and washed once with modified M9 medium. 1000 mL cultures were inoculated in 3L Fernbach-Design plastic flasks (Corning) at an OD_600_ of 0.05 and grown to an OD_600_ of 0.8 at 37°C. The exponential logarithmic growth rate should be above 0.6 h^-1^. Cells were collected at 2500 g, for 10 minutes at 4°C. They were washed twice with cold magnesium-free LiP buffer (detailed composition in Supplementary Table 7). In the final wash step a volume corresponding to an original culture volume of 33 mL was transferred to each 1.5 mL screw cap tube. After removal of the supernatant, pellets were snap-frozen in liquid nitrogen and stored at −80°C.

### Preparation of modified M9 medium

We observed that red iron precipitates form in M9 medium when prepared according to Kochanowski et al.^36^. We confirmed this using ICP-MS, showing that not only iron, but also copper and zinc concentrations were lower in the medium supernatant or medium filtrate (Supplementary Fig. 1A). Precipitating metal ions may cause problems during cell harvesting, as large amounts of metals co-pellet and potentially interfere with downstream assays. Therefore, we tested 10 µM ferrichrome and 1 mM citrate, as previously described by Hartmann and Braun^38^, to solubilise precipitation-prone metals. ICP-MS confirmed that, in both media, metals did not precipitate anymore (Supplementary Fig. 1A). Despite only adding 4 µM and 10 µM of FeCl_3_ to the citrate and ferrichrome media respectively, the final soluble concentration was higher than in the original M9 medium with 60 µM FeCl_3_ (∼1 µM soluble). When comparing the effect on growth as a function of the FeCl_3_ concentration in the three media, the citrate medium achieved the highest density, followed by the ferrichrome and original M9 media. The maximum growth rate was highest in the citrate medium (µ: ∼0.7), followed by the original M9 medium (µ: ∼0.6) and the ferrichrome medium (µ: ∼0.55) (Supplementary Fig. 1B). We therefore decided to use 1 mM citrate for our new modified M9 medium due to its superior growth characteristics. Higher citrate concentrations strongly impact the pH of the medium, the maximum density, and the maximum growth rate (IC_50_: 25.5 mM) and therefore could not be used (Supplementary Fig. 1C). We used a citrate assay kit (Sigma, MAK057) to determine the citrate concentration in the supernatant of the medium during growth to an OD_600_ of 0.8 and confirmed that citrate is not markedly consumed by the cells at this growth stage (Supplementary Fig. 1D).

We also noticed that the concentration of trace elements such as copper are very high^39^ in the original M9 medium, likely creating a less physiological environment with increased oxidative stress and resultant upregulation of copper efflux systems. To get a reference for more physiological metal concentrations, we determined metal concentrations in Luria-Bertani (Miller) medium by ICP-MS (Supplementary Fig. 1E). The metal concentrations are consistent with published measurements^19,40,41^ and indeed the trace metal concentrations were considerably lower in LB medium as compared to the original M9 medium by Kochanowski et al.^36^. For our new modified M9 medium, we therefore also adjusted all divalent metal concentrations to the levels in LB medium. In brief, calcium is the only metal with the same concentration and zinc (1.6-fold) is the only metal with an increased concentration, while magnesium (4-fold), iron (12-fold), manganese (23.6-fold), copper (12.6-fold) and cobalt (50.6-fold) have a decreased concentration in the modified M9 medium. The detailed composition of the new modified M9 medium can be found in Supplementary Table 8 along with preparation instructions. All solutions were filter sterilised using Millex 0.2 µm PTFE filters (Merck) in combination with seal free PP/PE syringes (Merck) or the Corning® bottle-top vacuum filter system. All media components that were not trace metal grade were filtered through a Chelex-100 column (Bio-Rad) in order to remove any contaminating divalent cations.

Finally, the original M9 medium had a maximum growth rate of 0.63 while the maximum growth rate of the modified M9 was 0.7 in addition to a shorter lag time (Supplementary Fig. 1F). As expected, a direct comparison of the proteomes (Supplementary Fig. 1G) shows a lower abundance of copper detoxification systems in the modified M9 medium, likely due to lower intracellular Cu^+^ and Cu^2+^ concentrations^42^. In addition, Fe-S-cluster assembly systems Isc and Suf, which are negatively regulated by intracellular Fe^2+43^, also have a decreased abundance. We also note that chelation dependent iron import systems are predominantly downregulated, likely due to the readily available Fe^3+^-citrate. In contrast, ferrous iron import system proteins^44,45^, such as feoA, feoC, efeB, efeO are slightly upregulated. At least the Efe system, which is non-functional in the K-12 strain because of a frameshift in efeU, has been reported to be induced by low pH^45^. Other proteins involved in a response to an acidic pH were also upregulated, likely in response to the citrate in the medium. In general many of the proteins increasing in abundance were iron-binding proteins that are transcriptionally regulated by e.g. Fur, indicating a higher intracellular Fe^2+^ concentration.

#### *E. coli* 96-well growth assay

A starter culture was inoculated from a single colony on an LB agar plate in LB medium and grown overnight at 37°C. Another pre-culture in modified M9 medium was inoculated and grown overnight at 30°C to not overcrowd the culture. The cells were once washed in LiP buffer and collected in pellets. We resuspended the pellets in the assay media so that the final OD_600_ is 0.05. Each replicate of 200 µL was transferred to a clear flat-bottom 96-well plate (Thermo, 136101). The four corner wells were used as negative controls and each well of the plate was either filled with a culture or water to minimise evaporation. The clear plastic lid was taped to the plate in two opposite positions to reduce rubbing during incubation while allowing gas exchange. The plate was incubated at 37°C with constant double orbital shaking at 1000 rpm in a CLARIOstar® Plus (BMG LABTECH). The OD_600_ was determined every 10 minutes without “Well Scan” for 24 hours or until stationary phases were reached.

### Native cell lysate preparation

Frozen pellets were thawed on ice. Approximately 1-2 pellet volumes of acid-washed ≤106 μm glass beads (Sigma, G4649) were added to the tube, as well as 500 µL of magnesium-free LiP buffer. The crude native protein lysate was extracted by bead beating at a speed of 6 m/s, cycle time of 30 seconds, 2 cycles and 300 seconds pause at 4°C using a FastPrep-24™ 5G bead beating grinder and lysis system (MP Biomedicals, 116005500). The crude lysate was cleared by centrifugation at 18000 g for 10 minutes at 4°C and the supernatant was transferred to a new tube.

### Chelator Stock Solution Preparation

We used the water-soluble chelators Ethylenediaminetetraacetic acid (EDTA, Sigma, #431788), Ethylene glycol-bis(2-aminoethylether)-N,N,N′,N′-tetraacetic acid (EGTA, Sigma, #E0396), Diethylenetriaminepentaacetic acid (DTPA, Sigma, #32319) and Trientine dihydrochloride (TETA, USP, #1683504) and the DMSO-soluble chelators N,N,N′,N′-Tetrakis(2-pyridylmethyl)ethylenediamine (TPEN, Sigma, #P4413) and 2,2′-Bipyridyl (DiP, Sigma, #D216305) (Fig. 1A). The stock solutions of all water soluble chelators were prepared to be 26x of the final concentration, while DMSO soluble chelators were prepared to be 51x concentrated to limit the amount of DMSO added to the sample. The maximum concentrations of chelators for dose-response curves were generally dictated by their stock solubility limits. We prepared a 520 mM stock of EDTA, EGTA and DTPA, which was solubilised by addition of KOH, to a final pH between 7.4 and 8.0. TETA was soluble at a 520 mM concentration without addition of base. The stock concentrations of DiP and TPEN were 510 mM and 51 mM, respectively. We confirmed that addition of those stocks in their designated treatment concentrations and volumes does not affect the pH of the buffered sample.

### Limited proteolysis

For samples subjected to chelator treatments, cell lysates were filtered through Zeba™ Spin Desalting Columns, 7K MWCO (Thermo Fisher Scientific, #89894) in order to remove most of the free metals from the lysate. To remove any metal impurities present in the filters, they were first equilibrated by two washes with 100 mM EDTA, five washes with H_2_O and three washes with magnesium-free LiP buffer prior to use. We confirmed by ICP-MS and LC-MS that contaminating metal concentrations were reduced and that EDTA was efficiently removed from the column by washing (data not shown). After filtration protein concentrations were determined using the Pierce™ Dilution-Free™ Rapid Gold BCA Protein Assay (Thermo Fisher Scientific, #A55860) for quick processing and subsequently adjusted to 1 mg ml^-1^.

The next steps were performed on a Hamilton Liquid handling platform using the CO-RE 96 head, allowing parallel and precisely time controlled processing of up to 96 samples. For the treatment the chelator was added in a 1:26 or 1:51 v/v ratio to the lysate (depending on solvent and maximum concentration, e.g. 2 µL to 100 µL lysate) and then incubated for 10 minutes at 4°C. The chelator was removed from the sample in a second filtration step with Zeba™ Spin Desalting Plates, 7K MWCO (Thermo Fisher Scientific, #89808). The filtration was performed with 100 µL allowing us to split the sample for processing in the LiP workflow as well as for a “tryptic digest only” control. For LiP samples, 5 µL of 0.1 µg/µL (1:100) Proteinase K from *Tritirachium album* (Sigma, P2308) was added to 50 µL of treated lysate and incubated for 1 before the reaction was stopped by heating to 99°C for 5 minutes and by the addition of sodium deoxycholate (DOC) to a final concentration of 5%. For samples subjected to the standard LiP-MS workflow (e.g. MgCl_2_ treatment) both filtration steps were skipped and incubation with the treatment was performed for 5 minutes at 25°C. Otherwise all other steps are identical.

### Trypsin digestion

We reduced denatured proteins by addition of 5 mM tris(2-carboxyethyl)phosphine (TCEP) for 40 minutes at 37°C. Alkylation was subsequently performed with 40 mM iodoacetamide (IAA) and incubation for 30 minutes in the dark at room temperature. Samples were then diluted with 100 mM ammonium bicarbonate to a final DOC concentration of 1% (1:5 dilution) and digested with trypsin (Promega, #V5113) and LysC (Wako Chemicals, #125-05061) (1:100 enzyme:substrate ratio) overnight at 37°C. For samples previously exposed to chelators, we added 3 mM CaCl_2_ to the digestion reaction to quench any leftover chelator and to provide extra calcium ions to trypsin. The digestion was stopped by addition of formic acid to a final concentration of 2% (pH < 3). The resultant DOC precipitate was then removed by filtration through an AcroPrep™ Advance, 0.2 µm PTFE membrane, 96-well filter plate (PALL, #8582) by centrifugation at 1000 g for 2 minutes.

### Sample desalting

Peptides were desalted using BioPureSPF EasyFlo PROTO 300 C18 Mini-96-Well (The Nest Group, #HNFR S18V) plates with a capacity of 7-70 µg according to the manufacturer’s instructions. After washing with 0.1% formic acid (FA), peptides were eluted in 80% acetonitrile (ACN) and 0.1% FA and dried in a vacuum centrifuge. This was followed by resuspension in 25 µL of 0.1% FA. For most samples we added 1:20 iRT peptides (Biognosys) into the resuspension solution for quality control of MS runs.

### Peptide ion exchange

For the proteinase K activity assay, chelators were removed from the samples prior to desalting to minimise the risk of ion suppression. In this workflow, our standard filtering approach is not possible because peptides would be removed from the samples as well as the chelator. After the addition of DOC, the samples were diluted with 1700 µL of water to reduce salt concentrations below 5 mM and then acidified with 5 µL of 50% phosphoric acid. The samples were filtered through a 0.2 µm filter plate to remove resultant DOC precipitates. The filtrates were then cleaned up using a HIL-SCX 96 well plate (The Nest Group, #HNS HIL-SCX). For the HIL-SCX cleaning, the wells were consecutively pre-washed via vacuum/negative pressure with 400 µL of methanol, 400 µL of water, twice with 400 µL of the high salt-containing Buffer D (5 mM KH_2_PO_4_, 25% acetonitrile, 300 mM KCl adjusted to pH 2.8 with phosphoric acid) - the last wash being done at by gravity ideally - and finally three times with 400 µL of the low salt-containing Buffer C (5 mM KH_2_PO_4_, 25% acetonitrile adjusted to pH 2.8 with phosphoric acid). The peptides were then loaded onto the HIL-SCX resin, washed twice with 400 µL of Buffer C and finally eluted in a clean plate with 400 µL of Buffer D. The HIL-SCX peptide eluates were vacuum dried to remove acetonitrile and then resuspended in 200 µL of Buffer A (0.1% formic acid) and further desalted as described above.

### Testing condition-specific effects on proteinase K activity

We prepared short peptides as previously described^46^ in order to test for condition specific effects in a proteinase K activity assay. Proteinase K binds calcium for stability^47^ making a negative effect of chelators on protease activity possible. Peptides were resuspended to a concentration of 0.5 µg/µL in 50 µL magnesium-free LiP buffer. The LiP step was performed with metals and chelators as described above with 1:50 proteinase K. The already peptide containing samples were not reduced, alkylated or digested another time, but simply desalted after removal of sodium deoxycholate as described above. For the chelator treated samples we added an ion exchange step before the C18 peptide desalting step as described above in order to remove chelators from the samples as much as possible. We do this because we had observed ion suppression for some chelators before. As described below, this step was only partially successful as we still observed ion suppression in the TPEN condition (Supplementary Fig. 2D). For the analysis we used the R package protti^48^. We filtered to only retain peptides of a maximum length of 10 amino acids to exclude any structure specific effects on peptide digestion. In addition, we only kept peptides for the analysis for which either all or no replicates were present for any given condition (no missing at random cases). Further, we excluded non-tryptic peptides. We identify significantly changing peptides by differential abundance analysis between the control (H_2_O treatment) and each metal or chelator treatment using a moderated t-test followed by Benjamini-Hochberg correction. We consider peptides to be significantly changing if they have an adjusted p-value ≤ 0.05 and a |log_2_(fold change)| ≥ 1. Since both fully and semi-tryptic peptides can be substrates and products of proteinase K, we used an additional control without proteinase K to broadly define which peptides are the substrates (increase in - PK condition) and which ones are the products (decrease in -PK condition). Any peptides that do not change in this condition but in other conditions are labelled as “unknown” since we do not know for sure if they report on an activity increase or decrease.

We could confirm that our inhibitory controls (pH 5 and pH 6) indeed decreased proteinase K activity, while pH 8 increased it (seen by a decrease in substrate peptides) as expected (Supplementary Fig. 2C and 3A). There was a clear dose dependent effect of all metals on proteinase K activity (Supplementary Fig. 3A). Interestingly, the strength of inhibition for the different metals follows the Irving-Williams series, likely indicating that stronger binders, such as copper, have a stronger inhibitory effect due to the stronger interacting with the protein. None of the metals inhibited proteinase K at a concentration of 1 µM, likely due to the fact that this concentration was lower than the proteinase K concentration in the reaction (3.45 µM). This could indicate that the metals are not denaturing proteinase K, but rather inhibit it by binding, however, this is hard to judge for the short 1 minute duration of the assay.

For chelator treatments, we saw a less strong effect on proteinase K activity than with metals with only EDTA and TETA having an effect at their highest concentrations (Supplementary Fig. 2C). TPEN caused ion suppression during MS acquisition at 1 mM and to a small extent at 100 µM (Supplementary Fig. 2D).

### Thermal proteome profiling

The TPP protocol was adapted from Pepelnjak et al.^46^. *E. coli* lysates (1 µg/µL) were treated for 10 minutes on 4°C with H_2_O (control) or a final concentration of 20 mM EDTA (1:26 dilution) and subsequently filtered through Zeba™ Spin Desalting Columns, 7K MWCO (Thermo Fisher Scientific, #89894). Samples were aliquoted into a 96-well plate (65 µL, 3 replicates) and subjected to a temperature gradient of 9 temperatures (37°C, 40.5°C, 44.4°C, 49.3°C, 54.1°C, 58.9°C, 63.8°C, 68.6°C, 76°C) for 5 minutes in a Biometra Tadvanced 96-well PCR machine. Subsequently, samples were filtered through a 0.2 µm PVDF membrane filter (Corning® FiltrEX™ 96 well, #CLS3505) by centrifugation at 800 g for 5 minutes. 50 µL of flowthrough was transferred and mixed with an equal amount of 10% sodium deoxycholate, followed by reduction, alkylation as described above. Digestion and TMT labelling was performed as described below.

### TMT labelling

TPP samples were digested with trypsin and LysC (1:100 enzyme:substrate ratio) in 100 mM HEPES pH 8.2 at 37°C overnight. Peptides were desalted as described above and dried in a vacuum centrifuge. We used TMTpro 18-plex (Thermo Fisher Scientific, #A52045, batch1-16 Lot Y1373062, batch 17-18 Lot YH378596) to label the 9 temperatures (increasing) of the treatment followed by the 9 temperatures (decreasing) of the control in increasing order of isotope weights. Each replicate consisted of a separate plex using the same labelling strategy. We resuspended the dried peptides in 100 mM HEPES pH 8.5 to a final concentration of 0.5 µg/µL and labelled 20 µg of peptides with 50 µg of TMT labels using the standard protocol. Each plex was pooled after quenching, containing equal amounts of every sample. After acidification, labelled peptides were briefly dried in a vacuum centrifuge to remove ACN, followed by high-pH fractionation as described below.

### High-pH reversed-phase fractionation

To improve protein identification the pooled TMT samples were fractionated into 8 fractions using the Pierce™ High pH Reversed-Phase Peptide Fractionation Kit (Thermo, 84868). We used the TMT workflow according to the manufacturer’s instructions but made further adjustments to the acetonitrile concentrations in the fractions to get a better distribution of peptides. 12.5% (Fraction 1), 15% (Fraction 2), 17.5% (Fraction 3), 20% (Fraction 4), 22.5% (Fraction 5), 25% (Fraction 6), 30% (Fraction 7), 50% (Fraction 8). After fractionation samples were dried in a SpeedVac concentrator and resuspended in 20 µL of 0.1% formic acid. 1 µL of each fraction was injected into the MS for the measurement.

#### Size exclusion chromatography (SEC) of *E. coli* lysate

*E. coli* lysate was first filtered with Costar® Spin-X® tube filters (Merck, # CLS8161), for 12000 g at 4°C, for 2 min and adjusted to a final concentration of 5 µg/µL with LiP buffer containing 1 mM MgCl_2_. 500 µg of the proteome extract were then treated with 0 mM, 1 mM, 5 mM and 10 mM EDTA for 15 min at 4°C and subsequently fractionated with a BioSep 5 µm SEC-s4000 500 Å, LC Column 300 x 7.8 mm. Each SEC run was carried out at 4°C, in LiP buffer, with a flow rate of 0.5 mL/min. 48 fractions were collected in a retention time window from 10.25 min to 25 min. The molecular weight calibration curve for SEC was obtained by running a protein standard mix (Phenomenex, #AL0-3042).

The collected fractions were then subjected to trypsin digestion using the FASAP protocol using 96-well plate MWCO filters (AcroPep Advance Filter Plates for Ultrafiltration 1mL Omega 10K MWCO, Pall Corporation, USA), as described in^49^. Briefly, each fraction was loaded onto the filters and concentrated through centrifugation to completely remove the LiP buffer. Subsequently, proteins were denatured and reduced via a 30 min incubation at 37°C with 5 mM of TCEP in 8 M urea/ 20 mM ammonium bicarbonate (AMBIC) (pH 8.8). Cysteine residues were then alkylated by adding 50 mM IAA in 20 mM AMBIC and incubating for 1h at room temperature in the dark. The reaction was stopped by removing the added buffers by centrifugation, and washing three times with 20 mM in AMBIC.

Proteins were then digested at 37°C for 16 h adding 1 μg of trypsin (Promega, #V5113) and 0.3 μg lysC (Wako Chemicals, #125-05061). The resulting peptides were collected on a 96-well PCR-plate (Hard-Shell® 96-Well PCR Plates, Biorad, #HSP9601) by centrifugation, followed by a filter wash with 100 μl of ddH_2_O. Peptides were then dried in Speedvac and resuspended with 15 μL of Buffer A (0.1% formic acid).

#### ICP-MS sample preparation, acquisition and data analysis

To each sample (∼200-500 µL) we added 200 µL of 67% nitric acid (VWR, 83879) and 100 µL of 30% hydrogen peroxide (Supelco, 16911). Samples were incubated at 40°C in a sonication bath for 2 hours. If there were still insoluble precipitates we incubated the sample further overnight. For analysis samples were diluted to a final volume of 4 mL with 2% nitric acid.

The ICP-MS measurements were performed with an Agilent 8800 Triple quad ICP-MS, equipped with a standard x-lens setting, nickel cones and an Agilent MicroMist quartz nebulizer. The feed was 0.1 ml/min and the RF power 1550 W. The tune settings were based on the Agilent General Purpose method and only slightly modified by an autotune procedure using an Agilent 10 ppb tuning solution containing Li, Y, Ce and Tl. The values are reported as the average of 60 sweeps x 3 replicates. Depending on the element they were measured with oxygen, helium and/or hydrogen as collision/reaction gases. All solutions were prepared with 60% HNO_3_ (Merck, 1.01518) and 18.2 MΩ Millipore water. The certified ICP multi-element standard IV (Merck, 1.11355) was used for the calibration that consisted of at least eight calibration points. The calibration standards were freshly prepared before a measurement series. The device performance was monitored with an internal standard (Yttrium, Merck, 1.70368) in the wash solution that was injected after nine samples. The data was processed with Agilent MassHunter^TM^ and the concentrations are stated in ppb. The calibration curves were of linear fashion and an R value of above 0.99 was achieved for each element, most of the time it was above 0.999. Data was further processed using an R shiny app (https://github.com/jpquast/ICP-MS-data-explorer) in order to select the most consistent gas modes and isotopes for each metal, to correct for sample dilution, convert to molar concentrations and to normalise measurements to input material weight.

### Chelator LC-MS sample preparation, acquisition and data analysis

In order to determine absolute chelator concentrations we prepared chelator standard curves in lysate with LiP buffer. All samples were measured using an Agilent 6546 mass spectrometer, coupled with an Agilent Infinity II chromatographic system. We used an Atlantis Premier BEH C18 AX 1.5 μm, 2.1×30 mm column. The solvent system was 2.5 μM medronic acid in water (buffer A), 0.1 % formic acid in methanol (buffer B) and 50 mM ammonium formate and 0.1% formic acid in water (buffer C). The column compartment was kept at 40 °C and the flow rate was 1.2 mL/min. The gradient was: 0-0.4 min, 100% A; 0.4-0.8 min, 40% A and 60% C; 0.8-1.5 min, 5% A, 40% B and 55% C; 1.5-2.05 min, 5% A, 40% B and 55% C; 2.05-2.5 min, 100 % B; 2.5-2.9, 100 % B; 2.9-3 min, 100% A, 3-4 min, 100% A. The injection volume was 2 μL. The data was processed with Agilent MassHunter^TM^. Data analysis was performed in Skyline v.23.1.0.455. We transformed intensities to the log2 space and concentrations to the log10 space and fitted a line to the linear portion of the standard curves. Concentrations were estimated from the linear model for each chelator.

### Proteomics LC-MS data acquisition

We performed the LC-MS analysis for full proteome LiP (120 min) and proteinase K activity assays (60 min) on an Orbitrap Exploris 480 mass spectrometer (Thermo Fisher Scientific), TMT analysis on an Orbitrap Eclipse Tribrid mass spectrometer (Thermo Fisher Scientific) and SEC-MS on an Orbitrap Astral mass spectrometer (Thermo Fisher Scientific).

For the Exploris 480 and Eclipse peptides were chromatographically separated using a 40 cm × 0.75 mm inner diameter column (CoAnn ICT36007515F-50-5) with a pulled emitter packed with 3 μm C18 beads (Dr. Maisch, Reprosil-Pur 120). For the Astral a 20 cm x 0.75 mm column was used instead. The LC settings for each instrument and method can be found in the table below. The injection amount for each sample was usually in the range of 1 μg.

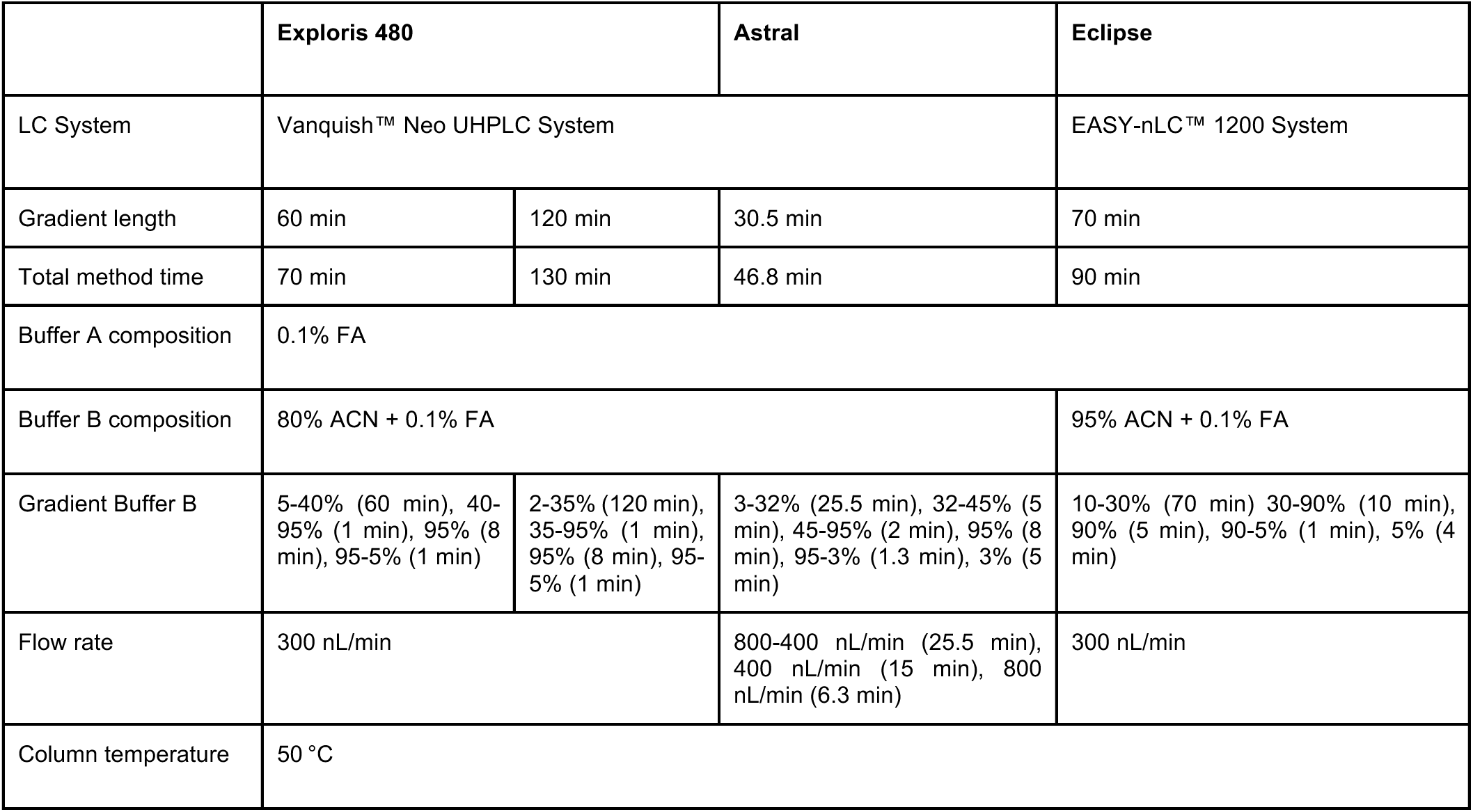

### Data-independent acquisition (DIA)

DIA measurements were acquired on the Exploris 480 and Astral mass spectrometer according to the settings below.

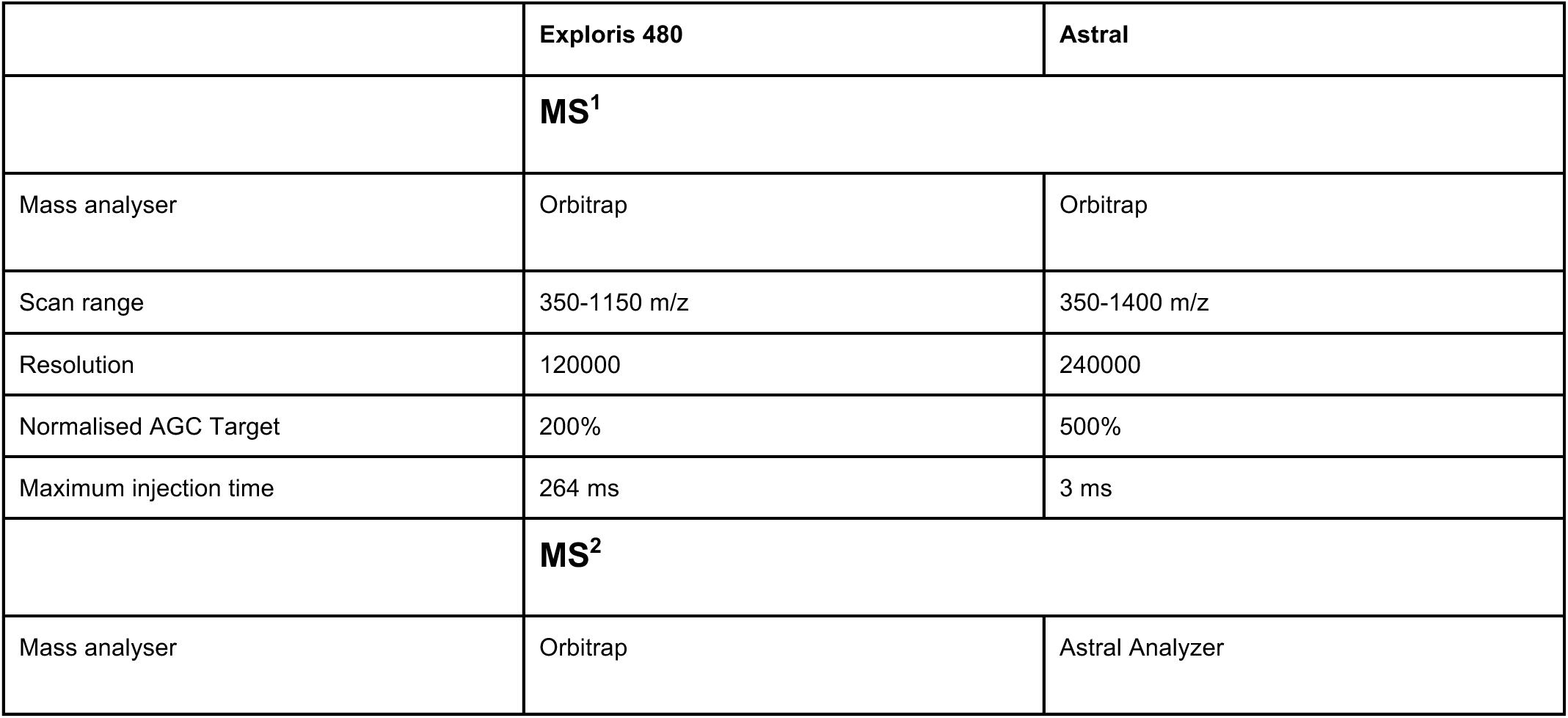
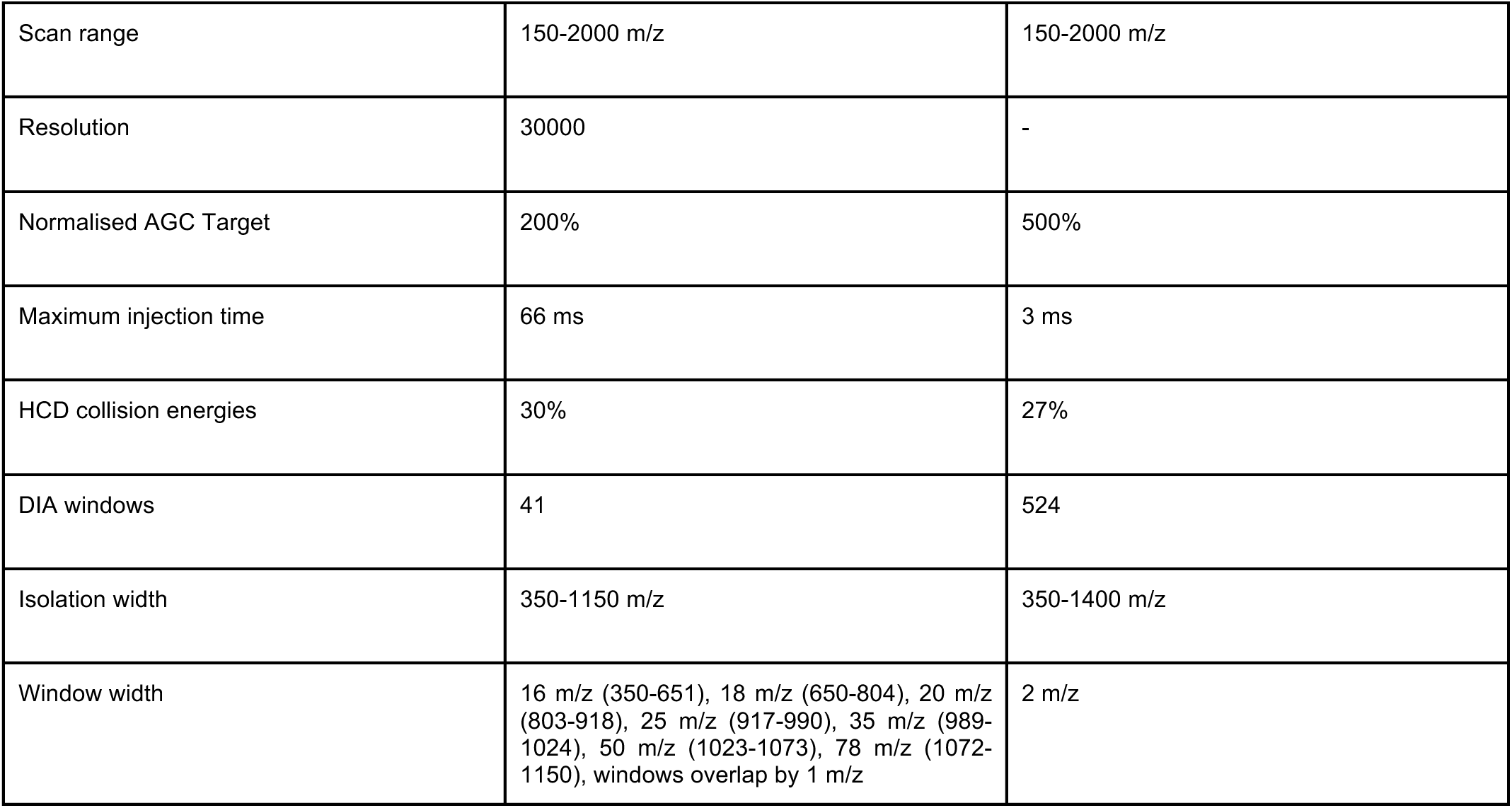

Data-independent acquisition (DIA) DIA measurements were acquired on the Exploris 480 and Astral mass spectrometer according to the settings below.
|  | Exploris 480 | Astral |
| --- | --- | --- |
|  | **MS<sup>1</sup>** |  |
| Mass analyser | Orbitrap | Orbitrap |
| Scan range | 350-1150 m/z | 350-1400 m/z |
| Resolution | 120000 | 240000 |
| Normalised AGC Target | 200% | 500% |
| Maximum injection time | 264 ms | 3 ms |
|  | **MS<sup>2</sup>** |  |
| Mass analyser | Orbitrap | Astral Analyzer |
| Scan range | 150-2000 m/z | 150-2000 m/z |
| Resolution | 30000 | - |
| Normalised AGC Target | 200% | 500% |
| Maximum injection time | 66 ms | 3 ms |
| HCD collision energies | 30% | 27% |
| DIA windows | 41 | 524 |
| Isolation width | 350-1150 m/z | 350-1400 m/z |
| Window width | 16 m/z (350-651), 18 m/z (650-804), 20 m/z (803-918), 25 m/z (917-990), 35 m/z (989-1024), 50 m/z (1023-1073), 78 m/z (1072-1150), windows overlap by 1 m/z | 2 m/z |

### Data-dependent acquisition (DDA) for Tandem mass tag (TMT)

TMT samples were analysed on the Eclipse mass spectrometer using the above mentioned chromatographic settings. We used a DDA MS3-based method with real time search and a 3 second cycle time adapted from Leutert et al.^50^. MS1 data was collected in the Orbitrap at a resolution of 120000 in a scan range of 400-1700 m/z, using an automatic maximum injection time and standard AGC target, with included charge states 2-6 and a dynamic exclusion of 45 seconds. The most intense precursors were selected using an isolation window of 0.7 Da and fragmented with CID fragmentation using 33% collision energy. MS2 scans were acquired in the ion trap with a maximum injection time of 35 ms and a standard AGC target. We performed online real-time search (RTS) of MS2 spectra against the E. coli proteome using a tryptic digestion pattern with maximum 1 missed cleavage, the static modifications carbamidomethyl on cysteines and TMTpro on lysines and N-terminus, the variable modification of oxidation on methionines and maximum 2 variable modifications. Scans that passed the standard RTS scoring thresholds (Charge State 2: Xcorr >= 1, dCn >= 0.1, Precursor PPM <= 25, Charge State 3: Xcorr >= 1.5, dCn >= 0.1, Precursor PPM <= 25) were sent for MS3. Using synchronous precursor selection (SPS) the 10 most intense matching fragments were selected with an isolation window of 1.2 Da, fragmented using HCD fragmentation at 45% collision energy and sent for an MS3 scan in the orbitrap at a resolution of 50000, scan range of 100-500 m/z, TurboTMT Off, 200% normalised AGC target and a maximum injection time of 200 ms.

### MS data search

#### DIA data

DIA files were searched individually in Spectronaut 18 or 19 using the file monitoring feature with a semi-specific (trypsin) peptide search. The resulting search archives were used to create spectral libraries for LiP and tryptic control separately. All LiP or tryptic control files of one experiment were then searched again for semi-specific peptides using the corresponding spectral library having the cross-run normalisation setting switched off. The same search pipeline was also used for the SEC-MS data, with the difference that only specific (trypsin) peptides were searched.

#### TMT data

The DDA raw files were converted to mzML files using MSConvert (v3.0). We used FragPipe (v21.1, MSFragger v4.0, Philosopher v5.1, IonQuant v1.10.12) for the peptide search with a TMT 16-MS3 workflow. For MSFragger^51^ a tryptic digest pattern with a maximum of 2 missed cleavages was used. We changed the variable N-terminal TMTpro modification to be constant but otherwise used the default settings. The results were filtered with an FDR of 1%. We use IonQuant for isobaric quantification of precursors, selecting TMT18 and quantification level 3. We group by peptide sequence and turn off normalisation as well as any ratio to abundance normalisation.

#### Compilation of annotated MBPs

We compiled a ground truth metal-binding protein (MBP) dataset using information from UniProt (Binding Site, Cofactor, Keyword, Catalytic activity) and QuickGO (molecular function). The information from all sources was combined in order to determine the most likely metal-ligand. We implemented this process in the protti R package^48^ through the “extract_metal_binders” function, which can be applied to any proteome in the UniProt database^52^ in order to extract all metal-binding information. In brief, we use ChEBI^53^ identifiers in combination with manual annotations in order to determine which ligands contain a metal atom. The protti R package contains a dataset of all existing metals. The symbol of each metal is used to determine which entries of the ChEBI database (2 and 3 star annotations) contain a metal atom in their formula. We also manually add all metal-ligand ChEBI identifiers that appear in UniProt, but that do not have an associated formula. This information is used to select all UniProt proteins with a metal-ligand in the “Binding Site”, “Cofactor” or “Catalytic activity” fields. In addition, keywords indicating the presence of a metal-ligand are extracted using the elemental metal names from the dataset in protti as well as the additional keywords “Metal-binding”, “Metalloprotease”, “2Fe-2S”, “3Fe-4S”, “4Fe-4S” and “Heme”.

We generated a metal-specific slim GO dataset containing 850 molecular function GO terms (“metal_go_slim_subset” in protti) by using the terms associated with metal-containing ChEBI identifiers. We also propagated ChEBI identifiers to any child terms that were not annotated. In addition, 230 terms of the complete dataset were manually added. This slim GO dataset was used to identify metal-binding proteins for each proteome by searching the QuickGO database.

In order to combine information from multiple sources, we made use of the hierarchical nature of ChEBI identifiers, always selecting the most specific one as the final ligand annotation if there were multiple associated ChEBI identifiers (e.g. “divalent metal cation” and “magnesium(2+)”). In addition, we use the formula of the ChEBI identifier to determine the bound metal (e.g. “heme b” contains “iron atom”).

Our computational method of metal-binding protein extraction could identify over 1000 metal-binding proteins for *E. coli* (Fig. 2A, **Supplementary Fig. 7B**). The most common annotation sources were “GO term” and “Keyword”, while “Catalytic activity” only accounted for a small number of annotations, since for those the metal needs to be directly involved in the reaction (Supplementary Fig. 7A).

For the functional assignment of metal-binding proteins to six major functional categories (catalytic, structural, regulator, transport, electron transfer, biosynthesis), information from UniProt was used. EC numbers were automatically extracted and the corresponding proteins assigned as “catalytic”. All sites in entries lacking an EC number were manually reviewed and assigned a role based on the contents of other selected UniProt entry fields.

#### Metal-binding site predictions

The structure-based predictors used in this study are Metal3D (M3D)^26^, GASS^27^, and Master of Metals (MoM)^28^. These predictors infer metal-binding sites using homology to known structures or distance-based features.

Metal3D is a deep learning-based tool that operates on a voxelized representation of the protein environment, predicting metal density on a per-residue basis. GASS employs a parallel genetic algorithm to identify candidate metal-binding sites that are structurally similar to curated templates from M-CSA^54^ and MetalPDB^1^. Master of Metals combines a neural network with a filtering step that compares the network’s output against local structural features of known binding sites, using distance matrices of Cα and Cβ atoms for comparison. For parameter selection, each metalloprotein predictor was configured following the criteria detailed in their respective original articles to ensure optimal performance.

Metal3D processes protein structures and specific residues, converting the local environment around each residue into a voxelized format. It predicts metal density per residue, and these values can be averaged to calculate an overall zinc density (ρ) for the entire protein. Only predictions with ρ ≥ 0.75 are retained for further analysis.

GASS identifies metal-binding sites using a parallel genetic algorithm and templates from M-CSA and MetalPDB. Candidate sites are scored based on a fitness value that quantifies the structural match between the candidate site and the reference template. This score is derived from geometric distances between residues in the candidate site and those in the template, with lower fitness values indicating higher similarity and better predictions. Only predictions with a fitness score ≤ 0.5 were considered for further analysis.

Master of Metals identifies triads or quadruplets of amino acids with appropriate spatial arrangements in protein structures provided as input. These candidate sites are then ranked by their structural similarity to templates in the MetalPDB database. Similarity is assessed using a distance parameter (d_max), with smaller values indicating higher similarity and more accurate predictions. Only predictions with d_max ≤ 0.35 Å were considered for further analysis.

To perform a proteome-wide prediction of metal-binding sites across the *E. coli* proteome we obtained the list of proteins from UniProt^52^. From this list, we retrieved all structural models of proteins available in the AlphaFold database^55–57^. AlphaFold models provide a per-residue confidence measure, plDDT, which reflects the local accuracy of the prediction. To avoid biasing the prediction results due to model quality, we retained only the structures with at least 90% of their residues having a plDDT > 0.7.

We consider a metal-binding site to be high confidence if it is predicted by at least two predictors. For the distance calculations we only use high confidence sites.

### Data analysis

#### Growth Curves

OD_600_ measurements of growth assays were exported from CLARIOstar MARS using the “Table View” and selecting “All Cycles”. We converted the data into the input format for the QurvE R package^58^ and ran the QurvE analysis. The negative controls were used as blanks and subtracted from the sample values for each time point. We mainly used the linear fits and if indicated spline fits for growth parameter estimation.

#### LiP-MS data

We used the protti R package^48^ for data analysis, following the dose-response workflow guidelines outlined in the protti documentation. In every experiment, we fit a dose-response curve to peptide precursor ions and computationally validate hits using a computational pipeline with multiple lines of evidence, as outlined in Supplementary Fig. 3 and below. In this pipeline, we use parameters of each precursor that we obtained from the following sources. From the 4-parameter log logistic regression curve fits, we obtain parameters that we use to assess the quality of the fit (Supplementary Fig. 3A). In addition, we calculate the pearson correlation between the curve and the measured data points to estimate the goodness of fit, as previously established^15^ (Supplementary Fig. 3B). Lastly, we also perform an ANOVA statistical test to find precursors that have a significant response larger than noise.

The principles for each validation step are the following. On the precursor level, we validate that any sibling precursors of a significant peptide are showing a similar behaviour. Precursors that are part of the same parent peptide group should behave in the same way, if the dose-response behaviour comes from a structural change on the peptide level. If this is not the case, this can be explained by a modification (e.g. oxidation), an artefact of the MS acquisition or a false peptide assignment during data search. On the protein level, we collect additional evidence for significant precursors by checking for precursor sequence overlaps and co-occurrence of multiple significant precursors in the same protein. Lastly, on the treatment level, we check if any significant precursors from one condition are also significant or close to significance in other conditions. This allows us to better compare conditions qualitatively and to not draw the wrong conclusions about the exclusivity of hits, because they were barely below our significance thresholds.

Before precursor-level validation, we look for instances of unrealistic parameters or systematic missingness and assign specific values for some of their parameters, as explained below. We specifically assign values to the correlation r and hill coefficient b parameters, because these are used for tests in the precursor level validation pipeline. We specifically assign not naturally occurring values such as 0 or NA to make sure that they are recognised and correctly handled in the automatic analysis pipeline. Precursors that have a partially systematic missingness (missing not at random, MNAR) on the curve level (Supplementary Fig. 4C), as classified by the protti package, are assigned with a correlation r = 1 and hill coefficient b = 0. Even though these cases are rare, excluding them could lead to an underestimation of potentially relevant effects. We only consider MNAR precursors qualitatively as hits meaning parameters from curve fits are not used for further analysis. For cases in which no curve could be fit, we annotate with r = 0 and b = NA, marking these precursors as non-significant. All precursors with an absolute r < 0.01 (a very flat slope) are annotated with b = NA. The same applies for precursors with an EC_50_ value c that is 20 times larger than the maximally used concentration for the dose-response curve. These cutoff values were empirically validated by us and usually appear for poor curve fits (low r). If we set b = NA for these cases, we make sure that whatever unrealistic b values were present before, are not actually considered for validation of a significant sibling precursor. A significant precursor with such a sibling precursor could be a false positive hit and is therefore considered only if further validated by additional evidence (protein and treatment level).

In the precursor level validation (Supplementary Fig. 4C) we check that the precursor passes a completeness criterion of at least 2 replicates in at least 4 concentrations. This is followed by the significance test in which we consider every precursor with a correlation r ≥ 0.8 and an adjusted p-value ≤ 0.05 of the ANOVA test as significant. We chose the cutoff of 0.8 because this consistently has a false discovery rate of 0.05 in simulated data (not shown). As outlined in the figure scheme we then move on to perform checks of each significant precursor with its sibling precursors of the same parent peptide group. This group importantly also contains modified versions of the same precursor. In our analysis we only look for oxidation as a variable modification, but technically this can also be expanded for other modifications. In brief, an annotation of “n1” indicates that only a single precursor is present for the parent peptide group, making cross-validation impossible. Thus, these are accepted as valid due to the absence of contradictory evidence. If there are any sibling precursors in the same group that have a low correlation r < 0.5, the annotation gets the suffix “_low_r”. If a precursor fails all standard checks it gets passed on to check for an oxidation change that could explain an unexpected opposing behaviour of modified vs. unmodified precursors (as outlined in Supplementary Fig. 4D). If this is also not the case the significant precursor gets the annotation “verify” indicating that it is potentially problematic. Taken together this means that with this type of validation, no precursors are removed from the significant list, problematic precursors are only annotated as “verify”. It is the subsequently performed protein and treatment (batch) level validation that aims to validate these “verify” precursors as well as give additional evidence to precursors that are labelled as “n1”.

For protein level validation, we check if a significant precursor either has other significant precursors overlapping with its sequence or if there are other significant precursors in the same protein (Supplementary Fig. 4E). For batch level validation, we check if the same significant precursor is also found in another related dataset (Supplementary Fig. 4F). Here we have two levels of validity, only precursors with r ≥ 0.8 and any with r ≥ 0.5 and r < 0.8. The only way to actually expand the original number of significant hits is by this second category, which also considers slightly less confident precursors as hits if they are highly confident in another treatment. This approach is particularly relevant for comparing hits across different datasets. To conclude that a hit is unique to a dataset we want to make sure that it does not barely miss the cutoff of significance in the dataset it is compared to.

The two new levels of evidence are then used to annotate “n1” precursors or “verify” precursors in the mentioned order. Any precursors that are still labelled as “verify” afterwards are removed and annotated as “not_valid”. Taken together the validity annotation is mainly meant to remove not trustworthy precursors from the hit list as well as to improve the overlap between datasets, by including hits that are slightly below the cutoff.

### TPP-TMT data

We first filtered for precursors that were confidently identified in both the online RTS and offline search. Subsequently, we correct for isotopic impurities using the R package MSnbase^59^. We manually remove all peptides that cannot be uniquely attributed to a specific protein. Then we used MSstatsTMT^60^ to consolidate precursors across fractions and to calculate protein intensities (without global normalisation). We calculated fold changes using the 37°C temperature as a reference point for each condition. The TPP R package was used for curve based normalisation as described in^12^, followed by nonparametric curve fitting and significance analysis with empirically determined degrees of freedom using the NPARC R package^61^. We only consider proteins that have no missing values in their melting curves for at least two of the three replicates for each condition.

### SEC data

We manually remove all peptides that can not be uniquely attributed to a specific protein and the precursors with intensity lower than 50. We then used the R package CCprofiler^62^ to further filter the data and perform the differential analysis between conditions. First, we imputed missing values by spline interpolation, then we normalized the peptide traces across replicates, conditions and fractions by Cyclic Loess. Next, we removed all the precursors that were identified in less than 3 consecutive fractions and those not having at least one high correlating (0.8) sibling peptide and absolute sibling peptide correlation lower than 0.6. Lastly, we kept only proteins with at least 5 precursors. Having completed the filtering steps, we moved on with performing the differential analysis at the protein-feature level, where a ’protein feature’ is a peak group of coeluting peptides. The fold change was calculated using the 0mM EDTA sample as reference. For the purpose of comparing the SEC data with LiP and TPP data, we considered as hits all the proteins that present a significant assembly change when treated with 10mM EDTA.

### Active Site Analysis

For the distance to enzyme active site comparison, we retrieved all annotated E.coli K-12 (UP000000625) active sites from UniprotKB. For each significant LiP peptide in enzymes with active-site annotations, we computed the minimum Cα–Cα distance between any peptide residue (start–end range) and any annotated active-site residue, using AFDBv4 models. Distances were summarized and compared by functional assignment.

To determine a minimum distance threshold value, we computed Youden’s J statistic between significant vs. non-significant peptides with respect to the minimal distance to a metal binding site. That value was 12.66 Å. However, we opted for a more conservative value of 10 Å.

### Structural Distance Analysis

For the identification of known and novel metal-binding sites we performed 3D distance measurements between peptides and annotated or predicted metal-binding sites in AlphaFold predicted structures.

Ribosomal proteins were excluded from this analysis due to their poorly predicted structures and because they undergo extensive structural destabilisation upon chelator treatment, which makes them less suitable for binding site inference. For each peptide, the minimum distance between any peptide residue and the metal-coordinating residues of the annotated site was determined. For proteins with multiple annotated sites we considered only the shortest distance. Statistical comparisons between distance distributions of hit and non-hit peptides were performed using the Wilcoxon rank-sum test, with p-value < 0.05 being considered as significant.

The average of median distances for hit peptides was 4.6 Å across chelators, which we used as a threshold to identify binding sites. “Additional metal-binding sites” are sites in which hit peptides are closer to a predicted site than to a previously annotated site. “Novel metal-binding sites” are sites in which a predicted site is within 4.6 Å of a hit peptide, but there is no previous metal-binding site for this protein.

### Cloning

The pCN24A plasmid of the ASKA library^63^ was used as the backbone for all overexpression plasmids. We changed the N-terminal tag from His_6_ to FLAG to minimise any non-specific metal-binding of the purified protein. For this we amplified the original plasmid with Q5^®^ High-Fidelity DNA Polymerase (NEB) using the reverse primer in order to introduce the FLAG tag (pCN24A_FLAG_rev: 5’-CTTGTCGTCATCGTCTTTGTAGTCcatagttaatttctcctctttaa tgaattctgtgtg-3’) and a standard forward primer (pCN24A_fw: 5’-taagggtcgacctgcagccaag-3’).

The gene inserts were amplified by PCR directly from genomic *E. coli* DNA with primers that have a 25 bp overlap with the backbone. The siriusGFP was amplified by PCR from the plasmid pMJ922 (Addgene, #78312). Final plasmids were generated using Gibson Assembly (NEB). Every plasmid was verified by full-plasmid sequencing.

### Protein expression and purification

For large-scale purification 4 L of transformed BL21 (DE3) competent cells (Thermo Fisher Scientific, #EC0114) were grown in LB medium containing 100 µg/mL chloramphenicol. When OD_600_ reached 0.5, we induced with 100 µM IPTG and transferred the cultures to 16°C overnight. TrhO was induced with 0.4% lactose^64^ instead, because IPTG only lead to low levels of expression.

We lysed cells according to the manufacturer’s instructions using CelLytic™ B 2x (Sigma, #B7310), 0.2 mg/mL Lysozyme (Sigma, #62971), 50 units/mL Benzonase (Millipore, #E1014) and cOmplete™, EDTA-free Protease Inhibitor Cocktail (Roche, #04693132001). The lysate was cleared by centrifugation at 20,000 g for 20 minutes and filtered through a 0.4 µm filter. The proteins were purified using 10 mL (bed volume) Pierce™ Anti-DYKDDDDK Affinity Resin (Thermo Fisher Scientific, #A36804), which should have a capacity of 30 mg. We pre-washed the resin with TBS (50 mM Tris pH 7.5, 150 mM NaCl) and added the cleared lysate for incubation for 1 hour at 4°C on a roller. The resin was washed with 20 column volumes (CV) of TBS. We eluted the protein with 1.5 mg/mL 3x FLAG peptide (MedChem, #HY-P0319) by incubation with 8 mL for 30 minutes at 4°C and repeated the process once followed by rinsing the resin with TBS. We also performed a subsequent elution using glycine pH 2.8 in order to remove any leftover protein from the column. The acid elution was processed separately and not pooled with the 3x FLAG peptide eluted protein. We concentrated the elution using an Amicon^®^ Ultra Centrifugal Filter, 10 kDa MWCO (Millipore, #UFC9010) until the volume reached 5 mL. We further purified the protein by size-exclusion chromatography using a HiLoad 16/600 Superdex 75 pg or 200 pg column (Cytiva) in TBS. We analysed elution fractions by SDS-PAGE (4-12% Bolt gels, MES buffer, Thermo Fisher Scientific) and stained with QuickBlue Protein Stain (LubioScience, #LU001000). Fractions containing pure protein were pooled, supplemented with 5% glycerol and snap frozen in liquid nitrogen before storage at −80°C.

### Western Blotting

We used the BioRad Trans-Blot Turbo Transfer System in combination with the Trans-Blot Turbo Transfer Pack (BioRad, #1704158), which contains a nitrocellulose membrane. We transferred using the standard program with constant 25 V, up to 1.0 A for 30 minutes. The membrane was blocked for 1 hour with 5% milk powder in PBST (137 mM NaCl, 2.7 mM KCl, 1.4 mM KH_2_PO_4_, 4.3 mM Na_2_HPO_4_, 0.1% Tween-20). We incubated with the FLAG M2 primary antibody (Sigma, #F1804) overnight at 4°C. The blot was washed three times with PBST. We incubated with an anti-mouse secondary antibody coupled to Alexa Fluor Plus 680 (Thermo Fisher Scientific, #A32729) for 30 minutes at RT. After three final washes, the blot was imaged with a LI-COR imaging system.

## Data availability

All MS proteomics data have been deposited at ProteomeXchange Consortium via the PRIDE partner repository. The LiP-MS chelator dataset has the identifier PXD061428. The EDTA TPP dataset has the identifier PXD061179. The magnesium LiP-MS dataset has the identifier PXD062222. The PK activity assay datasets have the identifier PXD062281. The modified M9 medium dataset has the identifier PXD062302. The EDTA SEC-MS dataset has the identifier PXD061480.

## Code availability

All data analysis scripts can be found on GitHub (https://github.com/jpquast/LiP-Metal-Map).

## Author Contributions

J.P.Q. designed and performed LiP and TPP experiments, designed the computational pipeline and performed bioinformatics analysis. A.P. performed the SEC-MS experiment and performed bioinformatics analysis. R.J.S., A.P., X.L., E.Z. and L.G. assisted with experiments. G.M. assisted with data analysis. M.S.P.C. performed the chelator concentration determination. J.P.Q. and M.D. purified proteins for validation. J.P.Q. and R.J.S. prepared ICP-MS samples and performed the computational analysis. C.C. performed computational predictions of metal-binding sites. N.d.S., J.P.Q., R.J.S. and P.P. wrote the manuscript with contributions from all authors. C.A., A.R., P.B. and N.Z. supervised parts of the project. P.P. conceived and supervised the project.

## Acknowledgements

The authors acknowledge Dr Mario Leutert (ETH Zurich) for assistance with TMT data analysis, and Roberta Florea (ETH Zurich) for preliminary protein purification experiments. We further want to thank Gary Sharples (Durham University) and Caryn Outten (University of South Carolina) for making raw data from previous studies available for comparisons. This work was supported by the Promedica Stiftung, Chur and the European Research Council [grant agreement number 866004].

## Supplementary Figures

**Supplementary Fig. 1:**
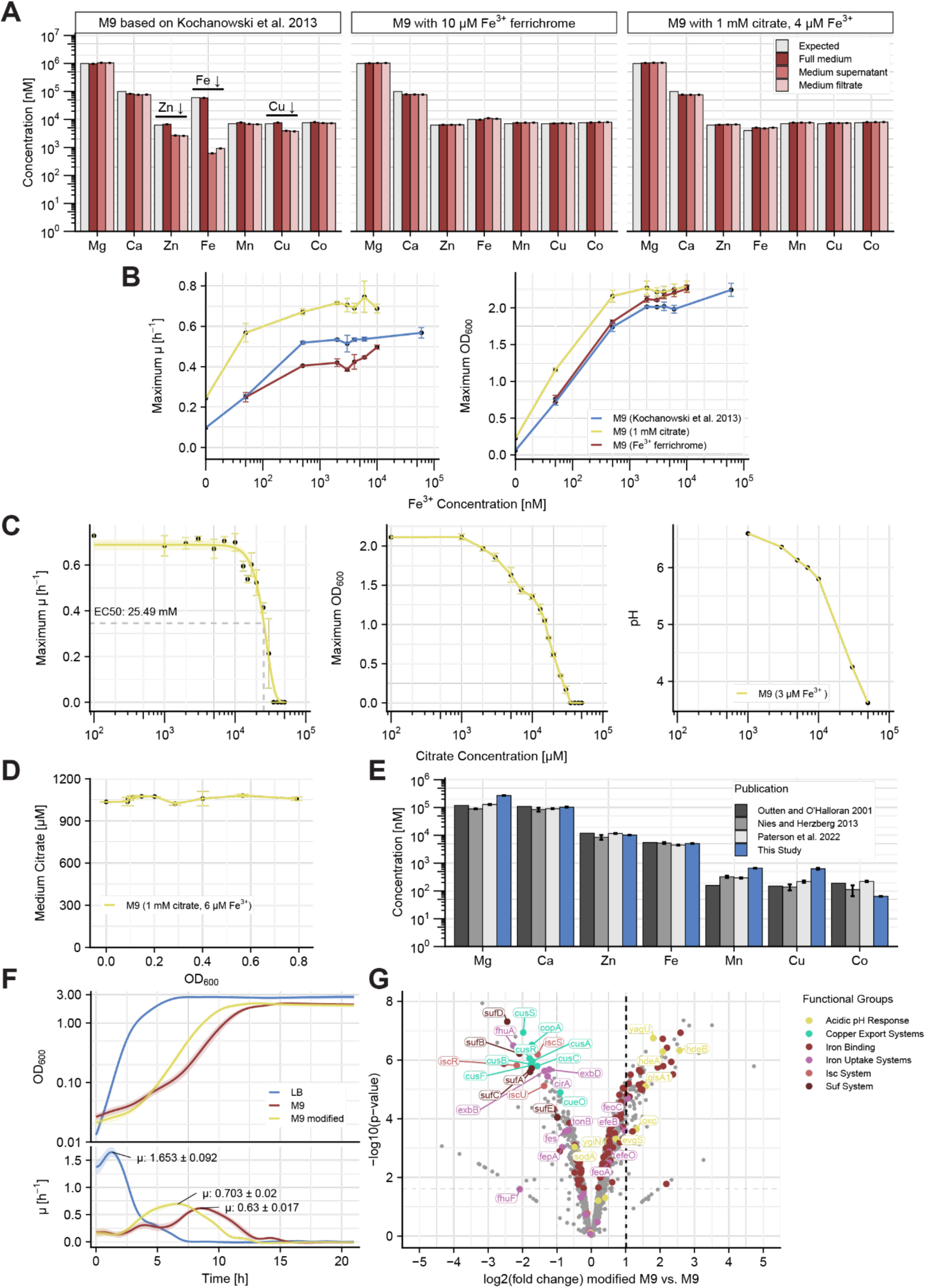
Optimisation of modified M9 medium composition for Metal-LiP. **A** Metal concentrations (ICP-MS) in three different M9 media. The full medium, supernatant of centrifuged medium and the filtered medium were compared to check for metal precipitation, which was only observed for the original M9 medium after Kochanowski et al. 2013^36^ (n = 3 technical replicates). **B** The maximum growth rate µ and maximum OD_600_ of *E. coli* grown in standard, citrate or ferrichrome M9 medium as a function of the FeCl_3_ concentration in the medium (n = 4 biological replicates). **C** The maximum growth rate µ, maximum OD_600_ and pH of M9 medium containing 3 µM FeCl_3_ as a function of the citrate concentration (n = 4 biological replicates). **D** The citrate concentration in M9 medium as a function of density shows that citrate is not consumed and stable over the growth to a density of OD_600_ = 0.8 (n = 2 technical replicates). **E** Metal concentrations (ICP-MS) in LB medium are comparable between this study (n = 3 technical replicates) and three previously published studies. **F** OD_600_ and growth rate µ over time for LB medium (n = 32 biological replicates), original M9 medium (n = 30 biological replicates) and modified M9 medium (n = 30 biological replicates). **G** Volcano Plot of protein abundance changes between the original and the modified M9 medium. Proteins are coloured based on several functional groups. Iron binding was annotated only after the other iron related groups were annotated. The horizontal cutoff line shows an adjusted p-value (Benjamini-Hochberg) of 0.05 (n = 4 biological replicates). The standard deviation is shown as error bars for panel A-E and as a shaded area around the curve for panel F.

**Supplementary Fig. 2:**
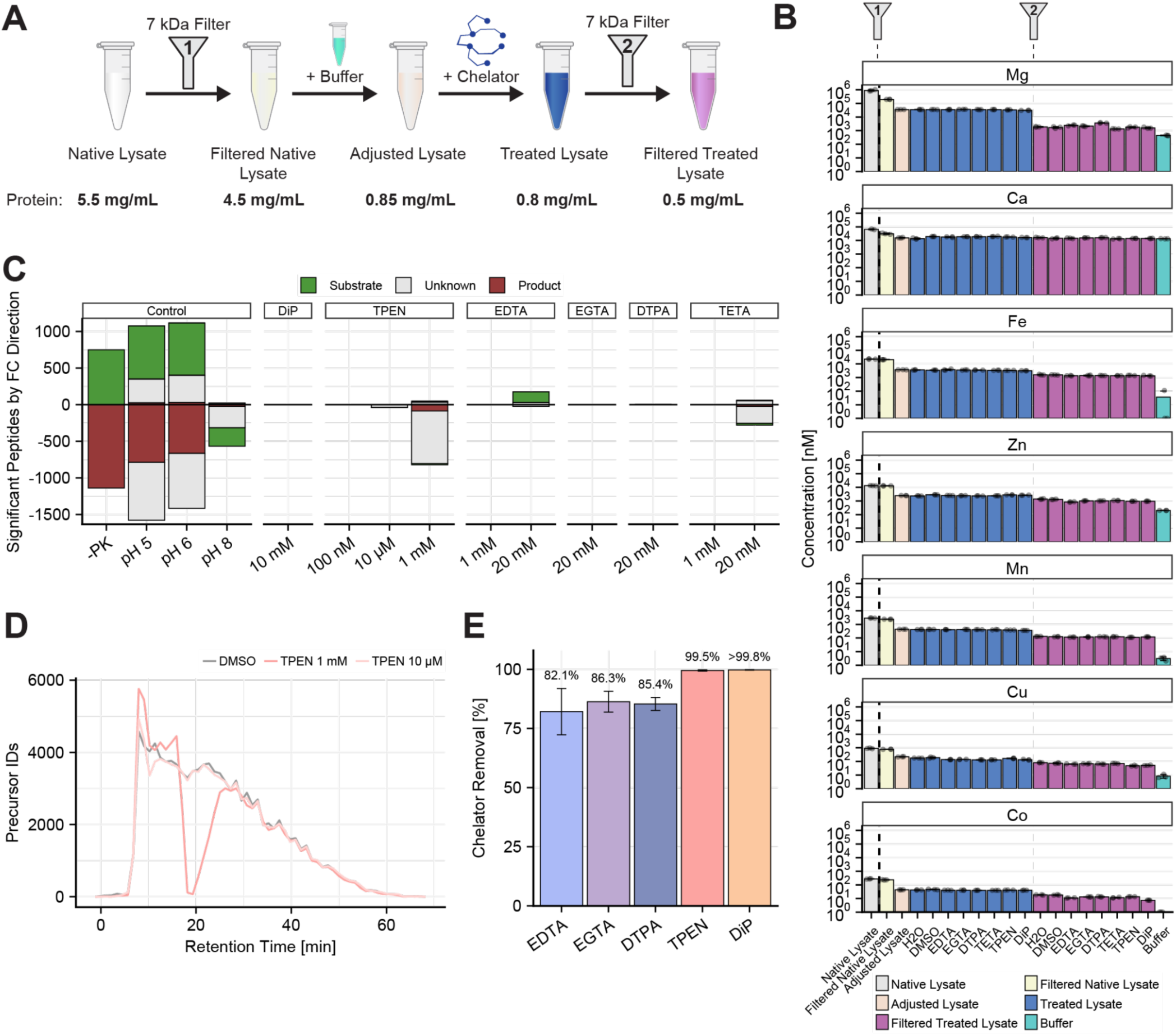
Development of the Metal-LiP workflow. **A** Detailed steps of lysate filtration and treatment including the protein concentrations determined at each step (n = 3 technical replicates). **B** Raw metal concentrations of the lysate at different steps of the workflow, as determined by ICP-MS (n = 3 technical replicates). The error bars show the standard deviation. **C** Effect of chelators on PK activity as determined by a PK activity assay (see Methods). Changes in PK activity are represented as changes in substrate (green) and product (red) peptides (annotated based on -PK control) (n = 3 technical replicates). “Unknown” peptides (grey) are not changing in abundance in the -PK control and can therefore not be labelled as substrate or product. pH 5 and 6 are controls expected to show a decreased activity while pH 8 is expected to show an increased activity. TPEN at 1 mM showed considerable ion suppression in the MS spectra at least partially explaining the peptide changes (see panel D). **D** Precursor ID plot over retention time showing the average for the DMSO control and two TPEN concentrations. There is clear ion suppression for 1 mM TPEN around the 20 minute mark and a small dip at 10 minutes for the 10 µM concentration. **E** Reduction in chelator concentrations by filtration with the 7 kDa size exclusion filter. The maximum chelator concentration (Fig. 1A) was used here. TETA could not be determined. n = 3 technical replicates, the error bars show the standard deviation.

**Supplementary Fig. 3:**
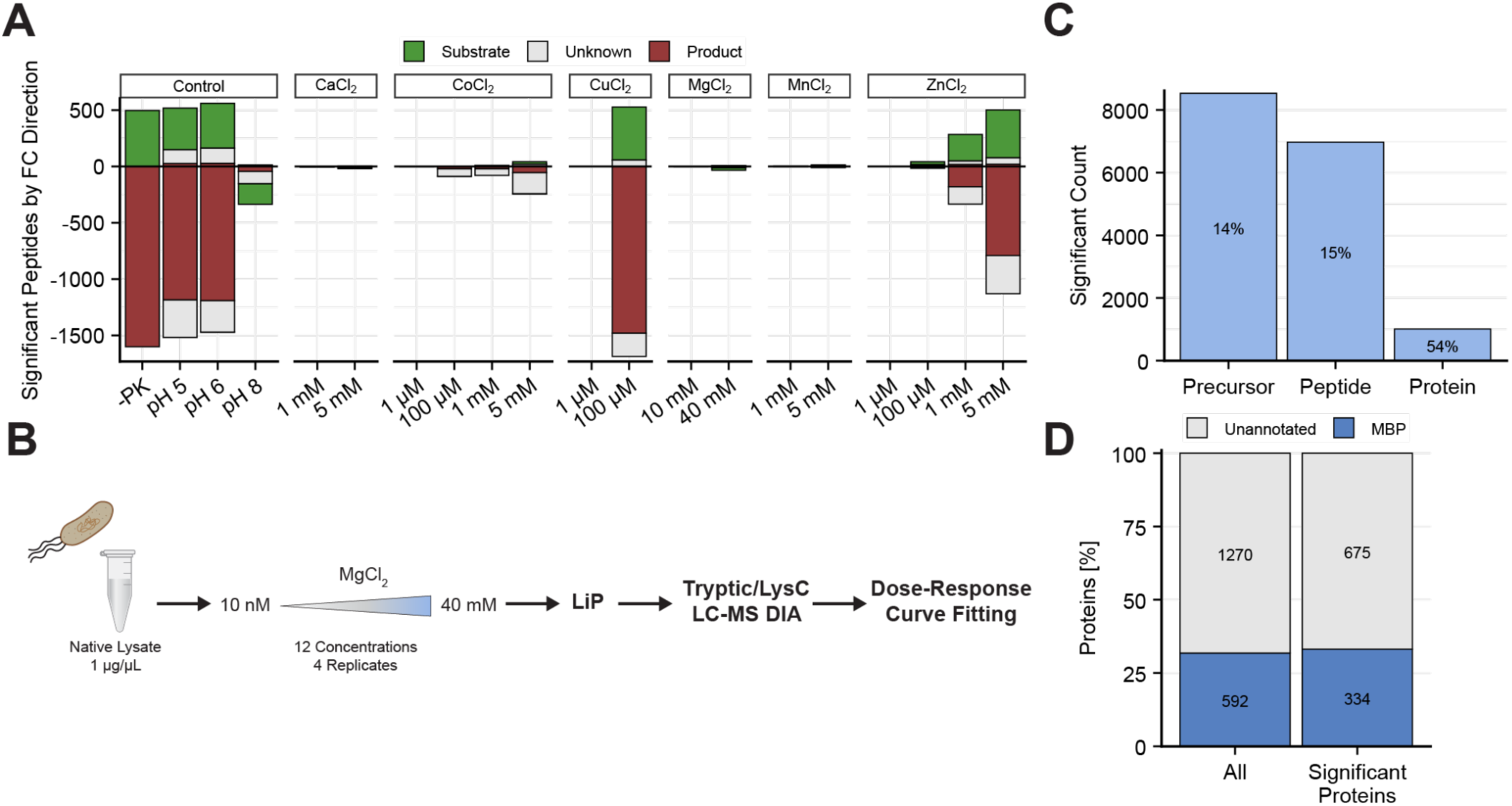
Chelator treatment is preferable to metal treatment in Metal-LiP. **A** Effect of metals on PK activity as shown by the changes in substrate and product peptides (based on -PK control) (n = 3). Unknown peptides are not changing in abundance in the -PK control and can therefore not be labelled as substrate or product. **B** Sample preparation pipeline for MgCl_2_ treated *E. coli* lysates. The 12 dose-response treatment includes a vehicle only control. **C** Number of significantly changing precursors, peptides and proteins with significant peptides. Percentages show the fraction of all detected precursors or proteins. **D** Percentage of MBPs for all detected proteins and for proteins with significantly changing precursors.

**Supplementary Fig. 4:**
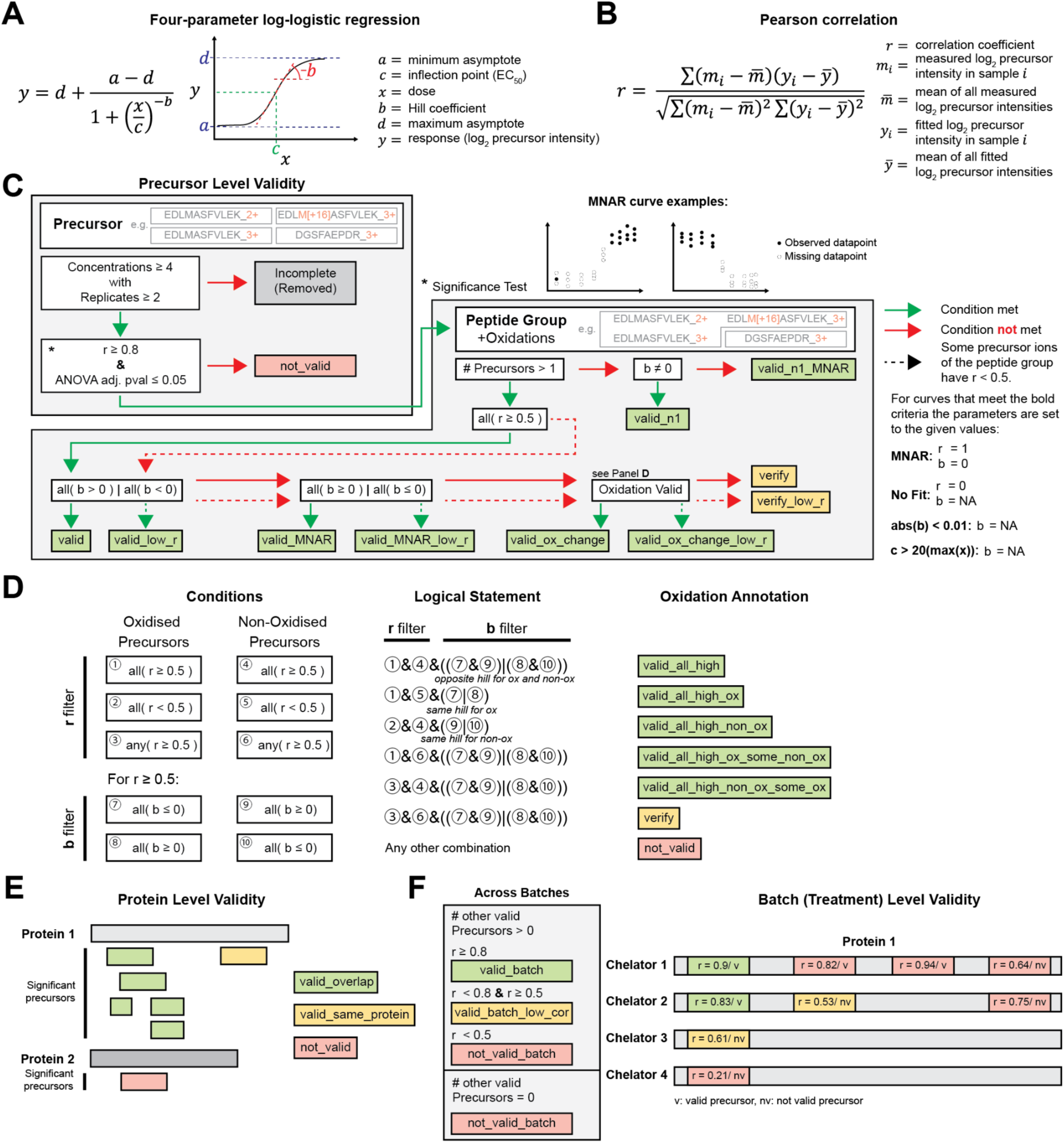
Data analysis pipeline for dose-response curve fitting and precursor hit identification. **A** Four-parameter log-logistic regression equation used for sigmoidal dose-response curve fitting. **B** Pearson correlation equation used to calculate the correlation between measured precursor intensities and the fitted dose-response curve. **C** Precursor level dose-response curve validity annotation. Prior to the annotation workflow, precursors belonging to one of the following groups are assigned a specific correlation and hill coefficient value: missing not at random (MNAR) doses, no successful curve fit, absolute hill coefficient of lower than 0.01 or EC50 that is 20 times higher than the highest dose point (see bottom right of panel). First, for each precursor ion a completeness and significance check (correlation and ANOVA adjusted p-value) is performed. Any precursor that passes the test is then evaluated alongside precursors of the same peptide group in terms of their correlations and hill coefficients (negative slope of dose-response curve). Lastly, a methionine oxidation change check is performed in “Oxidation Valid” (see Panel D) for precursors that have an oxidised peptide in the peptide group. **D** There are a given set of conditions for both the correlation (r) and hill coefficient (b) for oxidised and non-oxidised precursors. For example, if there are oxidised and non-oxidised precursors in the same peptide group and they have an opposite response (hill coefficient), this is considered a significant oxidation change in response to the treatment. Based on specific combinations of these statements we assign different levels of confidence for an oxidation change. All “valid” and “verify” outcomes are considered a pass of the “Oxidation Valid” test. **E** Protein level validity annotation. All significant precursors (r ≥ 0.8 and ANOVA adjusted p-value ≤ 0.05) of a chelator treatment are checked for overlap and if they are part of the same protein. This annotation is only considered as further evidence for precursors that got a “verify” flag in panel C and precursors that were “n1”. **F** Batch level validity annotation. Each distinct treatment is considered a batch. For each precursor we check if it is valid (based on panel C) in other batches. If there is at least one other valid precursor, depending on its own correlation it falls into different validity categories. For precursors that were already valid this boosts confidence for “n1”. Precursors that were not valid yet become valid due to this annotation, creating a hitlist that is more consistent across treatments.

**Supplementary Fig. 5:**
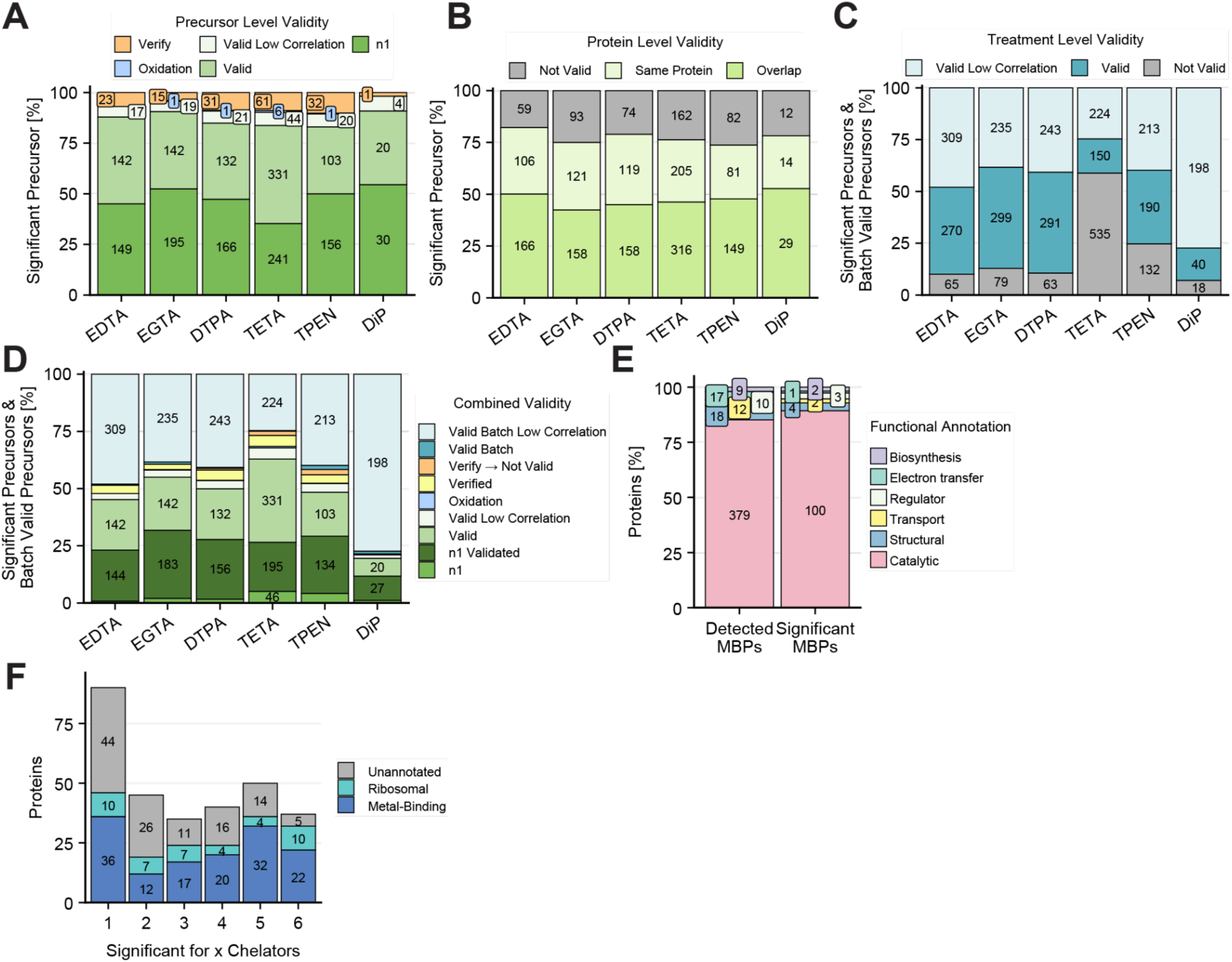
Validity of significant precursors in Metal-LiP. **A** Precursor level validity applied to the initial precursor hits based on our validity annotation approach (see Methods and Supplementary Fig. 4). **B** Protein level validity annotation applied to the initial precursor hits. **C** Treatment (batch) level validity, comparing the initial precursor hits of all chelators and checking for overlaps (“Valid”) and overlaps with lower correlation (r < 0.8 & r ≥ 0.5) precursors (“Valid Low Correlation”). This shows that TETA has a lot more unique precursor hits. **D** Combination of all three validity types combining initial precursor hits (panel A and B) with treatment (batch) level additional precursor hits (panel C). **E** Distribution of functional categories among MBPs with a binding-site annotation for LiP hits (n = 127) compared to all detected proteins (n = 437). No significant difference between distributions was observed (χ² = 4.04, p = 0.54). **F** The plot shows the number and types of proteins that were significant for a given number of chelators.

**Supplementary Fig. 6:**
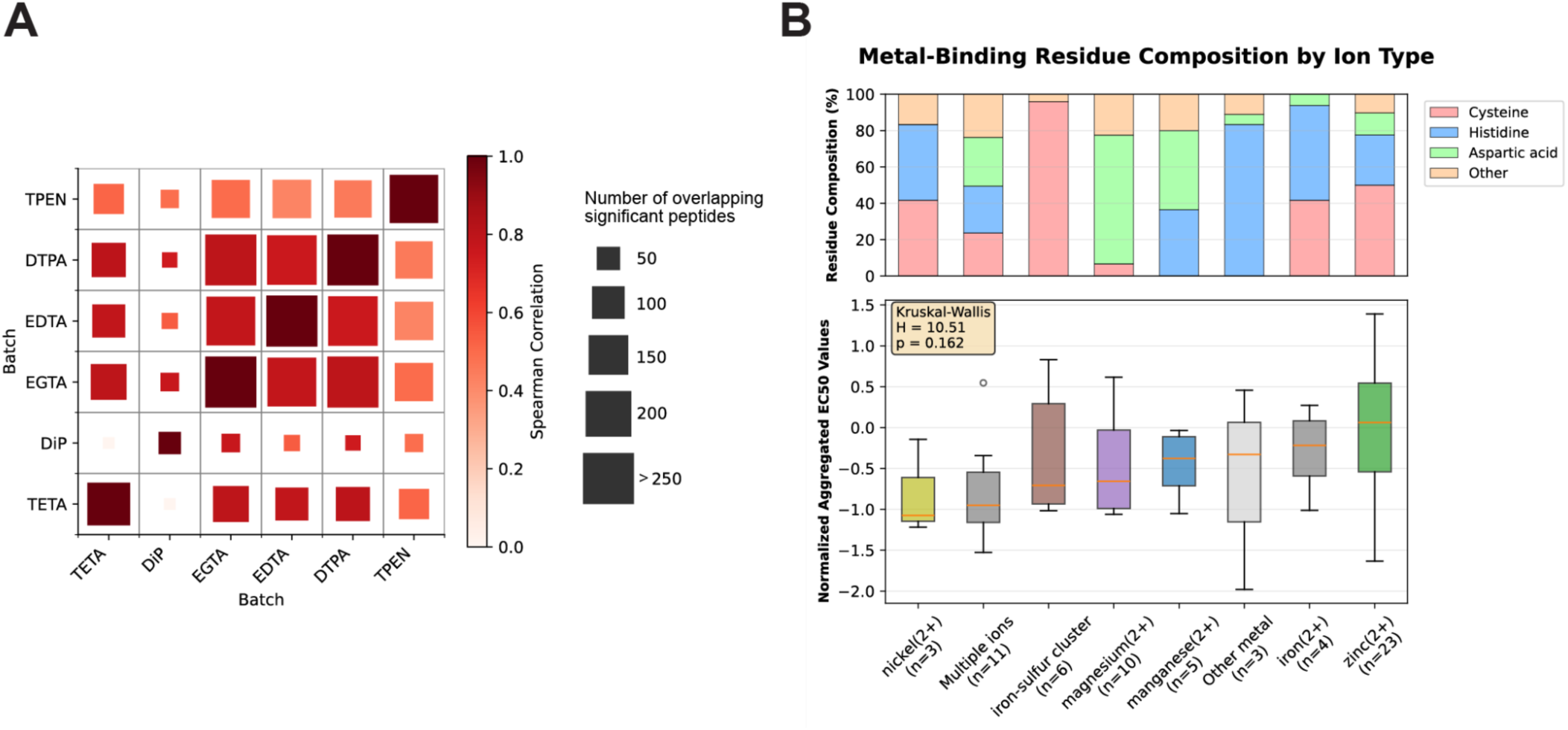
**Relative metal affinity analysis**. **A** Spearman correlation of precursor EC_50_ values extracted from the Metal-LiP experiment. Spearman correlation computed over shared significant precursors. Significant precursors are those with a Pearson correlation between the dose-response curve and the measured data points of r ≥ 0.8 and an adjusted p-value ≤ 0.05. Correlation of EC_50_ values between chelators is computed only over shared significant precursors. The number of shared significant precursors is indicated by the size of the cells in the matrix. **B** Metal-type to EC_50_ value relationship and residue composition. (**Upper**) Proportional stacked bar chart showing the compositions of residue types across different types of metal ions. (**Lower**) Boxplot showing trends in per-protein aggregated EC_50_ values across different types of metal ions. Annotated metal-binding proteins with at least one significant precursor are included. To aggregate precursor EC_50_ values from different chelators, within-chelator Z-scores are computed and then averaged. Only precursors that are significant across at least 2 chelators are included. To examine protein-level relationships between estimated EC_50_ values and metal type, precursor-averaged Z-scores are collapsed by taking per-protein medians.

**Supplementary Fig. 7:**
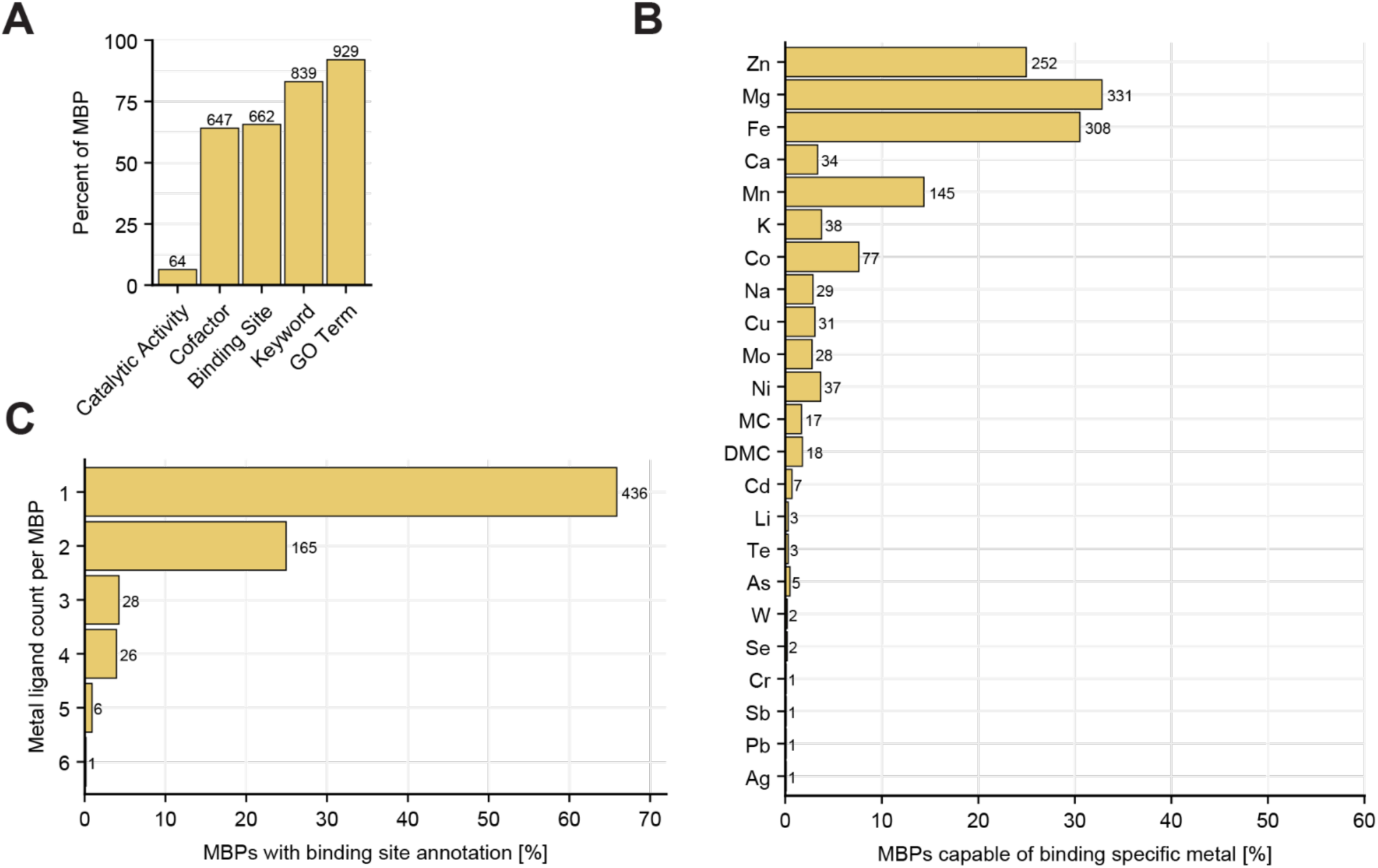
Known MBPs in the *E. coli* proteome. **A** Source composition of annotated metal-binding protein references for *E. coli* expressed as the percentage of all annotated metal-binding proteins for the given species. GO terms were obtained from QuickGO, “Catalytic Activity”, “Cofactor”, “Binding Site” and “Keyword” are metal-binding related terms from UniProt. **B** Percentage and number of metal-binding proteins capable of binding a specific metal. Some proteins bind multiple metals, therefore the percentage adds up to more than 100%. **C** Percentage and number of metal-binding proteins with an indicated number of metal-binding sites. Only uses proteins with “Binding Site” annotations.

**Supplementary Fig. 8:**
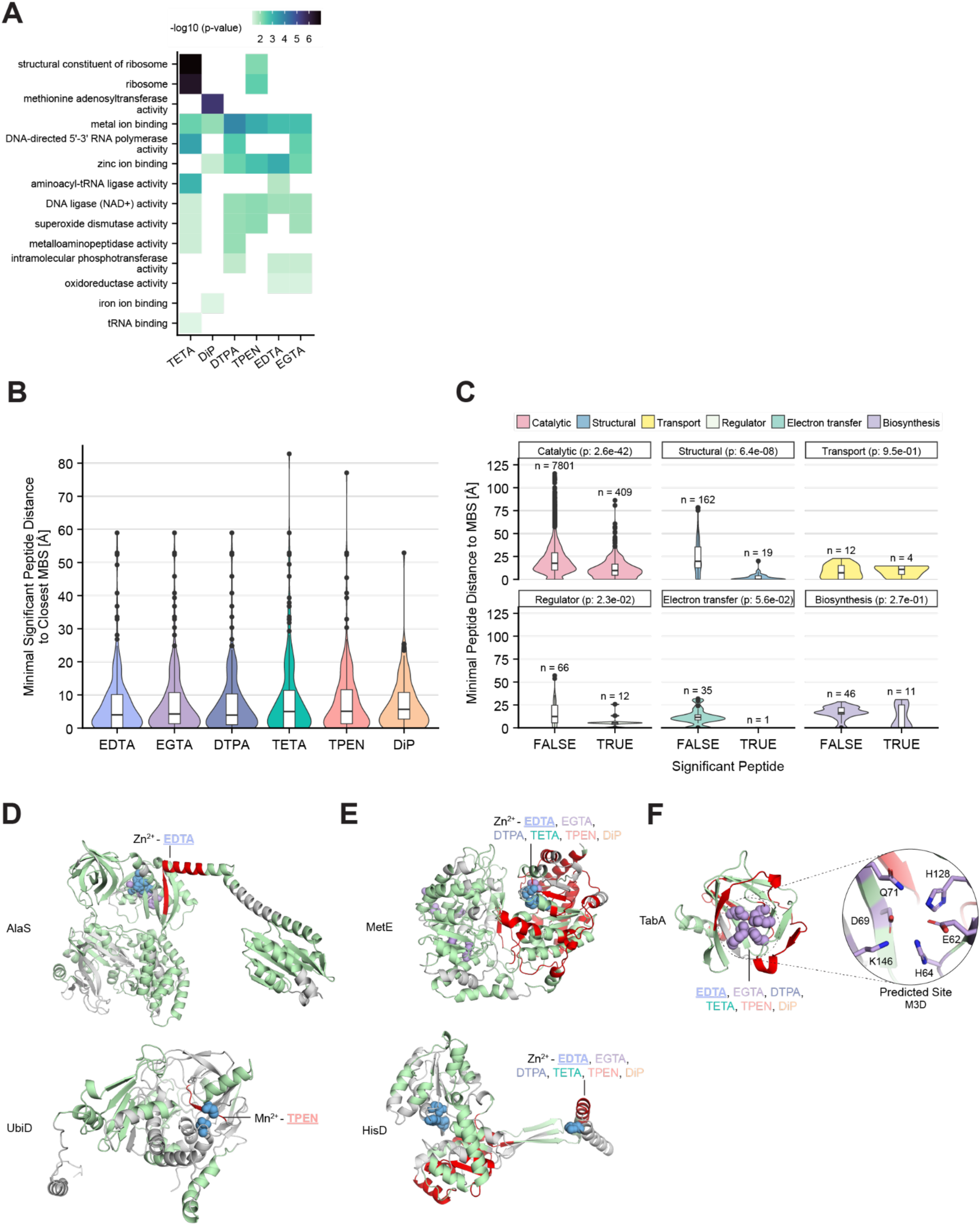
Contextualisation of Metal-LiP hits in 3D structures and protein domains. **A** InterPro domain level molecular function GO enrichment. The significance cutoff for the fisher’s exact test is a p-value of 0.05. Only terms with at least three significant proteins are plotted. **B** The violin plot shows the minimal distance distribution of significant peptides to the closest annotated metal-binding site of MBPs. **C** The violin plot shows the minimal distance distribution of peptides to the closest annotated metal-binding site of MBPs. For each functional MBP category, significantly changing (i.e., hit) and non-changing peptides are plotted separately. p-values comparing the ranks of changing and non-changing peptides are shown (Wilcoxon rank sum test). n indicates the number of peptides per distribution. **D-E** AlphaFold predictions of the indicated proteins. Grey: Regions not detected by MS, Green: Detected, but non-significant regions, Red: Significant peptides, Blue: Annotated metal-binding sites, Violet: metal-binding sites predicted by at least 2/3 predictors (C-D), or 1/3 predictors (E). We indicate which chelators have a significant peptide in the metal-binding site and underline the chelator of which the labeled AlphaFold prediction is displayed.

**Supplementary Fig. 9:**
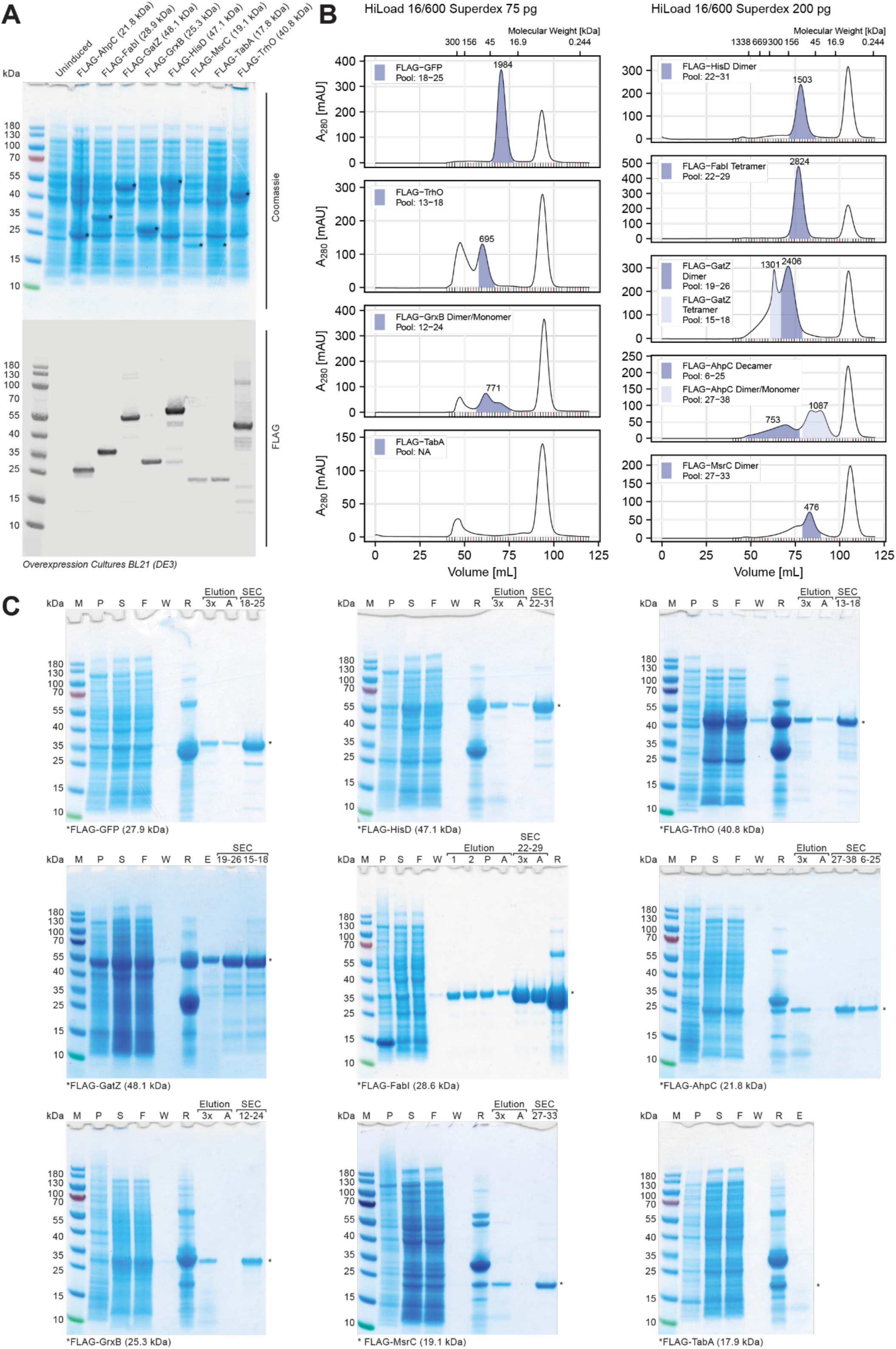
FLAG purification of candidate MBPs identified by Metal-LiP. **A** SDS page stained with coomassie (top) and western blot with anti-FLAG (bottom) for FLAG-tagged proteins overexpressed in BL21 (DE3) cells after induction with IPTG or Lactose (TrhO). Asterisks indicate the overexpressed proteins. **B** SEC profiles of proteins purified on either the 75 or 200 pg superdex columns. The shaded areas indicate the pooled fractions. The large peak on the right is the 3x FLAG peptide used for the protein elution from the FLAG-resin. **C** Complete stepwise protein purification for all 9 expressed proteins. M: PageRuler™ Prestained Protein Ladder, 10 to 180 kDa, P: Lysate Pellet, S: Lysate supernatant, F: Lysate after resin incubation, W: 4th wash step, R: resin after 3x FLAG peptide elution, E: pool of 3x FLAG peptide elution, Elution: 1: First round of 3x FLAG peptide elution, 2: Second round of 3x FLAG peptide elution, P/3x: Pool of 3x FLAG peptide elution, A: Pool of acid elution, SEC: Pooled fractions of 3x FLAG peptide elution SEC run or acid elution SEC run. If not otherwise indicated only the prior is shown.

**Supplementary Fig. 10:**
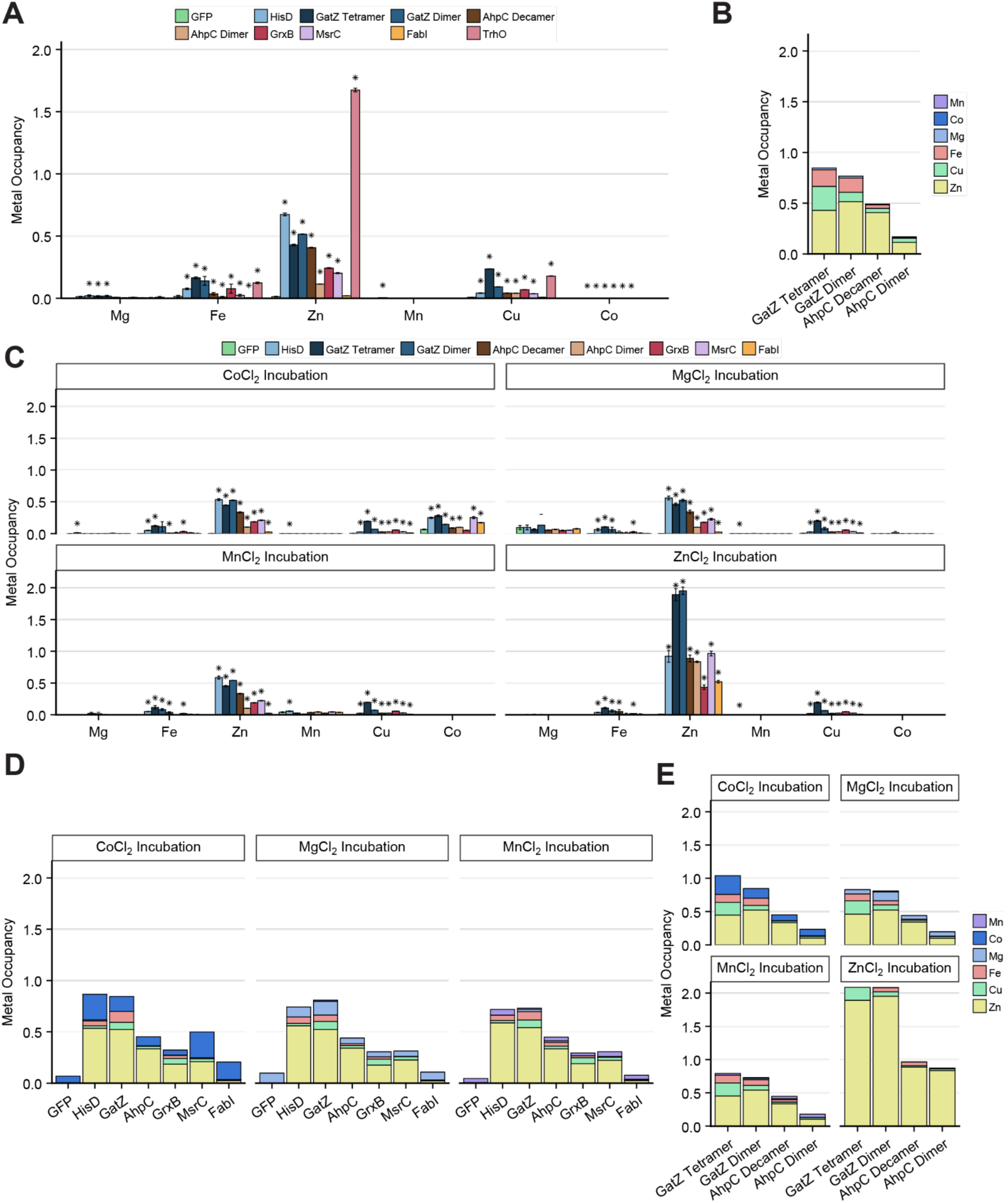
ICP-MS analysis of candidate MBPs *in vivo* and *in vitro.* **A** Metal abundance normalised to protein abundance (metal occupancy) for the indicated purified proteins, measured by ICP-MS. Error bars represent the standard deviation. Asterisks indicate a significant change (p-value <= 0.05) compared to the GFP control as determined by a one-sided Welch’s t-test. **B** Stacked metal occupancies for the indicated assembly states of GatZ and AhpC. **C** Same as panel A, but the purified proteins were incubated with a 2-fold molar excess of the indicated metal followed by filtration, prior to metal abundance determination. **D** Stacked plot of metal occupancies for purified proteins incubated with a 2-fold molar excess of the indicated metal. For GatZ and AhpC the dimer and decamer are shown respectively. **E** Same as panel D, but showing all assembly states of GatZ and AhpC. All metal abundances in panels A-E were determined by ICP-MS in triplicate.

**Supplementary Fig. 11:**
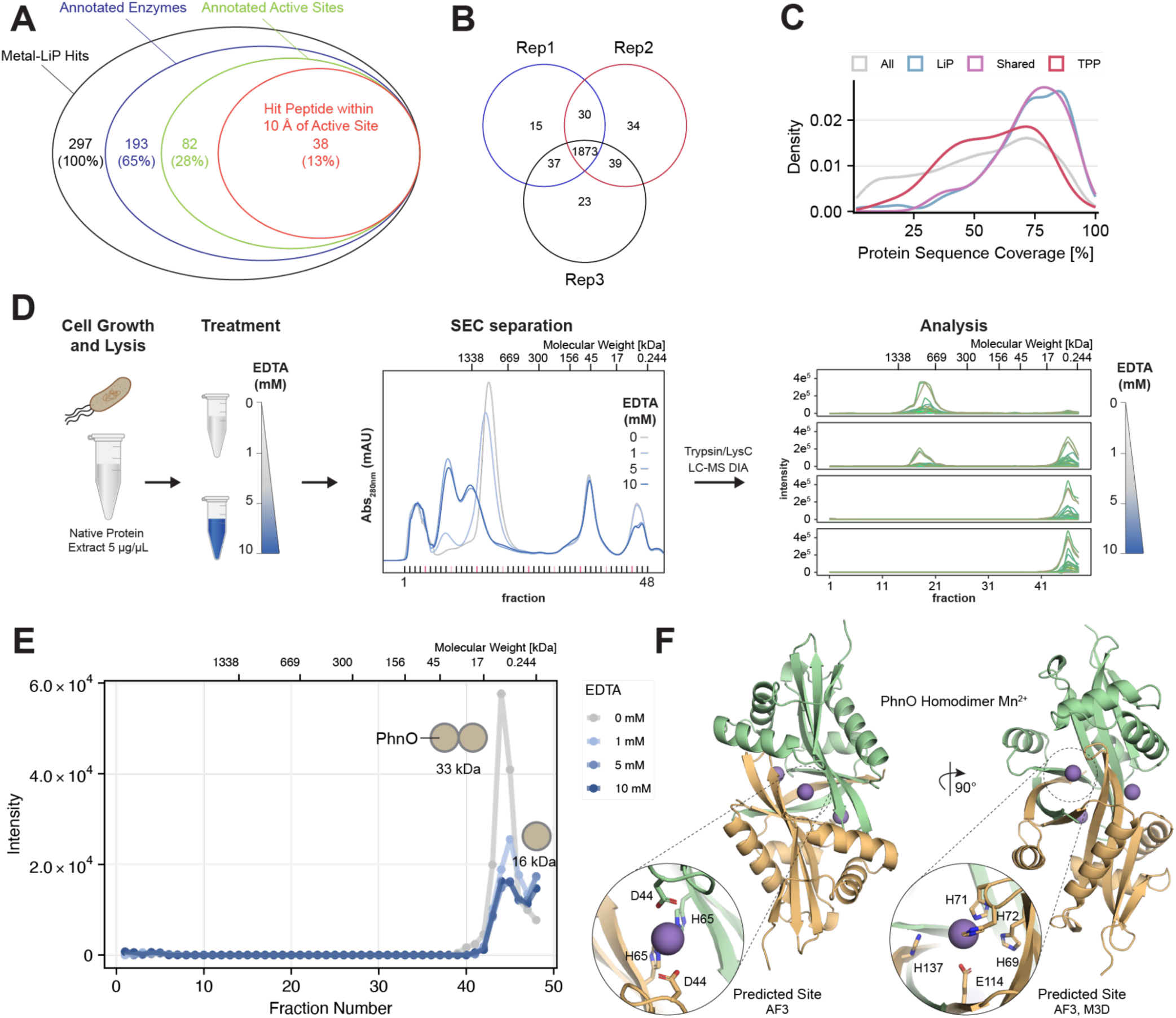
SEC-MS identifies chelator-induced changes in protein complex assembly states. **A** Subsets of Metal-LiP hit proteins annotated as enzymes (has EC number), annotated active site and a hit peptide within 10 Å of the active site. **B** Overlap of protein identifications between the three replicate TMT plexes. **C** Protein frequency by sequence coverage of all proteins detected in the LiP-MS experiment (“All”), as well as the TPP exclusive (“TPP”), LiP exclusive (“LiP”) and shared set (“Shared”) of significant proteins. **D** SEC-MS workflow for the identification of metal-regulated complex assemblies. Native lysates were treated with four EDTA concentrations (including control) and fractionated. Data analysis shows identified peptide precursor ions over all fractions. A shift indicates a complex assembly state change. **E** SEC-MS protein abundance traces for PhnO at different EDTA concentrations. The first peak is the homodimer, the second cut-off peak is the monomer. **F** AlphaFold 3 prediction of the PhnO homodimer in complex with three Mn^2+^ ions. Two metal-binding sites are created exclusively by ligands of each subunit, while an additional site is in the interface of both subunits.

## Supplementary Tables

**Supplementary Table 7:** Magnesium-free LiP buffer composition. All stock solutions were sterile filtered. The ddH_2_O was obtained from a Milli-Q^®^ system and confirmed to be metal-free by ICP-MS. The Chelex-100 was converted to the potassium form through KOH treatment as indicated by the manufacturers.

|  | Stock | Final | Comment | Product |
| --- | --- | --- | --- | --- |
| KCl | 1 M | 150 mM |  | Sigma, #409316 |
| HEPES pH 7.4 | 1 M | 100 mM | Run through a Chelex-100 column (potassium form) | Sigma, #H7523 |
| KOH | - | - | Used to adjust the HEPES Stock, prior to Chelex-100 treatment | Sigma, #P5958 |

**Supplementary Table 8:** Modified M9 medium composition. All stock solutions were sterile filtered. Stock solutions were mixed in the indicated order to prevent precipitation. The ddH_2_O was obtained from a Milli-Q^®^ system and confirmed to be metal-free by ICP-MS.

|  | Component | Stock | Final | Mixing Order | Comment | Product |
| --- | --- | --- | --- | --- | --- | --- |
| 5x base salts | Na<sub>2</sub>HPO<sub>4</sub> | 211 mM | 42.2 mM | 12 | Run through a Chelex-100 column (sodium form) | Supelco, #1.06580 |
|  | KH<sub>2</sub>PO<sub>4</sub> | 110 mM | 22 mM | 12 |  | Fluka, #60220 |
|  | NaCl | 42.8 mM | 8.6 mM | 12 |  | Supelco, #1.06404 |
|  | (NH<sub>4</sub>)<sub>2</sub>SO<sub>4</sub> | 56.7 mM | 11.3 mM | 12 |  | Fluka, #9982 |
| Carbon source | Glucose | 200 g/L | 5 g/L | 11 |  | Sigma, #49139 |
| Solubilizer | Citric acid | 1 M | 1 mM | 1 |  | Sigma, #27487 |
| Vitamines | Thiamine-HCl | 500 mg/L | 1 mg/L | 10 |  | Supelco, #47858 |
| Major divalent elements | MgCl<sub>2</sub> | 1 M | 250 µM | 8 | Trace metal grade | Sigma, #255777 |
|  | CaCl<sub>2</sub> | 100 mM | 100 µM | 7 |  | Sigma, #499609 |
| Trace elements | FeCl<sub>2</sub> | 5 mM | 5 µM | 2 |  | Sigma, #701122 |
|  | ZnCl<sub>2</sub> | 10 mM | 10 µM | 3 |  | Sigma, #229997 |
|  | CuCl<sub>2</sub> | 600 µM | 600 nM | 4 |  | Sigma, #459097 |
|  | MnCl<sub>2</sub> | 300 µM | 300 nM | 5 |  | Sigma, #203734 |
|  | CoCl<sub>2</sub> | 150 µM | 150 nM | 6 |  | Sigma, #409332 |
| Solvent | ddH<sub>2</sub>O | - | - | 9 | Metal-free (determined by ICP-MS) |  |

