## Supplementary Tables for "Systematic structural and functional analysis of metal-binding sites in native proteomes"

Supplementary Table 1: Metal-LiP Hits

| # | Gene Name | UniProt ID | LiP Positions | Chelators | Bound Metal | Metal Sites | Ribosome |
|---|---|---|---|---|---|---|---|
| 1 | iscU | P0ACD4 | 47-57; 60-78; 83-122 | DiP, DTPA, EDTA, EGTA, TETA, TPEN | copper, iron, zinc | - | - |
| 2 | rpmF | P0A7N4 | 2-10; 17-24; 43-50 | DiP, DTPA, EDTA, EGTA, TETA, TPEN | - | - | + |
| 3 | gatZ | P0C8J8 | 10-35; 66-74; 93-105; 121-161; 181-198; 257-264; 358-374 | DiP, DTPA, EDTA, EGTA, TETA, TPEN | - | Site 1: 44, 88, 122, 124, 171, 175, 257 (MoM, M3D), Site 2: 226, 228, 231 (GASS), Site 3: 189, 191, 192 (M3D), Site 4: 322, 355 (M3D) | - |
| 4 | pepT | P29745 | 78-88; 169-177; 181-187 | DiP, DTPA, EDTA, EGTA, TETA, TPEN | zinc | Site 1: 78, 140, 174, 196, 379 (M3D, Annotated) | - |
| 5 | gloA | P0AC81 | 30-38; 66-88; 105-127 | DiP, DTPA, EDTA, EGTA, TETA, TPEN | nickel | Site 1: 5, 56, 74, 122 (Annotated) | - |
| 6 | ybgI | P0AFP6 | 18-32; 38-54; 168-220 | DiP, DTPA, EDTA, EGTA, TETA, TPEN | divalent metal cation | Site 1: 63, 64, 97, 101, 108, 215, 219 (M3D, Annotated) | - |
| 7 | ycfH | P0AFQ7 | 65-82; 147-154 | DiP, DTPA, EDTA, EGTA, TETA, TPEN | cobalt, manganese, nickel | Site 1: 7, 9, 94, 130, 155, 205 (MoM, M3D, Annotated) | - |
| 8 | rpmH | P0A7P5 | 3-11 | DiP, DTPA, EDTA, EGTA, TETA, TPEN | - | - | + |
| 9 | luxS | P45578 | 91-99; 114-144; 153-163 | DiP, DTPA, EDTA, EGTA, TETA, TPEN | iron | Site 1: 54, 58, 128, 134 (MoM, M3D, Annotated) | - |

Supplementary Table 1: Metal-LiP Hits *(continued)*

| # | Gene Name | UniProt ID | LiP Positions | Chelators | Bound Metal | Metal Sites | Ribosome |
|---|---|---|---|---|---|---|---|
| 10 | hisD | P06988 | 4-18; 28-34; 61-79; 94-116; 144-174; 189-202; 206-218; 312-333; 337-345; 377-388; 390-415 | DiP, DTPA, EDTA, EGTA, TETA, TPEN | zinc, manganese | Site 1: 259, 262, 360, 419 (Annotated) | - |
| 11 | rplM | P0AA10 | 3-12; 40-61; 73-95; 124-142 | DiP, DTPA, EDTA, EGTA, TETA, TPEN | zinc | - | + |
| 12 | rpsD | P0A7V8 | 13-22; 32-44; 48-54 | DiP, DTPA, EDTA, EGTA, TETA, TPEN | - | - | + |
| 13 | argE | P23908 | 191-201; 339-364; 371-380 | DiP, DTPA, EDTA, EGTA, TETA, TPEN | zinc, cobalt | Site 1: 80, 112, 113, 144, 145, 169, 355 (M3D, Annotated) | - |
| 14 | rpmE | P0A7M9 | 9-23; 26-47; 63-70 | DiP, DTPA, EDTA, EGTA, TETA, TPEN | zinc | Site 1: 16 (Annotated) | + |
| 15 | rpsU | P68679 | 6-18; 26-33; 35-42 | DiP, DTPA, EDTA, EGTA, TETA, TPEN | - | - | + |
| 16 | rpsL | P0A7S3 | 20-31; 36-43; 67-83; 100-105 | DiP, DTPA, EDTA, EGTA, TETA, TPEN | - | - | + |
| 17 | ybeY | P0A898 | 28-44; 81-90 | DiP, DTPA, EDTA, EGTA, TETA, TPEN | zinc, nickel | Site 1: 114, 118, 124 (MoM, M3D, Annotated) | - |
| 18 | gpmI | P37689 | 11-31; 51-59; 304-317; 365-382; 389-397; 422-442; 453-476; 486-492 | DiP, DTPA, EDTA, EGTA, TETA, TPEN | manganese | Site 1: 336, 403, 407, 463 (M3D, MoM, Annotated), Site 2: 14, 64, 444, 445 (Annotated) | - |

Supplementary Table 1: Metal-LiP Hits *(continued)*

| # | Gene Name | UniProt ID | LiP Positions | Chelators | Bound Metal | Metal Sites | Ribosome |
|---|---|---|---|---|---|---|---|
| 19 | rpmI | P0A7Q1 | 53-65 | DiP, DTPA, EDTA, EGTA, TETA, TPEN | - | - | + |
| 20 | rplN | P0ADY3 | 79-98; 114-123 | DiP, DTPA, EDTA, EGTA, TETA, TPEN | - | - | + |
| 21 | pepQ | P21165 | 209-227; 392-398 | DiP, DTPA, EDTA, EGTA, TETA, TPEN | manganese | Site 1: 246, 257, 339, 346, 382, 384, 423 (Annotated, M3D) | - |
| 22 | deoB | P0A6K6 | 2-41; 180-189; 227-243; 265-271; 275-287; 302-312; 314-336; 338-361; 382-393 | DiP, DTPA, EDTA, EGTA, TETA, TPEN | manganese, cobalt, magnesium | Site 1: 10, 11, 98, 306, 311, 347, 348, 359 (M3D, Annotated, MoM, GASS), Site 2: 18, 90, 363, 380 (M3D, MoM) | - |
| 23 | rpmJ | P0A7Q6 | 19-32 | DiP, DTPA, EDTA, EGTA, TETA, TPEN | - | Site 1: 11, 14, 27, 33 (MoM, GASS, M3D) | + |
| 24 | iscR | P0AGK8 | 40-48; 70-91; 104-111 | DiP, DTPA, EDTA, EGTA, TETA, TPEN | iron | Site 1: 92, 98, 104 (Annotated) | - |
| 25 | yobA | P0AA57 | 51-64; 87-104 | DiP, DTPA, EDTA, EGTA, TETA, TPEN | copper | Site 1: 25, 27, 54, 113 (GASS, Annotated) | - |
| 26 | metE | P25665 | 315-337; 395-413; 421-441; 449-466; 468-484; 507-535; 541-549; 560-565; 611-619; 627-633; 656-668; 692-710; 732-751 | DiP, DTPA, EDTA, EGTA, TETA, TPEN | zinc | Site 1: 641, 643, 665, 725, 726 (MoM, M3D, Annotated), Site 2: 6, 321, 353 (GASS), Site 3: 146, 148, 197 (GASS) | - |
| 27 | tabA | P0AF96 | 6-18; 48-59; 83-98; 115-129 | DiP, DTPA, EDTA, EGTA, TETA, TPEN | - | Site 1: 62, 64, 69, 71, 128, 146 (M3D) | - |

Supplementary Table 1: Metal-LiP Hits *(continued)*

| # | Gene Name | UniProt ID | LiP Positions | Chelators | Bound Metal | Metal Sites | Ribosome |
|---|---|---|---|---|---|---|---|
| 28 | pgm | P36938 | 54-72; 133-156; 519-529 | DiP, DTPA, EDTA, EGTA, TETA, TPEN | magnesium | Site 1: 146, 304, 306, 308, 309 (M3D, Annotated, GASS), Site 2: 473, 474, 476, 527 (M3D) | - |
| 29 | pyrI | P0A7F3 | 18-28; 95-128 | DiP, DTPA, EDTA, EGTA, TETA, TPEN | zinc | Site 1: 109, 114, 117, 138, 141 (MoM, M3D, Annotated) | - |
| 30 | rpmA | P0A7L8 | 25-39; 45-55 | DiP, DTPA, EDTA, EGTA, TETA, TPEN | - | - | + |
| 31 | zur | P0AC51 | 4-15; 44-78 | DiP, DTPA, EDTA, EGTA, TETA, TPEN | zinc | Site 1: 152, 154, 158, 160, 162 (MoM, M3D), Site 2: 103, 106, 143, 146 (MoM, M3D), Site 3: 77, 88, 96, 98, 111 (M3D) | - |
| 32 | acnB | P36683 | 78-85; 318-331; 443-450; 779-791; 807-819 | DiP, DTPA, EDTA, EGTA, TETA, TPEN | iron | Site 1: 710, 769, 772 (Annotated) | - |
| 33 | aroG | P0AB91 | 54-61; 76-92; 174-186; 328-339 | DiP, DTPA, EDTA, EGTA, TETA, TPEN | - | Site 1: 61, 92, 268, 302, 304, 326, 328 (MoM, M3D) | - |
| 34 | rpsS | P0A7U3 | 7-17; 37-43; 70-78 | DiP, DTPA, EDTA, EGTA, TETA, TPEN | - | - | + |
| 35 | hisI | P06989 | 51-56; 76-112; 115-125 | DiP, DTPA, EDTA, EGTA, TETA, TPEN | - | Site 1: 80, 81, 82, 84, 85 (GASS, M3D), Site 2: 153, 156, 172 (M3D) | - |
| 36 | metF | P0AEZ1 | 2-7; 91-100 | DiP, DTPA, EDTA, EGTA, TETA, TPEN | - | Site 1: 28, 57, 88, 273 (M3D) | - |

Supplementary Table 1: Metal-LiP Hits *(continued)*

| # | Gene Name | UniProt ID | LiP Positions | Chelators | Bound Metal | Metal Sites | Ribosome |
|---|---|---|---|---|---|---|---|
| 37 | maeB | P76558 | 633-647 | DiP, DTPA, EDTA, EGTA, TETA, TPEN | magnesium, manganese | Site 1: 136, 137, 162 (Annotated) | - |
| 38 | metK | P0A817 | 2-18; 40-47; 61-70; 91-105; 178-186; 332-343 | DiP, DTPA, EDTA, EGTA, TETA | magnesium, potassium, manganese, cobalt | Site 1: 12, 143, 167 (GASS), Site 2: 17 (Annotated) | - |
| 39 | iscA | P0AAC8 | 13-20; 90-107 | DiP, DTPA, EDTA, EGTA, TPEN | iron | Site 1: 35, 99, 101 (Annotated) | - |
| 40 | lpxC | P0A725 | 223-237; 271-278 | DiP, DTPA, EDTA, TETA, TPEN | zinc, iron | Site 1: 62, 79, 191, 238, 239, 242, 265 (M3D, MoM, Annotated) | - |
| 41 | uspD | P0AAB8 | 131-142 | DiP, DTPA, EDTA, EGTA, TETA | - | - | - |
| 42 | trxB | P0A9P4 | 137-145 | DiP, DTPA, EDTA, EGTA, TETA | - | Site 1: 6, 107 (M3D) | - |
| 43 | ahpC | P0AE08 | 178-187 | DiP, DTPA, EDTA, EGTA, TPEN | - | - | - |
| 44 | yfeX | P76536 | 2-24; 66-77; 125-131; 142-155; 162-170; 176-181; 186-197; 200-217; 233-273 | DiP, DTPA, EDTA, EGTA, TPEN | iron | Site 1: 215 (Annotated) | - |
| 45 | bolA | P0ABE2 | 15-28 | DiP, DTPA, EDTA, TETA, TPEN | - | - | - |
| 46 | yhhW | P46852 | 83-100; 182-189 | DiP, DTPA, EDTA, EGTA, TPEN | copper, iron, zinc, cobalt | Site 1: 57, 59, 101, 103 (MoM, GASS, M3D, Annotated) | - |
| 47 | pepD | P15288 | 431-438; 473-480 | DiP, EDTA, EGTA, TETA, TPEN | zinc, cobalt | Site 1: 24, 28, 76, 115, 116, 146, 169, 457 (GASS, M3D, Annotated) | - |

Supplementary Table 1: Metal-LiP Hits *(continued)*

| # | Gene Name | UniProt ID | LiP Positions | Chelators | Bound Metal | Metal Sites | Ribosome |
|---|---|---|---|---|---|---|---|
| 48 | leuA | P09151 | 29-37; 201-223; 286-295 | DiP, EDTA, EGTA, TETA, TPEN | manganese | Site 1: 14, 202, 204, 238 (MoM, M3D, Annotated) | - |
| 49 | pepA | P68767 | 15-26; 49-72; 342-356 | DiP, DTPA, EDTA, EGTA, TETA | manganese | Site 1: 270, 275, 282, 293, 352, 354 (M3D, Annotated) | - |
| 50 | rpsK | P0A7R9 | 14-46; 58-69; 99-106 | DiP, DTPA, EGTA, TETA, TPEN | - | - | + |
| 51 | thrB | P00547 | 62-71; 206-217; 270-278 | DiP, DTPA, EDTA, EGTA, TETA | - | - | - |
| 52 | accD | P0A9Q5 | 27-36 | DiP, DTPA, EDTA, EGTA, TPEN | zinc | Site 1: 27, 30, 46, 49, 51 (MoM, M3D, Annotated) | - |
| 53 | yajD | P0AAQ2 | 22-29 | DiP, DTPA, EDTA, EGTA, TPEN | zinc | - | - |
| 54 | pgk | P0A799 | 128-146; 161-176; 249-258 | DiP, DTPA, EDTA, EGTA, TETA | - | - | - |
| 55 | queC | P77756 | 170-177; 194-206 | DTPA, EDTA, EGTA, TETA, TPEN | zinc | Site 1: 188, 197, 200, 203 (MoM, M3D, Annotated) | - |
| 56 | dksA | P0ABS1 | 105-139 | DTPA, EDTA, EGTA, TETA, TPEN | zinc | Site 1: 114, 117, 135, 138 (Annotated) | - |
| 57 | topA | P06612 | 596-604; 684-700 | DTPA, EDTA, EGTA, TETA, TPEN | magnesium, zinc, manganese, calcium | Site 1: 662, 665, 683, 689 (MoM, M3D), Site 2: 599, 602, 619, 630 (MoM, GASS, M3D), Site 3: 711, 714, 731, 736 (MoM, M3D), Site 4: 9, 111 (Annotated) | - |

Supplementary Table 1: Metal-LiP Hits *(continued)*

| # | Gene Name | UniProt ID | LiP Positions | Chelators | Bound Metal | Metal Sites | Ribosome |
|---|---|---|---|---|---|---|---|
| 58 | ileS | P00956 | 184-192; 334-348; 746-761; 831-851; 854-860; 906-923; 925-934 | DTPA, EDTA, EGTA, TETA, TPEN | zinc | Site 1: 901, 904, 906, 921, 924 (MoM, GASS, M3D, Annotated), Site 2: 177, 179, 461 (MoM), Site 3: 188, 191, 406, 409 (MoM, M3D) | - |
| 59 | trhO | P24188 | 259-298 | DTPA, EDTA, EGTA, TETA, TPEN | - | Site 1: 280, 285, 302, 303, 307, 308 (MoM, M3D), Site 2: 267, 270, 274, 292, 295 (MoM, GASS, M3D), Site 3: 124, 126, 215 (MoM) | - |
| 60 | fbaA | P0AB71 | 21-52; 55-61; 76-85; 195-212; 243-251; 253-276 | DTPA, EDTA, EGTA, TETA, TPEN | zinc | Site 1: 111, 227, 265, 289 (MoM, Annotated), Site 2: 108, 110, 142 (MoM), Site 3: 145, 175 (Annotated) | - |
| 61 | msrB | P0A746 | 41-79 | DTPA, EDTA, EGTA, TETA, TPEN | zinc, iron | Site 1: 46, 49, 95, 98 (MoM, M3D, Annotated) | - |
| 62 | copA | Q59385 | 2-19; 23-69 | DTPA, EDTA, EGTA, TETA, TPEN | copper, magnesium | Site 1: 14, 17 (Annotated), Site 2: 110, 113 (Annotated), Site 3: 720, 724 (Annotated) | - |
| 63 | metG | P00959 | 151-175; 272-283 | DTPA, EDTA, EGTA, TETA, TPEN | zinc | Site 1: 146, 149, 159, 162 (MoM, M3D, Annotated), Site 2: 22, 81, 84 (GASS), Site 3: 55, 71, 99 (GASS) | - |
| 64 | sodA | P00448 | 31-60; 62-68; 101-119; 125-138 | DTPA, EDTA, EGTA, TETA, TPEN | manganese | Site 1: 27, 31, 32, 82, 168, 172 (M3D, Annotated, MoM) | - |
| 65 | ghrB | P37666 | 300-312; 317-324 | DTPA, EDTA, EGTA, TETA, TPEN | - | - | - |
| 66 | glmS | P17169 | 2-11; 203-218 | DTPA, EDTA, EGTA, TETA, TPEN | - | Site 1: 2, 72, 98 (GASS) | - |
| 67 | argG | P0A6E4 | 87-107; 366-386; 409-417 | DTPA, EDTA, EGTA, TETA, TPEN | - | Site 1: 245, 257 (M3D) | - |

Supplementary Table 1: Metal-LiP Hits *(continued)*

| # | Gene Name | UniProt ID | LiP Positions | Chelators | Bound Metal | Metal Sites | Ribosome |
|---|---|---|---|---|---|---|---|
| 68 | rpoC | P0A8T7 | 89-96; 884-901 | DTPA, EDTA, EGTA, TETA, TPEN | zinc, magnesium | Site 1: 70, 72, 85, 88 (Annotated), Site 2: 460, 462, 464 (Annotated), Site 3: 814, 888, 895, 898 (Annotated) | - |
| 69 | recR | P0A7H6 | 62-76 | DTPA, EDTA, EGTA, TETA, TPEN | zinc | Site 1: 56, 57, 60, 65, 69, 72 (GASS, MoM, M3D) | - |
| 70 | nikR | P0A6Z6 | 38-65; 71-84 | DTPA, EDTA, EGTA, TETA, TPEN | nickel | Site 1: 62, 76, 87, 89, 95 (MoM, Annotated, M3D) | - |
| 71 | hemB | P0ACB2 | 80-101; 117-149; 303-313 | DTPA, EDTA, EGTA, TETA, TPEN | zinc, magnesium | Site 1: 118, 120, 122, 123, 130, 216 (M3D, MoM, Annotated), Site 2: 232 (Annotated) | - |
| 72 | ligA | P15042 | 407-425 | DTPA, EDTA, EGTA, TETA, TPEN | zinc, magnesium | Site 1: 408, 411, 426, 432 (Annotated) | - |
| 73 | slyD | P0A9K9 | 17-45; 62-78; 141-158 | DTPA, EDTA, EGTA, TETA, TPEN | nickel, copper, zinc, cobalt | Site 1: 167, 168, 184, 185, 193, 195 (Annotated) | - |
| 74 | phoA | P00634 | 296-303 | DTPA, EDTA, EGTA, TETA, TPEN | magnesium, zinc | Site 1: 73, 124, 175, 177, 344, 349, 353, 391, 392, 394, 434 (M3D, Annotated, MoM) | - |
| 75 | speD | P0A7F6 | 20-27; 39-68; 82-90; 116-140 | DTPA, EDTA, EGTA, TETA, TPEN | magnesium | - | - |
| 76 | ychJ | P37052 | 2-25 | DTPA, EDTA, EGTA, TETA, TPEN | - | Site 1: 99, 138, 140, 142, 149, 150 (M3D, MoM), Site 2: 5, 7, 16, 17, 39 (MoM, M3D), Site 3: 54, 118 (M3D) | - |
| 77 | rpmG | P0A7N9 | 11-25; 29-35; 38-45 | DTPA, EDTA, EGTA, TETA, TPEN | - | - | + |
| 78 | asnB | P22106 | 2-11 | DTPA, EDTA, EGTA, TETA, TPEN | - | Site 1: 30, 161, 163, 315 (M3D) | - |

Supplementary Table 1: Metal-LiP Hits *(continued)*

| # | Gene Name | UniProt ID | LiP Positions | Chelators | Bound Metal | Metal Sites | Ribosome |
|---|---|---|---|---|---|---|---|
| 79 | fusA | P0A6M8 | 84-101; 114-128; 398-408; 578-594 | DTPA, EDTA, EGTA, TETA, TPEN | - | - | - |
| 80 | dmsA | P18775 | 567-575 | DTPA, EDTA, EGTA, TETA, TPEN | iron, molybdenum | Site 1: 63, 67, 71, 104 (Annotated), Site 2: 205 (Annotated) | - |
| 81 | rpsJ | P0A7R5 | 17-31; 36-59; 71-89; 95-103 | DTPA, EDTA, EGTA, TETA, TPEN | - | - | + |
| 82 | map | P0AE18 | 21-43; 68-86 | DTPA, EDTA, EGTA, TETA, TPEN | cobalt, zinc, manganese, iron, sodium | Site 1: 97, 108, 109, 171, 202, 204, 235 (M3D, Annotated) | - |
| 83 | ppa | P0A7A9 | 36-45; 61-87; 95-105; 135-143; 153-162 | DTPA, EDTA, EGTA, TETA, TPEN | magnesium, zinc | Site 1: 66, 71, 103 (Annotated) | - |
| 84 | fur | P0A9A9 | 78-98; 131-148 | DTPA, EDTA, EGTA, TETA, TPEN | zinc, iron | Site 1: 32, 33, 71, 81, 84, 88, 90, 93, 96, 101, 133, 138, 145 (M3D, Annotated, MoM), Site 2: 87, 89, 108, 125 (GASS, M3D, Annotated) | - |
| 85 | dnaX | P06710 | 61-80 | DTPA, EDTA, EGTA, TETA, TPEN | zinc | Site 1: 64, 73, 76, 79 (Annotated) | - |
| 86 | rplJ | P0A7J3 | 21-31; 62-72 | DTPA, EDTA, EGTA, TETA, TPEN | - | - | + |
| 87 | gapA | P0A9B2 | 150-160 | DTPA, EDTA, EGTA, TETA, TPEN | - | Site 1: 150, 151, 177 (M3D) | - |
| 88 | nfo | P0A6C1 | 3-34; 41-53; 116-136; 158-171; 174-184 | DiP, DTPA, EDTA, EGTA | zinc, manganese | Site 1: 182, 229, 231 (MoM, M3D, Annotated), Site 2: 69, 107, 109, 145, 179, 216, 261 (M3D, Annotated) | - |
| 89 | erpA | P0ACC3 | 2-9; 21-32; 78-94; 104-112 | DiP, DTPA, EDTA, TPEN | iron | Site 1: 42, 106, 108 (Annotated) | - |

Supplementary Table 1: Metal-LiP Hits *(continued)*

| # | Gene Name | UniProt ID | LiP Positions | Chelators | Bound Metal | Metal Sites | Ribosome |
|---|---|---|---|---|---|---|---|
| 90 | rpsN | P0AG59 | 29-43; 76-81 | DiP, DTPA, EDTA, EGTA | - | - | + |
| 91 | glnA | P0A9C5 | 210-225; 457-469 | DiP, EGTA, TETA, TPEN | magnesium | Site 1: 132, 210, 212, 213, 221 (Annotated, GASS), Site 2: 130, 270, 272, 345, 358 (M3D, Annotated) | - |
| 92 | uspG | P39177 | 136-142 | DiP, DTPA, TETA, TPEN | - | - | - |
| 93 | tsf | P0A6P1 | 78-85; 168-193 | DTPA, EDTA, EGTA, TETA | zinc | - | - |
| 94 | carB | P00968 | 601-615 | DTPA, EDTA, EGTA, TETA | magnesium, manganese | Site 1: 932, 1039, 1041, 1050 (M3D), Site 2: 285, 299, 301 (Annotated), Site 3: 829, 841, 843 (Annotated) | - |
| 95 | adhE | P0A9Q7 | 69-84; 370-381; 464-481; 729-750 | DTPA, EDTA, EGTA, TETA | iron | Site 1: 653, 657, 723, 737, 741 (MoM, M3D, Annotated), Site 2: 561, 563 (M3D) | - |
| 96 | hemL | P23893 | 209-224; 245-265 | DTPA, EDTA, EGTA, TETA | - | - | - |
| 97 | mdh | P61889 | 251-262 | DTPA, EDTA, EGTA, TETA | - | - | - |
| 98 | fabI | P0AEK4 | 63-80; 210-218 | DTPA, EDTA, EGTA, TETA | - | - | - |
| 99 | yfgJ | P76575 | 32-45 | DTPA, EDTA, EGTA, TPEN | - | - | - |
| 100 | lpdA | P0A9P0 | 38-54 | DTPA, EDTA, EGTA, TETA | zinc | - | - |
| 101 | deoC | P0A6L0 | 38-51 | DTPA, EDTA, EGTA, TETA | - | - | - |
| 102 | gdhA | P00370 | 413-436 | DTPA, EDTA, EGTA, TETA | - | - | - |

Supplementary Table 1: Metal-LiP Hits *(continued)*

| # | Gene Name | UniProt ID | LiP Positions | Chelators | Bound Metal | Metal Sites | Ribosome |
|---|---|---|---|---|---|---|---|
| 103 | rplQ | P0AG44 | 23-30 | DTPA, EDTA, EGTA, TPEN | - | - | + |
| 104 | gmhA | P63224 | 61-69 | DTPA, EGTA, TETA, TPEN | zinc | Site 1: 61, 65, 172, 180 (Annotated) | - |
| 105 | msrC | P76270 | 100-115 | DTPA, EDTA, EGTA, TETA | - | - | - |
| 106 | yceD | P0AB28 | 67-75 | DTPA, EDTA, EGTA, TPEN | - | Site 1: 73, 76, 137, 141, 142 (MoM, M3D) | - |
| 107 | yrdA | P0A9W9 | 33-57; 72-92; 149-158; 171-184 | DTPA, EDTA, TETA, TPEN | zinc | Site 1: 48, 50, 67, 68, 79, 96, 98 (MoM, M3D, GASS) | - |
| 108 | dnaJ | P08622 | 154-171 | DTPA, EDTA, EGTA, TETA | zinc | Site 1: 144, 147, 197, 200 (Annotated), Site 2: 161, 164, 183, 186 (Annotated) | - |
| 109 | rpsG | P02359 | 37-76; 79-95; 119-131; 171-179 | DTPA, EDTA, EGTA, TETA | - | - | + |
| 110 | rapZ | P0A894 | 28-42 | DTPA, EDTA, EGTA, TETA | - | - | - |
| 111 | btuE | P06610 | 83-96 | DTPA, EDTA, EGTA, TETA | - | Site 1: 37, 71, 146 (GASS) | - |
| 112 | eno | P0A6P9 | 247-254; 271-282; 312-324; 350-357; 370-393 | DTPA, EDTA, EGTA, TETA | magnesium | Site 1: 42 (Annotated), Site 2: 246, 290, 317 (Annotated) | - |
| 113 | murA | P0A749 | 381-391 | DTPA, EDTA, EGTA, TETA | - | - | - |
| 114 | grxB | P0AC59 | 7-14; 132-145 | DTPA, EDTA, EGTA, TPEN | - | - | - |
| 115 | mrp | P0AF08 | 279-300 | DTPA, EDTA, EGTA, TETA | iron | Site 1: 283, 286, 288 (MoM), Site 2: 26, 65 (M3D) | - |
| 116 | rne | P21513 | 734-762; 832-841 | DTPA, EDTA, EGTA, TETA | magnesium, zinc | Site 1: 303, 346 (Annotated), Site 2: 404, 407 (Annotated) | - |

Supplementary Table 1: Metal-LiP Hits *(continued)*

| # | Gene Name | UniProt ID | LiP Positions | Chelators | Bound Metal | Metal Sites | Ribosome |
|---|---|---|---|---|---|---|---|
| 117 | groEL | P0A6F5 | 458-470; 519-526 | DTPA, EDTA, EGTA, TETA | magnesium | - | - |
| 118 | ybjQ | P0A8C1 | 59-70 | DTPA, EDTA, TETA, TPEN | - | - | - |
| 119 | aceA | P0A9G6 | 245-257 | DTPA, EDTA, EGTA, TETA | magnesium, manganese | Site 1: 244, 246, 249 (GASS), Site 2: 195, 196, 197, 289, 321 (M3D), Site 3: 157 (Annotated) | - |
| 120 | efeO | P0AB24 | 260-275 | DTPA, EDTA, EGTA, TPEN | - | - | - |
| 121 | rplV | P61175 | 99-110 | DTPA, EDTA, EGTA, TETA | - | - | + |
| 122 | yciF | P21362 | 5-13; 99-108 | DTPA, EDTA, EGTA, TETA | - | - | - |
| 123 | gltB | P09831 | 12-27; 541-549; 570-586; 1406-1428 | DTPA, EGTA, TETA, TPEN | iron | Site 1: 1102, 1108, 1113 (Annotated) | - |
| 124 | hcxA | P45579 | 331-351 | DTPA, EDTA, EGTA, TPEN | zinc | Site 1: 70, 75, 135 (GASS), Site 2: 173, 257, 274 (M3D, Annotated) | - |
| 125 | rbsK | P0A9J6 | 63-74 | DTPA, EDTA, EGTA, TETA | potassium, magnesium, manganese | - | - |
| 126 | glpX | P0A9C9 | 145-152 | DTPA, EDTA, EGTA, TPEN | manganese | Site 1: 236, 237, 246 (M3D), Site 2: 33, 57 (Annotated), Site 3: 85, 88, 213 (Annotated) | - |
| 127 | hisB | P06987 | 177-183 | EDTA, EGTA, TETA, TPEN | magnesium, zinc | Site 1: 93, 95, 101, 103, 104 (MoM, M3D, Annotated), Site 2: 203, 323, 324, 327 (MoM, M3D), Site 3: 229, 233, 300 (M3D), Site 4: 9, 11, 130 (Annotated) | - |
| 128 | letB | P76272 | 632-646 | DiP, TETA, TPEN | - | - | - |
| 129 | gloC | P75849 | 5-11; 98-109 | DiP, EDTA, EGTA | zinc | Site 1: 30, 56, 58, 60, 61, 83, 129, 132, 136, 151, 162, 168, 192 (MoM, M3D, Annotated, GASS) | - |
| 130 | wrbA | P0A8G6 | 156-164 | DiP, TETA, TPEN | - | - | - |

Supplementary Table 1: Metal-LiP Hits *(continued)*

| # | Gene Name | UniProt ID | LiP Positions | Chelators | Bound Metal | Metal Sites | Ribosome |
|---|---|---|---|---|---|---|---|
| 131 | yjdM | P0AFJ1 | 2-8 | DTPA, EDTA, TPEN | - | - | - |
| 132 | rplE | P62399 | 49-64; 136-145; 151-179 | DTPA, EDTA, TETA | - | - | + |
| 133 | purA | P0A7D4 | 319-332; 344-349 | DTPA, EDTA, EGTA | magnesium | Site 1: 14, 41 (Annotated) | - |
| 134 | ptsI | P08839 | 502-510 | DTPA, EGTA, TETA | magnesium | Site 1: 431, 455 (Annotated) | - |
| 135 | rplW | P0ADZ0 | 13-20; 27-33; 49-64; 88-100 | DTPA, EDTA, TETA | - | - | + |
| 136 | hisC | P06986 | 94-100 | DTPA, EDTA, EGTA | - | - | - |
| 137 | xthA | P09030 | 241-251 | DTPA, EGTA, TETA | magnesium, manganese | Site 1: 34, 258 (Annotated), Site 2: 151, 153 (Annotated) | - |
| 138 | tyrB | P04693 | 263-271 | DTPA, EGTA, TETA | - | - | - |
| 139 | rplR | P0C018 | 103-111 | DTPA, EDTA, TPEN | - | - | + |
| 140 | ycdX | P75914 | 78-88; 151-166 | DTPA, TETA, TPEN | zinc | Site 1: 7, 9, 15, 39, 40, 73, 101, 131, 166, 192, 194 (MoM, GASS, M3D, Annotated), Site 2: 49, 50 (M3D) | - |
| 141 | rpoA | P0A7Z4 | 176-182 | DTPA, EGTA, TETA | - | - | - |
| 142 | glgP | P0AC86 | 381-389 | DTPA, EGTA, TETA | - | - | - |
| 143 | gloB | P0AC84 | 121-136 | DTPA, EDTA, EGTA | zinc | Site 1: 12, 28, 53, 55, 57, 58, 110, 114, 127, 165 (M3D, MoM, GASS, Annotated) | - |
| 144 | ispB | P0AD57 | 181-194 | DTPA, TETA, TPEN | magnesium | Site 1: 242, 246, 278 (MoM), Site 2: 84, 88 (Annotated) | - |
| 145 | rpsI | P0A7X3 | 2-9; 33-41; 48-61; 69-77; 88-99 | DTPA, EDTA, TETA | - | - | + |
| 146 | thrA | P00561 | 103-125 | DTPA, EGTA, TETA | sodium | - | - |
| 147 | ppiB | P23869 | 121-135 | EDTA, EGTA, TETA | - | - | - |
| 148 | cpdB | P08331 | 569-580 | EDTA, EGTA, TPEN | divalent metal cation | Site 1: 31, 33, 76, 116, 117, 225, 257, 259 (MoM, Annotated) | - |

Supplementary Table 1: Metal-LiP Hits *(continued)*

| # | Gene Name | UniProt ID | LiP Positions | Chelators | Bound Metal | Metal Sites | Ribosome |
|---|---|---|---|---|---|---|---|
| 149 | grxD | P0AC69 | 27-46 | EDTA, TETA, TPEN | iron | Site 1: 30 (Annotated) | - |
| 150 | pta | P0A9M8 | 432-443 | EDTA, EGTA, TETA | zinc | - | - |
| 151 | yfhM | P76578 | 998-1018 | EDTA, EGTA, TPEN | - | - | - |
| 152 | rpsM | P0A7S9 | 4-27; 32-71; 79-87 | EDTA, TETA, TPEN | - | - | + |
| 153 | ahpF | P35340 | 223-233 | EDTA, EGTA, TPEN | - | - | - |
| 154 | ttcA | P76055 | 290-304 | EDTA, EGTA, TETA | iron, magnesium | Site 1: 122, 125, 213 (Annotated) | - |
| 155 | trpE | P00895 | 238-250 | EDTA, EGTA, TETA | magnesium | Site 1: 361, 498 (Annotated) | - |
| 156 | yejK | P33920 | 230-243 | EDTA, EGTA, TETA | - | Site 1: 211, 215, 218 (GASS) | - |
| 157 | glyA | P0A825 | 24-42 | EDTA, EGTA, TETA | zinc | - | - |
| 158 | add | P22333 | 14-29; 49-62; 273-302 | EDTA, EGTA, TPEN | zinc | Site 1: 12, 14, 197, 200, 221, 278 (MoM, M3D, Annotated) | - |
| 159 | pepN | P04825 | 326-337; 391-402 | EDTA, EGTA, TPEN | zinc | Site 1: 121, 264, 297, 298, 301, 319, 320, 381 (M3D, MoM, GASS, Annotated), Site 2: 705, 734 (M3D) | - |
| 160 | rplY | P68919 | 2-9; 52-68 | EGTA, TETA, TPEN | - | - | + |
| 161 | rplB | P60422 | 72-81; 134-140; 157-163; 190-199; 225-238 | EGTA, TETA, TPEN | zinc | - | + |
| 162 | rplD | P60723 | 14-21; 80-86; 108-114; 116-130; 171-180 | EGTA, TETA, TPEN | - | - | + |
| 163 | ispG | P62620 | 276-290 | DiP, TPEN | iron | Site 1: 34, 38, 44 (GASS), Site 2: 270, 273, 305, 312 (Annotated) | - |
| 164 | cmoA | P76290 | 111-119 | DiP, DTPA | - | - | - |
| 165 | rplO | P02413 | 3-13 | DiP, TPEN | - | - | + |
| 166 | aroH | P00887 | 12-24 | DTPA, EDTA | - | - | - |
| 167 | purH | P15639 | 495-504 | DTPA, TETA | - | - | - |
| 168 | lysA | P00861 | 342-359 | DTPA, TETA | - | Site 1: 137, 181, 182 (MoM) | - |

Supplementary Table 1: Metal-LiP Hits *(continued)*

| # | Gene Name | UniProt ID | LiP Positions | Chelators | Bound Metal | Metal Sites | Ribosome |
|---|---|---|---|---|---|---|---|
| 169 | dosC | P0AA89 | 435-446 | DTPA, EDTA | iron, magnesium | Site 1: 98 (Annotated), Site 2: 333, 376 (Annotated) | - |
| 170 | avtA | P09053 | 2-10 | DTPA, EGTA | - | - | - |
| 171 | atpD | P0ABB4 | 231-247 | DTPA, TETA | - | - | - |
| 172 | degP | P0C0V0 | 408-425 | DTPA, EGTA | - | - | - |
| 173 | pheS | P08312 | 170-176 | DTPA, TETA | magnesium | Site 1: 252 (Annotated) | - |
| 174 | rpsC | P0A7V3 | 54-62; 70-79; 96-107; 212-225 | DTPA, TETA | - | - | + |
| 175 | ycfP | P0A8E1 | 77-84 | DTPA, TPEN | - | - | - |
| 176 | nrdR | P0A8D0 | 42-50 | EDTA, EGTA | zinc | Site 1: 3, 6, 31, 34 (MoM, GASS, M3D) | - |
| 177 | yeaD | P39173 | 21-37 | EDTA, TETA | - | Site 1: 150, 233, 235 (GASS) | - |
| 178 | pgl | P52697 | 296-314 | EDTA, TETA | - | Site 1: 180, 233, 235 (MoM) | - |
| 179 | ispF | P62617 | 3-24; 143-156 | EDTA, TPEN | zinc, manganese | Site 1: 8, 10, 42 (MoM, M3D, Annotated) | - |
| 180 | rplX | P60624 | 5-17; 44-61 | EDTA, TETA | - | - | + |
| 181 | gltX | P04805 | 127-141 | EDTA, TETA | zinc | Site 1: 98, 99, 100, 121, 125, 127 (MoM, M3D, Annotated) | - |
| 182 | rpsO | P0ADZ4 | 2-8; 18-30; 34-48 | EDTA, TETA | - | - | + |
| 183 | argB | P0A6C8 | 94-111 | EDTA, TETA | - | - | - |
| 184 | sra | P68191 | 9-16; 22-39 | EDTA, TETA | - | - | - |
| 185 | gatB | P37188 | 27-40 | EDTA, TETA | - | - | - |
| 186 | mreB | P0A9X4 | 110-123 | EDTA, EGTA | - | - | - |
| 187 | mmuM | Q47690 | 61-81 | EDTA, TPEN | zinc | Site 1: 229, 255, 295, 296 (MoM, M3D, Annotated), Site 2: 53, 55, 97 (GASS) | - |
| 188 | acuI | P26646 | 215-222 | EDTA, EGTA | - | - | - |
| 189 | bfr | P0ABD3 | 108-117 | EDTA, EGTA | iron | Site 1: 18, 25, 51, 54, 94, 123, 127, 130 (M3D, Annotated), Site 2: 46, 50 (Annotated), Site 3: 52 (Annotated) | - |
| 190 | glsA1 | P77454 | 48-61 | EGTA, TETA | - | - | - |
| 191 | gcvR | P0A9I3 | 166-190 | EGTA, TETA | - | - | - |
| 192 | yciK | P31808 | 191-198 | EGTA, TETA | - | - | - |
| 193 | maoP | P0ADN2 | 64-76 | EGTA, TETA | - | - | - |
| 194 | cfa | P0A9H7 | 5-18 | EGTA, TETA | - | Site 1: 90, 91, 94 (MoM) | - |

Supplementary Table 1: Metal-LiP Hits *(continued)*

| # | Gene Name | UniProt ID | LiP Positions | Chelators | Bound Metal | Metal Sites | Ribosome |
|---|---|---|---|---|---|---|---|
| 195 | lon | P0A9M0 | 691-698 | EGTA, TETA | - | - | - |
| 196 | ilvB | P08142 | 83-99 | EGTA, TETA | magnesium | Site 1: 444, 471 (Annotated) | - |
| 197 | ybcJ | P0AAS7 | 20-31 | EGTA, TPEN | - | - | - |
| 198 | amyA | P26612 | 92-106 | EGTA, TETA | calcium, sodium | Site 1: 104, 198, 239 (Annotated) | - |
| 199 | cysK | P0ABK5 | 56-84; 194-226 | EGTA, TETA | - | - | - |
| 200 | rpmB | P0A7M2 | 29-37; 64-72 | EGTA, TPEN | - | - | + |
| 201 | yahK | P75691 | 143-168; 193-211 | EGTA, TETA | zinc | Site 1: 91, 93, 94, 96, 99, 104, 107, 109 (GASS, MoM, M3D, Annotated), Site 2: 40, 62, 63, 88, 158, 339 (MoM, GASS, M3D, Annotated) | - |
| 202 | carA | P0A6F1 | 186-202 | EGTA, TETA | - | Site 1: 290, 291, 310, 312, 338, 353 (MoM, M3D), Site 2: 54, 115, 119 (GASS) | - |
| 203 | rplC | P60438 | 14-26; 47-53; 60-70; 84-93; 97-112 | TETA, TPEN | - | - | + |
| 204 | rplK | P0A7J7 | 73-80; 104-113; 135-142 | TETA, TPEN | - | - | + |
| 205 | speA | P21170 | 59-69; 383-396; 619-625; 630-640; 653-658 | TETA, TPEN | magnesium | - | - |
| 206 | fruB | P69811 | 101-111 | TETA, TPEN | - | - | - |
| 207 | bcp | P0AE52 | 41-61 | TETA, TPEN | - | - | - |
| 208 | selO | P77649 | 333-349 | DiP | magnesium | Site 1: 247, 256 (Annotated) | - |
| 209 | rpmD | P0AG51 | 36-45 | DiP | - | - | + |
| 210 | radD | P33919 | 433-446 | DTPA | zinc | Site 1: 384, 387, 411, 425 (Annotated) | - |
| 211 | ybiI | P41039 | 59-67 | DTPA | zinc | - | - |
| 212 | guaC | P60560 | 178-186 | EDTA | potassium | - | - |
| 213 | ybeL | P0AAT9 | 103-122 | EDTA | - | - | - |
| 214 | alaS | P00957 | 694-713 | EDTA | zinc | Site 1: 564, 568, 571, 664, 666, 668, 670 (MoM, M3D, Annotated), Site 2: 75, 87, 88 (MoM), Site 3: 188, 191, 207 (GASS), Site 4: 581, 600 (M3D) | - |
| 215 | pnp | P05055 | 611-623 | EDTA | magnesium, manganese | Site 1: 486, 492 (Annotated) | - |
| 216 | rpsA | P0AG67 | 91-100 | EDTA | - | - | + |
| 217 | mukF | P60293 | 74-83 | EGTA | calcium | - | - |
| 218 | ftsH | P0AAI3 | 352-370 | EGTA | zinc | Site 1: 414, 418, 492 (Annotated) | - |
| 219 | grpE | P09372 | 8-35 | EGTA | - | - | - |
| 220 | aroB | P07639 | 177-188 | EGTA | zinc, cobalt | Site 1: 136, 184, 247, 264, 268 (M3D, Annotated) | - |
| 221 | ygiW | P0ADU5 | 54-64; 71-93; 103-113 | TETA | - | - | - |

Supplementary Table 1: Metal-LiP Hits *(continued)*

| # | Gene Name | UniProt ID | LiP Positions | Chelators | Bound Metal | Metal Sites | Ribosome |
|---|---|---|---|---|---|---|---|
| 222 | speE | P09158 | 30-38 | TETA | - | Site 1: 64, 65, 69, 88, 90, 91, 158 (MoM, M3D) | - |
| 223 | asnS | P0A8M0 | 115-122; 326-343; 425-440 | TETA | - | - | - |
| 224 | rpsT | P0A7U7 | 50-61; 77-85 | TETA | - | - | + |
| 225 | rplA | P0A7L0 | 38-53; 63-71; 142-150 | TETA | - | - | + |
| 226 | pdeK | P37649 | 526-535 | TETA | - | - | - |
| 227 | rffH | P61887 | 143-152 | TETA | magnesium | Site 1: 108, 223 (Annotated) | - |
| 228 | thiI | P77718 | 303-316 | TETA | iron | - | - |
| 229 | pepP | P15034 | 81-92 | TETA | manganese | Site 1: 230, 261, 272, 274, 355, 382, 384, 407 (M3D, Annotated) | - |
| 230 | rpsR | P0A7T7 | 14-24 | TETA | - | - | + |
| 231 | purD | P15640 | 115-136; 386-399 | TETA | magnesium, manganese | Site 1: 286, 288 (Annotated) | - |
| 232 | clpB | P63284 | 668-681; 739-756 | TETA | - | - | - |
| 233 | aspS | P21889 | 40-58 | TETA | - | Site 1: 433, 448, 475 (M3D) | - |
| 234 | icd | P08200 | 188-207 | TETA | magnesium, manganese | Site 1: 307, 311 (Annotated) | - |
| 235 | rpsE | P0A7W1 | 46-62; 112-138; 160-167 | TETA | - | - | + |
| 236 | pyrB | P0A786 | 9-30 | TETA | - | - | - |
| 237 | ycjY | P76049 | 158-167 | TETA | - | - | - |
| 238 | guaA | P04079 | 174-188; 426-437 | TETA | - | - | - |
| 239 | can | P61517 | 174-189 | TETA | zinc | Site 1: 42, 44, 98, 101, 147 (MoM, M3D, Annotated, GASS) | - |
| 240 | pldB | P07000 | 31-38 | TETA | - | Site 1: 52, 53 (M3D) | - |
| 241 | katE | P21179 | 541-560 | TETA | iron | Site 1: 10, 12, 33 (MoM), Site 2: 415 (Annotated) | - |
| 242 | rpsH | P0A7W7 | 42-64 | TETA | - | - | + |
| 243 | aceE | P0AFG8 | 128-151; 249-264; 612-621 | TETA | magnesium | Site 1: 107, 143, 408 (MoM), Site 2: 449, 453 (M3D), Site 3: 231, 261, 263 (Annotated) | - |
| 244 | aceB | P08997 | 158-164; 456-466 | TETA | - | - | - |
| 245 | trmJ | P0AE01 | 7-16 | TETA | - | - | - |
| 246 | rpsP | P0A7T3 | 13-25 | TETA | - | - | + |
| 247 | pyrG | P0A7E5 | 105-124 | TETA | magnesium | Site 1: 72, 140 (Annotated) | - |
| 248 | gpt | P0A9M5 | 76-101 | TETA | magnesium | Site 1: 89 (Annotated) | - |
| 249 | rplF | P0AG55 | 110-134; 163-170 | TETA | - | - | + |
| 250 | yahJ | P77554 | 135-149 | TETA | - | Site 1: 121, 123, 125, 182, 184, 186, 213, 273, 275, 308, 362 (MoM, GASS, M3D) | - |
| 251 | manA | P00946 | 345-354 | TETA | zinc | Site 1: 97, 99, 134, 249, 255, 264 (M3D, Annotated) | - |
| 252 | atpC | P0A6E6 | 100-124 | TETA | - | - | - |
| 253 | purC | P0A7D7 | 164-177 | TETA | - | - | - |
| 254 | proS | P16659 | 96-119 | TETA | - | - | - |
| 255 | metAS | P07623 | 164-181 | TETA | - | - | - |
| 256 | gyrA | P0AES4 | 77-91 | TETA | - | - | - |
| 257 | puuE | P50457 | 123-140 | TETA | - | - | - |
| 258 | yhhX | P46853 | 27-36 | TETA | - | - | - |
| 259 | fre | P0AEN1 | 84-99 | TETA | iron | - | - |
| 260 | bipA | P0DTT0 | 510-529 | TETA | - | - | - |

Supplementary Table 1: Metal-LiP Hits *(continued)*

| # | Gene Name | UniProt ID | LiP Positions | Chelators | Bound Metal | Metal Sites | Ribosome |
|---|---|---|---|---|---|---|---|
| 261 | rlmI | P75876 | 18-29 | TETA | - | Site 1: 181, 195, 197, 198 (M3D) | - |
| 262 | gstA | P0A9D2 | 132-150 | TETA | - | - | - |
| 263 | glf | P37747 | 33-52 | TETA | - | - | - |
| 264 | aas | P31119 | 709-719 | TETA | - | - | - |
| 265 | folP | P0AC13 | 90-104 | TETA | magnesium | Site 1: 22 (Annotated) | - |
| 266 | suhB | P0ADG4 | 95-106 | TETA | magnesium, lithium | Site 1: 67, 84, 86 (Annotated) | - |
| 267 | lysC | P08660 | 423-442 | TETA | - | - | - |
| 268 | ndk | P0A763 | 86-104 | TETA | magnesium | - | - |
| 269 | rsfS | P0AAT6 | 53-60 | TETA | - | - | - |
| 270 | cbpM | P63264 | 59-74 | TETA | - | - | - |
| 271 | cysS | P21888 | 114-131 | TETA | zinc | Site 1: 28, 209, 224, 234, 235, 238 (MoM, M3D, Annotated), Site 2: 37, 40 (M3D) | - |
| 272 | yaeH | P62768 | 56-66 | TETA | - | - | - |
| 273 | gcvP | P33195 | 384-398 | TETA | - | - | - |
| 274 | tpx | P0A862 | 82-93 | TETA | - | - | - |
| 275 | tas | P0A9T4 | 189-200; 283-297 | TETA | - | Site 1: 191, 194, 196 (MoM), Site 2: 4, 12, 229 (GASS) | - |
| 276 | nusA | P0AFF6 | 81-104 | TETA | - | - | - |
| 277 | rnc | P0A7Y0 | 192-208 | TETA | magnesium | Site 1: 41, 114, 117 (Annotated) | - |
| 278 | trpGD | P00904 | 159-181 | TETA | magnesium | - | - |
| 279 | dsbC | P0AEG6 | 108-123 | TETA | - | Site 1: 118, 121, 125 (MoM) | - |
| 280 | katG | P13029 | 206-214 | TETA | iron | Site 1: 267 (Annotated) | - |
| 281 | pfkA | P0A796 | 268-274 | TETA | magnesium | Site 1: 104 (Annotated) | - |
| 282 | ade | P31441 | 516-538 | TETA | manganese, iron | Site 1: 90, 92, 118, 120, 121, 185, 214, 235, 236, 284, 473, 474 (MoM, M3D), Site 2: 415, 550, 554 (GASS) | - |
| 283 | dhaL | P76014 | 131-142 | TETA | magnesium | Site 1: 30, 35, 37 (Annotated) | - |
| 284 | kgtP | P0AEX3 | 2-9 | TETA | - | - | - |
| 285 | rplI | P0A7R1 | 12-22 | TPEN | - | - | + |
| 286 | yghU | Q46845 | 172-178 | TPEN | - | Site 1: 43, 76, 273 (MoM) | - |
| 287 | ubiD | P0AAB4 | 174-180 | TPEN | manganese | Site 1: 160, 320, 322 (GASS), Site 2: 175, 241 (Annotated) | - |
| 288 | frsA | P04335 | 274-281 | TPEN | - | Site 1: 139, 204, 319 (GASS), Site 2: 172, 249, 251 (GASS) | - |
| 289 | miaB | P0AEI1 | 160-170 | TPEN | iron | Site 1: 12, 49, 83 (Annotated), Site 2: 157, 161, 164 (Annotated) | - |
| 290 | degQ | P39099 | 266-272 | TPEN | - | - | - |
| 291 | feoC | P64638 | 49-62 | TPEN | iron | Site 1: 56, 61, 64, 70 (Annotated) | - |
| 292 | fruK | P0AEW9 | 179-189 | TPEN | magnesium | - | - |
| 293 | tsaD | P05852 | 35-49 | TPEN | iron, magnesium | Site 1: 111, 113, 115, 139, 300 (MoM, M3D, Annotated, GASS) | - |
| 294 | ycgL | P0AB43 | 38-44 | TPEN | - | - | - |
| 295 | minD | P0AEZ3 | 36-44 | TPEN | - | - | - |
| 296 | hrpA | P43329 | 148-157 | TPEN | - | - | - |
| 297 | yccX | P0AB65 | 29-37 | TPEN | - | - | - |

Supplementary Table 2: E. coli annotated metal-binding proteome

| # | Gene Name | UniProt ID | Ligand Name | Ligand Identifier | Ligand Position | Source |
|---|---|---|---|---|---|---|
| 1 | ndh | P00393 | copper(2+), copper(1+) | - | - | catalytic activity & go term, catalytic activity & go term |
| 2 | sodA | P00448 | manganese(2+) | 1(Mn(2+)) | 27, 82, 168, 172 | Keyword & binding & cofactor & go term |
| 3 | thrA | P00561 | sodium(1+) | 1(Na(+)) | 606, 609, 611, 613 | Keyword & binding & cofactor & go term |
| 4 | metL | P00562 | sodium(1+) | 1(Na(+)) | 603, 605, 607 | Keyword & binding & cofactor & go term |
| 5 | phoA | P00634 | magnesium(2+); zinc(2+); zinc(2+) | 1(Mg(2+)); 1(Zn(2+)); 2(Zn(2+)) | 73, 175, 177, 344; 349, 353, 434; 73, 391, 392 | Keyword & binding & cofactor & go term; Keyword & binding & cofactor & go term; Keyword & binding & cofactor & go term |
| 6 | lacZ | P00722 | magnesium(2+); magnesium(2+); manganese(2+); sodium(1+) | 1(Mg(2+)); 2(Mg(2+)); -; 1(Na(+)) | 417, 419, 462; 598; -; 202, 602, 605 | Keyword & binding & cofactor & go term; Keyword & binding & cofactor & go term; Keyword & cofactor & go term; Keyword & binding & cofactor & go term |
| 7 | ppc | P00864 | magnesium(2+) | - | - | Keyword & cofactor & go term |
| 8 | aroF | P00888 | divalent metal cation | - | - | cofactor |
| 9 | ilvI | P00893 | magnesium(2+) | 1(Mg(2+)) | 448, 475 | Keyword & binding & cofactor & go term |
| 10 | trpE | P00895 | magnesium(2+) | 1(Mg(2+)) | 361, 498 | Keyword & binding & cofactor & go term |
| 11 | trpGD | P00904 | magnesium(2+) | - | - | go term |
| 12 | xylA | P00944 | magnesium(2+); magnesium(2+) | 1(Mg(2+)); 2(Mg(2+)) | 232, 268, 296, 339; 268, 271, 307, 309 | Keyword & binding & cofactor & go term; Keyword & binding & cofactor & go term |
| 13 | manA | P00946 | zinc(2+) | 1(Zn(2+)) | 97, 99, 134, 255 | Keyword & binding & cofactor & go term |
| 14 | ileS | P00956 | zinc(2+) | 1(Zn(2+)) | 901, 904, 921, 924 | Keyword & binding & cofactor & go term |
| 15 | alaS | P00957 | zinc(2+) | 1(Zn(2+)) | 564, 568, 666, 670 | Keyword & binding & cofactor & go term |
| 16 | metG | P00959 | zinc(2+) | 1(Zn(2+)) | 146, 149, 159, 162 | Keyword & binding & cofactor & go term |

Supplementary Table 2: E. coli annotated metal-binding proteome *(continued)*

| # | Gene Name | UniProt ID | Ligand Name | Ligand Identifier | Ligand Position | Source |
|---|---|---|---|---|---|---|
| 17 | carB | P00968 | magnesium(2+); magnesium(2+); magnesium(2+); magnesium(2+); manganese(2+); manganese(2+); manganese(2+); manganese(2+) | 1(Mg(2+)); 2(Mg(2+)); 3(Mg(2+)); 4(Mg(2+)); 1(Mn(2+)); 2(Mn(2+)); 3(Mn(2+)); 4(Mn(2+)) | 285, 299; 299, 301; 829, 841; 841, 843; 285, 299; 299, 301; 829, 841; 841, 843 | Keyword & binding & cofactor & go term; Keyword & binding & cofactor & go term; Keyword & binding & cofactor & go term; Keyword & binding & cofactor & go term; Keyword & binding & cofactor & go term; Keyword & binding & cofactor & go term; Keyword & binding & cofactor & go term; Keyword & binding & cofactor & go term |
| 18 | melB | P02921 | sodium(1+), lithium(1+) | - | - | catalytic activity & go term, catalytic activity |
| 19 | tolC | P02930 | enterobactin(1-) | - | - | go term |
| 20 | dnaQ | P03007 | magnesium(2+), manganese(2+); magnesium(2+), manganese(2+) | 1(a divalent metal cation); 2(a divalent metal cation) | 12, 14, 167; 12 | Keyword & binding & cofactor & go term, Keyword & binding & cofactor & go term; Keyword & binding & cofactor & go term, Keyword & binding & cofactor & go term |
| 21 | kefC | P03819 | potassium(1+) | - | - | Keyword & go term |
| 22 | kdpA | P03959 | potassium(1+) | - | - | Keyword & go term |
| 23 | kdpB | P03960 | magnesium(2+); potassium(1+) | 1(Mg(2+)); - | 518, 522; - | Keyword & binding & go term; Keyword & catalytic activity & go term |
| 24 | kdpC | P03961 | potassium(1+) | - | - | Keyword & go term |
| 25 | argI | P04391 | zinc(2+) | 1(Zn(2+)) | 274 | Keyword & binding & go term |
| 26 | gshB | P04425 | magnesium(2+); manganese(2+) | 1(Mg(2+)); - | 281, 283; - | Keyword & binding & cofactor & go term; Keyword & cofactor & go term |
| 27 | gltX | P04805 | zinc(2+) | 1(Zn(2+)) | 98, 100, 125, 127 | Keyword & binding & cofactor & go term |
| 28 | pepN | P04825 | zinc(2+) | 1(Zn(2+)) | 297, 301, 320 | Keyword & binding & cofactor & go term |
| 29 | kdsB | P04951 | magnesium(2+) | - | - | Keyword & cofactor & go term |
| 30 | sbcB | P04995 | magnesium(2+); magnesium(2+); zinc cation | 1(Mg(2+)); 2(Mg(2+)); - | 15; 17, 186; - | Keyword & binding & cofactor & go term; Keyword & binding & cofactor & go term; Keyword & go term |
| 31 | pyrC | P05020 | zinc(2+); zinc(2+) | 1(Zn(2+)); 2(Zn(2+)) | 17, 19, 103, 251; 103, 140, 178 | Keyword & binding & cofactor & go term; Keyword & binding & cofactor & go term |
| 32 | pabB | P05041 | magnesium(2+) | - | - | Keyword & cofactor & go term |

Supplementary Table 2: E. coli annotated metal-binding proteome *(continued)*

| # | Gene Name | UniProt ID | Ligand Name | Ligand Identifier | Ligand Position | Source |
|---|---|---|---|---|---|---|
| 33 | alkB | P05050 | iron(2+) | 1(Fe cation) | 131, 133, 187 | Keyword & binding & cofactor & go term |
| 34 | pnp | P05055 | magnesium(2+); manganese(2+) | 1(Mg(2+)); - | 486, 492; - | Keyword & binding & cofactor & go term; Keyword & cofactor & go term |
| 35 | tag | P05100 | zinc(2+) | 1(Zn(2+)) | 4, 17, 175, 179 | Keyword & binding & go term |
| 36 | ptrA | P05458 | magnesium cation; zinc(2+) | -; 1(Zn(2+)) | -; 88, 92, 169 | Keyword & go term; Keyword & binding & cofactor & go term |
| 37 | mutM | P05523 | zinc(2+) | - | - | Keyword & cofactor & go term |
| 38 | ilvD | P05791 | di-mu-sulfido-diiron; iron cation, Fe4S4 iron-sulfur cluster; magnesium(2+) | 1([2Fe-2S] cluster); -; 1(Mg(2+)) | 122, 195; -; 81, 123, 124, 491 | Keyword & binding & cofactor & go term; Keyword & go term, go term; Keyword & binding & cofactor & go term |
| 39 | ilvC | P05793 | magnesium(2+); magnesium(2+) | 1(Mg(2+)); 2(Mg(2+)) | 217, 221; 217, 389, 393 | Keyword & binding & cofactor & go term; Keyword & binding & cofactor & go term |
| 40 | fepA | P05825 | iron cation, siderophore, ferrienterobactin(3-), enterobactin(1-) | - | - | Keyword, go term, go term, go term |
| 41 | ttdA | P05847 | iron cation; tetra-mu3-sulfido-tetrairon | -; 1(iron-sulfur cluster) | -; 71, 190, 277 | Keyword & go term; Keyword & binding & cofactor & go term |
| 42 | tsaD | P05852 | iron(2+); magnesium(2+) | 1(Fe cation); - | 111, 115, 300; - | Keyword & binding & cofactor & go term; Keyword & go term |
| 43 | btuB | P06129 | R-cob(III)alamin; calcium(2+); calcium(2+); cyanocob(III)alamin | -; 1(Ca(2+)); 2(Ca(2+)); 1(cyanocob(III)alamin) | -; 199, 211, 213, 215, 250; 213, 215, 249, 250, 261; 83, 85, 92, 110, 111, 251, 309, 517, 551 | go term; Keyword & binding & go term; Keyword & binding & go term; binding |
| 44 | ada | P06134 | zinc(2+) | 1(Zn(2+)) | 38, 42, 69, 72 | Keyword & binding & cofactor & go term |
| 45 | btuC | P06609 | R-cob(III)alamin | - | - | go term |
| 46 | btuD | P06611 | R-cob(III)alamin | - | - | catalytic activity & go term |
| 47 | topA | P06612 | magnesium(2+); zinc cation, manganese(2+), calcium(2+) | 1(Mg(2+)); - | 9, 111; - | Keyword & binding & go term; Keyword & go term, Keyword & cofactor & go term, Keyword & cofactor & go term |
| 48 | dnaX | P06710 | zinc(2+) | 1(Zn(2+)) | 64, 73, 76, 79 | Keyword & binding & go term |
| 49 | melA | P06720 | manganese(2+) | 1(Mn(2+)) | 173, 203 | Keyword & binding & cofactor & go term |
| 50 | argF | P06960 | zinc cation | - | - | Keyword & go term |

Supplementary Table 2: E. coli annotated metal-binding proteome *(continued)*

| # | Gene Name | UniProt ID | Ligand Name | Ligand Identifier | Ligand Position | Source |
|---|---|---|---|---|---|---|
| 51 | cca | P06961 | magnesium(2+); nickel(2+) | 1(Mg(2+)); - | 21, 23; - | Keyword & binding & cofactor & go term; Keyword & cofactor & go term |
| 52 | dut | P06968 | magnesium(2+) | - | - | Keyword & cofactor & go term |
| 53 | fhuA | P06971 | ferrichrome; iron cation, siderophore | 1(ferrichrome); - | 114, 133, 148, 149, 277, 278, 279, 346, 347, 348, 424, 735; - | binding; Keyword & go term, go term |
| 54 | fhuB | P06972 | iron cation, siderophore | - | - | Keyword, go term |
| 55 | hisB | P06987 | magnesium(2+); zinc(2+) | 1(Mg(2+)); 1(Zn(2+)) | 9, 11, 130; 93, 95, 101, 103 | Keyword & binding & cofactor & go term; Keyword & binding & cofactor & go term |
| 56 | hisD | P06988 | manganese(2+); zinc(2+) | -; 1(Zn(2+)) | -; 259, 262, 360, 419 | Keyword & go term; Keyword & binding & cofactor & go term |
| 57 | ompC | P06996 | magnesium(2+) | 1(Mg(2+)) | 340, 342, 355 | Keyword & binding & go term |
| 58 | pfkB | P06999 | magnesium(2+); potassium(1+) | 1(Mg(2+)); 1(K(+)) | 190; 250, 252, 286, 289, 291, 293 | Keyword & binding & cofactor & go term; Keyword & binding & go term |
| 59 | poxB | P07003 | magnesium(2+) | 1(Mg(2+)) | 433, 460, 462 | Keyword & binding & cofactor & go term |
| 60 | sdhB | P07014 | di-mu-sulfido-diiron; iron cation, Fe3S4 iron-sulfur cluster; tetra-mu3-sulfido-tetrairon; tri-mu-sulfido-mu3-sulfido-triiron(0) | 1([2Fe-2S] cluster); -; 1([4Fe-4S] cluster); 1([3Fe-4S] cluster) | 55, 60, 75; -; 149, 152, 155, 216; 159, 206, 212 | Keyword & binding & cofactor & go term; Keyword & go term, go term; Keyword & binding & cofactor & go term; Keyword & binding & cofactor & go term |
| 61 | ushA | P07024 | zinc(2+); zinc(2+) | 1(Zn(2+)); 2(Zn(2+)) | 41, 43, 84, 254; 84, 116, 217, 252 | Keyword & binding & cofactor & go term; Keyword & binding & cofactor & go term |
| 62 | putP | P07117 | sodium(1+) | - | - | Keyword & catalytic activity & go term |
| 63 | valS | P07118 | magnesium(2+) | - | - | go term |
| 64 | pheT | P07395 | magnesium(2+) | 1(Mg(2+)) | 454, 460, 463, 464 | Keyword & binding & cofactor & go term |
| 65 | aroB | P07639 | cobalt(2+); zinc(2+) | -; 1(Zn(2+)) | -; 184, 247, 264 | Keyword & cofactor & go term; Keyword & binding & cofactor & go term |

Supplementary Table 2: E. coli annotated metal-binding proteome *(continued)*

| # | Gene Name | UniProt ID | Ligand Name | Ligand Identifier | Ligand Position | Source |
|---|---|---|---|---|---|---|
| 66 | fdhF | P07658 | Mo(=O)-bis(molybdopterin guanine dinucleotide)(4-); iron cation, molybdenum cation; tetra-mu3-sulfido-tetrairon | 1(Mo-bis(molybdopterin guanine dinucleotide)); -; 1([4Fe-4S] cluster) | 44, 110, 111, 112, 140, 173, 174, 175, 176, 177, 178, 179, 180, 201, 202, 203, 204, 221, 222, 223, 297, 301, 335, 402, 403, 404, 405, 406, 407, 408, 409, 410, 428, 429, 445, 478, 579, 580, 581, 582, 583, 584, 585, 586, 587, 588, 654, 655, 661, 662, 678, 679; -; 8, 10, 11, 15, 42 | binding & cofactor; Keyword & go term, Keyword & go term; Keyword & binding & cofactor & go term |
| 67 | fhuC | P07821 | iron(3+), siderophore, iron(III) hydroxamate | - | - | Keyword & go term, go term, go term |
| 68 | fhuD | P07822 | coprogen; iron cation | 1(Fe(III)-coprogen); - | 68, 84, 103, 106, 124, 217, 273, 274, 275; - | binding; Keyword |
| 69 | ddlB | P07862 | magnesium(2+); magnesium(2+); manganese(2+) | 1(Mg(2+)); 2(Mg(2+)); - | 257, 270; 270, 272; - | Keyword & binding & cofactor & go term; Keyword & binding & cofactor & go term; Keyword & cofactor & go term |
| 70 | tdh | P07913 | cobalt(2+), cadmium(2+), iron(2+), manganese(2+); zinc(2+); zinc(2+) | -; 1(Zn(2+)); 2(Zn(2+)) | -; 38, 63, 64; 93, 96, 99, 107 | Keyword & cofactor & go term, Keyword & cofactor & go term, Keyword & go term, Keyword & go term; Keyword & binding & cofactor & go term; Keyword & binding & cofactor & go term |
| 71 | ilvB | P08142 | magnesium(2+) | 1(Mg(2+)) | 444, 471 | Keyword & binding & cofactor & go term |
| 72 | folC | P08192 | magnesium(2+); magnesium(2+) | 1(Mg(2+)); 2(Mg(2+)) | 83, 146; 173 | Keyword & binding & cofactor & go term; Keyword & binding & cofactor & go term |
| 73 | icd | P08200 | magnesium(2+); manganese(2+) | 1(Mg(2+)); 1(Mn(2+)) | 307; 307, 311 | Keyword & binding & cofactor & go term; Keyword & binding & cofactor & go term |
| 74 | nirB | P08201 | di-mu-sulfido-diiron; iron cation; siroheme(8-); tetra-mu3-sulfido-tetrairon | 1([2Fe-2S] cluster); -; 1(siroheme); 1([4Fe-4S] cluster) | 425, 427, 459, 462; -; 685; 641, 647, 681, 685 | Keyword & binding & cofactor & go term; Keyword & go term; Keyword & binding & cofactor & go term; Keyword & binding & cofactor & go term |
| 75 | araA | P08202 | manganese(2+) | 1(Mn(2+)) | 306, 333, 350, 450 | Keyword & binding & cofactor & go term |

Supplementary Table 2: E. coli annotated metal-binding proteome *(continued)*

| # | Gene Name | UniProt ID | Ligand Name | Ligand Identifier | Ligand Position | Source |
|---|---|---|---|---|---|---|
| 76 | araD | P08203 | cobalt cation, manganese cation; zinc(2+) | -; 1(Zn(2+)) | -; 76, 95, 97, 171 | Keyword & go term, Keyword & go term; Keyword & binding & cofactor & go term |
| 77 | pheS | P08312 | magnesium(2+) | 1(Mg(2+)) | 252 | Keyword & binding & cofactor & go term |
| 78 | cpdB | P08331 | divalent metal cation; divalent metal cation | 1(a divalent metal cation); 2(a divalent metal cation) | 31, 33, 76, 259; 76, 116, 225, 257 | Keyword & binding & cofactor & go term; Keyword & binding & cofactor & go term |
| 79 | mutT | P08337 | magnesium(2+); manganese(2+) | 1(Mg(2+)); - | 37, 57; - | Keyword & binding & cofactor & go term; Keyword & go term |
| 80 | recB | P08394 | magnesium(2+) | 1(Mg(2+)) | 956, 1067, 1080, 1081 | Keyword & binding & cofactor & go term |
| 81 | dnaJ | P08622 | zinc(2+); zinc(2+) | 1(Zn(2+)); 2(Zn(2+)) | 144, 147, 197, 200; 161, 164, 183, 186 | Keyword & binding & cofactor & go term; Keyword & binding & cofactor & go term |
| 82 | ptsI | P08839 | magnesium(2+) | 1(Mg(2+)) | 431, 455 | Keyword & binding & cofactor & go term |
| 83 | purK | P09029 | metal cation | - | - | go term |
| 84 | xthA | P09030 | magnesium(2+); magnesium(2+); manganese(2+) | 1(Mg(2+)); 2(Mg(2+)); - | 34, 258; 151, 153; - | Keyword & binding & cofactor & go term; Keyword & binding & cofactor & go term; Keyword & cofactor & go term |
| 85 | galT | P09148 | iron(2+); zinc(2+) | 1(Fe cation); 1(Zn(2+)) | 182, 281, 296, 298; 52, 55, 115, 164 | Keyword & binding & go term; Keyword & binding & cofactor & go term |
| 86 | leuA | P09151 | manganese(2+) | 1(Mn(2+)) | 14, 202, 204, 238 | Keyword & binding & cofactor & go term |
| 87 | narG | P09152 | Mo(=O)-bis(molybdopterin guanine dinucleotide)(4-); iron cation, molybdenum cation; tetra-mu3-sulfido-tetrairon | 1(Mo-bis(molybdopterin guanine dinucleotide)); -; 1([4Fe-4S] cluster) | 223; -; 50, 54, 58, 93 | binding & cofactor; Keyword & go term, Keyword & go term; Keyword & binding & cofactor & go term |
| 88 | rnd | P09155 | divalent metal cation | - | - | cofactor |
| 89 | vsr | P09184 | magnesium(2+); magnesium(2+); zinc(2+) | 1(Mg(2+)); 2(Mg(2+)); - | 51, 63; 51; - | Keyword & binding & cofactor & go term; Keyword & binding & cofactor & go term; Keyword & cofactor & go term |
| 90 | nagE | P09323 | zinc(2+) | 1(Zn(2+)) | 554, 569 | Keyword & binding & cofactor & go term |
| 91 | glpQ | P09394 | calcium(2+) | 1(Ca(2+)) | 63, 65, 171 | Keyword & binding & cofactor & go term |

Supplementary Table 2: E. coli annotated metal-binding proteome *(continued)*

| # | Gene Name | UniProt ID | Ligand Name | Ligand Identifier | Ligand Position | Source |
|---|---|---|---|---|---|---|
| 92 | gltB | P09831 | iron cation, Fe3S4 iron-sulfur cluster; tri-mu-sulfido-mu3-sulfido-triiron(0) | -; 1([3Fe-4S] cluster) | -; 1102, 1108, 1113 | Keyword & go term, go term; Keyword & binding & cofactor |
| 93 | gltD | P09832 | iron cation; tetra-mu3-sulfido-tetrairon | -; 1([4Fe-4S] cluster) | -; 47, 50, 55, 59 | Keyword & go term; Keyword & binding & cofactor & go term |
| 94 | modC | P09833 | molybdenum cation, molybdate | - | - | Keyword, catalytic activity & go term |
| 95 | uvrA | P0A698 | zinc(2+) | - | - | Keyword & go term |
| 96 | ackA | P0A6A3 | magnesium(2+); manganese(2+), zinc(2+) | 1(Mg(2+)); - | 10, 387; - | Keyword & binding & cofactor & go term; Keyword & cofactor & go term, Keyword & go term |
| 97 | leuC | P0A6A6 | iron cation; tetra-mu3-sulfido-tetrairon | -; 1([4Fe-4S] cluster) | -; 347, 407, 410 | Keyword & go term; Keyword & binding & cofactor & go term |
| 98 | iscS | P0A6B7 | di-mu-sulfido-diiron; iron cation | 1([2Fe-2S] cluster); - | 328; - | Keyword & binding & go term; Keyword & go term |
| 99 | nfo | P0A6C1 | manganese(2+); zinc(2+); zinc(2+); zinc(2+) | -; 1(Zn(2+)); 2(Zn(2+)); 3(Zn(2+)) | -; 69, 109, 145; 145, 179, 216, 261; 182, 229, 231 | Keyword & cofactor & go term; Keyword & binding & cofactor & go term; Keyword & binding & cofactor & go term; Keyword & binding & cofactor & go term |
| 100 | aroK | P0A6D7 | magnesium(2+) | 1(Mg(2+)) | 18 | Keyword & binding & cofactor & go term |
| 101 | aroL | P0A6E1 | magnesium(2+) | 1(Mg(2+)) | 16, 32 | Keyword & binding & cofactor & go term |
| 102 | bioD2 | P0A6E9 | magnesium(2+) | 1(Mg(2+)) | 17, 55, 112 | Keyword & binding & cofactor & go term |
| 103 | glpK | P0A6F3 | zinc(2+) | 1(Zn(2+)) | 479 | Keyword & binding & go term |
| 104 | groEL | P0A6F5 | magnesium(2+) | - | - | go term |
| 105 | groES | P0A6F9 | metal cation | - | - | go term |
| 106 | clpX | P0A6H1 | zinc(2+) | 1(Zn(2+)) | 15, 18, 37, 40 | Keyword & binding & cofactor & go term |
| 107 | hslU | P0A6H5 | magnesium(2+) | - | - | go term |
| 108 | coaD | P0A6I6 | magnesium(2+) | - | - | Keyword & cofactor |
| 109 | ddlA | P0A6J8 | magnesium(2+); magnesium(2+); manganese(2+) | 1(Mg(2+)); 2(Mg(2+)); - | 302, 315; 315, 317; - | Keyword & binding & cofactor & go term; Keyword & binding & cofactor & go term; Keyword & cofactor & go term |
| 110 | def | P0A6K3 | iron(2+); zinc(2+) | 1(Fe cation); - | 91, 133, 137; - | Keyword & binding & cofactor & go term; Keyword & cofactor & go term |

Supplementary Table 2: E. coli annotated metal-binding proteome *(continued)*

| # | Gene Name | UniProt ID | Ligand Name | Ligand Identifier | Ligand Position | Source |
|---|---|---|---|---|---|---|
| 111 | deoB | P0A6K6 | cobalt(2+), magnesium(2+); manganese(2+); manganese(2+) | -; 1(Mn(2+)); 2(Mn(2+)) | -; 10, 347, 348; 306, 311, 359 | Keyword & cofactor & go term, Keyword & go term; Keyword & binding & cofactor & go term; Keyword & binding & cofactor & go term |
| 112 | tsf | P0A6P1 | zinc(2+) | - | - | go term |
| 113 | engB | P0A6P7 | magnesium(2+) | 1(Mg(2+)) | 40, 62 | Keyword & binding & cofactor & go term |
| 114 | eno | P0A6P9 | magnesium(2+); magnesium(2+) | 1(Mg(2+)); 2(Mg(2+)) | 246, 290, 317; 42 | Keyword & binding & cofactor & go term; Keyword & binding & cofactor & go term |
| 115 | galK | P0A6T3 | magnesium(2+) | 1(Mg(2+)) | 130, 162 | Keyword & binding & go term |
| 116 | folE | P0A6T5 | zinc(2+) | 1(Zn(2+)) | 111, 114, 182 | Keyword & binding & go term |
| 117 | glgC | P0A6V1 | magnesium(2+) | - | - | go term |
| 118 | mraY | P0A6W3 | magnesium(2+) | - | - | Keyword & cofactor & go term |
| 119 | gshA | P0A6W9 | metal cation | - | - | go term |
| 120 | hslO | P0A6Y5 | zinc(2+) | - | - | Keyword & go term |
| 121 | dnaK | P0A6Y8 | zinc(2+) | - | - | go term |
| 122 | nikR | P0A6Z6 | nickel(2+) | 1(Ni(2+)) | 76, 87, 89, 95 | Keyword & binding & cofactor & go term |
| 123 | hypA | P0A700 | nickel(2+); zinc(2+) | 1(Ni(2+)); 1(Zn(2+)) | 2; 73, 76, 90, 93 | Keyword & binding & go term; Keyword & binding & go term |
| 124 | hybF | P0A703 | nickel(2+); zinc(2+) | 1(Ni(2+)); 1(Zn(2+)) | 2, 3; 73, 76, 89, 92 | Keyword & binding & go term; Keyword & binding & go term |
| 125 | prs | P0A717 | magnesium(2+); magnesium(2+); manganese cation | 1(Mg(2+)); 2(Mg(2+)); - | 131; 170; - | Keyword & binding & cofactor & go term; Keyword & binding & cofactor & go term; Keyword & go term |
| 126 | lpxC | P0A725 | iron(2+); zinc(2+) | -; 1(Zn(2+)) | -; 79, 238, 242 | Keyword & cofactor & go term; Keyword & binding & cofactor & go term |
| 127 | yceF | P0A729 | divalent metal cation | - | - | cofactor |
| 128 | msrB | P0A746 | iron cation; zinc(2+) | -; 1(Zn(2+)) | -; 46, 49, 95, 98 | Keyword & go term; Keyword & binding & cofactor & go term |
| 129 | ndk | P0A763 | magnesium(2+) | - | - | Keyword & cofactor & go term |
| 130 | mntH | P0A769 | iron cation, manganese(2+), cadmium(2+) | - | - | Keyword & go term, Keyword & go term, go term |
| 131 | rppH | P0A776 | magnesium(2+), zinc(2+), manganese(2+) | - | - | Keyword & cofactor & go term, Keyword & cofactor, Keyword & cofactor |
| 132 | orn | P0A784 | zinc(2+) | 1(Zn(2+)) | 12, 14, 163 | Keyword & binding & go term |
| 133 | pfkA | P0A796 | magnesium(2+) | 1(Mg(2+)) | 104 | Keyword & binding & cofactor & go term |

Supplementary Table 2: E. coli annotated metal-binding proteome *(continued)*

| # | Gene Name | UniProt ID | Ligand Name | Ligand Identifier | Ligand Position | Source |
|---|---|---|---|---|---|---|
| 134 | ppa | P0A7A9 | magnesium(2+); magnesium(2+); zinc(2+) | 1(Mg(2+)); 2(Mg(2+)); - | 66, 71, 103; 71; - | Keyword & binding & cofactor & go term; Keyword & binding & cofactor & go term; Keyword & go term |
| 135 | ppk | P0A7B1 | magnesium(2+) | 1(Mg(2+)) | 375, 405 | Keyword & binding & cofactor & go term |
| 136 | nadK | P0A7B3 | divalent metal cation | - | - | cofactor & go term |
| 137 | proB | P0A7B5 | magnesium(2+) | - | - | go term |
| 138 | hslV | P0A7B8 | magnesium(2+); sodium(1+) | -; 1(Na(+)) | -; 157, 160, 163 | Keyword & go term; Keyword & binding & go term |
| 139 | purA | P0A7D4 | magnesium(2+) | 1(Mg(2+)) | 14, 41 | Keyword & binding & cofactor & go term |
| 140 | pyrE | P0A7E3 | magnesium(2+) | - | - | Keyword & cofactor & go term |
| 141 | pyrG | P0A7E5 | magnesium(2+) | 1(Mg(2+)) | 72, 140 | Keyword & binding & go term |
| 142 | pyrI | P0A7F3 | zinc(2+) | 1(Zn(2+)) | 109, 114, 138, 141 | Keyword & binding & cofactor & go term |
| 143 | speD | P0A7F6 | magnesium(2+) | - | - | Keyword & go term |
| 144 | recR | P0A7H6 | zinc cation | - | - | Keyword & go term |
| 145 | ribA | P0A7I7 | magnesium(2+); zinc(2+) | -; 1(Zn(2+)) | -; 54, 65, 67 | Keyword & go term; Keyword & binding & cofactor & go term |
| 146 | ribB | P0A7J0 | magnesium(2+); magnesium(2+); manganese(2+) | 1(Mg(2+)); 2(Mg(2+)); - | 38; 38, 153; - | Keyword & binding & cofactor & go term; Keyword & binding & cofactor & go term; Keyword & cofactor & go term |
| 147 | rpmE | P0A7M9 | zinc(2+) | 1(Zn(2+)) | 16 | Keyword & binding & cofactor & go term |
| 148 | rpsB | P0A7V0 | zinc(2+) | - | - | go term |
| 149 | rnc | P0A7Y0 | magnesium(2+) | 1(Mg(2+)) | 41, 114, 117 | Keyword & binding & cofactor & go term |
| 150 | rnhA | P0A7Y4 | magnesium(2+); magnesium(2+) | 1(Mg(2+)); 2(Mg(2+)) | 10, 48, 70; 10, 134 | Keyword & binding & cofactor & go term; Keyword & binding & cofactor & go term |
| 151 | ruvA | P0A809 | magnesium(2+) | - | - | cofactor |
| 152 | ruvB | P0A812 | magnesium(2+) | 1(Mg(2+)) | 69 | binding & cofactor |
| 153 | ruvC | P0A814 | magnesium(2+); magnesium(2+) | 1(Mg(2+)); 2(Mg(2+)) | 8, 139; 67 | Keyword & binding & cofactor & go term; Keyword & binding & cofactor & go term |
| 154 | metK | P0A817 | magnesium(2+); manganese(2+), cobalt(2+); potassium(1+) | 1(Mg(2+)); -; 1(K(+)) | 17; -; 43 | Keyword & binding & cofactor & go term; Keyword & cofactor & go term, Keyword & cofactor & go term; Keyword & binding & cofactor & go term |
| 155 | glyA | P0A825 | zinc(2+) | - | - | go term |
| 156 | sucC | P0A836 | magnesium(2+) | 1(Mg(2+)) | 199, 213 | Keyword & binding & cofactor & go term |

Supplementary Table 2: E. coli annotated metal-binding proteome *(continued)*

| # | Gene Name | UniProt ID | Ligand Name | Ligand Identifier | Ligand Position | Source |
|---|---|---|---|---|---|---|
| 157 | surE | P0A840 | cobalt(2+), magnesium(2+), manganese(2+), nickel(2+), zinc(2+) | 1(a divalent metal cation) | 8, 9, 39, 92 | Keyword & binding & cofactor & go term, Keyword & binding & cofactor & go term, Keyword & binding & cofactor & go term, Keyword & binding & cofactor & go term, Keyword & binding & cofactor & go term |
| 158 | tgt | P0A847 | zinc(2+) | 1(Zn(2+)) | 302, 304, 307, 333 | Keyword & binding & cofactor & go term |
| 159 | tnaA | P0A853 | potassium(1+) | - | - | go term |
| 160 | trmD | P0A873 | magnesium(2+) | - | - | go term |
| 161 | thyA | P0A884 | magnesium(2+) | - | - | go term |
| 162 | ybeY | P0A898 | nickel cation;<br>zinc(2+) | -;<br>1(Zn(2+)) | -;<br>114, 118, 124 | Keyword & go term;<br>Keyword & binding & cofactor & go term |
| 163 | nrdR | P0A8D0 | zinc(2+) | - | - | Keyword & cofactor & go term |
| 164 | upp | P0A8F0 | magnesium(2+) | - | - | Keyword & cofactor & go term |
| 165 | zupT | P0A8H3 | cadmium(2+), cobalt(2+), manganese(2+), copper(2+);<br>iron(2+);<br>zinc(2+) | -;<br>1(Fe(2+));<br>1(Zn(2+)) | -;<br>120, 123, 149, 152, 181;<br>123, 148, 152 | Keyword & catalytic activity & go term, Keyword & catalytic activity & go term, Keyword & catalytic activity & go term, Keyword & catalytic activity & go term;<br>Keyword & binding & catalytic activity & go term;<br>Keyword & binding & catalytic activity & go term |
| 166 | yacG | P0A8H8 | zinc(2+) | 1(Zn(2+)) | 9, 12, 28, 32 | Keyword & binding & cofactor & go term |
| 167 | dnaT | P0A8J2 | magnesium(2+) | - | - | go term |
| 168 | serS | P0A8L1 | magnesium(2+) | - | - | go term |
| 169 | thrS | P0A8M3 | zinc(2+) | 1(Zn(2+)) | 334, 385, 511 | Keyword & binding & cofactor & go term |
| 170 | lysS | P0A8N3 | magnesium(2+);<br>magnesium(2+) | 1(Mg(2+));<br>2(Mg(2+)) | 415, 422;<br>422 | Keyword & binding & cofactor & go term;<br>Keyword & binding & cofactor & go term |
| 171 | lysU | P0A8N5 | magnesium(2+);<br>magnesium(2+) | 1(Mg(2+));<br>2(Mg(2+)) | 415, 422;<br>422 | Keyword & binding & cofactor & go term;<br>Keyword & binding & cofactor & go term |
| 172 | yggX | P0A8P3 | iron cation | - | - | Keyword & go term |
| 173 | rpoC | P0A8T7 | magnesium(2+);<br>zinc(2+);<br>zinc(2+) | 1(Mg(2+));<br>1(Zn(2+));<br>2(Zn(2+)) | 460, 462, 464;<br>70, 72, 85, 88;<br>814, 888, 895, 898 | Keyword & binding & cofactor & go term;<br>Keyword & binding & cofactor & go term;<br>Keyword & binding & cofactor & go term |
| 174 | rbn | P0A8V0 | zinc(2+);<br>zinc(2+) | 1(Zn(2+));<br>2(Zn(2+)) | 64, 66, 141, 212;<br>68, 69, 212, 270 | Keyword & binding & cofactor & go term;<br>Keyword & binding & cofactor & go term |

Supplementary Table 2: E. coli annotated metal-binding proteome *(continued)*

| # | Gene Name | UniProt ID | Ligand Name | Ligand Identifier | Ligand Position | Source |
|---|---|---|---|---|---|---|
| 175 | yjjG | P0A8Y1 | manganese(2+), magnesium(2+), cobalt(2+) | - | - | Keyword & cofactor & go term, Keyword & cofactor & go term, Keyword & cofactor & go term |
| 176 | yihX | P0A8Y3 | magnesium(2+); manganese(2+), cobalt(2+), zinc(2+) | 1(Mg(2+)); - | 6, 166; - | Keyword & binding & cofactor & go term; Keyword & cofactor & go term, Keyword & cofactor & go term, Keyword & cofactor & go term |
| 177 | yidA | P0A8Y5 | magnesium(2+); manganese(2+), cobalt(2+), zinc(2+) | 1(Mg(2+)); - | 9, 11, 220; - | Keyword & binding & cofactor & go term; Keyword & cofactor & go term, Keyword & cofactor & go term, Keyword & cofactor & go term |
| 178 | pldA | P0A921 | calcium(2+); calcium(2+); calcium(2+) | 1(Ca(2+)); 2(Ca(2+)); 3(Ca(2+)) | 126; 167, 172; 204 | Keyword & binding & cofactor & go term; Keyword & binding & cofactor & go term; Keyword & binding & cofactor & go term |
| 179 | speG | P0A951 | magnesium(2+) | 1(Mg(2+)) | 35, 76 | Keyword & binding & go term |
| 180 | fbp | P0A993 | magnesium(2+); magnesium(2+) | 1(Mg(2+)); 2(Mg(2+)) | 89, 110, 112; 110, 113, 275 | Keyword & binding & cofactor & go term; Keyword & binding & cofactor & go term |
| 181 | glpC | P0A996 | iron cation; tetra-mu3-sulfido-tetrairon; tetra-mu3-sulfido-tetrairon | -; 1([4Fe-4S] cluster); 2([4Fe-4S] cluster) | -; 9, 12, 15, 66; 19, 56, 59, 62 | Keyword & go term; Keyword & binding & go term; Keyword & binding & go term |
| 182 | ftnA | P0A998 | iron(2+), iron(3+); iron(2+), iron(3+); iron(2+), iron(3+); iron(III) oxide-hydroxide(1-) | 1(Fe cation); 2(Fe cation); 3(Fe cation); - | 17, 50, 53; 50, 94, 127, 130; 49, 126, 130; - | Keyword & binding & catalytic activity & go term, Keyword & binding & go term; Keyword & binding & catalytic activity & go term, Keyword & binding & go term; Keyword & binding & catalytic activity & go term, Keyword & binding & go term; catalytic activity |
| 183 | ftnB | P0A9A2 | iron(3+), iron(2+) | - | - | go term, go term |
| 184 | fur | P0A9A9 | iron cation; zinc(2+) | 1(Fe cation); 1(Zn(2+)) | 87, 89, 108, 125; 33, 81, 90, 93, 96, 101 | Keyword & binding & go term; Keyword & binding & go term |
| 185 | glnA | P0A9C5 | magnesium(2+); magnesium(2+) | 1(Mg(2+)); 2(Mg(2+)) | 130, 270, 358; 132, 213, 221 | Keyword & binding & cofactor & go term; Keyword & binding & cofactor & go term |

Supplementary Table 2: E. coli annotated metal-binding proteome *(continued)*

| # | Gene Name | UniProt ID | Ligand Name | Ligand Identifier | Ligand Position | Source |
|---|---|---|---|---|---|---|
| 186 | glpX | P0A9C9 | manganese(2+); manganese(2+) | 1(Mn(2+)); 2(Mn(2+)) | 33, 57; 85, 88, 213 | Keyword & binding & cofactor & go term; Keyword & binding & cofactor & go term |
| 187 | fnr | P0A9E5 | iron cation; tetra-mu3-sulfido-tetrairon | -; 1([4Fe-4S] cluster) | -; 20, 23, 29, 122 | Keyword & go term; Keyword & binding & cofactor & go term |
| 188 | yeiL | P0A9E9 | iron cation; tetra-mu3-sulfido-tetrairon | -; 1([4Fe-4S] cluster) | -; 68, 91, 93, 116 | Keyword & go term; Keyword & binding & cofactor & go term |
| 189 | mntR | P0A9F1 | manganese(2+) | - | - | Keyword & go term |
| 190 | cueR | P0A9G4 | copper(1+) | 1(Cu(+)) | 112, 120 | Keyword & binding & go term |
| 191 | aceA | P0A9G6 | magnesium(2+); manganese cation | 1(Mg(2+)); - | 157; - | Keyword & binding & cofactor & go term; Keyword & go term |
| 192 | modE | P0A9G8 | molybdate; molybdenum cation | 1(molybdate); - | 126, 128, 163, 166, 183, 184; - | binding; Keyword & go term |
| 193 | btuR | P0A9H5 | adenosylcob(III)yrinate a,c-diamide(4-), cob(II)yrinic acid a,c diamide(4-), cob(II)alamin, cobamamide | | - | catalytic activity, catalytic activity, catalytic activity, catalytic activity |
| 194 | citE | P0A9I1 | magnesium(2+) | 1(Mg(2+)) | 139, 166 | Keyword & binding & cofactor & go term |
| 195 | nirD | P0A9I8 | Fe2S2 iron-sulfur cluster | - | - | go term |
| 196 | rng | P0A9J0 | magnesium(2+) | 1(Mg(2+)) | 304, 347 | Keyword & binding & cofactor & go term |
| 197 | rbsK | P0A9J6 | magnesium(2+), manganese(2+); potassium(1+) | -; 1(K(+)) | -; 249, 251, 285, 288, 290, 294 | Keyword & cofactor & go term, Keyword & cofactor & go term; Keyword & binding & go term |
| 198 | phoU | P0A9K7 | magnesium(2+), manganese(2+) | - | - | go term, go term |
| 199 | slyD | P0A9K9 | copper cation, zinc(2+), cobalt(2+); nickel(2+) | -; 1(Ni(2+)) | -; 167, 168, 184, 185, 193, 195 | Keyword & go term, Keyword & go term, Keyword & go term; Keyword & binding & go term |
| 200 | hpt | P0A9M2 | magnesium(2+) | 1(Mg(2+)) | 99, 100 | Keyword & binding & cofactor & go term |
| 201 | gpt | P0A9M5 | magnesium(2+) | 1(Mg(2+)) | 89 | Keyword & binding & cofactor & go term |
| 202 | pta | P0A9M8 | zinc(2+) | - | - | go term |
| 203 | pflA | P0A9N4 | iron cation, potassium(1+); tetra-mu3-sulfido-tetrairon | -; 1([4Fe-4S] cluster) | -; 30, 34, 37 | Keyword & go term, Keyword & go term; Keyword & binding & cofactor & go term |
| 204 | nrdG | P0A9N8 | iron cation; tetra-mu3-sulfido-tetrairon | -; 1([4Fe-4S] cluster) | -; 26, 30, 33 | Keyword & go term; Keyword & binding & cofactor & go term |
| 205 | lpdA | P0A9P0 | zinc(2+) | - | - | go term |
| 206 | accD | P0A9Q5 | zinc(2+) | 1(Zn(2+)) | 27, 30, 46, 49 | Keyword & binding & cofactor & go term |

Supplementary Table 2: E. coli annotated metal-binding proteome *(continued)*

| # | Gene Name | UniProt ID | Ligand Name | Ligand Identifier | Ligand Position | Source |
|---|---|---|---|---|---|---|
| 207 | adhE | P0A9Q7 | iron(2+) | 1(Fe cation) | 653, 657, 723, 737 | Keyword & binding & cofactor & go term |
| 208 | fdx | P0A9R4 | di-mu-sulfido-diiron; iron cation | 1([2Fe-2S] cluster); - | 42, 48, 51, 87; - | Keyword & binding & cofactor & go term; Keyword & go term |
| 209 | fucO | P0A9S1 | iron(2+) | 1(Fe cation) | 195, 199, 262, 276 | Keyword & binding & cofactor & go term |
| 210 | gatD | P0A9S3 | zinc(2+); zinc(2+) | 1(Zn(2+)); 2(Zn(2+)) | 38, 59, 144; 89, 92, 95, 103 | Keyword & binding & cofactor & go term; Keyword & binding & cofactor & go term |
| 211 | gldA | P0A9S5 | zinc(2+) | 1(Zn(2+)) | 171, 254, 271 | Keyword & binding & cofactor & go term |
| 212 | ibaG | P0A9W6 | Fe2S2 iron-sulfur cluster | - | - | go term |
| 213 | yrdA | P0A9W9 | zinc(2+) | - | - | go term |
| 214 | znuC | P0A9X1 | zinc(2+) | - | - | Keyword & catalytic activity & go term |
| 215 | rplM | P0AA10 | zinc(2+) | - | - | go term |
| 216 | yobA | P0AA57 | copper cation | 1(Cu cation) | 27, 113 | Keyword & binding & go term |
| 217 | dosC | P0AA89 | heme; iron cation; magnesium(2+) | 1(heme); -; 1(Mg(2+)) | 98; -; 333, 376 | Keyword & binding & cofactor & go term; Keyword & go term; Keyword & binding & cofactor & go term |
| 218 | zraP | P0AAA9 | copper cation, zinc(2+), nickel cation, cobalt(2+) | - | - | go term, Keyword & go term, go term, go term |
| 219 | wzb | P0AAB2 | metal cation | - | - | go term |
| 220 | ubiD | P0AAB4 | manganese(2+) | 1(Mn(2+)) | 175, 241 | Keyword & binding & cofactor & go term |
| 221 | ybhL | P0AAC4 | calcium(2+) | - | - | go term |
| 222 | yccA | P0AAC6 | calcium(2+) | - | - | go term |
| 223 | iscA | P0AAC8 | Fe2S2 iron-sulfur cluster; iron(2+) | -; 1(Fe cation) | -; 35, 99, 101 | go term; Keyword & binding & cofactor & go term |
| 224 | dppD | P0AAG0 | heme, Fe4S4 iron-sulfur cluster | - | - | go term, go term |
| 225 | sapD | P0AAH4 | potassium(1+) | - | - | Keyword |
| 226 | ftsH | P0AAI3 | zinc(2+) | 1(Zn(2+)) | 414, 418, 492 | Keyword & binding & cofactor & go term |
| 227 | ynfG | P0AAJ1 | iron cation; tetra-mu3-sulfido-tetrairon; tetra-mu3-sulfido-tetrairon; tetra-mu3-sulfido-tetrairon; tetra-mu3-sulfido-tetrairon | -; 1([4Fe-4S] cluster); 2([4Fe-4S] cluster); 3([4Fe-4S] cluster); 4([4Fe-4S] cluster) | -; 14, 17, 20, 145; 24, 126, 129, 141; 67, 70, 75, 109; 79, 99, 102, 105 | Keyword & go term; Keyword & binding & cofactor & go term; Keyword & binding & cofactor & go term; Keyword & binding & cofactor & go term; Keyword & binding & cofactor & go term |

Supplementary Table 2: E. coli annotated metal-binding proteome *(continued)*

| # | Gene Name | UniProt ID | Ligand Name | Ligand Identifier | Ligand Position | Source |
|---|---|---|---|---|---|---|
| 228 | fdnH | P0AAJ3 | iron cation; tetra-mu3-sulfido-tetrairon; tetra-mu3-sulfido-tetrairon; tetra-mu3-sulfido-tetrairon; tetra-mu3-sulfido-tetrairon | -; 1([4Fe-4S] cluster); 2([4Fe-4S] cluster); 3([4Fe-4S] cluster); 4([4Fe-4S] cluster) | -; 39, 42, 45, 179; 49, 160, 163, 175; 100, 103, 108, 143; 112, 133, 136, 139 | Keyword & go term; Keyword & binding & cofactor & go term; Keyword & binding & cofactor & go term; Keyword & binding & cofactor & go term; Keyword & binding & cofactor & go term |
| 229 | fdoH | P0AAJ5 | iron cation; tetra-mu3-sulfido-tetrairon; tetra-mu3-sulfido-tetrairon; tetra-mu3-sulfido-tetrairon; tetra-mu3-sulfido-tetrairon | -; 1([4Fe-4S] cluster); 2([4Fe-4S] cluster); 3([4Fe-4S] cluster); 4([4Fe-4S] cluster) | -; 39, 42, 45, 179; 49, 160, 163, 175; 100, 103, 108, 143; 112, 133, 136, 139 | Keyword & go term; Keyword & binding & cofactor & go term; Keyword & binding & cofactor & go term; Keyword & binding & cofactor & go term; Keyword & binding & cofactor & go term |
| 230 | hybA | P0AAJ8 | iron cation; tetra-mu3-sulfido-tetrairon; tetra-mu3-sulfido-tetrairon; tetra-mu3-sulfido-tetrairon; tetra-mu3-sulfido-tetrairon | -; 1([4Fe-4S] cluster); 2([4Fe-4S] cluster); 3([4Fe-4S] cluster); 4([4Fe-4S] cluster) | -; 47, 50, 53, 197; 57, 174, 177, 193; 112, 115, 120, 155; 124, 145, 148, 151 | Keyword & go term; Keyword & binding & cofactor & go term; Keyword & binding & cofactor & go term; Keyword & binding & cofactor & go term; Keyword & binding & cofactor & go term |
| 231 | hycB | P0AAK1 | iron cation; tetra-mu3-sulfido-tetrairon; tetra-mu3-sulfido-tetrairon; tetra-mu3-sulfido-tetrairon; tetra-mu3-sulfido-tetrairon | -; 1([4Fe-4S] cluster); 2([4Fe-4S] cluster); 3([4Fe-4S] cluster); 4([4Fe-4S] cluster) | -; 12, 15, 18, 159; 22, 143, 146, 155; 51, 54, 59, 92; 63, 82, 85, 88 | Keyword & go term; Keyword & binding & cofactor & go term; Keyword & binding & cofactor & go term; Keyword & binding & cofactor & go term; Keyword & binding & cofactor & go term |
| 232 | hydN | P0AAK4 | iron cation; tetra-mu3-sulfido-tetrairon; tetra-mu3-sulfido-tetrairon; tetra-mu3-sulfido-tetrairon; tetra-mu3-sulfido-tetrairon | -; 1([4Fe-4S] cluster); 2([4Fe-4S] cluster); 3([4Fe-4S] cluster); 4([4Fe-4S] cluster) | -; 12, 15, 18, 147; 22, 131, 134, 143; 58, 61, 66, 99; 70, 89, 92, 95 | Keyword & go term; Keyword & binding & cofactor & go term; Keyword & binding & cofactor & go term; Keyword & binding & cofactor & go term; Keyword & binding & cofactor & go term |
| 233 | nrfC | P0AAK7 | iron(2+), iron(3+); tetra-mu3-sulfido-tetrairon; tetra-mu3-sulfido-tetrairon; tetra-mu3-sulfido-tetrairon; tetra-mu3-sulfido-tetrairon | -; 1([4Fe-4S] cluster); 2([4Fe-4S] cluster); 3([4Fe-4S] cluster); 4([4Fe-4S] cluster) | -; 46, 49, 52, 172; 56, 152, 155, 168; 92, 95, 100, 135; 104, 125, 128, 131 | Keyword & go term, Keyword & go term; Keyword & binding & go term; Keyword & binding & go term; Keyword & binding & go term; Keyword & binding & go term |

Supplementary Table 2: E. coli annotated metal-binding proteome *(continued)*

| # | Gene Name | UniProt ID | Ligand Name | Ligand Identifier | Ligand Position | Source |
|---|---|---|---|---|---|---|
| 234 | napF | P0AAL0 | iron cation; tetra-mu3-sulfido-tetrairon; tetra-mu3-sulfido-tetrairon; tetra-mu3-sulfido-tetrairon | -; 1([4Fe-4S] cluster); 2([4Fe-4S] cluster); 3([4Fe-4S] cluster) | -; 37, 40, 43, 47; 69, 72, 75, 79; 141, 144, 147, 151 | Keyword & go term; Keyword & binding & cofactor & go term; Keyword & binding & cofactor & go term; Keyword & binding & cofactor & go term |
| 235 | napG | P0AAL3 | iron cation; tetra-mu3-sulfido-tetrairon; tetra-mu3-sulfido-tetrairon; tetra-mu3-sulfido-tetrairon; tetra-mu3-sulfido-tetrairon | -; 1([4Fe-4S] cluster); 2([4Fe-4S] cluster); 3([4Fe-4S] cluster); 4([4Fe-4S] cluster) | -; 61, 64, 67, 71; 99, 102, 107, 111; 139, 147, 150, 154; 186, 189, 192, 196 | Keyword & go term; Keyword & binding & cofactor & go term; Keyword & binding & cofactor & go term; Keyword & binding & cofactor & go term; Keyword & binding & cofactor & go term |
| 236 | ydhY | P0AAL6 | iron cation; tetra-mu3-sulfido-tetrairon; tetra-mu3-sulfido-tetrairon; tetra-mu3-sulfido-tetrairon; tetra-mu3-sulfido-tetrairon | -; 1([4Fe-4S] cluster); 2([4Fe-4S] cluster); 3([4Fe-4S] cluster); 4([4Fe-4S] cluster) | -; 68, 71, 74, 193; 78, 183, 186, 189; 123, 126, 131, 166; 135, 156, 159, 162 | Keyword & go term; Keyword & binding & go term; Keyword & binding & go term; Keyword & binding & go term; Keyword & binding & go term |
| 237 | ykgJ | P0AAL9 | iron cation, iron-sulfur cluster | - | - | Keyword & go term, go term |
| 238 | hyaC | P0AAM1 | heme, iron cation | - | - | Keyword & go term, Keyword & go term |
| 239 | hypC | P0AAM3 | iron cation | - | - | go term |
| 240 | hybG | P0AAM7 | iron cation | - | - | go term |
| 241 | hypB | P0AAN3 | nickel(2+); nickel(2+); zinc(2+) | 1(Ni(2+)); 2(Ni(2+)); 1(Zn(2+)) | 2, 5, 7; 166, 167, 198; 166, 167, 198 | Keyword & binding & go term; Keyword & binding & go term; Keyword & binding & go term |
| 242 | dgcC | P0AAP1 | magnesium(2+) | 1(Mg(2+)) | 248, 249, 291 | Keyword & binding & cofactor & go term |
| 243 | frmR | P0AAP3 | metal cation | - | - | go term |
| 244 | yajD | P0AAQ2 | zinc(2+) | - | - | go term |
| 245 | bhsA | P0AB40 | copper cation | - | - | Keyword |
| 246 | lapB | P0AB58 | iron cation | 1(Fe cation) | 357, 360, 371, 374 | Keyword & binding & go term |
| 247 | fbaA | P0AB71 | zinc(2+); zinc(2+) | 1(Zn(2+)); 2(Zn(2+)) | 111, 227, 265; 145, 175 | Keyword & binding & cofactor & go term; Keyword & binding & cofactor & go term |
| 248 | kbaY | P0AB74 | zinc(2+) | 1(Zn(2+)) | 83, 180, 208 | Keyword & binding & cofactor & go term |
| 249 | kbl | P0AB77 | metal cation | - | - | go term |
| 250 | nth | P0AB83 | iron cation; tetra-mu3-sulfido-tetrairon | -; 1([4Fe-4S] cluster) | -; 187, 194, 197, 203 | Keyword & go term; Keyword & binding & cofactor & go term |
| 251 | apbE | P0AB85 | magnesium(2+) | 1(Mg(2+)) | 185, 299, 302, 303 | Keyword & binding & cofactor & go term |
| 252 | fucA | P0AB87 | zinc(2+) | 1(Zn(2+)) | 73, 92, 94, 155 | Keyword & binding & cofactor & go term |

Supplementary Table 2: E. coli annotated metal-binding proteome *(continued)*

| # | Gene Name | UniProt ID | Ligand Name | Ligand Identifier | Ligand Position | Source |
|---|---|---|---|---|---|---|
| 253 | arsB | P0AB93 | arsenite(1-), antimonous acid | - | - | go term, go term |
| 254 | arsC | P0AB96 | arsenite(1-), arsenate(2-) | - | - | catalytic activity & go term, catalytic activity & go term |
| 255 | mgtA | P0ABB8 | magnesium(2+) | 1(Mg(2+)) | 331, 641, 645, 709, 734, 738 | Keyword & binding & catalytic activity & go term |
| 256 | bfr | P0ABD3 | ferroheme b(2-); iron(2+), iron(3+); iron(2+), iron(3+); iron(2+), iron(3+) | 1(heme b); 1(Fe cation); 2(Fe cation); 3(Fe cation) | 52; 18, 51, 54, 127; 51, 94, 127, 130; 46, 50 | Keyword & binding & cofactor & go term; Keyword & binding & catalytic activity & cofactor & go term, Keyword & binding & catalytic activity & cofactor & go term; Keyword & binding & catalytic activity & cofactor & go term, Keyword & binding & catalytic activity & cofactor & go term; Keyword & binding & catalytic activity & cofactor & go term, Keyword & binding & catalytic activity & cofactor & go term |
| 257 | cybB | P0ABE5 | ferroheme b(2-); ferroheme b(2-); iron cation | 1(heme b); 2(heme b); - | 13, 151; 45, 137; - | Keyword & binding & cofactor & go term; Keyword & binding & cofactor & go term; Keyword & go term |
| 258 | cynT | P0ABE9 | zinc(2+) | 1(Zn(2+)) | 39, 41, 98, 101 | Keyword & binding & cofactor & go term |
| 259 | cdd | P0ABF6 | zinc(2+) | 1(Zn(2+)) | 102, 129, 132 | Keyword & binding & cofactor & go term |
| 260 | corA | P0ABI4 | magnesium(2+), cobalt(2+), manganese(2+), nickel(2+) | - | - | Keyword & catalytic activity & go term, Keyword & catalytic activity & go term, Keyword & catalytic activity, go term |
| 261 | cyoB | P0ABI8 | copper(2+); ferroheme b(2-); ferroheme o(2-); iron(2+), iron(3+) | 1(Cu(2+)); 1(heme b); 1(Fe(II)-heme o); - | 284, 333, 334; 106, 170, 421, 481, 482; 288, 411, 419; - | Keyword & binding & cofactor & go term; Keyword & binding & cofactor & go term; Keyword & binding & cofactor & go term; Keyword & go term, Keyword & go term |
| 262 | cyoA | P0ABJ1 | iron(2+), iron(3+), copper cation | - | - | go term, go term, go term |
| 263 | cyoC | P0ABJ3 | iron(2+), iron(3+) | - | - | go term, go term |
| 264 | cydA | P0ABJ9 | ferroheme b(2-); ferroheme b(2-); iron cation, heme d cis-diol(2-) | b558(heme b); b595(heme b); - | 186, 393; 19; - | Keyword & binding & cofactor & go term; Keyword & binding & cofactor & go term; Keyword & go term, Keyword & cofactor & go term |

Supplementary Table 2: E. coli annotated metal-binding proteome *(continued)*

| # | Gene Name | UniProt ID | Ligand Name | Ligand Identifier | Ligand Position | Source |
|---|---|---|---|---|---|---|
| 265 | cydB | P0ABK2 | iron cation,<br>ferroheme b(2-),<br>heme d cis-diol(2-) | - | - | Keyword & go term,<br>Keyword & cofactor,<br>Keyword & cofactor |
| 266 | nrfA | P0ABK9 | calcium(2+);<br>ferroheme c(2-);<br>ferroheme c(2-);<br>ferroheme c(2-);<br>ferroheme c(2-);<br>ferroheme c(2-);<br>iron(2+), iron(3+) | 1(Ca(2+));<br>1(heme c);<br>2(heme c);<br>3(heme c);<br>4(heme c);<br>5(heme c);<br>- | 215, 216, 261, 263;<br>122, 125, 126;<br>160, 163, 164, 301;<br>94, 209, 212, 213;<br>282, 285, 286, 393;<br>275, 314, 317, 318;<br>- | Keyword & binding &<br>cofactor & go term;<br>Keyword & binding &<br>cofactor & go term;<br>Keyword & binding &<br>cofactor & go term;<br>Keyword & binding &<br>cofactor & go term;<br>Keyword & binding &<br>cofactor & go term;<br>Keyword & binding &<br>cofactor & go term;<br>Keyword & catalytic activity<br>& go term, Keyword &<br>catalytic activity & go term |
| 267 | nrfB | P0ABL1 | heme;<br>heme;<br>heme;<br>heme;<br>heme;<br>iron(2+), iron(3+) | 1(heme);<br>2(heme);<br>3(heme);<br>4(heme);<br>5(heme);<br>- | 49, 52, 53;<br>78, 81, 82;<br>113, 116, 117;<br>138, 141, 142;<br>163, 166, 167;<br>- | Keyword & binding & go<br>term;<br>Keyword & binding & go<br>term;<br>Keyword & binding & go<br>term;<br>Keyword & binding & go<br>term;<br>Keyword & binding & go<br>term;<br>Keyword & go term,<br>Keyword & go term |
| 268 | napB | P0ABL3 | ferroheme c(2-);<br>ferroheme c(2-);<br>iron cation | 1(heme c);<br>2(heme c);<br>- | 69, 83, 86, 87;<br>104, 123, 126, 127;<br>- | Keyword & binding;<br>Keyword & binding;<br>Keyword & go term |
| 269 | napC | P0ABL5 | heme;<br>heme;<br>heme;<br>heme;<br>iron cation | 1(heme);<br>2(heme);<br>3(heme);<br>4(heme);<br>- | 57, 60, 63, 109;<br>87, 90, 91, 188;<br>147, 150, 151;<br>179, 182, 183;<br>- | Keyword & binding & go<br>term;<br>Keyword & binding & go<br>term;<br>Keyword & binding & go<br>term;<br>Keyword & binding & go<br>term;<br>Keyword & go term |
| 270 | ccmB | P0ABL8 | heme | - | - | go term |
| 271 | ccmC | P0ABM1 | heme | - | - | go term |
| 272 | ccmH | P0ABM9 | heme;<br>iron cation | 1(heme);<br>- | 43, 46;<br>- | Keyword & binding;<br>Keyword & go term |
| 273 | dgkA | P0ABN1 | magnesium(2+) | 1(a divalent metal<br>cation) | 29, 77 | Keyword & binding &<br>cofactor & go term |
| 274 | coaBC | P0ABQ0 | magnesium(2+) | - | - | Keyword & cofactor & go<br>term |
| 275 | hcaE | P0ABR5 | di-mu-sulfido-<br>diiron;<br>iron cation | 1([2Fe-2S] cluster);<br>1(Fe cation) | 85, 87, 105, 108;<br>213, 218 | Keyword & binding &<br>cofactor & go term;<br>Keyword & binding &<br>cofactor & go term |

Supplementary Table 2: E. coli annotated metal-binding proteome *(continued)*

| # | Gene Name | UniProt ID | Ligand Name | Ligand Identifier | Ligand Position | Source |
|---|---|---|---|---|---|---|
| 276 | yeaW | P0ABR7 | di-mu-sulfido-diiron; iron cation | 1([2Fe-2S] cluster); 1(Fe cation) | 89, 91, 109, 112; 211, 216, 325 | Keyword & binding & cofactor & go term; Keyword & binding & cofactor & go term |
| 277 | mhpB | P0ABR9 | iron(2+) | - | - | Keyword & go term |
| 278 | dksA | P0ABS1 | zinc(2+) | 1(Zn(2+)) | 114, 117, 135, 138 | Keyword & binding & go term |
| 279 | dnaG | P0ABS5 | magnesium(2+); magnesium(2+); zinc(2+) | 1(Mg(2+)); 2(Mg(2+)); - | 265, 309; 309, 311; - | Keyword & binding & cofactor & go term; Keyword & binding & cofactor & go term; Keyword & cofactor & go term |
| 280 | dps | P0ABT2 | iron(2+), iron(3+); iron(2+), iron(3+) | 1(Fe cation); 2(Fe cation) | 51, 78, 82; 82 | Keyword & binding & catalytic activity & go term, Keyword & binding & catalytic activity & go term; Keyword & binding & catalytic activity & go term, Keyword & binding & catalytic activity & go term |
| 281 | ychF | P0ABU2 | magnesium(2+) | 1(Mg(2+)) | 16, 36 | Keyword & binding & cofactor & go term |
| 282 | hcaC | P0ABW0 | di-mu-sulfido-diiron; iron cation | 1([2Fe-2S] cluster); - | 42, 44, 62, 65; - | Keyword & binding & go term; Keyword & go term |
| 283 | yfaE | P0ABW3 | di-mu-sulfido-diiron; iron cation | 1([2Fe-2S] cluster); - | 37, 42, 45, 74; - | Keyword & binding & cofactor & go term; Keyword & go term |
| 284 | flhC | P0ABY7 | zinc(2+) | 1(Zn(2+)) | 137, 140, 157, 160 | Keyword & binding & cofactor & go term |
| 285 | kdsC | P0ABZ4 | magnesium(2+) | 1(Mg(2+)) | 32, 34, 125 | Keyword & binding & cofactor & go term |
| 286 | folP | P0AC13 | magnesium(2+) | 1(Mg(2+)) | 22 | Keyword & binding & cofactor & go term |
| 287 | fumA | P0AC33 | iron cation; tetra-mu3-sulfido-tetrairon | -; 1([4Fe-4S] cluster) | -; 105, 224, 318 | Keyword & go term; Keyword & binding & cofactor & go term |
| 288 | sdhD | P0AC44 | heme; iron cation | 1(heme); - | 71; - | Keyword & binding & cofactor & go term; Keyword & go term |
| 289 | frdB | P0AC47 | di-mu-sulfido-diiron; iron cation, Fe3S4 iron-sulfur cluster; tetra-mu3-sulfido-tetrairon; tri-mu-sulfido-mu3-sulfido-triiron(0) | 1([2Fe-2S] cluster); -; 1([4Fe-4S] cluster); 1([3Fe-4S] cluster) | 58, 63, 66, 78; -; 149, 152, 155, 215; 159, 205, 211 | Keyword & binding & cofactor & go term; Keyword & go term, go term; Keyword & binding & cofactor & go term; Keyword & binding & cofactor & go term |
| 290 | zur | P0AC51 | zinc(2+) | - | - | Keyword & go term |
| 291 | grxD | P0AC69 | di-mu-sulfido-diiron; iron cation | 1([2Fe-2S] cluster); - | 30; - | Keyword & binding & go term; Keyword & go term |
| 292 | wecA | P0AC78 | magnesium(2+), manganese(2+) | - | - | Keyword & cofactor & go term, Keyword & cofactor & go term |

Supplementary Table 2: E. coli annotated metal-binding proteome *(continued)*

| # | Gene Name | UniProt ID | Ligand Name | Ligand Identifier | Ligand Position | Source |
|---|---|---|---|---|---|---|
| 293 | gloA | P0AC81 | nickel(2+) | 1(Ni(2+)) | 5, 56, 74, 122 | Keyword & binding & cofactor & go term |
| 294 | gloB | P0AC84 | zinc(2+); zinc(2+) | 1(Zn(2+)); 2(Zn(2+)) | 53, 55, 110, 127; 57, 58, 127, 165 | Keyword & binding & cofactor & go term; Keyword & binding & cofactor & go term |
| 295 | dnaB | P0ACB0 | magnesium(2+) | - | - | cofactor |
| 296 | hemB | P0ACB2 | magnesium(2+); zinc(2+) | 1(Mg(2+)); 1(Zn(2+)) | 232; 120, 122, 130 | Keyword & binding & go term; Keyword & binding & cofactor & go term |
| 297 | erpA | P0ACC3 | Fe2S2 iron-sulfur cluster, Fe4S4 iron-sulfur cluster; iron cation | 1(iron-sulfur cluster); - | 42, 106, 108; - | binding & cofactor & go term, binding & cofactor & go term; Keyword & go term |
| 298 | glmU | P0ACC7 | cobalt(2+); magnesium(2+) | 1(Co(2+)); 1(Mg(2+)) | 105, 227; 105, 227 | Keyword & binding & cofactor & go term; Keyword & binding & cofactor & go term |
| 299 | iscU | P0ACD4 | copper cation, iron(2+), zinc(2+), Fe2S2 iron-sulfur cluster, Fe4S4 iron-sulfur cluster | - | - | go term, go term, go term, go term, go term |
| 300 | hyaB | P0ACD8 | di-mu-sulfido-diiron(2+), di-mu-sulfido-diiron(1+); nickel(2+) | -; 1(Ni(2+)) | -; 76, 79, 576, 579 | go term, go term; Keyword & binding & cofactor & go term |
| 301 | hybC | P0ACE0 | di-mu-sulfido-diiron(2+), di-mu-sulfido-diiron(1+); nickel(2+) | -; 1(Ni(2+)) | -; 61, 64, 546, 549 | go term, go term; Keyword & binding & cofactor & go term |
| 302 | soxR | P0ACS2 | di-mu-sulfido-diiron; iron cation | 1([2Fe-2S] cluster); - | 119, 122, 124, 130; - | Keyword & binding & go term; Keyword & go term |
| 303 | zntR | P0ACS5 | zinc(2+) | 1(Zn(2+)) | 114, 115, 119, 124 | Keyword & binding & go term |
| 304 | mpaA | P0ACV6 | zinc(2+) | 1(Zn(2+)) | 49, 52, 157 | Keyword & binding & cofactor & go term |
| 305 | etp | P0ACZ2 | metal cation | - | - | go term |
| 306 | cusR | P0ACZ8 | copper cation | - | - | Keyword |
| 307 | yecA | P0AD05 | iron cation | - | - | go term |
| 308 | pgpC | P0AD42 | magnesium(2+) | - | - | Keyword & cofactor & go term |
| 309 | ispB | P0AD57 | magnesium(2+); magnesium(2+) | 1(Mg(2+)); 2(Mg(2+)) | 84, 88; 84, 88 | Keyword & binding & cofactor & go term; Keyword & binding & cofactor & go term |
| 310 | pykF | P0AD61 | magnesium(2+); potassium(1+) | 1(Mg(2+)); 1(K(+)) | 222, 246; 34, 36, 66, 67 | Keyword & binding & cofactor & go term; Keyword & binding & cofactor & go term |
| 311 | nlpD | P0ADA3 | metal cation | - | - | go term |
| 312 | kbp | P0ADE6 | potassium(1+) | - | - | Keyword & go term |

Supplementary Table 2: E. coli annotated metal-binding proteome *(continued)*

| # | Gene Name | UniProt ID | Ligand Name | Ligand Identifier | Ligand Position | Source |
|---|---|---|---|---|---|---|
| 313 | edd | P0ADF6 | iron cation; tetra-mu3-sulfido-tetrairon | -; 1([4Fe-4S] cluster) | -; 154, 221 | Keyword & go term; Keyword & binding & cofactor & go term |
| 314 | ilvM | P0ADG1 | magnesium(2+) | - | - | Keyword & cofactor |
| 315 | suhB | P0ADG4 | lithium(1+); magnesium(2+) | -; 1(Mg(2+)) | -; 67, 84, 86 | Keyword & go term; Keyword & binding & cofactor & go term |
| 316 | guaB | P0ADG7 | potassium(1+) | 1(K(+)) | 300, 302, 305, 469, 470, 471 | Keyword & binding & cofactor & go term |
| 317 | entB | P0ADI4 | enterobactin(1-); magnesium(2+) | -; 1(Mg(2+)) | -; 227, 242, 244 | go term; Keyword & binding & cofactor & go term |
| 318 | yieE | P0ADM8 | magnesium(2+) | - | - | go term |
| 319 | yigB | P0ADP0 | magnesium(2+); manganese(2+), cobalt(2+), zinc(2+) | 1(Mg(2+)); - | 16, 18, 188; - | Keyword & binding & cofactor & go term; Keyword & cofactor & go term, Keyword & cofactor & go term, Keyword & cofactor & go term |
| 320 | ygiC | P0ADT5 | magnesium(2+); magnesium(2+) | 1(Mg(2+)); 2(Mg(2+)) | 102, 115; 115, 117 | Keyword & binding & go term; Keyword & binding & go term |
| 321 | yhcC | P0ADW6 | iron cation; tetra-mu3-sulfido-tetrairon | -; 1([4Fe-4S] cluster) | -; 33, 45, 48 | Keyword & go term; Keyword & binding & cofactor & go term |
| 322 | map | P0AE18 | cobalt(2+), zinc(2+), manganese(2+), iron(2+); cobalt(2+), zinc(2+), manganese(2+), iron(2+); sodium(1+) | 1(a divalent metal cation); 2(a divalent metal cation); - | 97, 108, 235; 108, 171, 204, 235; - | Keyword & binding & cofactor & go term, Keyword & binding & cofactor & go term, Keyword & binding & cofactor & go term, Keyword & binding & cofactor & go term; Keyword & binding & cofactor & go term, Keyword & binding & cofactor & go term, Keyword & binding & cofactor & go term, Keyword & binding & cofactor & go term; Keyword & cofactor & go term |
| 323 | aphA | P0AE22 | magnesium(2+) | 1(Mg(2+)) | 69, 71, 192 | Keyword & binding & cofactor & go term |
| 324 | bfd | P0AE56 | di-mu-sulfido-diiron; iron cation | 1([2Fe-2S] cluster); - | 4, 6, 39, 42; - | Keyword & binding & cofactor & go term; Keyword & go term |
| 325 | cheY | P0AE67 | magnesium(2+) | 1(Mg(2+)) | 12, 13, 57, 59 | Keyword & binding & cofactor & go term |
| 326 | cobU | P0AE76 | adenosylcobinamide, adenosylcobi-namide phosphate(1-), adenosylcobi-namide guanosyl diphosphate(1-) | - | - | catalytic activity, catalytic activity & go term, catalytic activity & go term |

Supplementary Table 2: E. coli annotated metal-binding proteome *(continued)*

| # | Gene Name | UniProt ID | Ligand Name | Ligand Identifier | Ligand Position | Source |
|---|---|---|---|---|---|---|
| 327 | corC | P0AE78 | cobalt cation,<br>magnesium cation | - | - | Keyword, Keyword |
| 328 | cpxA | P0AE82 | magnesium(2+) | - | - | cofactor |
| 329 | cyoE | P0AEA5 | ferroheme b(2-),<br>ferroheme o(2-) | - | - | catalytic activity & go term,<br>catalytic activity & go term |
| 330 | cysG | P0AEA8 | iron(2+),<br>siroheme(8-),<br>ferroheme b(2-) | - | - | catalytic activity & go term,<br>catalytic activity & go term,<br>go term |
| 331 | dapE | P0AED7 | cobalt(2+);<br>zinc(2+);<br>zinc(2+) | -;<br>1(Zn(2+));<br>2(Zn(2+)) | -;<br>66, 99, 162;<br>99, 134, 348 | Keyword & cofactor & go term;<br>Keyword & binding & cofactor & go term;<br>Keyword & binding & cofactor & go term |
| 332 | mglB | P0AEE5 | calcium(2+) | 1(Ca(2+)) | 157, 159, 161, 163, 165, 228 | Keyword & binding & go term |
| 333 | rseP | P0AEH1 | zinc(2+) | 1(Zn(2+)) | 22, 26 | Keyword & binding & cofactor & go term |
| 334 | miaB | P0AEI1 | iron cation;<br>tetra-mu3-sulfido-tetrairon;<br>tetra-mu3-sulfido-tetrairon | -;<br>1([4Fe-4S] cluster);<br>2([4Fe-4S] cluster) | -;<br>12, 49, 83;<br>157, 161, 164 | Keyword & go term;<br>Keyword & binding & cofactor & go term;<br>Keyword & binding & cofactor & go term |
| 335 | rimO | P0AEI4 | iron cation;<br>tetra-mu3-sulfido-tetrairon;<br>tetra-mu3-sulfido-tetrairon | -;<br>1([4Fe-4S] cluster);<br>2([4Fe-4S] cluster) | -;<br>17, 53, 82;<br>150, 154, 157 | Keyword & go term;<br>Keyword & binding & cofactor & go term;<br>Keyword & binding & cofactor & go term |
| 336 | nudJ | P0AEI6 | magnesium(2+) | - | - | Keyword & cofactor |
| 337 | entC | P0AEJ2 | magnesium(2+);<br>magnesium(2+) | 1(Mg(2+));<br>2(Mg(2+)) | 140, 142, 145, 146;<br>241, 376 | Keyword & binding & cofactor & go term;<br>Keyword & binding & cofactor & go term |
| 338 | eutB | P0AEJ6 | cobalt cation;<br>cobamamide | -;<br>1(adenosylcob(III)alamin) | -;<br>194, 246, 295, 401 | Keyword;<br>binding & cofactor |
| 339 | exoX | P0AEK0 | magnesium(2+) | - | - | Keyword & cofactor |
| 340 | fabG | P0AEK2 | calcium(2+);<br>calcium(2+);<br>calcium(2+) | 1(Ca(2+));<br>2(Ca(2+));<br>3(Ca(2+)) | 50, 53;<br>145;<br>233, 234 | Keyword & binding & go term;<br>Keyword & binding & go term;<br>Keyword & binding & go term |
| 341 | fdnI | P0AEK7 | ferroheme b(2-);<br>ferroheme b(2-);<br>iron cation | 1(heme b);<br>2(heme b);<br>- | 18, 169;<br>57, 155;<br>- | Keyword & binding & cofactor & go term;<br>Keyword & binding & cofactor & go term;<br>Keyword & go term |
| 342 | fdoI | P0AEL0 | ferroheme b(2-);<br>ferroheme b(2-);<br>iron cation | 1(heme b);<br>2(heme b);<br>- | 18, 167;<br>57, 153;<br>- | Keyword & binding & cofactor;<br>Keyword & binding & cofactor;<br>Keyword & go term |
| 343 | feoA | P0AEL3 | iron cation | - | - | Keyword & go term |
| 344 | fepB | P0AEL6 | iron cation | - | - | Keyword |
| 345 | fre | P0AEN1 | iron cation | - | - | Keyword |
| 346 | galU | P0AEP3 | magnesium(2+) | - | - | Keyword & cofactor & go term |

Supplementary Table 2: E. coli annotated metal-binding proteome *(continued)*

| # | Gene Name | UniProt ID | Ligand Name | Ligand Identifier | Ligand Position | Source |
|---|---|---|---|---|---|---|
| 347 | gcl | P0AEP7 | magnesium(2+) | - | - | Keyword & cofactor & go term |
| 348 | gltS | P0AER8 | sodium(1+) | - | - | Keyword & go term |
| 349 | gss | P0AES0 | magnesium(2+); magnesium(2+) | 1(Mg(2+)); 2(Mg(2+)) | 318, 330; 330, 332 | Keyword & binding & go term; Keyword & binding & go term |
| 350 | gudD | P0AES2 | magnesium(2+) | 1(Mg(2+)) | 235, 266, 289 | Keyword & binding & cofactor & go term |
| 351 | gyrB | P0AES6 | magnesium(2+); magnesium(2+); manganese(2+), calcium(2+); potassium(1+); sodium(1+) | 1(Mg(2+)); 2(Mg(2+)); -; 1(K(+)); 1(Na(+)) | 424, 498; 498, 500; -; 94, 97, 100, 117, 121; 103, 105 | Keyword & binding & cofactor & go term; Keyword & binding & cofactor & go term; Keyword & cofactor & go term, Keyword & cofactor & go term; Keyword & binding & cofactor & go term; Keyword & binding & cofactor & go term |
| 352 | hycI | P0AEV9 | nickel(2+) | 1(Ni(2+)) | 16, 62, 90 | Keyword & binding & go term |
| 353 | cpdA | P0AEW4 | iron(2+); iron(2+) | 1(Fe cation); 2(Fe cation) | 22, 24, 64, 205; 64, 94, 164, 203 | Keyword & binding & cofactor & go term; Keyword & binding & cofactor & go term |
| 354 | gsk | P0AEW6 | magnesium(2+) | - | - | Keyword & cofactor |
| 355 | fruK | P0AEW9 | magnesium(2+) | - | - | Keyword & cofactor |
| 356 | mazG | P0AEY3 | magnesium(2+) | 1(Mg(2+)) | 172, 175, 193, 196 | Keyword & binding & cofactor & go term |
| 357 | mdfA | P0AEY8 | sodium(1+), potassium(1+) | - | - | go term, go term |
| 358 | modB | P0AF01 | molybdenum cation, molybdate | - | - | Keyword, go term |
| 359 | mrp | P0AF08 | iron cation, Fe4S4 iron-sulfur cluster | - | - | Keyword & go term, go term |
| 360 | nagA | P0AF18 | cobalt(2+), manganese(2+), cadmium(2+), iron(2+), nickel(2+); zinc(2+) | -; 1(Zn(2+)) | -; 131, 195, 216 | Keyword & cofactor & go term, Keyword & cofactor & go term, Keyword & cofactor & go term, Keyword & cofactor & go term, Keyword & cofactor & go term; Keyword & binding & cofactor & go term |
| 361 | nagD | P0AF24 | magnesium(2+); manganese(2+), cobalt(2+), zinc(2+) | 1(Mg(2+)); - | 9, 11, 201; - | Keyword & binding & cofactor & go term; Keyword & cofactor & go term, Keyword & cofactor & go term, Keyword & cofactor & go term |
| 362 | narJ | P0AF26 | metal cation | - | - | go term |
| 363 | narV | P0AF32 | ferroheme b(2-); ferroheme b(2-); iron cation | 1(heme b); 2(heme b); - | 57, 206; 67, 188; - | Keyword & binding & cofactor & go term; Keyword & binding & cofactor & go term; Keyword & go term |
| 364 | yjbB | P0AF43 | sodium(1+) | - | - | go term |

Supplementary Table 2: E. coli annotated metal-binding proteome *(continued)*

| # | Gene Name | UniProt ID | Ligand Name | Ligand Identifier | Ligand Position | Source |
|---|---|---|---|---|---|---|
| 365 | nsrR | P0AF63 | di-mu-sulfido-diiron; iron cation | 1([2Fe-2S] cluster); - | 91, 96, 102; - | Keyword & binding & cofactor & go term; Keyword & go term |
| 366 | tsaE | P0AF67 | magnesium(2+) | 1(Mg(2+)) | 42, 108 | Keyword & binding & go term |
| 367 | nhaB | P0AFA7 | sodium(1+) | - | - | Keyword & catalytic activity & go term |
| 368 | nikC | P0AFA9 | nickel(2+) | - | - | Keyword & go term |
| 369 | nudB | P0AFC0 | magnesium(2+) | 1(Mg(2+)) | 56, 60, 117 | Keyword & binding & cofactor & go term |
| 370 | nuoB | P0AFC7 | iron cation; tetra-mu3-sulfido-tetrairon | -; 1([4Fe-4S] cluster) | -; 63, 64, 129, 158 | Keyword & go term; Keyword & binding & cofactor & go term |
| 371 | nuoE | P0AFD1 | di-mu-sulfido-diiron; iron cation | 1([2Fe-2S] cluster); - | 92, 97, 133, 137; - | Keyword & binding & cofactor & go term; Keyword & go term |
| 372 | nuoI | P0AFD6 | iron cation; tetra-mu3-sulfido-tetrairon; tetra-mu3-sulfido-tetrairon | -; 1([4Fe-4S] cluster); 2([4Fe-4S] cluster) | -; 60, 63, 66, 109; 70, 99, 102, 105 | Keyword & go term; Keyword & binding & cofactor & go term; Keyword & binding & cofactor & go term |
| 373 | sucA | P0AFG3 | magnesium(2+) | - | - | go term |
| 374 | aceE | P0AFG8 | magnesium(2+) | 1(Mg(2+)) | 231, 261, 263 | Keyword & binding & cofactor & go term |
| 375 | oxc | P0AFI0 | magnesium(2+) | 1(Mg(2+)) | 447, 474, 476 | Keyword & binding & cofactor & go term |
| 376 | pitA | P0AFJ7 | magnesium cation, arsenate(2-), zinc(2+), tellurite | - | - | Keyword, catalytic activity & go term, Keyword & go term, go term |
| 377 | pmbA | P0AFK0 | divalent metal cation | - | - | Keyword & go term |
| 378 | ppx | P0AFL6 | magnesium(2+) | - | - | Keyword & cofactor |
| 379 | ybgI | P0AFP6 | divalent metal cation; divalent metal cation | 1(a divalent metal cation); 2(a divalent metal cation) | 63, 101, 219; 64, 215, 219 | Keyword & binding & go term; Keyword & binding & go term |
| 380 | ycfH | P0AFQ7 | cobalt(2+), manganese(2+), nickel(2+); cobalt(2+), manganese(2+), nickel(2+) | 1(a divalent metal cation); 2(a divalent metal cation) | 7, 9, 94, 205; 94, 130, 155 | Keyword & binding & cofactor & go term, Keyword & binding & cofactor & go term, Keyword & binding & cofactor & go term; Keyword & binding & cofactor & go term, Keyword & binding & cofactor & go term, Keyword & binding & cofactor & go term |
| 381 | mepM | P0AFS9 | calcium(2+); zinc(2+) | -; 1(Zn(2+)) | -; 314 | Keyword & cofactor & go term; Keyword & binding & cofactor & go term |
| 382 | yhiD | P0AFV2 | magnesium cation | - | - | Keyword |
| 383 | trkH | P0AFZ7 | potassium(1+) | 1(K(+)) | 111, 112, 220, 221, 318, 319, 435, 436 | Keyword & binding & go term |

Supplementary Table 2: E. coli annotated metal-binding proteome *(continued)*

| # | Gene Name | UniProt ID | Ligand Name | Ligand Identifier | Ligand Position | Source |
|---|---|---|---|---|---|---|
| 384 | rpe | P0AG07 | cobalt(2+), iron(2+), manganese(2+), zinc(2+) | 1(a divalent metal cation) | 34, 36, 68, 177 | Keyword & binding & cofactor & go term, Keyword & binding & cofactor & go term, Keyword & binding & cofactor & go term, Keyword & binding & cofactor & go term |
| 385 | purF | P0AG16 | magnesium(2+) | 1(Mg(2+)) | 305, 367, 368 | Keyword & binding & cofactor & go term |
| 386 | spoT | P0AG24 | manganese(2+) | - | - | Keyword & cofactor |
| 387 | rpsQ | P0AG63 | zinc(2+) | - | - | go term |
| 388 | rusA | P0AG74 | magnesium(2+) | 1(Mg(2+)) | 70, 72, 91 | Keyword & binding & cofactor & go term |
| 389 | serB | P0AGB0 | magnesium(2+); manganese(2+), cobalt(2+), zinc(2+) | 1(Mg(2+)); - | 12, 116, 118, 272; - | Keyword & binding & cofactor & go term; Keyword & cofactor & go term, Keyword & cofactor & go term, Keyword & cofactor & go term |
| 390 | sodC | P0AGD1 | copper cation; zinc(2+) | 1(Cu cation); 1(Zn(2+)) | 67, 69, 92, 147; 92, 101, 109, 112 | Keyword & binding & cofactor & go term; Keyword & binding & cofactor & go term |
| 391 | sodB | P0AGD3 | iron cation | 1(Fe cation) | 27, 74, 157, 161 | Keyword & binding & cofactor & go term |
| 392 | sstT | P0AGE4 | sodium(1+) | - | - | catalytic activity & go term |
| 393 | chrR | P0AGE6 | chromium(6+), chromium(3+) | - | - | catalytic activity, catalytic activity |
| 394 | thiL | P0AGG0 | magnesium(2+); magnesium(2+); magnesium(2+); magnesium(2+); magnesium(2+) | 1(Mg(2+)); 2(Mg(2+)); 3(Mg(2+)); 4(Mg(2+)); 5(Mg(2+)) | 46, 47, 122; 47, 75; 30, 75, 212; 30, 45, 75; 215 | Keyword & binding & go term; Keyword & binding & go term; Keyword & binding & go term; Keyword & binding & go term; Keyword & binding & go term |
| 395 | trxC | P0AGG4 | zinc(2+) | - | - | Keyword & go term |
| 396 | tldD | P0AGG8 | iron cation, zinc(2+) | - | - | go term, Keyword & go term |
| 397 | trkA | P0AGI8 | potassium(1+) | - | - | Keyword & go term |
| 398 | ubiA | P0AGK1 | magnesium(2+) | - | - | Keyword & cofactor |
| 399 | iscR | P0AGK8 | di-mu-sulfido-diiron; iron cation | 1([2Fe-2S] cluster); - | 92, 98, 104; - | Keyword & binding & cofactor & go term; Keyword & go term |
| 400 | srkA | P0C0K3 | magnesium(2+) | 1(Mg(2+)) | 206, 217 | Keyword & binding & cofactor & go term |
| 401 | iscX | P0C0L9 | iron(2+) | - | - | go term |
| 402 | rlmE | P0C0R7 | magnesium(2+) | - | - | cofactor |
| 403 | mepA | P0C0T5 | zinc(2+); zinc(2+) | 1(Zn(2+)); 2(Zn(2+)) | 110, 113, 120, 211; 147, 150 | Keyword & binding & cofactor & go term; Keyword & binding & cofactor & go term |

Supplementary Table 2: E. coli annotated metal-binding proteome *(continued)*

| # | Gene Name | UniProt ID | Ligand Name | Ligand Identifier | Ligand Position | Source |
|---|---|---|---|---|---|---|
| 404 | rimK | P0C0U4 | magnesium(2+); magnesium(2+); manganese(2+); manganese(2+) | 1(Mg(2+)); 2(Mg(2+)); 1(Mn(2+)); 2(Mn(2+)) | 248, 260; 260, 262; 248, 260; 260, 262 | Keyword & binding & cofactor & go term; Keyword & binding & cofactor & go term; Keyword & binding & cofactor & go term; Keyword & binding & cofactor & go term |
| 405 | gatY | P0C8J6 | zinc(2+) | 1(Zn(2+)) | 83, 180, 208 | Keyword & binding & cofactor & go term |
| 406 | yfjU | P0CF87 | arsenite(1-), arsenate(2-) | - | - | catalytic activity & go term, catalytic activity & go term |
| 407 | ilvG | P0DP89 | magnesium(2+) | - | - | go term |
| 408 | ilvG | P0DP90 | magnesium(2+) | 1(Mg(2+)) | 428, 455 | Keyword & binding & cofactor & go term |
| 409 | ynfU | P0DSF4 | zinc(2+) | 1(Zn(2+)) | 19, 22, 41, 44 | Keyword & binding & cofactor & go term |
| 410 | entE | P10378 | enterobactin(1-) | - | - | go term |
| 411 | secA | P10408 | zinc(2+) | 1(Zn(2+)) | 885, 887, 896, 897 | Keyword & binding & cofactor & go term |
| 412 | iap | P10423 | zinc(2+); zinc(2+) | 1(Zn(2+)); 2(Zn(2+)) | 117, 143, 204; 143, 176 | Keyword & binding & go term; Keyword & binding & go term |
| 413 | rnhB | P10442 | manganese(2+), magnesium(2+) | 1(a divalent metal cation) | 16, 17, 108 | Keyword & binding & cofactor & go term, Keyword & binding & cofactor & go term |
| 414 | ugpQ | P10908 | cobalt(2+), magnesium(2+), manganese(2+) | 1(a divalent metal cation) | 39, 41, 114 | Keyword & binding & cofactor & go term, Keyword & binding & cofactor & go term, Keyword & binding & cofactor & go term |
| 415 | aceK | P11071 | magnesium(2+), manganese(2+) | - | - | go term, go term |
| 416 | narH | P11349 | iron cation, Fe3S4 iron-sulfur cluster; tetra-mu3-sulfido-tetrairon; tetra-mu3-sulfido-tetrairon; tetra-mu3-sulfido-tetrairon; tri-mu-sulfido-mu3-sulfido-triiron(0) | -; 1([4Fe-4S] cluster); 2([4Fe-4S] cluster); 3([4Fe-4S] cluster); 1([3Fe-4S] cluster) | -; 16, 19, 22, 263; 26, 244, 247, 259; 184, 187, 192, 227; 196, 217, 223 | Keyword & go term, go term; Keyword & binding & cofactor & go term; Keyword & binding & cofactor & go term; Keyword & binding & cofactor & go term; Keyword & binding & cofactor |
| 417 | narI | P11350 | ferroheme b(2-); ferroheme b(2-); iron cation | 1(heme b); 2(heme b); - | 56, 205; 66, 187; - | Keyword & binding & cofactor & go term; Keyword & binding & cofactor & go term; Keyword & go term |
| 418 | entF | P11454 | enterobactin(1-) | - | - | go term |
| 419 | nadA | P11458 | iron cation; tetra-mu3-sulfido-tetrairon | -; 1([4Fe-4S] cluster) | -; 113, 200, 297 | Keyword & go term; Keyword & binding & cofactor & go term |
| 420 | fucK | P11553 | divalent metal cation | - | - | cofactor |

Supplementary Table 2: E. coli annotated metal-binding proteome *(continued)*

| # | Gene Name | UniProt ID | Ligand Name | Ligand Identifier | Ligand Position | Source |
|---|---|---|---|---|---|---|
| 421 | tdcD | P11868 | magnesium(2+) | 1(Mg(2+)) | 11 | Keyword & binding & cofactor & go term |
| 422 | moeA | P12281 | molybdenum cation, molybdate, Mo(VI)O2(OH)-molybdopterin cofactor(4-), magnesium(2+) | - | - | Keyword & go term, catalytic activity & go term, catalytic activity & go term, Keyword & cofactor & go term |
| 423 | moeB | P12282 | zinc(2+) | 1(Zn(2+)) | 172, 175, 244, 247 | Keyword & binding & cofactor & go term |
| 424 | udp | P12758 | potassium(1+) | - | - | go term |
| 425 | bioB | P12996 | di-mu-sulfido-diiron(2+), di-mu-sulfido-diiron(1+); iron cation; tetra-mu3-sulfido-tetrairon | 1([2Fe-2S] cluster); -; 1([4Fe-4S] cluster) | 97, 128, 188, 260; -; 53, 57, 60 | Keyword & binding & catalytic activity & cofactor & go term, Keyword & binding & catalytic activity & cofactor & go term; Keyword & go term; Keyword & binding & cofactor & go term |
| 426 | bioD1 | P13000 | magnesium(2+); magnesium(2+) | 1(Mg(2+)); 2(Mg(2+)) | 13; 17, 55, 116 | Keyword & binding & cofactor & go term; Keyword & binding & cofactor & go term |
| 427 | metH | P13009 | cobalt cation; methylcobalamin; zinc(2+) | -; 1(methylcob(III)alamin); 1(Zn(2+)) | -; 694, 756, 757, 758, 759, 760, 804, 808, 860; 247, 310, 311 | Keyword & go term; binding & cofactor; Keyword & binding & cofactor & go term |
| 428 | ampD | P13016 | zinc(2+) | 1(Zn(2+)) | 34, 154, 164 | Keyword & binding & cofactor & go term |
| 429 | fdhE | P13024 | iron(3+) | - | - | go term |
| 430 | katG | P13029 | ferroheme b(2-); iron cation | 1(heme b); - | 267; - | Keyword & binding & cofactor & go term; Keyword & go term |
| 431 | fecA | P13036 | iron cation, siderophore | - | - | Keyword, go term |
| 432 | fes | P13039 | ferrienterobactin(3-), Fe(III)-[N-(2,3-dihydroxybenzoyl)-L-serine], Fe(III)-[N-(2,3-dihydroxybenzoyl)-L-serine]3, Fe(III)-[N-(2,3-dihydroxybenzoyl)-L-serine]2, iron cation, enterobactin(1-) | - | - | catalytic activity, catalytic activity, catalytic activity, catalytic activity, go term, go term |
| 433 | nhaA | P13738 | sodium(1+), lithium(1+) | - | - | Keyword & catalytic activity & go term, catalytic activity |

Supplementary Table 2: E. coli annotated metal-binding proteome *(continued)*

| # | Gene Name | UniProt ID | Ligand Name | Ligand Identifier | Ligand Position | Source |
|---|---|---|---|---|---|---|
| 434 | topB | P14294 | magnesium(2+); magnesium(2+); manganese(2+), calcium(2+) | 1(Mg(2+)); 2(Mg(2+)); - | 7, 103; 103, 105; - | Keyword & binding & cofactor & go term; Keyword & binding & cofactor & go term; Keyword & cofactor & go term, Keyword & cofactor & go term |
| 435 | zraS | P14377 | zinc cation | - | - | Keyword |
| 436 | fumB | P14407 | iron cation; tetra-mu3-sulfido-tetrairon | -; 1([4Fe-4S] cluster) | -; 105, 224, 318 | Keyword & go term; Keyword & binding & cofactor & go term |
| 437 | traI | P14565 | magnesium(2+) | 1(Mg(2+)) | 146, 157, 159 | Keyword & binding & cofactor & go term |
| 438 | murD | P14900 | magnesium(2+) | - | - | cofactor |
| 439 | fecB | P15028 | iron cation | - | - | Keyword |
| 440 | fecD | P15029 | iron cation, iron chelate | - | - | Keyword, go term |
| 441 | fecC | P15030 | iron cation, iron chelate | - | - | Keyword, go term |
| 442 | fecE | P15031 | iron cation, iron(III) dicitrate(3-) | - | - | Keyword, catalytic activity |
| 443 | pepP | P15034 | manganese(2+); manganese(2+) | 1(Mn(2+)); 2(Mn(2+)) | 272, 355, 384, 407; 261, 272, 407 | Keyword & binding & cofactor & go term; Keyword & binding & cofactor & go term |
| 444 | ligA | P15042 | magnesium(2+); zinc(2+) | -; 1(Zn(2+)) | -; 408, 411, 426, 432 | Keyword & cofactor & go term; Keyword & binding & go term |
| 445 | recQ | P15043 | magnesium(2+); manganese(2+); zinc(2+) | -; 1(Mn(2+)); 1(Zn(2+)) | -; 10, 31, 35; 380, 397, 400, 403 | Keyword & cofactor & go term; Keyword & binding & go term; Keyword & binding & cofactor & go term |
| 446 | purL | P15254 | magnesium(2+) | 1(Mg(2+)) | 718, 722, 884 | Keyword & binding & go term |
| 447 | pepD | P15288 | cobalt(2+); zinc(2+); zinc(2+) | -; 1(Zn(2+)); 2(Zn(2+)) | -; 115, 146, 457; 76, 115, 169 | Keyword & cofactor & go term; Keyword & binding & cofactor & go term; Keyword & binding & cofactor & go term |
| 448 | purD | P15640 | magnesium(2+); manganese(2+) | 1(Mg(2+)); - | 286, 288; - | Keyword & binding & cofactor & go term; Keyword & cofactor & go term |
| 449 | dgt | P15723 | magnesium(2+), manganese(2+), cobalt(2+) | - | - | Keyword & cofactor & go term, go term, go term |
| 450 | gcd | P15877 | magnesium(2+) | - | - | go term |
| 451 | sdaA | P16095 | tetra-mu3-sulfido-tetrairon, iron cation | - | - | Keyword & cofactor & go term, Keyword & go term |
| 452 | panF | P16256 | sodium(1+) | - | - | Keyword & catalytic activity & go term |
| 453 | miaA | P16384 | magnesium(2+) | - | - | Keyword & cofactor |

Supplementary Table 2: E. coli annotated metal-binding proteome *(continued)*

| # | Gene Name | UniProt ID | Ligand Name | Ligand Identifier | Ligand Position | Source |
|---|---|---|---|---|---|---|
| 454 | hycE | P16431 | tetra-mu3-sulfido-tetrairon, iron cation, nickel(2+) | - | - | Keyword & cofactor & go term, Keyword & go term, Keyword & cofactor & go term |
| 455 | hycF | P16432 | iron cation; tetra-mu3-sulfido-tetrairon; tetra-mu3-sulfido-tetrairon | -; 1([4Fe-4S] cluster); 2([4Fe-4S] cluster) | -; 40, 43, 46, 50; 75, 78, 81, 85 | Keyword & go term; Keyword & binding & go term; Keyword & binding & go term |
| 456 | hycG | P16433 | iron cation; tetra-mu3-sulfido-tetrairon | -; 1([4Fe-4S] cluster) | -; 45, 51, 115, 145 | Keyword & go term; Keyword & binding & cofactor & go term |
| 457 | selD | P16456 | magnesium(2+) | 1(Mg(2+)) | 51, 91, 227 | Keyword & binding & cofactor & go term |
| 458 | phnJ | P16688 | tetra-mu3-sulfido-tetrairon, iron cation | - | - | Keyword & cofactor & go term, Keyword & go term |
| 459 | phnO | P16691 | divalent metal cation | - | - | cofactor |
| 460 | phnP | P16692 | manganese(2+); manganese(2+); zinc(2+) | 1(Mn(2+)); 2(Mn(2+)); 1(Zn(2+)) | 76, 78, 143, 164; 80, 81, 164, 222; 21, 23, 26, 225 | Keyword & binding & cofactor & go term; Keyword & binding & cofactor & go term; Keyword & binding & cofactor & go term |
| 461 | fhuE | P16869 | coprogen; iron(3+), siderophore | 1(Fe(III)-coprogen); - | 117, 142, 275, 357, 373, 416; - | binding; Keyword & go term, go term |
| 462 | menD | P17109 | magnesium(2+), manganese(2+) | - | - | Keyword & cofactor & go term, Keyword & cofactor & go term |
| 463 | gutQ | P17115 | zinc(2+) | 1(Zn(2+)) | 75 | Keyword & binding & go term |
| 464 | cirA | P17315 | iron cation, siderophore | - | - | Keyword, go term |
| 465 | chbF | P17411 | cobalt(2+), nickel(2+); manganese(2+) | -; 1(Mn(2+)) | -; 172, 203 | Keyword & cofactor & go term, Keyword & cofactor & go term; Keyword & binding & cofactor & go term |
| 466 | betB | P17445 | potassium(1+); potassium(1+) | 1(K(+)); 2(K(+)) | 26, 27, 93, 180; 246, 457, 460 | Keyword & binding & cofactor & go term; Keyword & binding & cofactor & go term |
| 467 | mutY | P17802 | iron cation; tetra-mu3-sulfido-tetrairon | -; 1([4Fe-4S] cluster) | -; 192, 199, 202, 208 | Keyword & go term; Keyword & binding & cofactor & go term |
| 468 | cysI | P17846 | iron cation, di-mu-sulfido-diiron(2+), di-mu-sulfido-diiron(1+); siroheme(8-); tetra-mu3-sulfido-tetrairon | -; 1(siroheme); 1([4Fe-4S] cluster) | -; 483; 434, 440, 479, 483 | Keyword & go term, go term, go term; Keyword & binding & cofactor & go term; Keyword & binding & cofactor & go term |
| 469 | priA | P17888 | zinc(2+) | - | - | Keyword & go term |
| 470 | murC | P17952 | magnesium(2+) | - | - | go term |

Supplementary Table 2: E. coli annotated metal-binding proteome *(continued)*

| # | Gene Name | UniProt ID | Ligand Name | Ligand Identifier | Ligand Position | Source |
|---|---|---|---|---|---|---|
| 471 | pgpA | P18200 | magnesium(2+) | - | - | Keyword & cofactor & go term |
| 472 | dmsA | P18775 | Mo(=O)-bis(molybdopterin guanine dinucleotide)(4-); iron cation, molybdenum cation; tetra-mu3-sulfido-tetrairon | 1(Mo-bis(molybdopterin guanine dinucleotide)); -; 1([4Fe-4S] cluster) | 172, 173, 174, 175, 176, 205, 244, 245, 270, 271, 291, 292, 293, 386, 387, 390, 488, 512, 513, 701, 707, 708, 709, 788, 804, 805; -; 63, 67, 71, 104 | binding & cofactor; Keyword & go term, Keyword & go term; Keyword & binding & cofactor & go term |
| 473 | dmsB | P18776 | iron cation; tetra-mu3-sulfido-tetrairon; tetra-mu3-sulfido-tetrairon; tetra-mu3-sulfido-tetrairon; tetra-mu3-sulfido-tetrairon | -; 1([4Fe-4S] cluster); 2([4Fe-4S] cluster); 3([4Fe-4S] cluster); 4([4Fe-4S] cluster) | -; 14, 17, 20, 145; 24, 126, 129, 141; 67, 70, 75, 109; 79, 99, 102, 105 | Keyword & go term; Keyword & binding & cofactor & go term; Keyword & binding & cofactor & go term; Keyword & binding & cofactor & go term; Keyword & binding & cofactor & go term |
| 474 | nadE | P18843 | magnesium(2+) | 1(Mg(2+)) | 52, 165 | Keyword & binding & go term |
| 475 | narW | P19317 | metal cation | - | - | go term |
| 476 | narY | P19318 | iron cation, Fe3S4 iron-sulfur cluster; tetra-mu3-sulfido-tetrairon; tetra-mu3-sulfido-tetrairon; tetra-mu3-sulfido-tetrairon; tri-mu-sulfido-mu3-sulfido-triiron(0) | -; 1([4Fe-4S] cluster); 2([4Fe-4S] cluster); 3([4Fe-4S] cluster); 1([3Fe-4S] cluster) | -; 16, 19, 22, 262; 26, 243, 246, 258; 183, 186, 191, 226; 195, 216, 222 | Keyword & go term, go term; Keyword & binding & cofactor & go term; Keyword & binding & cofactor & go term; Keyword & binding & cofactor & go term; Keyword & binding & cofactor & go term |
| 477 | narZ | P19319 | Mo(=O)-bis(molybdopterin guanine dinucleotide)(4-); iron cation, molybdenum cation; tetra-mu3-sulfido-tetrairon | 1(Mo-bis(molybdopterin guanine dinucleotide)); -; 1([4Fe-4S] cluster) | 223; -; 50, 54, 58, 93 | binding & cofactor; Keyword & go term, Keyword & go term; Keyword & binding & cofactor & go term |
| 478 | pdxA | P19624 | cobalt(2+), magnesium(2+), zinc(2+) | 1(a divalent metal cation) | 166, 211, 266 | Keyword & binding & cofactor & go term, Keyword & binding & cofactor & go term, Keyword & binding & cofactor & go term |
| 479 | eutC | P19636 | cobalt cation; cobamamide | -; 1(adenosylcob(III)alamin) | -; 207, 228, 258 | Keyword; binding & cofactor |
| 480 | entD | P19925 | magnesium(2+) | 1(Mg(2+)) | 107, 109, 152 | Keyword & binding & cofactor & go term |
| 481 | hyaD | P19930 | nickel(2+) | 1(Ni(2+)) | 19, 65, 96 | Keyword & binding & go term |
| 482 | hyaF | P19932 | nickel cation | - | - | Keyword |

Supplementary Table 2: E. coli annotated metal-binding proteome *(continued)*

| # | Gene Name | UniProt ID | Ligand Name | Ligand Identifier | Ligand Position | Source |
|---|---|---|---|---|---|---|
| 483 | parE | P20083 | magnesium(2+); magnesium(2+); manganese(2+), calcium(2+) | 1(Mg(2+)); 2(Mg(2+)); - | 418, 490; 490, 492; - | Keyword & binding & cofactor & go term; Keyword & binding & cofactor & go term; Keyword & cofactor & go term, Keyword & cofactor & go term |
| 484 | bisC | P20099 | Mo(=O)-bis(molybdopterin guanine dinucleotide)(4-); molybdenum cation | 1(Mo-bis(molybdopterin guanine dinucleotide)); - | 148; - | binding & cofactor; Keyword & go term |
| 485 | pepQ | P21165 | manganese(2+); manganese(2+) | 1(Mn(2+)); 2(Mn(2+)) | 257, 339, 384, 423; 246, 257, 423 | Keyword & binding & cofactor & go term; Keyword & binding & cofactor & go term |
| 486 | speA | P21170 | magnesium(2+) | - | - | Keyword & cofactor & go term |
| 487 | katE | P21179 | heme; iron cation | 1(heme); - | 415; - | Keyword & binding & cofactor & go term; Keyword & go term |
| 488 | pncA | P21369 | zinc(2+) | 1(Zn(2+)) | 52, 54, 86 | Keyword & binding & go term |
| 489 | yggF | P21437 | manganese(2+); manganese(2+) | 1(Mn(2+)); 2(Mn(2+)) | 32, 56; 84, 87, 212 | Keyword & binding & cofactor & go term; Keyword & binding & cofactor & go term |
| 490 | rnr | P21499 | magnesium(2+) | - | - | cofactor |
| 491 | rne | P21513 | magnesium(2+); zinc(2+) | 1(Mg(2+)); 1(Zn(2+)) | 303, 346; 404, 407 | Keyword & binding & cofactor & go term; Keyword & binding & cofactor & go term |
| 492 | pdeL | P21514 | magnesium(2+); manganese(2+) | 1(Mg(2+)); - | 141, 200, 232, 262; - | Keyword & binding & cofactor & go term; Keyword & cofactor & go term |
| 493 | pykA | P21599 | magnesium(2+); potassium(1+) | 1(Mg(2+)); 1(K(+)) | 225, 252; 38, 40, 70 | Keyword & binding & cofactor & go term; Keyword & binding & cofactor & go term |
| 494 | ybhA | P21829 | magnesium(2+); manganese(2+), cobalt(2+), zinc(2+) | 1(Mg(2+)); - | 9, 11, 223; - | Keyword & binding & cofactor & go term; Keyword & cofactor & go term, Keyword & cofactor & go term, Keyword & cofactor & go term |
| 495 | cysS | P21888 | zinc(2+) | 1(Zn(2+)) | 28, 209, 234, 238 | Keyword & binding & cofactor & go term |
| 496 | murE | P22188 | magnesium(2+) | - | - | Keyword & cofactor & go term |
| 497 | cysQ | P22255 | magnesium(2+); magnesium(2+) | 1(Mg(2+)); 2(Mg(2+)) | 64, 83, 85; 83, 86, 205 | Keyword & binding & cofactor & go term; Keyword & binding & cofactor & go term |

Supplementary Table 2: E. coli annotated metal-binding proteome *(continued)*

| # | Gene Name | UniProt ID | Ligand Name | Ligand Identifier | Ligand Position | Source |
|---|---|---|---|---|---|---|
| 498 | pckA | P22259 | calcium(2+); magnesium(2+); manganese(2+) | 1(Ca(2+)); -; 1(Mn(2+)) | 149, 150, 152, 283; -; 213, 232, 269 | Keyword & binding & go term; Keyword & go term; Keyword & binding & cofactor & go term |
| 499 | add | P22333 | zinc(2+) | 1(Zn(2+)) | 12, 14, 197, 278 | Keyword & binding & cofactor & go term |
| 500 | ispA | P22939 | magnesium(2+); magnesium(2+) | 1(Mg(2+)); 2(Mg(2+)) | 84, 90; 84, 90 | Keyword & binding & cofactor & go term; Keyword & binding & cofactor & go term |
| 501 | tdk | P23331 | zinc(2+) | 1(Zn(2+)) | 145, 147, 182, 185 | Keyword & binding & go term |
| 502 | hyfA | P23481 | iron cation; tetra-mu3-sulfido-tetrairon; tetra-mu3-sulfido-tetrairon; tetra-mu3-sulfido-tetrairon; tetra-mu3-sulfido-tetrairon | -; 1([4Fe-4S] cluster); 2([4Fe-4S] cluster); 3([4Fe-4S] cluster); 4([4Fe-4S] cluster) | -; 12, 15, 18, 162; 22, 146, 149, 158; 51, 54, 59, 92; 63, 82, 85, 88 | Keyword & go term; Keyword & binding & cofactor & go term; Keyword & binding & cofactor & go term; Keyword & binding & cofactor & go term; Keyword & binding & cofactor & go term |
| 503 | fecI | P23484 | iron cation | - | - | Keyword |
| 504 | fecR | P23485 | iron cation | - | - | Keyword |
| 505 | garL | P23522 | cobalt(2+), iron(2+), manganese(2+); magnesium(2+) | -; 1(Mg(2+)) | -; 153, 179 | Keyword & cofactor & go term, Keyword & cofactor & go term, Keyword & cofactor & go term; Keyword & binding & cofactor & go term |
| 506 | ppsA | P23538 | magnesium(2+) | 1(Mg(2+)) | 680, 704 | Keyword & binding & cofactor & go term |
| 507 | phoQ | P23837 | magnesium(2+); magnesium(2+) | 1(Mg(2+)); 1(a divalent metal cation) | 385, 442; 151, 152 | Keyword & binding & go term; Keyword & binding & go term |
| 508 | yicC | P23839 | divalent metal cation | - | - | cofactor |
| 509 | dppA | P23847 | heme | - | - | go term |
| 510 | trkG | P23849 | potassium(1+) | 1(K(+)) | 114, 115, 223, 224, 320, 321, 437, 438 | Keyword & binding & go term |
| 511 | hemH | P23871 | ferroheme b(2-); iron(2+) | -; 1(Fe cation) | -; 194, 275 | catalytic activity & go term; Keyword & binding & catalytic activity & go term |
| 512 | hipA | P23874 | magnesium(2+) | - | - | go term |
| 513 | fepD | P23876 | iron cation, ferrienterobactin(3-) | - | - | Keyword, go term |
| 514 | fepG | P23877 | iron cation, ferrienterobactin(3-) | - | - | Keyword, go term |
| 515 | fepC | P23878 | iron cation, ferrienterobactin(3-) | - | - | Keyword, catalytic activity & go term |
| 516 | cydC | P23886 | ferroheme b(2-) | - | - | go term |
| 517 | htpX | P23894 | zinc(2+) | 1(Zn(2+)) | 139, 143, 222 | Keyword & binding & cofactor & go term |

Supplementary Table 2: E. coli annotated metal-binding proteome *(continued)*

| # | Gene Name | UniProt ID | Ligand Name | Ligand Identifier | Ligand Position | Source |
|---|---|---|---|---|---|---|
| 518 | argE | P23908 | cobalt(2+); zinc(2+); zinc(2+) | -; 1(Zn(2+)); 2(Zn(2+)) | -; 80, 112, 169; 112, 145, 355 | Keyword & cofactor & go term; Keyword & binding & cofactor & go term; Keyword & binding & cofactor & go term |
| 519 | entS | P24077 | enterobactin(1-) | - | - | go term |
| 520 | dcp | P24171 | calcium cation; zinc(2+) | -; 1(Zn(2+)) | -; 470, 474, 477 | Keyword & go term; Keyword & binding & cofactor & go term |
| 521 | manB | P24175 | magnesium(2+) | 1(Mg(2+)) | 98, 245, 247, 249 | Keyword & binding & cofactor & go term |
| 522 | accC | P24182 | magnesium(2+); magnesium(2+); manganese(2+); manganese(2+) | 1(Mg(2+)); 2(Mg(2+)); 1(Mn(2+)); 2(Mn(2+)) | 276, 288; 288, 290; 276, 288; 288, 290 | Keyword & binding & cofactor & go term; Keyword & binding & cofactor & go term; Keyword & binding & cofactor & go term; Keyword & binding & cofactor & go term |
| 523 | fdnG | P24183 | Mo(=O)-bis(molybdopterin guanine dinucleotide)(4-); molybdenum cation, iron(2+), iron(3+); tetra-mu3-sulfido-tetrairon | 1(Mo-bis(molybdopterin guanine dinucleotide)); -; 1([4Fe-4S] cluster) | 196; -; 50, 53, 57, 92 | binding & cofactor; Keyword & go term, Keyword & go term, Keyword & go term; Keyword & binding & cofactor & go term |
| 524 | hypD | P24192 | iron cation; tetra-mu3-sulfido-tetrairon | 1(Fe cation); - | 41, 69, 72; - | Keyword & binding & go term; Keyword & cofactor & go term |
| 525 | ygiD | P24197 | iron(2+); zinc(2+) | -; 1(Zn(2+)) | -; 22, 57, 177, 234 | Keyword & go term; Keyword & binding & cofactor & go term |
| 526 | mcrA | P24200 | zinc(2+) | - | - | go term |
| 527 | yjiA | P24203 | zinc(2+); zinc(2+); zinc(2+); zinc(2+) | 1(Zn(2+)); 2(Zn(2+)); 3(Zn(2+)); 4(Zn(2+)) | 37, 42, 66; 74, 114; 167, 170, 187; 167, 170, 187 | Keyword & binding & go term; Keyword & binding & go term; Keyword & binding & go term; Keyword & binding & go term |
| 528 | uxuA | P24215 | iron(2+), manganese(2+) | - | - | Keyword & cofactor & go term, Keyword & cofactor & go term |
| 529 | acpS | P24224 | magnesium(2+) | 1(Mg(2+)) | 9, 58 | Keyword & binding & cofactor & go term |
| 530 | hmp | P24232 | ferroheme b(2-); iron cation | 1(heme b); - | 85; - | Keyword & binding & cofactor & go term; Keyword & go term |
| 531 | radA | P24554 | zinc cation | - | - | Keyword & go term |
| 532 | tehB | P25397 | tellurite, methanetelluronate(1-) | - | - | catalytic activity, catalytic activity |

Supplementary Table 2: E. coli annotated metal-binding proteome *(continued)*

| # | Gene Name | UniProt ID | Ligand Name | Ligand Identifier | Ligand Position | Source |
|---|---|---|---|---|---|---|
| 533 | frmA | P25437 | zinc(2+); zinc(2+) | 1(Zn(2+)); 2(Zn(2+)) | 40, 62, 169; 92, 95, 98, 106 | Keyword & binding & cofactor & go term; Keyword & binding & cofactor & go term |
| 534 | acnA | P25516 | iron cation; tetra-mu3-sulfido-tetrairon | -; 1([4Fe-4S] cluster) | -; 435, 501, 504 | Keyword & go term; Keyword & binding & cofactor & go term |
| 535 | hflX | P25519 | magnesium(2+) | 1(Mg(2+)) | 211, 231 | Keyword & binding & cofactor & go term |
| 536 | mnmE | P25522 | magnesium(2+); potassium(1+) | 1(Mg(2+)); 1(K(+)) | 230, 251; 226, 245, 247, 250 | Keyword & binding & go term; Keyword & binding & cofactor & go term |
| 537 | codA | P25524 | iron(2+); manganese(2+); zinc(2+) | 1(Fe(2+)); -; 1(Zn(2+)) | 62, 64, 215, 314; -; 62, 64, 215, 314 | Keyword & binding & cofactor & go term; Keyword & cofactor & go term; Keyword & binding & cofactor & go term |
| 538 | yicR | P25531 | zinc(2+) | 1(Zn(2+)) | 171, 173, 184 | Keyword & binding & go term |
| 539 | yhdE | P25536 | manganese(2+) | - | - | cofactor & go term |
| 540 | ribD | P25539 | zinc(2+) | 1(Zn(2+)) | 50, 75, 84 | Keyword & binding & cofactor & go term |
| 541 | aslA | P25549 | calcium(2+) | 1(Ca(2+)) | 94, 95, 136, 356, 357 | Keyword & binding & cofactor & go term |
| 542 | aslB | P25550 | iron cation; tetra-mu3-sulfido-tetrairon; tetra-mu3-sulfido-tetrairon; tetra-mu3-sulfido-tetrairon | -; 1([4Fe-4S] cluster); 2([4Fe-4S] cluster); 3([4Fe-4S] cluster) | -; 21, 25, 28; 276, 282, 297, 352; 339, 342, 348, 371 | Keyword & go term; Keyword & binding & cofactor & go term; Keyword & binding & cofactor & go term; Keyword & binding & cofactor & go term |
| 543 | metE | P25665 | zinc(2+) | 1(Zn(2+)) | 641, 643, 665, 726 | Keyword & binding & cofactor & go term |
| 544 | malS | P25718 | calcium(2+) | 1(Ca(2+)) | 314, 464 | Keyword & binding & cofactor & go term |
| 545 | waaP | P25741 | magnesium(2+) | - | - | Keyword & cofactor |
| 546 | ligB | P25772 | magnesium(2+), manganese(2+) | - | - | Keyword & cofactor, Keyword & cofactor |
| 547 | preA | P25889 | iron cation; tetra-mu3-sulfido-tetrairon; tetra-mu3-sulfido-tetrairon | -; 1([4Fe-4S] cluster); 2([4Fe-4S] cluster) | -; 344, 347, 350, 388; 354, 378, 381, 384 | Keyword & go term; Keyword & binding & cofactor & go term; Keyword & binding & cofactor & go term |
| 548 | loiP | P25894 | zinc(2+) | 1(Zn(2+)) | 130, 134, 189 | Keyword & binding & cofactor & go term |
| 549 | fepE | P26266 | iron cation, ferrienterobactin(3-) | - | - | Keyword, go term |
| 550 | folK | P26281 | magnesium(2+) | - | - | go term |
| 551 | appB | P26458 | heme, iron cation | - | - | Keyword & cofactor, Keyword & go term |

Supplementary Table 2: E. coli annotated metal-binding proteome *(continued)*

| # | Gene Name | UniProt ID | Ligand Name | Ligand Identifier | Ligand Position | Source |
|---|---|---|---|---|---|---|
| 552 | appC | P26459 | heme; heme; iron cation | 1(heme); 2(heme); - | 19; 186, 393; - | Keyword & binding & cofactor & go term; Keyword & binding & cofactor & go term; Keyword & go term |
| 553 | amyA | P26612 | calcium(2+); sodium(1+) | 1(Ca(2+)); - | 104, 198, 239; - | Keyword & binding & cofactor & go term; Keyword & go term |
| 554 | maeA | P26616 | magnesium(2+), manganese(2+) | 1(a divalent metal cation) | 246, 247, 270 | Keyword & binding & cofactor & go term, Keyword & binding & cofactor & go term |
| 555 | ftsP | P26648 | copper cation | - | - | go term |
| 556 | waaO | P27128 | magnesium(2+) | 1(Mg(2+)) | 131, 133, 265 | Keyword & binding & cofactor & go term |
| 557 | waaJ | P27129 | magnesium(2+) | 1(Mg(2+)) | 130, 132, 264 | Keyword & binding & cofactor & go term |
| 558 | glnD | P27249 | magnesium(2+) | - | - | Keyword & cofactor |
| 559 | ahr | P27250 | zinc(2+); zinc(2+) | 1(Zn(2+)); 2(Zn(2+)) | 38, 63, 152; 96, 99, 102, 110 | Keyword & binding & cofactor & go term; Keyword & binding & cofactor & go term |
| 560 | scpA | P27253 | cobalt cation; cobamamide | -; 1(adenosylcob(III)alamin) | -; 597 | Keyword & go term; binding & cofactor |
| 561 | nadR | P27278 | magnesium(2+) | - | - | go term |
| 562 | dinG | P27296 | iron cation, magnesium(2+); tetra-mu3-sulfido-tetrairon | -; 1([4Fe-4S] cluster) | -; 120, 194, 199, 205 | Keyword & go term, Keyword & cofactor & go term; Keyword & binding & cofactor & go term |
| 563 | prlC | P27298 | zinc(2+) | 1(Zn(2+)) | 469, 473, 476 | Keyword & binding & cofactor & go term |
| 564 | tktA | P27302 | calcium(2+), manganese(2+), cobalt(2+); magnesium(2+) | -; 1(Mg(2+)) | -; 155, 185, 187 | Keyword & cofactor & go term, Keyword & cofactor & go term, Keyword & cofactor & go term; Keyword & binding & cofactor & go term |
| 565 | gluQ | P27305 | zinc(2+) | 1(Zn(2+)) | 111, 113, 125, 129 | Keyword & binding & cofactor & go term |
| 566 | roxA | P27431 | iron(2+) | 1(Fe cation) | 125, 127, 187 | Keyword & binding & cofactor & go term |
| 567 | acs | P27550 | magnesium(2+) | 1(Mg(2+)) | 537, 539, 542 | Keyword & binding & cofactor & go term |
| 568 | cyaY | P27838 | iron(2+), iron(3+) | - | - | Keyword & go term, Keyword & go term |
| 569 | yigL | P27848 | magnesium(2+); manganese(2+), cobalt(2+), zinc(2+) | 1(Mg(2+)); - | 8, 10, 214; - | Keyword & binding & cofactor & go term; Keyword & cofactor & go term, Keyword & cofactor & go term, Keyword & cofactor & go term |
| 570 | tatD | P27859 | magnesium(2+), manganese(2+) | 1(a divalent metal cation) | 91, 127, 152 | Keyword & binding & cofactor & go term, Keyword & binding & cofactor & go term |

Supplementary Table 2: E. coli annotated metal-binding proteome *(continued)*

| # | Gene Name | UniProt ID | Ligand Name | Ligand Identifier | Ligand Position | Source |
|---|---|---|---|---|---|---|
| 571 | bcr | P28246 | sodium(1+) | - | - | go term |
| 572 | qorA | P28304 | zinc(2+) | - | - | go term |
| 573 | fpr | P28861 | di-mu-sulfido-diiron(2+), di-mu-sulfido-diiron(1+) | - | - | catalytic activity & go term, catalytic activity & go term |
| 574 | nrdD | P28903 | zinc(2+) | 1(Zn(2+)) | 644, 647, 662, 665 | Keyword & binding & go term |
| 575 | cydD | P29018 | ferroheme b(2-) | - | - | go term |
| 576 | menC | P29208 | magnesium(2+) | 1(Mg(2+)) | 161, 190, 213 | Keyword & binding & cofactor & go term |
| 577 | pepT | P29745 | zinc(2+); zinc(2+) | 1(Zn(2+)); 2(Zn(2+)) | 78, 140, 196; 140, 174, 379 | Keyword & binding & cofactor & go term; Keyword & binding & cofactor & go term |
| 578 | rnt | P30014 | magnesium(2+); magnesium(2+) | 1(Mg(2+)); 2(Mg(2+)) | 23; 23, 25, 181, 186 | Keyword & binding & cofactor & go term; Keyword & binding & cofactor & go term |
| 579 | leuB | P30125 | magnesium(2+); manganese(2+) | 1(Mg(2+)); - | 227, 251, 255; - | Keyword & binding & cofactor & go term; Keyword & cofactor & go term |
| 580 | hypF | P30131 | zinc(2+) | - | - | Keyword & go term |
| 581 | thiC | P30136 | iron cation; tetra-mu3-sulfido-tetrairon; zinc(2+) | -; 1([4Fe-4S] cluster); 1(Zn(2+)) | -; 581, 584, 589; 437, 501 | Keyword & go term; Keyword & binding & cofactor & go term; Keyword & binding & go term |
| 582 | thiE | P30137 | magnesium(2+) | 1(Mg(2+)) | 70, 89 | Keyword & binding & cofactor & go term |
| 583 | thiF | P30138 | zinc(2+) | 1(Zn(2+)) | 169, 172, 240, 243 | Keyword & binding & cofactor & go term |
| 584 | thiH | P30140 | iron cation; tetra-mu3-sulfido-tetrairon | -; 1([4Fe-4S] cluster) | -; 85, 89, 92 | Keyword & go term; Keyword & binding & cofactor & go term |
| 585 | yaaJ | P30143 | sodium(1+) | - | - | go term |
| 586 | sdaB | P30744 | tetra-mu3-sulfido-tetrairon, iron cation | - | - | Keyword & cofactor & go term, Keyword & go term |
| 587 | moaA | P30745 | iron cation; tetra-mu3-sulfido-tetrairon; tetra-mu3-sulfido-tetrairon | -; 1([4Fe-4S] cluster); 2([4Fe-4S] cluster) | -; 24, 28, 31; 257, 260, 274 | Keyword & go term; Keyword & binding & cofactor & go term; Keyword & binding & cofactor & go term |
| 588 | rnb | P30850 | magnesium(2+) | - | - | cofactor |
| 589 | glnE | P30870 | magnesium(2+) | - | - | Keyword & cofactor & go term |
| 590 | ygiF | P30871 | metal cation | - | - | go term |
| 591 | panB | P31057 | magnesium(2+); manganese(2+), cobalt(2+), zinc(2+) | 1(Mg(2+)); - | 45, 84, 114; - | Keyword & binding & cofactor & go term; Keyword & cofactor & go term, Keyword & cofactor & go term, Keyword & cofactor & go term |
| 592 | kch | P31069 | potassium(1+) | - | - | go term |
| 593 | glmM | P31120 | magnesium(2+) | 1(Mg(2+)) | 102, 241, 243, 245 | Keyword & binding & cofactor & go term |

Supplementary Table 2: E. coli annotated metal-binding proteome *(continued)*

| # | Gene Name | UniProt ID | Ligand Name | Ligand Identifier | Ligand Position | Source |
|---|---|---|---|---|---|---|
| 594 | dgcZ | P31129 | magnesium(2+); zinc(2+) | 1(Mg(2+)); 1(Zn(2+)) | 165, 166, 208; 22, 52, 79, 83 | Keyword & binding & cofactor & go term; Keyword & binding & go term |
| 595 | acrB | P31224 | enterobactin(1-) | - | - | go term |
| 596 | ade | P31441 | manganese(2+), iron(2+) | - | - | Keyword & cofactor & go term, go term |
| 597 | yidJ | P31447 | calcium(2+) | 1(Ca(2+)) | 12, 13, 52, 284, 285 | Keyword & binding & cofactor & go term |
| 598 | yidK | P31448 | sodium(1+) | - | - | Keyword & go term |
| 599 | glvG | P31450 | manganese(2+) | 1(Mn(2+)) | 169, 200 | Keyword & binding & go term |
| 600 | dgoR | P31460 | zinc(2+) | 1(Zn(2+)) | 146, 150, 195 | Keyword & binding & go term |
| 601 | yieH | P31467 | magnesium(2+); manganese(2+), cobalt(2+), zinc(2+) | 1(Mg(2+)); - | 10, 12, 167; - | Keyword & binding & cofactor & go term; Keyword & cofactor & go term, Keyword & cofactor & go term, Keyword & cofactor & go term |
| 602 | efeB | P31545 | ferroheme b(2-); iron(2+) | 1(heme b); - | 236, 237, 238, 329, 334, 335, 336, 347; - | Keyword & binding & catalytic activity & cofactor & go term; Keyword & catalytic activity & go term |
| 603 | hchA | P31658 | zinc(2+) | 1(Zn(2+)) | 86, 91, 123 | Keyword & binding & go term |
| 604 | rpnC | P31665 | magnesium cation | - | - | Keyword |
| 605 | rpnA | P31667 | magnesium(2+) | - | - | Keyword & cofactor |
| 606 | otsA | P31677 | potassium(1+) | - | - | Keyword |
| 607 | otsB | P31678 | magnesium(2+); manganese(2+), cobalt(2+), zinc(2+) | 1(Mg(2+)); - | 20, 22, 198; - | Keyword & binding & cofactor & go term; Keyword & cofactor & go term, Keyword & cofactor & go term, Keyword & cofactor & go term |
| 608 | chaA | P31801 | potassium(1+), sodium(1+), calcium(2+) | - | - | Keyword & catalytic activity & go term, Keyword & catalytic activity & go term, Keyword & catalytic activity |
| 609 | nnr | P31806 | potassium(1+) | 1(K(+)) | 72, 135, 171 | Keyword & binding & cofactor & go term |
| 610 | pqqL | P31828 | zinc(2+) | 1(Zn(2+)) | 80, 84, 160 | Keyword & binding & cofactor & go term |
| 611 | nuoF | P31979 | iron cation; tetra-mu3-sulfido-tetrairon | -; 1([4Fe-4S] cluster) | -; 351, 354, 357, 398 | Keyword & go term; Keyword & binding & cofactor & go term |
| 612 | gmm | P32056 | magnesium(2+); manganese(2+) | 1(Mg(2+)); - | 49, 69, 122; - | Keyword & binding & cofactor & go term; Keyword & go term |
| 613 | hemN | P32131 | iron cation; tetra-mu3-sulfido-tetrairon | -; 1([4Fe-4S] cluster) | -; 62, 66, 69 | Keyword & go term; Keyword & binding & cofactor & go term |
| 614 | frvX | P32153 | divalent metal cation; divalent metal cation | 1(a divalent metal cation); 2(a divalent metal cation) | 61, 175, 228; 175, 206, 316 | Keyword & binding & cofactor & go term; Keyword & binding & cofactor & go term |
| 615 | yiiM | P32157 | molybdenum cation | - | - | go term |
| 616 | menA | P32166 | magnesium(2+) | - | - | go term |

Supplementary Table 2: E. coli annotated metal-binding proteome *(continued)*

| # | Gene Name | UniProt ID | Ligand Name | Ligand Identifier | Ligand Position | Source |
|---|---|---|---|---|---|---|
| 617 | rhaD | P32169 | zinc(2+) | 1(Zn(2+)) | 141, 143, 212 | Keyword & binding & cofactor & go term |
| 618 | rhaA | P32170 | manganese(2+); zinc(2+) | 1(Mn(2+)); 1(Zn(2+)) | 262, 294, 296; 226, 259, 286, 326 | Keyword & binding & cofactor & go term; Keyword & binding & go term |
| 619 | rhaB | P32171 | magnesium(2+) | - | - | Keyword & cofactor |
| 620 | mobA | P32173 | Mo(VI)O2(OH)-molybdopterin cofactor(4-), Mo(VI)-molybdopterin guanine dinucleotide(4-), manganese(2+); magnesium(2+) | -; 1(Mg(2+)) | -; 101 | catalytic activity & go term, catalytic activity & go term, Keyword & cofactor & go term; Keyword & binding & cofactor & go term |
| 621 | fdoG | P32176 | molybdenum cation, Mo(=O)-bis(molybdopterin guanine dinucleotide)(4-), iron(2+), iron(3+); tetra-mu3-sulfido-tetrairon | -; 1([4Fe-4S] cluster) | -; 50, 53, 57, 92 | Keyword & go term, cofactor, Keyword & go term, Keyword & go term; Keyword & binding & cofactor & go term |
| 622 | fdhD | P32177 | Mo(=O)-bis(molybdopterin guanine dinucleotide)(4-) | 1(Mo-bis(molybdopterin guanine dinucleotide)) | 260, 261, 262, 263, 264, 265 | binding |
| 623 | gph | P32662 | magnesium(2+) | 1(Mg(2+)) | 13, 15, 192 | Keyword & binding & cofactor & go term |
| 624 | nudC | P32664 | magnesium(2+), manganese(2+), zinc(2+); magnesium(2+), manganese(2+), zinc(2+); magnesium(2+), manganese(2+), zinc(2+); zinc(2+) | 1(a divalent metal cation); 2(a divalent metal cation); 3(a divalent metal cation); 1(Zn(2+)) | 158, 178, 219; 174; 174, 178, 219; 98, 101, 116, 119 | Keyword & binding & cofactor & go term, Keyword & binding & cofactor & go term, Keyword & binding & cofactor & go term; Keyword & binding & cofactor & go term, Keyword & binding & cofactor & go term, Keyword & binding & cofactor & go term; Keyword & binding & cofactor & go term, Keyword & binding & cofactor & go term, Keyword & binding & cofactor & go term; Keyword & binding & cofactor & go term |
| 625 | ptsA | P32670 | magnesium(2+) | 1(Mg(2+)) | 543, 567 | Keyword & binding & cofactor & go term |
| 626 | pflC | P32675 | iron cation; tetra-mu3-sulfido-tetrairon | -; 1([4Fe-4S] cluster) | -; 47, 51, 54 | Keyword & go term; Keyword & binding & cofactor & go term |
| 627 | yjcE | P32703 | sodium(1+), potassium(1+) | - | - | Keyword & go term, go term |
| 628 | actP | P32705 | sodium(1+), tellurite | - | - | Keyword, go term |
| 629 | nrfE | P32710 | heme | - | - | go term |

Supplementary Table 2: E. coli annotated metal-binding proteome *(continued)*

| # | Gene Name | UniProt ID | Ligand Name | Ligand Identifier | Ligand Position | Source |
|---|---|---|---|---|---|---|
| 630 | nrfF | P32711 | heme; iron cation | 1(heme); - | 44, 47; - | Keyword & binding; Keyword & go term |
| 631 | yjcS | P32717 | zinc(2+); zinc(2+) | 1(Zn(2+)); 2(Zn(2+)) | 180, 182, 291, 310; 184, 185, 310, 355 | Keyword & binding & cofactor & go term; Keyword & binding & cofactor & go term |
| 632 | alsE | P32719 | cobalt(2+), manganese(2+), zinc(2+) | 1(a divalent metal cation) | 30, 32, 63, 173 | Keyword & binding & cofactor & go term, Keyword & binding & cofactor & go term, Keyword & binding & cofactor & go term |
| 633 | rihB | P33022 | calcium(2+) | 1(Ca(2+)) | 11, 16, 124, 240 | Keyword & binding & cofactor & go term |
| 634 | psuG | P33025 | iron(2+), cobalt(2+); manganese(2+) | -; 1(Mn(2+)) | -; 145 | Keyword & cofactor & go term, Keyword & cofactor & go term; Keyword & binding & cofactor & go term |
| 635 | yeiR | P33030 | zinc(2+) | - | - | Keyword & go term |
| 636 | purT | P33221 | magnesium(2+) | 1(Mg(2+)) | 267, 279 | Keyword & binding & go term |
| 637 | yjfC | P33222 | magnesium(2+); magnesium(2+) | 1(Mg(2+)); 2(Mg(2+)) | 103, 116; 116, 118 | Keyword & binding & go term; Keyword & binding & go term |
| 638 | torA | P33225 | Mo(=O)-bis(molybdopterin guanine dinucleotide)(4-); molybdenum cation, iron(2+), iron(3+) | 1(Mo-bis(molybdopterin guanine dinucleotide)); - | 191; - | binding & cofactor; Keyword & go term, Keyword & catalytic activity & go term, Keyword & catalytic activity & go term |
| 639 | torC | P33226 | heme; heme; heme; heme; heme; iron cation | 1(heme); 2(heme); 3(heme); 4(heme); 5(heme); - | 48, 51, 52; 77, 80, 81; 138, 141, 142; 170, 173, 174; 329, 332, 333; - | Keyword & binding & go term; Keyword & binding & go term; Keyword & binding & go term; Keyword & binding & go term; Keyword & binding & go term; Keyword & go term |
| 640 | ralR | P33229 | calcium(2+), magnesium(2+) | - | - | Keyword & cofactor, Keyword & cofactor |
| 641 | yehQ | P33353 | zinc(2+) | - | - | Keyword & go term |
| 642 | tktB | P33570 | calcium(2+), manganese(2+), cobalt(2+); magnesium(2+) | -; 1(Mg(2+)) | -; 154, 184, 186 | Keyword & cofactor & go term, Keyword & cofactor & go term, Keyword & cofactor & go term; Keyword & binding & cofactor & go term |
| 643 | nikA | P33590 | nickel cation, heme, cluster | - | - | Keyword & go term, go term, go term |
| 644 | nikB | P33591 | nickel(2+) | - | - | Keyword & go term |
| 645 | nikD | P33593 | nickel(2+) | - | - | Keyword & catalytic activity & go term |

Supplementary Table 2: E. coli annotated metal-binding proteome *(continued)*

| # | Gene Name | UniProt ID | Ligand Name | Ligand Identifier | Ligand Position | Source |
|---|---|---|---|---|---|---|
| 646 | nikE | P33594 | nickel(2+) | - | - | Keyword & catalytic activity & go term |
| 647 | nuoG | P33602 | di-mu-sulfido-diiron; iron cation; tetra-mu3-sulfido-tetrairon; tetra-mu3-sulfido-tetrairon; tetra-mu3-sulfido-tetrairon | 1([2Fe-2S] cluster); -; 1([4Fe-4S] cluster); 2([4Fe-4S] cluster); 3([4Fe-4S] cluster) | 34, 45, 48, 67; -; 99, 103, 106, 112; 151, 154, 157, 201; 228, 231, 235, 263 | Keyword & binding & cofactor & go term; Keyword & go term; Keyword & binding & cofactor & go term; Keyword & binding & cofactor & go term; Keyword & binding & cofactor & go term |
| 648 | yfiH | P33644 | copper(2+); zinc(2+) | -; 1(Zn(2+)) | -; 71, 107, 124 | Keyword & cofactor & go term; Keyword & binding & cofactor & go term |
| 649 | feoB | P33650 | iron(2+) | - | - | Keyword & go term |
| 650 | radD | P33919 | zinc(2+) | 1(Zn(2+)) | 384, 387, 411, 425 | Keyword & binding & cofactor & go term |
| 651 | ccmF | P33927 | heme | - | - | go term |
| 652 | ccmA | P33931 | ferroheme b(2-) | - | - | catalytic activity & go term |
| 653 | napH | P33934 | iron cation; tetra-mu3-sulfido-tetrairon; tetra-mu3-sulfido-tetrairon | -; 1([4Fe-4S] cluster); 2([4Fe-4S] cluster) | -; 226, 229, 232, 236; 260, 263, 266, 270 | Keyword & go term; Keyword & binding & cofactor & go term; Keyword & binding & cofactor & go term |
| 654 | napA | P33937 | Mo(=O)-bis(molybdopterin guanine dinucleotide)(4-); molybdenum cation, iron(2+), iron(3+); tetra-mu3-sulfido-tetrairon | 1(Mo-bis(molybdopterin guanine dinucleotide)); -; 1([4Fe-4S] cluster) | 83, 150, 175, 179, 212, 213, 214, 215, 216, 217, 218, 219, 243, 244, 245, 246, 247, 262, 263, 264, 372, 376, 482, 508, 509, 531, 558, 718, 719, 720, 721, 722, 723, 724, 725, 726, 727, 802, 819; -; 46, 49, 53, 81 | binding & cofactor; Keyword & go term, Keyword & catalytic activity & go term, Keyword & catalytic activity & go term; Keyword & binding & cofactor & go term |
| 655 | amiA | P36548 | zinc(2+) | - | - | go term |
| 656 | hemF | P36553 | manganese(2+) | 1(Mn(2+)) | 96, 106, 145, 175 | Keyword & binding & cofactor & go term |
| 657 | cobS | P36561 | cobamamide, adenosylcobinamide guanosyl diphosphate(1-), adenosylcobalamin 5'-phosphate(2-), magnesium(2+) | - | - | catalytic activity & go term, catalytic activity & go term, catalytic activity & go term, Keyword & cofactor |

Supplementary Table 2: E. coli annotated metal-binding proteome *(continued)*

| # | Gene Name | UniProt ID | Ligand Name | Ligand Identifier | Ligand Position | Source |
|---|---|---|---|---|---|---|
| 658 | cueO | P36649 | copper(2+), copper(1+); copper(2+), copper(1+); copper(2+), copper(1+); copper(2+), copper(1+); iron(3+), iron(2+) | 1(Cu cation); 2(Cu cation); 3(Cu cation); 4(Cu cation); - | 101, 446; 103, 141, 501; 143, 448, 499; 443, 500, 505; - | Keyword & binding & catalytic activity & cofactor & go term, Keyword & binding & catalytic activity & cofactor & go term; Keyword & binding & catalytic activity & cofactor & go term, Keyword & binding & catalytic activity & cofactor & go term; Keyword & binding & catalytic activity & cofactor & go term, Keyword & binding & catalytic activity & cofactor & go term; Keyword & binding & catalytic activity & cofactor & go term, Keyword & binding & catalytic activity & cofactor & go term; Keyword & go term, Keyword & go term |
| 659 | acnB | P36683 | iron cation; tetra-mu3-sulfido-tetrairon | -; 1([4Fe-4S] cluster) | -; 710, 769, 772 | Keyword & go term; Keyword & binding & cofactor & go term |
| 660 | kdpF | P36937 | potassium(1+) | - | - | Keyword & go term |
| 661 | pgm | P36938 | magnesium(2+) | 1(Mg(2+)) | 146, 304, 306, 308 | Keyword & binding & cofactor & go term |
| 662 | rlmN | P36979 | iron cation, di-mu-sulfido-diiron(2+), di-mu-sulfido-diiron(1+); tetra-mu3-sulfido-tetrairon | -; 1([4Fe-4S] cluster) | -; 125, 129, 132 | Keyword & go term, catalytic activity & go term, catalytic activity & go term; Keyword & binding & cofactor & go term |
| 663 | rlmA | P36999 | zinc(2+) | 1(Zn(2+)) | 5, 8, 21, 25 | Keyword & binding & go term |
| 664 | fluC | P37002 | sodium(1+) | 1(Na(+)) | 75, 78 | Keyword & binding & go term |
| 665 | fbpC | P37009 | iron(3+) | - | - | Keyword & catalytic activity & go term |
| 666 | norR | P37013 | iron(2+) | - | - | go term |
| 667 | btuF | P37028 | cyanocob(III)alamin | 1(cyanocob(III)alamin) | 50, 242, 243, 244, 245, 246 | binding |
| 668 | yaeI | P37049 | divalent metal cation; divalent metal cation | 1(a divalent metal cation); 2(a divalent metal cation) | 56, 58, 88, 211; 88, 120, 209 | Keyword & binding & cofactor & go term; Keyword & binding & cofactor & go term |
| 669 | pepB | P37095 | manganese(2+); manganese(2+) | 1(Mn(2+)); 2(Mn(2+)) | 200, 277, 279; 195, 200, 218, 279 | Keyword & binding & cofactor & go term; Keyword & binding & cofactor & go term |

Supplementary Table 2: E. coli annotated metal-binding proteome *(continued)*

| # | Gene Name | UniProt ID | Ligand Name | Ligand Identifier | Ligand Position | Source |
|---|---|---|---|---|---|---|
| 670 | aegA | P37127 | iron cation; tetra-mu3-sulfido-tetrairon; tetra-mu3-sulfido-tetrairon; tetra-mu3-sulfido-tetrairon; tetra-mu3-sulfido-tetrairon; tetra-mu3-sulfido-tetrairon | -; 1([4Fe-4S] cluster); 2([4Fe-4S] cluster); 3([4Fe-4S] cluster); 4([4Fe-4S] cluster); 5([4Fe-4S] cluster) | -; 12, 15, 18, 137; 22, 121, 124, 133; 56, 59, 64, 97; 68, 87, 90, 93; 227, 230, 236, 240 | Keyword & go term; Keyword & binding & cofactor & go term; Keyword & binding & cofactor & go term; Keyword & binding & cofactor & go term; Keyword & binding & cofactor & go term; Keyword & binding & cofactor & go term |
| 671 | nudK | P37128 | magnesium(2+); magnesium(2+) | 1(Mg(2+)); 2(Mg(2+)) | 85, 104; 100, 104, 151 | Keyword & binding & cofactor & go term; Keyword & binding & cofactor & go term |
| 672 | nrdF | P37146 | iron cation; iron cation; manganese(2+) | 1(Fe cation); 2(Fe cation); - | 67, 98, 101; 98, 158, 192, 195; - | Keyword & binding & cofactor & go term; Keyword & binding & cofactor & go term; Keyword & go term |
| 673 | ptsP | P37177 | magnesium(2+) | 1(Mg(2+)) | 597, 621 | Keyword & binding & cofactor & go term |
| 674 | hybB | P37180 | heme, iron cation | - | - | Keyword, Keyword & go term |
| 675 | hybD | P37182 | nickel(2+) | 1(Ni(2+)) | 16, 62, 93 | Keyword & binding & cofactor & go term |
| 676 | ccp | P37197 | ferroheme c(2-); ferroheme c(2-); ferroheme c(2-); iron(2+), iron(3+) | 1(heme c); 2(heme c); 3(heme c); - | 59, 62, 63, 195; 207, 210, 211, 415; 351, 354, 355, 429; - | Keyword & binding & go term; Keyword & binding & go term; Keyword & binding & go term; Keyword & catalytic activity & go term, Keyword & catalytic activity & go term |
| 677 | dppF | P37313 | heme | - | - | go term |
| 678 | modA | P37329 | molybdate; molybdenum cation, tungsten cation, tungstate | 1(molybdate); - | 36, 63, 149, 176, 194; - | binding & go term; Keyword & go term, Keyword & go term, go term |
| 679 | glcB | P37330 | magnesium(2+) | 1(Mg(2+)) | 427, 455 | Keyword & binding & cofactor & go term |
| 680 | mdtK | P37340 | sodium(1+) | - | - | Keyword |
| 681 | norW | P37596 | iron(2+), iron(3+) | - | - | catalytic activity & go term, catalytic activity & go term |
| 682 | tauD | P37610 | iron(2+) | 1(Fe cation) | 99, 101, 255 | Keyword & binding & cofactor & go term |
| 683 | zntA | P37617 | lead(2+), cadmium(2+); magnesium(2+); zinc(2+); zinc(2+) | -; 1(Mg(2+)); 1(Zn(2+)); 2(Zn(2+)) | -; 436, 438, 628; 58, 59, 62; 392, 394, 714 | Keyword & catalytic activity & go term, Keyword & catalytic activity & go term; Keyword & binding & go term; Keyword & binding & catalytic activity & go term; Keyword & binding & catalytic activity & go term |

Supplementary Table 2: E. coli annotated metal-binding proteome *(continued)*

| # | Gene Name | UniProt ID | Ligand Name | Ligand Identifier | Ligand Position | Source |
|---|---|---|---|---|---|---|
| 684 | acpT | P37623 | magnesium(2+) | - | - | go term |
| 685 | yhjJ | P37648 | divalent metal cation, zinc cation | - | - | Keyword & go term, Keyword & go term |
| 686 | bcsA | P37653 | magnesium(2+) | - | - | cofactor |
| 687 | eptB | P37661 | calcium(2+) | - | - | cofactor |
| 688 | sgbH | P37678 | magnesium(2+) | 1(Mg(2+)) | 33, 62 | Keyword & binding & cofactor & go term |
| 689 | sgbU | P37679 | zinc(2+) | - | - | go term |
| 690 | sgbE | P37680 | zinc(2+) | 1(Zn(2+)) | 76, 95, 97, 171 | Keyword & binding & cofactor & go term |
| 691 | aldB | P37685 | magnesium cation | - | - | Keyword |
| 692 | yiaY | P37686 | iron cation | - | - | Keyword & cofactor & go term |
| 693 | gpmI | P37689 | manganese(2+); manganese(2+) | 1(Mn(2+)); 2(Mn(2+)) | 403, 407, 463; 14, 64, 444, 445 | Keyword & binding & cofactor & go term; Keyword & binding & cofactor & go term |
| 694 | envC | P37690 | metal cation | - | - | go term |
| 695 | rfbA | P37744 | magnesium(2+) | 1(Mg(2+)) | 111, 226 | Keyword & binding & cofactor & go term |
| 696 | rfbD | P37760 | magnesium(2+) | - | - | Keyword & cofactor & go term |
| 697 | mpl | P37773 | magnesium(2+) | - | - | Keyword & cofactor |
| 698 | chbG | P37794 | magnesium(2+) | 1(Mg(2+)) | 61, 125 | Keyword & binding & cofactor & go term |
| 699 | ygcB | P38036 | magnesium(2+) | 1(Mg(2+)) | 75, 160 | Keyword & binding & cofactor & go term |
| 700 | menF | P38051 | magnesium(2+) | 1(Mg(2+)) | 284, 416 | Keyword & binding & cofactor & go term |
| 701 | cusA | P38054 | copper(2+), silver(1+) | - | - | Keyword & go term, go term |
| 702 | dgcE | P38097 | magnesium(2+) | 1(Mg(2+)) | 720, 763 | Keyword & binding & cofactor & go term |
| 703 | rspA | P38104 | magnesium(2+) | - | - | go term |
| 704 | rspB | P38105 | zinc(2+); zinc(2+) | 1(Zn(2+)); 2(Zn(2+)) | 37, 59, 144; 89, 92, 95, 103 | Keyword & binding & cofactor & go term; Keyword & binding & cofactor & go term |
| 705 | fadK | P38135 | magnesium(2+) | - | - | Keyword & cofactor |
| 706 | ydaE | P38394 | zinc(2+) | - | - | go term |
| 707 | ygdG | P38506 | magnesium(2+); potassium(1+) | 1(Mg(2+)); 1(K(+)) | 104; 171, 172, 180, 182, 185 | Keyword & binding & cofactor & go term; Keyword & binding & cofactor & go term |
| 708 | znuA | P39172 | zinc(2+) | 1(Zn(2+)) | 59, 60, 143, 207 | Keyword & binding & go term |
| 709 | lplT | P39196 | sodium(1+) | - | - | go term |
| 710 | epmB | P39280 | iron cation; tetra-mu3-sulfido-tetrairon | -; 1([4Fe-4S] cluster) | -; 120, 124, 127 | Keyword & go term; Keyword & binding & cofactor & go term |
| 711 | rsgA | P39286 | zinc(2+) | 1(Zn(2+)) | 297, 302, 304, 310 | Keyword & binding & cofactor & go term |

Supplementary Table 2: E. coli annotated metal-binding proteome *(continued)*

| # | Gene Name | UniProt ID | Ligand Name | Ligand Identifier | Ligand Position | Source |
|---|---|---|---|---|---|---|
| 712 | queG | P39288 | iron cation, cob(II)alamin; tetra-mu3-sulfido-tetrairon; tetra-mu3-sulfido-tetrairon | -; 1([4Fe-4S] cluster); 2([4Fe-4S] cluster) | -; 193, 196, 199, 253; 203, 219, 246, 249 | Keyword & go term, cofactor; Keyword & binding & cofactor & go term; Keyword & binding & cofactor & go term |
| 713 | ulaG | P39300 | manganese(2+) | - | - | cofactor & go term |
| 714 | ulaD | P39304 | magnesium(2+) | 1(Mg(2+)) | 33, 62 | Keyword & binding & cofactor & go term |
| 715 | ulaF | P39306 | zinc(2+) | 1(Zn(2+)) | 74, 93, 95, 167 | Keyword & binding & cofactor & go term |
| 716 | ytfB | P39310 | metal cation | - | - | go term |
| 717 | idnD | P39346 | zinc(2+); zinc(2+) | 1(Zn(2+)); 2(Zn(2+)) | 40, 65, 153; 93, 96, 99, 107 | Keyword & binding & cofactor & go term; Keyword & binding & cofactor & go term |
| 718 | sgcE | P39362 | cobalt(2+), iron(2+), manganese(2+), zinc(2+) | 1(a divalent metal cation) | 31, 33, 64, 169 | Keyword & binding & cofactor & go term, Keyword & binding & cofactor & go term, Keyword & binding & cofactor & go term, Keyword & binding & cofactor & go term |
| 719 | sgcX | P39366 | divalent metal cation; divalent metal cation | 1(a divalent metal cation); 2(a divalent metal cation) | 67, 180, 235; 180, 213, 329 | Keyword & binding & cofactor & go term; Keyword & binding & cofactor & go term |
| 720 | iadA | P39377 | cobalt(2+); zinc(2+); zinc(2+) | -; 1(Zn(2+)); 2(Zn(2+)) | -; 68, 70, 162, 285; 162, 201, 230 | Keyword & cofactor & go term; Keyword & binding & cofactor & go term; Keyword & binding & cofactor & go term |
| 721 | yjiL | P39383 | iron cation; tetra-mu3-sulfido-tetrairon | -; 1([4Fe-4S] cluster) | -; 122, 160 | Keyword & go term; Keyword & binding & cofactor & go term |
| 722 | mdtM | P39386 | potassium(1+), sodium(1+) | - | - | Keyword & catalytic activity & go term, Keyword & catalytic activity & go term |
| 723 | lgoD | P39400 | zinc(2+); zinc(2+) | 1(Zn(2+)); 2(Zn(2+)) | 40, 65, 146; 92, 95, 98, 106 | Keyword & binding & cofactor & go term; Keyword & binding & cofactor & go term |
| 724 | fhuF | P39405 | di-mu-sulfido-diiron; iron(3+), iron(2+) | 1([2Fe-2S] cluster); - | 244, 245, 256, 259; - | Keyword & binding & cofactor & go term; Keyword & go term, Keyword & go term |
| 725 | rsmC | P39406 | magnesium(2+) | - | - | Keyword & cofactor |
| 726 | yjjV | P39408 | divalent metal cation; divalent metal cation | 1(a divalent metal cation); 2(a divalent metal cation) | 9, 11, 97, 207; 97, 133, 157 | Keyword & binding & cofactor & go term; Keyword & binding & cofactor & go term |

Supplementary Table 2: E. coli annotated metal-binding proteome *(continued)*

| # | Gene Name | UniProt ID | Ligand Name | Ligand Identifier | Ligand Position | Source |
|---|---|---|---|---|---|---|
| 727 | yjjW | P39409 | iron cation; tetra-mu3-sulfido-tetrairon; tetra-mu3-sulfido-tetrairon; tetra-mu3-sulfido-tetrairon | -; 1([4Fe-4S] cluster); 2([4Fe-4S] cluster); 3([4Fe-4S] cluster) | -; 31, 35, 38; 47, 50, 53, 86; 57, 76, 79, 82 | Keyword & go term; Keyword & binding & cofactor & go term; Keyword & binding & cofactor & go term; Keyword & binding & cofactor & go term |
| 728 | yjjX | P39411 | magnesium(2+); manganese(2+) | 1(Mg(2+)); - | 38, 68; - | Keyword & binding & cofactor & go term; Keyword & cofactor & go term |
| 729 | adhP | P39451 | zinc(2+); zinc(2+) | 1(Zn(2+)); 2(Zn(2+)) | 37, 58, 145; 89, 92, 95, 103 | Keyword & binding & cofactor & go term; Keyword & binding & cofactor & go term |
| 730 | garD | P39829 | iron(2+) | - | - | Keyword & cofactor & go term |
| 731 | znuB | P39832 | zinc(2+) | - | - | Keyword & go term |
| 732 | sprT | P39902 | zinc(2+) | 1(Zn(2+)) | 78, 82 | Keyword & binding & cofactor & go term |
| 733 | pdxK | P40191 | magnesium(2+); zinc(2+) | 1(Mg(2+)); - | 136, 162; - | Keyword & binding & cofactor & go term; Keyword & cofactor & go term |
| 734 | nlpE | P40710 | copper cation | - | - | Keyword |
| 735 | ybiI | P41039 | zinc(2+) | - | - | Keyword & go term |
| 736 | mltB | P41052 | calcium(2+), sodium(1+) | - | - | go term, go term |
| 737 | traR | P41065 | zinc(2+) | - | - | Keyword & go term |
| 738 | rihA | P41409 | calcium(2+) | - | - | go term |
| 739 | gspF | P41441 | calcium(2+) | 1(Ca(2+)) | 90, 144, 148 | Keyword & binding & go term |
| 740 | ygjK | P42592 | calcium(2+) | 1(Ca(2+)) | 454, 456, 458, 460, 462, 572 | Keyword & binding & go term |
| 741 | fadH | P42593 | iron cation; tetra-mu3-sulfido-tetrairon | -; 1([4Fe-4S] cluster) | -; 335, 338, 342, 354 | Keyword & go term; Keyword & binding & cofactor & go term |
| 742 | uxaA | P42604 | iron(2+), manganese(2+) | - | - | Keyword & cofactor & go term, Keyword & cofactor |
| 743 | tdcG | P42630 | tetra-mu3-sulfido-tetrairon, iron cation | - | - | Keyword & cofactor & go term, Keyword & go term |
| 744 | obgE | P42641 | magnesium(2+) | 1(Mg(2+)) | 173, 193 | Keyword & binding & cofactor & go term |
| 745 | nudL | P43337 | magnesium(2+); manganese(2+) | 1(Mg(2+)); - | 83, 87; - | Keyword & binding & cofactor & go term; Keyword & cofactor & go term |
| 746 | lpxH | P43341 | manganese(2+); manganese(2+) | 1(Mn(2+)); 2(Mn(2+)) | 8, 10, 41, 197; 41, 79, 114, 195 | Keyword & binding & cofactor & go term; Keyword & binding & cofactor & go term |
| 747 | ycaL | P43674 | zinc(2+) | 1(Zn(2+)) | 134, 138, 193 | Keyword & binding & cofactor & go term |
| 748 | yrbG | P45394 | calcium(2+), potassium(1+), sodium(1+) | - | - | go term, go term, go term |

Supplementary Table 2: E. coli annotated metal-binding proteome *(continued)*

| # | Gene Name | UniProt ID | Ligand Name | Ligand Identifier | Ligand Position | Source |
|---|---|---|---|---|---|---|
| 749 | kdsD | P45395 | zinc(2+) | 1(Zn(2+)) | 82 | Keyword & binding & go term |
| 750 | nanQ | P45424 | zinc(2+) | - | - | Keyword & cofactor & go term |
| 751 | nanK | P45425 | zinc(2+) | 1(Zn(2+)) | 156, 166, 168, 173 | Keyword & binding & go term |
| 752 | ubiV | P45475 | iron cation; tetra-mu3-sulfido-tetrairon | -; 1([4Fe-4S] cluster) | -; 39, 180, 193, 197 | Keyword & go term; Keyword & binding & cofactor & go term |
| 753 | kefB | P45522 | potassium(1+) | - | - | Keyword & go term |
| 754 | ubiU | P45527 | iron cation; tetra-mu3-sulfido-tetrairon | -; 1([4Fe-4S] cluster) | -; 169, 176, 193, 232 | Keyword & go term; Keyword & binding & cofactor & go term |
| 755 | frlC | P45541 | cobalt(2+), nickel(2+) | 1(a divalent metal cation) | 148, 181, 207, 247 | Keyword & binding & cofactor & go term, Keyword & binding & cofactor & go term |
| 756 | php | P45548 | zinc(2+); zinc(2+) | 1(Zn(2+)); 2(Zn(2+)) | 12, 14, 125, 243; 125, 158, 186 | Keyword & binding & cofactor & go term; Keyword & binding & cofactor & go term |
| 757 | yhfW | P45549 | magnesium(2+); manganese(2+) | -; 1(Mn(2+)) | -; 10, 302, 338, 339, 350 | Keyword & go term; Keyword & binding & cofactor & go term |
| 758 | dxr | P45568 | magnesium(2+), cobalt(2+); manganese(2+) | -; 1(Mn(2+)) | -; 150, 152, 231 | Keyword & cofactor & go term, Keyword & cofactor & go term; Keyword & binding & cofactor & go term |
| 759 | luxS | P45578 | iron cation | 1(Fe cation) | 54, 58, 128 | Keyword & binding & cofactor & go term |
| 760 | hcxA | P45579 | zinc(2+) | 1(Zn(2+)) | 173, 257, 274 | Keyword & binding & cofactor & go term |
| 761 | gspE | P45759 | zinc(2+) | 1(Zn(2+)) | 382, 385, 415, 418 | Keyword & binding & cofactor & go term |
| 762 | yrdD | P45771 | iron cation, zinc(2+) | - | - | go term, go term |
| 763 | nudE | P45799 | magnesium(2+) | 1(a divalent metal cation) | 95, 99 | Keyword & binding & cofactor & go term |
| 764 | dgcN | P46139 | magnesium(2+) | 1(Mg(2+)) | 286, 329 | Keyword & binding & cofactor & go term |
| 765 | rtcB | P46850 | manganese(2+); manganese(2+) | 1(Mn(2+)); 2(Mn(2+)) | 75, 78, 168; 78, 185, 281 | Keyword & binding & cofactor & go term; Keyword & binding & cofactor & go term |
| 766 | yhhW | P46852 | copper(2+), iron(2+), zinc(2+), cobalt(2+) | 1(a divalent metal cation) | 57, 59, 101, 103 | Keyword & binding & go term, Keyword & binding & cofactor & go term, Keyword & binding & cofactor & go term, Keyword & binding & cofactor & go term |

Supplementary Table 2: E. coli annotated metal-binding proteome *(continued)*

| # | Gene Name | UniProt ID | Ligand Name | Ligand Identifier | Ligand Position | Source |
|---|---|---|---|---|---|---|
| 767 | tynA | P46883 | calcium(2+); calcium(2+); copper cation; manganese(2+); zinc(2+) | 1(Ca(2+)); 2(Ca(2+)); 1(Cu cation); 1(Mn(2+)); - | 563, 564, 565, 708, 709; 603, 697, 700, 702, 709; 554, 556, 719; 563, 565, 708; - | Keyword & binding & cofactor & go term; Keyword & binding & cofactor & go term; Keyword & binding & cofactor & go term; Keyword & binding & cofactor & go term; Keyword & cofactor & go term |
| 768 | cof | P46891 | magnesium(2+); manganese(2+), cobalt(2+), zinc(2+) | 1(Mg(2+)); - | 8, 10, 212; - | Keyword & binding & cofactor & go term; Keyword & cofactor & go term, Keyword & cofactor & go term, Keyword & cofactor & go term |
| 769 | torZ | P46923 | Mo(=O)-bis(molybdopterin guanine dinucleotide)(4-); molybdenum cation, iron(2+), iron(3+) | 1(Mo-bis(molybdopterin guanine dinucleotide)); - | 176; - | binding & cofactor; Keyword & go term, Keyword & catalytic activity & go term, Keyword & catalytic activity & go term |
| 770 | mlc | P50456 | zinc(2+) | 1(Zn(2+)) | 247, 257, 259, 264 | Keyword & binding & go term |
| 771 | nei | P50465 | zinc(2+) | - | - | Keyword & cofactor & go term |
| 772 | mhpE | P51020 | manganese(2+) | 1(Mn(2+)) | 15, 197, 199 | Keyword & binding & go term |
| 773 | ycjG | P51981 | magnesium(2+) | 1(Mg(2+)) | 176, 202, 225 | Keyword & binding & cofactor & go term |
| 774 | torY | P52005 | heme; heme; heme; heme; heme; iron cation | 1(heme); 2(heme); 3(heme); 4(heme); 5(heme); - | 39, 42, 43; 68, 71, 72; 129, 132, 133; 161, 164, 165; 313, 316, 317; - | Keyword & binding & go term; Keyword & binding & go term; Keyword & binding & go term; Keyword & binding & go term; Keyword & binding & go term; Keyword & go term |
| 775 | nudI | P52006 | magnesium(2+) | - | - | Keyword & cofactor & go term |
| 776 | rdgB | P52061 | magnesium(2+); manganese(2+), nickel(2+) | 1(Mg(2+)); - | 40, 69; - | Keyword & binding & cofactor & go term; Keyword & cofactor & go term, Keyword & cofactor & go term |
| 777 | hemW | P52062 | heme, iron cation; tetra-mu3-sulfido-tetrairon | -; 1([4Fe-4S] cluster) | -; 16, 20, 23 | Keyword & go term, Keyword & go term; Keyword & binding & cofactor & go term |
| 778 | glcF | P52074 | iron cation; tetra-mu3-sulfido-tetrairon; tetra-mu3-sulfido-tetrairon | -; 1([4Fe-4S] cluster); 2([4Fe-4S] cluster) | -; 25, 28, 31, 85; 35, 75, 78, 81 | Keyword & go term; Keyword & binding & cofactor & go term; Keyword & binding & cofactor & go term |

Supplementary Table 2: E. coli annotated metal-binding proteome *(continued)*

| # | Gene Name | UniProt ID | Ligand Name | Ligand Identifier | Ligand Position | Source |
|---|---|---|---|---|---|---|
| 779 | cobC | P52086 | cobamamide, adenosylcobalamin 5'-phosphate(2-) | - | - | catalytic activity, catalytic activity |
| 780 | yaeR | P52096 | divalent metal cation | 1(a divalent metal cation) | 9, 57, 78, 125 | Keyword & binding & go term |
| 781 | yfhL | P52102 | iron cation; tetra-mu3-sulfido-tetrairon; tetra-mu3-sulfido-tetrairon | -; 1([4Fe-4S] cluster); 2([4Fe-4S] cluster) | -; 9, 12, 15, 54; 19, 38, 41, 50 | Keyword & go term; Keyword & binding & cofactor & go term; Keyword & binding & cofactor & go term |
| 782 | yfjY | P52140 | zinc(2+) | 1(Zn(2+)) | 109, 111, 122 | Keyword & binding & go term |
| 783 | yccM | P52636 | iron cation; tetra-mu3-sulfido-tetrairon; tetra-mu3-sulfido-tetrairon | -; 1([4Fe-4S] cluster); 2([4Fe-4S] cluster) | -; 251, 254, 257, 288; 261, 278, 281, 284 | Keyword & go term; Keyword & binding & go term; Keyword & binding & go term |
| 784 | ydbK | P52647 | iron cation; tetra-mu3-sulfido-tetrairon; tetra-mu3-sulfido-tetrairon; tetra-mu3-sulfido-tetrairon | -; 1([4Fe-4S] cluster); 2([4Fe-4S] cluster); 3([4Fe-4S] cluster) | -; 689, 692, 695, 755; 699, 745, 748, 751; 819, 822, 847, 1071 | Keyword & go term; Keyword & binding & cofactor & go term; Keyword & binding & cofactor & go term; Keyword & binding & cofactor & go term |
| 785 | mngB | P54746 | divalent metal cation | 1(a divalent metal cation) | 14, 16, 126, 338 | Keyword & binding & cofactor & go term |
| 786 | rlmD | P55135 | iron cation; tetra-mu3-sulfido-tetrairon | -; 1([4Fe-4S] cluster) | -; 81, 87, 90, 162 | Keyword & go term; Keyword & binding & go term |
| 787 | pphA | P55798 | manganese(2+); manganese(2+) | 1(Mn(2+)); 2(Mn(2+)) | 24, 26, 53; 53, 79, 187 | Keyword & binding & cofactor & go term; Keyword & binding & cofactor & go term |
| 788 | pphB | P55799 | manganese(2+); manganese(2+) | 1(Mn(2+)); 2(Mn(2+)) | 22, 24, 51; 51, 77, 187 | Keyword & binding & cofactor & go term; Keyword & binding & cofactor & go term |
| 789 | ysaA | P56256 | iron cation; tetra-mu3-sulfido-tetrairon; tetra-mu3-sulfido-tetrairon; tetra-mu3-sulfido-tetrairon; tetra-mu3-sulfido-tetrairon | -; 1([4Fe-4S] cluster); 2([4Fe-4S] cluster); 3([4Fe-4S] cluster); 4([4Fe-4S] cluster) | -; 12, 15, 18, 134; 22, 118, 121, 130; 58, 61, 66, 99; 70, 89, 92, 95 | Keyword & go term; Keyword & binding & go term; Keyword & binding & go term; Keyword & binding & go term; Keyword & binding & go term |
| 790 | mukF | P60293 | calcium(2+) | - | - | Keyword & go term |
| 791 | rplB | P60422 | zinc(2+) | - | - | go term |
| 792 | ispU | P60472 | magnesium(2+) | 1(Mg(2+)) | 26, 199, 213 | Keyword & binding & cofactor & go term |
| 793 | guaC | P60560 | potassium(1+) | 1(K(+)) | 181, 183 | Keyword & binding & go term |
| 794 | speB | P60651 | manganese(2+); manganese(2+) | 1(Mn(2+)); 2(Mn(2+)) | 126, 149, 153, 230; 149, 151, 230, 232 | Keyword & binding & cofactor & go term; Keyword & binding & cofactor & go term |

Supplementary Table 2: E. coli annotated metal-binding proteome *(continued)*

| # | Gene Name | UniProt ID | Ligand Name | Ligand Identifier | Ligand Position | Source |
|---|---|---|---|---|---|---|
| 795 | lipA | P60716 | iron(3+), di-mu-sulfido-diiron(2+), di-mu-sulfido-diiron(1+); tetra-mu3-sulfido-tetrairon(2+); tetra-mu3-sulfido-tetrairon(2+) | -; 1([4Fe-4S] cluster); 2([4Fe-4S] cluster) | -; 68, 73, 79, 308; 94, 98, 101 | Keyword & catalytic activity & go term, catalytic activity & go term, catalytic activity & go term; Keyword & binding & catalytic activity & cofactor & go term; Keyword & binding & catalytic activity & cofactor & go term |
| 796 | hisG | P60757 | magnesium(2+) | - | - | Keyword & cofactor & go term |
| 797 | can | P61517 | zinc(2+) | 1(Zn(2+)) | 42, 44, 98, 101 | Keyword & binding & cofactor & go term |
| 798 | rffH | P61887 | magnesium(2+) | 1(Mg(2+)) | 108, 223 | Keyword & binding & cofactor & go term |
| 799 | ispF | P62617 | zinc(2+), manganese(2+) | 1(a divalent metal cation) | 8, 10, 42 | Keyword & binding & cofactor & go term, Keyword & binding & cofactor & go term |
| 800 | ispG | P62620 | iron cation, di-mu-sulfido-diiron(1+), di-mu-sulfido-diiron(2+); tetra-mu3-sulfido-tetrairon | -; 1([4Fe-4S] cluster) | -; 270, 273, 305, 312 | Keyword & go term, go term, go term; Keyword & binding & cofactor & go term |
| 801 | ispH | P62623 | iron cation, di-mu-sulfido-diiron(2+), di-mu-sulfido-diiron(1+), Fe3S4 iron-sulfur cluster; tetra-mu3-sulfido-tetrairon | -; 1([4Fe-4S] cluster) | -; 12, 96, 197 | Keyword & go term, catalytic activity, catalytic activity, go term; Keyword & binding & cofactor & go term |
| 802 | nfuA | P63020 | iron cation; tetra-mu3-sulfido-tetrairon | -; 1([4Fe-4S] cluster) | -; 149, 152 | Keyword & go term; Keyword & binding & cofactor & go term |
| 803 | kup | P63183 | potassium(1+) | - | - | Keyword & catalytic activity & go term |
| 804 | gmhA | P63224 | zinc(2+) | 1(Zn(2+)) | 61, 65, 172, 180 | Keyword & binding & cofactor & go term |
| 805 | gmhB | P63228 | magnesium(2+); manganese(2+), cobalt(2+); zinc(2+) | 1(Mg(2+)); -; 1(Zn(2+)) | 11, 13, 136, 137; -; 92, 94, 107, 109 | Keyword & binding & cofactor & go term; Keyword & cofactor & go term, Keyword & cofactor & go term; Keyword & binding & cofactor & go term |
| 806 | zntB | P64423 | zinc(2+), magnesium(2+), cobalt(2+) | - | - | Keyword & catalytic activity & go term, go term, go term |
| 807 | ypjD | P64432 | heme | - | - | go term |
| 808 | rcnR | P64530 | metal cation | - | - | go term |
| 809 | rcnB | P64534 | copper cation | - | - | go term |

Supplementary Table 2: E. coli annotated metal-binding proteome *(continued)*

| # | Gene Name | UniProt ID | Ligand Name | Ligand Identifier | Ligand Position | Source |
|---|---|---|---|---|---|---|
| 810 | queE | P64554 | iron cation;<br>magnesium(2+);<br>tetra-mu3-sulfido-tetrairon | -;<br>1(Mg(2+));<br>1([4Fe-4S] cluster) | -;<br>40;<br>31, 35, 38 | Keyword & go term;<br>Keyword & binding & cofactor & go term;<br>Keyword & binding & cofactor & go term |
| 811 | yrfG | P64636 | magnesium(2+);<br>manganese(2+), cobalt(2+), zinc(2+) | 1(Mg(2+));<br>- | 9, 11, 174;<br>- | Keyword & binding & cofactor & go term;<br>Keyword & cofactor & go term, Keyword & cofactor & go term, Keyword & cofactor & go term |
| 812 | feoC | P64638 | Fe4S4 iron-sulfur cluster;<br>iron cation | 1(iron-sulfur cluster);<br>- | 56, 61, 64, 70;<br>- | binding & go term;<br>Keyword & go term |
| 813 | yfcD | P65556 | magnesium(2+) | 1(Mg(2+)) | 88, 92 | Keyword & binding & cofactor & go term |
| 814 | eutT | P65643 | cob(II)alamin, cobamamide, adenosylcobinamide, cob(II)inamide;<br>divalent metal cation | -;<br>1(a divalent metal cation) | -;<br>80, 83 | catalytic activity, catalytic activity, catalytic activity, catalytic activity;<br>Keyword & binding & cofactor & go term |
| 815 | ygeY | P65807 | cobalt(2+);<br>zinc(2+);<br>zinc(2+) | -;<br>1(Zn(2+));<br>2(Zn(2+)) | -;<br>81, 114, 174;<br>114, 149, 374 | Keyword & cofactor & go term;<br>Keyword & binding & cofactor & go term;<br>Keyword & binding & cofactor & go term |
| 816 | queD | P65870 | zinc(2+) | 1(Zn(2+)) | 16, 31, 33 | Keyword & binding & cofactor & go term |
| 817 | bepA | P66948 | zinc(2+) | 1(Zn(2+)) | 136, 140, 201 | Keyword & binding & cofactor & go term |
| 818 | yfcE | P67095 | manganese(2+);<br>manganese(2+) | 1(Mn(2+));<br>2(Mn(2+)) | 9, 11, 37, 129;<br>37, 73, 105, 127 | Keyword & binding & cofactor & go term;<br>Keyword & binding & cofactor & go term |
| 819 | cutC | P67826 | copper cation | - | - | go term |
| 820 | tadA | P68398 | zinc(2+) | 1(Zn(2+)) | 57, 87, 90 | Keyword & binding & cofactor & go term |
| 821 | fixX | P68646 | tetra-mu3-sulfido-tetrairon, iron cation | - | - | Keyword & go term, Keyword & go term |
| 822 | ybcO | P68661 | zinc(2+) | 1(Zn(2+)) | 13, 21, 54, 57 | Keyword & binding & go term |
| 823 | nfi | P68739 | magnesium(2+) | 1(Mg(2+)) | 35, 103 | Keyword & binding & cofactor & go term |
| 824 | pepA | P68767 | manganese(2+);<br>manganese(2+) | 1(Mn(2+));<br>2(Mn(2+)) | 275, 352, 354;<br>270, 275, 293, 354 | Keyword & binding & cofactor & go term;<br>Keyword & binding & cofactor & go term |
| 825 | sdhC | P69054 | heme;<br>iron cation | 1(heme);<br>- | 84;<br>- | Keyword & binding & cofactor & go term;<br>Keyword & go term |

Supplementary Table 2: E. coli annotated metal-binding proteome *(continued)*

| # | Gene Name | UniProt ID | Ligand Name | Ligand Identifier | Ligand Position | Source |
|---|---|---|---|---|---|---|
| 826 | fieF | P69380 | cadmium(2+);<br>iron(2+);<br>zinc(2+);<br>zinc(2+);<br>zinc(2+);<br>zinc(2+) | 1(Cd(2+));<br>-;<br>1(Zn(2+));<br>2(Zn(2+));<br>3(Zn(2+));<br>4(Zn(2+)) | 45, 49, 153, 157;<br>-;<br>45, 49, 153, 157;<br>68, 71, 75;<br>232, 248, 285;<br>261, 283, 285 | Keyword & binding & catalytic activity & go term;<br>Keyword & catalytic activity & go term;<br>Keyword & binding & catalytic activity & go term;<br>Keyword & binding & catalytic activity & go term;<br>Keyword & binding & catalytic activity & go term;<br>Keyword & binding & catalytic activity & go term |
| 827 | adk | P69441 | magnesium(2+) | - | - | go term |
| 828 | fadD | P69451 | magnesium(2+) | - | - | Keyword & cofactor |
| 829 | cutA | P69488 | copper cation | 1(Cu cation) | 16, 83, 84 | Keyword & binding & cofactor & go term |
| 830 | ccmE | P69490 | heme;<br>iron cation | 1(heme);<br>- | 130, 134;<br>- | Keyword & binding & go term;<br>Keyword & go term |
| 831 | apt | P69503 | magnesium(2+) | - | - | go term |
| 832 | ytfE | P69506 | iron cation | - | - | Keyword & go term |
| 833 | hyaA | P69739 | iron cation, di-mu-sulfido-diiron(2+), di-mu-sulfido-diiron(1+), Fe3S4 iron-sulfur cluster;<br>tetra-mu3-sulfido-tetrairon;<br>tetra-mu3-sulfido-tetrairon;<br>tri-mu-sulfido-mu3-sulfido-triiron(0) | -;<br>1([4Fe-4S] cluster);<br>2([4Fe-4S] cluster);<br>1([3Fe-4S] cluster) | -;<br>62, 65, 160, 194;<br>232, 235, 260, 266;<br>275, 294, 297 | Keyword & go term, go term, go term, go term;<br>Keyword & binding & cofactor & go term;<br>Keyword & binding & cofactor & go term;<br>Keyword & binding & cofactor & go term |
| 834 | hybO | P69741 | iron cation, di-mu-sulfido-diiron(2+), di-mu-sulfido-diiron(1+), Fe3S4 iron-sulfur cluster;<br>tetra-mu3-sulfido-tetrairon;<br>tetra-mu3-sulfido-tetrairon;<br>tri-mu-sulfido-mu3-sulfido-triiron(0) | -;<br>1([4Fe-4S] cluster);<br>2([4Fe-4S] cluster);<br>1([3Fe-4S] cluster) | -;<br>59, 62, 157, 191;<br>229, 232, 257, 263;<br>272, 292, 295 | Keyword & go term, go term, go term, go term;<br>Keyword & binding & cofactor & go term;<br>Keyword & binding & cofactor & go term;<br>Keyword & binding & cofactor & go term |
| 835 | crr | P69783 | zinc(2+) | 1(Zn(2+)) | 76, 91 | Keyword & binding & cofactor & go term |
| 836 | chbA | P69791 | magnesium(2+) | - | - | cofactor |
| 837 | fucI | P69922 | manganese(2+) | 1(Mn(2+)) | 337, 361, 528 | Keyword & binding & cofactor & go term |
| 838 | nrdB | P69924 | iron cation;<br>iron cation | 1(Fe cation);<br>2(Fe cation) | 85, 116, 119;<br>116, 205, 239, 242 | Keyword & binding & cofactor & go term;<br>Keyword & binding & cofactor & go term |
| 839 | yahK | P75691 | zinc(2+);<br>zinc(2+) | 1(Zn(2+));<br>2(Zn(2+)) | 40, 62, 158;<br>93, 96, 99, 107 | Keyword & binding & cofactor & go term;<br>Keyword & binding & cofactor & go term |

Supplementary Table 2: E. coli annotated metal-binding proteome *(continued)*

| # | Gene Name | UniProt ID | Ligand Name | Ligand Identifier | Ligand Position | Source |
|---|---|---|---|---|---|---|
| 840 | allE | P75713 | manganese(2+) | 1(Mn(2+)) | 196, 198, 202, 236 | Keyword & binding & cofactor & go term |
| 841 | zitB | P75757 | zinc(2+) | - | - | Keyword & go term |
| 842 | ybhJ | P75764 | iron cation; tetra-mu3-sulfido-tetrairon | -; 1([4Fe-4S] cluster) | -; 360, 421, 424 | Keyword & go term; binding & cofactor & go term |
| 843 | ybiX | P75779 | iron(2+) | 1(Fe cation) | 96, 98, 158 | Keyword & binding & cofactor & go term |
| 844 | fiu | P75780 | iron cation, siderophore | - | - | Keyword, go term |
| 845 | ybiV | P75792 | magnesium(2+); manganese(2+), cobalt(2+), zinc(2+) | 1(Mg(2+)); - | 9, 11, 215, 216; - | Keyword & binding & cofactor & go term; Keyword & cofactor & go term, Keyword & cofactor & go term, Keyword & cofactor & go term |
| 846 | ybiY | P75794 | iron cation; tetra-mu3-sulfido-tetrairon | -; 1([4Fe-4S] cluster) | -; 25, 29, 32 | Keyword & go term; Keyword & binding & cofactor & go term |
| 847 | dgcI | P75801 | magnesium(2+) | 1(Mg(2+)) | 327, 371 | Keyword & binding & cofactor & go term |
| 848 | yliI | P75804 | calcium(2+) | 1(Ca(2+)) | 240, 250 | Keyword & binding & cofactor & go term |
| 849 | ybjI | P75809 | magnesium(2+); manganese(2+), cobalt(2+), zinc(2+) | 1(Mg(2+)); - | 9, 11, 215; - | Keyword & binding & cofactor & go term; Keyword & cofactor & go term, Keyword & cofactor & go term, Keyword & cofactor & go term |
| 850 | rlmC | P75817 | iron cation; tetra-mu3-sulfido-tetrairon | -; 1([4Fe-4S] cluster) | -; 3, 11, 14, 87 | Keyword & go term; Keyword & binding & go term |
| 851 | amiD | P75820 | zinc(2+) | 1(Zn(2+)) | 50, 166, 176 | Keyword & binding & cofactor & go term |
| 852 | hcr | P75824 | di-mu-sulfido-diiron; iron cation | 1([2Fe-2S] cluster); - | 273, 278, 281, 311; - | Keyword & binding & cofactor & go term; Keyword & go term |
| 853 | hcp | P75825 | Fe4S2O2 iron-sulfur-oxygen cluster; di-mu-sulfido-diiron; iron cation | 1(hybrid [4Fe-2O-2S] cluster); 1([2Fe-2S] cluster); - | 249, 273, 317, 405, 433, 458, 492, 494; 3, 6, 18, 25; - | binding & cofactor; Keyword & binding & cofactor & go term; Keyword & go term |
| 854 | ycaO | P75838 | magnesium(2+) | - | - | go term |
| 855 | gloC | P75849 | zinc(2+); zinc(2+) | 1(Zn(2+)); 2(Zn(2+)) | 56, 58, 132, 151; 60, 61, 151, 192 | Keyword & binding & cofactor & go term; Keyword & binding & cofactor & go term |
| 856 | ycbX | P75863 | molybdenum cation, Fe2S2 iron-sulfur cluster | - | - | go term, go term |
| 857 | efeU | P75901 | iron(2+) | - | - | go term |
| 858 | dgcT | P75908 | magnesium(2+) | 1(Mg(2+)) | 318, 319, 361 | Keyword & binding & cofactor & go term |

Supplementary Table 2: E. coli annotated metal-binding proteome *(continued)*

| # | Gene Name | UniProt ID | Ligand Name | Ligand Identifier | Ligand Position | Source |
|---|---|---|---|---|---|---|
| 859 | ycdX | P75914 | zinc(2+); zinc(2+); zinc(2+) | 1(Zn(2+)); 2(Zn(2+)); 3(Zn(2+)) | 7, 9, 73, 192; 15, 40, 194; 73, 101, 131 | Keyword & binding & cofactor & go term; Keyword & binding & cofactor & go term; Keyword & binding & cofactor & go term |
| 860 | yceJ | P75925 | ferroheme b(2-); ferroheme b(2-); iron cation | 1(heme b); 2(heme b); - | 18, 159; 52, 145; - | Keyword & binding & cofactor & go term; Keyword & binding & cofactor & go term; Keyword & go term |
| 861 | comR | P75952 | copper cation | - | - | Keyword |
| 862 | nagK | P75959 | zinc(2+) | 1(Zn(2+)) | 157, 177, 179, 184 | Keyword & binding & go term |
| 863 | cobB | P75960 | zinc(2+) | 1(Zn(2+)) | 155, 174 | Keyword & binding & cofactor & go term |
| 864 | ycgM | P76004 | magnesium(2+) | 1(Mg(2+)) | 70, 72, 101 | Keyword & binding & cofactor & go term |
| 865 | cvrA | P76007 | potassium(1+) | - | - | Keyword & catalytic activity & go term |
| 866 | dhaL | P76014 | magnesium(2+) | 1(Mg(2+)) | 30, 35, 37 | Keyword & binding & cofactor & go term |
| 867 | ycjQ | P76043 | zinc(2+) | - | - | Keyword & cofactor & go term |
| 868 | ycjR | P76044 | manganese(2+) | 1(Mn(2+)) | 146, 179, 205, 240 | Keyword & binding & cofactor & go term |
| 869 | ycjX | P76046 | magnesium(2+) | - | - | cofactor |
| 870 | abgB | P76052 | manganese(2+) | - | - | cofactor |
| 871 | ttcA | P76055 | iron cation, magnesium(2+); tetra-mu3-sulfido-tetrairon | -; 1([4Fe-4S] cluster) | -; 122, 125, 213 | Keyword & go term, Keyword & cofactor & go term; Keyword & binding & cofactor & go term |
| 872 | paaA | P76077 | iron cation | - | - | cofactor |
| 873 | paaE | P76081 | di-mu-sulfido-diiron; iron cation | 1([2Fe-2S] cluster); - | 299, 304, 307, 337; - | Keyword & binding & cofactor & go term; Keyword & go term |
| 874 | ydcJ | P76097 | iron(2+) | 1(Fe(2+)) | 68, 224, 290 | Keyword & binding & cofactor & go term |
| 875 | insQ | P76102 | zinc(2+) | 1(Zn(2+)) | 334, 337, 353, 356 | Keyword & binding & go term |
| 876 | pqqU | P76115 | siderophore | - | - | go term |
| 877 | dosP | P76129 | heme; iron cation, magnesium(2+) | 1(heme); - | 69, 87; - | Keyword & binding & cofactor & go term; Keyword & go term, Keyword & cofactor & go term |
| 878 | ydeM | P76134 | iron cation; tetra-mu3-sulfido-tetrairon; tetra-mu3-sulfido-tetrairon; tetra-mu3-sulfido-tetrairon | -; 1([4Fe-4S] cluster); 2([4Fe-4S] cluster); 3([4Fe-4S] cluster) | -; 12, 16, 19; 257, 263, 278, 328; 315, 318, 324, 345 | Keyword & go term; Keyword & binding & cofactor & go term; Keyword & binding & cofactor & go term; Keyword & binding & cofactor & go term |
| 879 | dgcF | P76147 | magnesium(2+) | 1(Mg(2+)) | 181, 182, 224 | Keyword & binding & cofactor & go term |

Supplementary Table 2: E. coli annotated metal-binding proteome *(continued)*

| # | Gene Name | UniProt ID | Ligand Name | Ligand Identifier | Ligand Position | Source |
|---|---|---|---|---|---|---|
| 880 | ydhV | P76192 | iron cation, Mo(VI)O2(OH)-molybdopterin cofactor(4-), bis(molybdopterin)tungsten cofactor; tetra-mu3-sulfido-tetrairon | -; 1([4Fe-4S] cluster) | -; 307, 310, 314, 558 | Keyword & go term, cofactor, cofactor; Keyword & binding & cofactor & go term |
| 881 | astE | P76215 | zinc(2+) | 1(Zn(2+)) | 53, 56, 147 | Keyword & binding & cofactor & go term |
| 882 | cdgI | P76236 | magnesium(2+) | 1(Mg(2+)) | 364, 365, 407 | Keyword & binding & cofactor & go term |
| 883 | dgcJ | P76237 | magnesium(2+) | 1(Mg(2+)) | 382, 425 | Keyword & binding & cofactor & go term |
| 884 | dgcP | P76245 | magnesium(2+) | 1(Mg(2+)) | 212, 255 | Keyword & binding & cofactor & go term |
| 885 | dmlA | P76251 | magnesium(2+); manganese(2+) | -; 1(Mn(2+)) | -; 224, 248, 252 | Keyword & cofactor & go term; Keyword & binding & cofactor & go term |
| 886 | yeaX | P76254 | di-mu-sulfido-diiron; iron cation | 1([2Fe-2S] cluster); - | 270, 275, 278, 308; - | Keyword & binding & cofactor & go term; Keyword & go term |
| 887 | tsaB | P76256 | metal cation | - | - | go term |
| 888 | yoaA | P76257 | iron cation, magnesium(2+); tetra-mu3-sulfido-tetrairon | -; 1([4Fe-4S] cluster) | -; 108, 168, 173, 179 | Keyword & go term, Keyword & cofactor & go term; binding & cofactor & go term |
| 889 | mntP | P76264 | manganese(2+) | - | - | Keyword & go term |
| 890 | yedP | P76329 | magnesium(2+) | 1(Mg(2+)) | 13, 15, 214 | Keyword & binding & cofactor & go term |
| 891 | dgcQ | P76330 | magnesium(2+) | 1(Mg(2+)) | 436, 479 | Keyword & binding & cofactor & go term |
| 892 | msrP | P76342 | Mo(VI)O2(OH)-molybdopterin cofactor(4-); molybdenum cation | 1(Mo-molybdopterin); - | 88, 91, 92, 146, 181, 233, 238, 249, 250, 251; - | binding & cofactor; Keyword & go term |
| 893 | msrQ | P76343 | iron cation, ferroheme b(2-) | - | - | Keyword & go term, Keyword & cofactor & go term |
| 894 | zinT | P76344 | nickel cation, cadmium(2+); zinc(2+); zinc(2+) | -; 1(Zn(2+)); 2(Zn(2+)) | -; 167, 176, 178; 176, 212, 216 | Keyword & go term, Keyword & go term; Keyword & binding & go term; Keyword & binding & go term |
| 895 | yodB | P76345 | ferroheme b(2-); ferroheme b(2-); iron cation | 1(heme b); 2(heme b); - | 12, 152; 46, 138; - | Keyword & binding & cofactor & go term; Keyword & binding & cofactor & go term; Keyword & go term |
| 896 | mtfA | P76346 | zinc(2+) | 1(Zn(2+)) | 111, 148, 152, 211 | Keyword & binding & cofactor & go term |
| 897 | yeeS | P76362 | zinc(2+) | 1(Zn(2+)) | 97, 99, 110 | Keyword & binding & go term |
| 898 | cbeA | P76364 | magnesium(2+) | 1(Mg(2+)) | 86, 88, 102 | Keyword & binding & cofactor & go term |
| 899 | pphC | P76395 | manganese(2+) | - | - | Keyword & cofactor |

Supplementary Table 2: E. coli annotated metal-binding proteome *(continued)*

| # | Gene Name | UniProt ID | Ligand Name | Ligand Identifier | Ligand Position | Source |
|---|---|---|---|---|---|---|
| 900 | yegS | P76407 | calcium(2+); magnesium(2+) | -; 1(Mg(2+)) | -; 215, 218, 220 | Keyword & cofactor & go term; Keyword & binding & cofactor & go term |
| 901 | thiM | P76423 | magnesium(2+) | - | - | Keyword & cofactor & go term |
| 902 | rcnA | P76425 | cobalt cation, nickel(2+) | - | - | Keyword, Keyword & go term |
| 903 | preT | P76440 | iron-sulfur cluster | - | - | go term |
| 904 | rhmA | P76469 | magnesium(2+); nickel(2+) | 1(Mg(2+)); - | 153, 179; - | Keyword & binding & cofactor & go term; Keyword & cofactor & go term |
| 905 | yfbL | P76482 | metal cation | - | - | go term |
| 906 | yfbR | P76491 | cobalt(2+); manganese(2+), copper(2+) | 1(Co(2+)); - | 33, 68, 69, 137; - | Keyword & binding & cofactor & go term; Keyword & cofactor & go term, Keyword & cofactor & go term |
| 907 | yfdR | P76514 | cobalt(2+) | - | - | go term |
| 908 | ypdF | P76524 | cobalt(2+), manganese(2+), nickel(2+) | - | - | cofactor & go term, cofactor & go term, cofactor & go term |
| 909 | yfeX | P76536 | ferroheme b(2-); iron cation | 1(heme); - | 215; - | Keyword & binding & cofactor & go term; Keyword & go term |
| 910 | eutL | P76541 | zinc(2+) | 1(Zn(2+)) | 157 | Keyword & binding & go term |
| 911 | eutG | P76553 | iron cation | 1(Fe cation) | 212, 216, 281, 295 | Keyword & binding & cofactor & go term |
| 912 | maeB | P76558 | magnesium(2+), manganese(2+) | 1(a divalent metal cation) | 136, 137, 162 | Keyword & binding & cofactor & go term, Keyword & binding & cofactor & go term |
| 913 | yfgD | P76569 | arsenate(2-), arsenite(1-) | - | - | go term, go term |
| 914 | patZ | P76594 | metal cation | - | - | go term |
| 915 | glaH | P76621 | iron(2+) | 1(Fe cation) | 160, 162, 292 | Keyword & binding & cofactor & go term |
| 916 | guaD | P76641 | zinc(2+) | 1(Zn(2+)) | 82, 84, 237, 327 | Keyword & binding & cofactor & go term |
| 917 | epmC | P76938 | metal cation | - | - | go term |
| 918 | pdxY | P77150 | magnesium(2+) | - | - | Keyword & cofactor & go term |
| 919 | paoA | P77165 | di-mu-sulfido-diiron; di-mu-sulfido-diiron; iron cation | 1([2Fe-2S] cluster); 2([2Fe-2S] cluster); - | 99, 104, 105, 107, 119; 158, 161, 208, 210; - | Keyword & binding & cofactor & go term; Keyword & binding & cofactor & go term; Keyword & go term |
| 920 | pdeF | P77172 | magnesium(2+), manganese(2+) | - | - | cofactor, cofactor |
| 921 | cusC | P77211 | copper(2+) | - | - | go term |
| 922 | rclA | P77212 | copper(1+), copper(2+) | - | - | go term, go term |
| 923 | cusF | P77214 | copper cation | - | - | Keyword & go term |
| 924 | rhmD | P77215 | magnesium(2+) | 1(Mg(2+)) | 222, 248, 276 | Keyword & binding & cofactor & go term |

Supplementary Table 2: E. coli annotated metal-binding proteome *(continued)*

| # | Gene Name | UniProt ID | Ligand Name | Ligand Identifier | Ligand Position | Source |
|---|---|---|---|---|---|---|
| 925 | rsxB | P77223 | iron cation; tetra-mu3-sulfido-tetrairon; tetra-mu3-sulfido-tetrairon; tetra-mu3-sulfido-tetrairon | -; 1([4Fe-4S] cluster); 2([4Fe-4S] cluster); 3([4Fe-4S] cluster) | -; 49, 52, 57, 74; 117, 120, 123, 157; 127, 147, 150, 153 | Keyword & go term; Keyword & binding & cofactor & go term; Keyword & binding & cofactor & go term; Keyword & binding & cofactor & go term |
| 926 | ydfJ | P77228 | potassium(1+) | - | - | go term |
| 927 | cusB | P77239 | copper(2+) | - | - | Keyword & go term |
| 928 | prpD | P77243 | Fe2S2 iron-sulfur cluster | - | - | go term |
| 929 | hxpB | P77247 | cobalt(2+), magnesium(2+), manganese(2+), zinc(2+) | 1(a divalent metal cation) | 13, 15, 173 | Keyword & binding & cofactor & go term, Keyword & binding & cofactor & go term, Keyword & binding & cofactor & go term, Keyword & binding & cofactor & go term |
| 930 | fetA | P77279 | iron cation | - | - | Keyword |
| 931 | ydjJ | P77280 | zinc(2+); zinc(2+) | 1(Zn(2+)); 2(Zn(2+)) | 39, 65, 152; 95, 98, 101, 109 | Keyword & binding & cofactor & go term; Keyword & binding & cofactor & go term |
| 932 | dgcM | P77302 | magnesium(2+) | 1(Mg(2+)) | 291, 334 | Keyword & binding & cofactor & go term |
| 933 | fetB | P77307 | iron cation | - | - | Keyword |
| 934 | ybdR | P77316 | zinc(2+); zinc(2+) | 1(Zn(2+)); 2(Zn(2+)) | 38, 60; 90, 93, 96, 104 | Keyword & binding & cofactor & go term; Keyword & binding & cofactor & go term |
| 935 | ydeN | P77318 | calcium(2+) | 1(Ca(2+)) | 66, 67, 132, 345, 346 | Keyword & binding & cofactor & go term |
| 936 | paoB | P77324 | iron cation; tetra-mu3-sulfido-tetrairon | -; 1([4Fe-4S] cluster) | -; 119, 129, 138, 157 | Keyword & go term; Keyword & binding & cofactor & go term |
| 937 | hyfG | P77329 | tetra-mu3-sulfido-tetrairon, iron cation, nickel cation | - | - | Keyword & cofactor & go term, Keyword & go term, Keyword & go term |
| 938 | mscK | P77338 | potassium(1+) | - | - | Keyword |
| 939 | abgA | P77357 | manganese(2+) | - | - | cofactor |
| 940 | yphC | P77360 | zinc(2+); zinc(2+) | 1(Zn(2+)); 2(Zn(2+)) | 40, 70, 158; 100, 103, 106, 114 | Keyword & binding & cofactor & go term; Keyword & binding & cofactor & go term |
| 941 | glxK | P77364 | magnesium(2+), cobalt(2+), manganese(2+), iron(2+), calcium(2+) | - | - | cofactor, cofactor, cofactor, cofactor, cofactor |
| 942 | ycjU | P77366 | magnesium(2+) | 1(Mg(2+)) | 9, 11, 172 | Keyword & binding & cofactor & go term |

Supplementary Table 2: E. coli annotated metal-binding proteome *(continued)*

| # | Gene Name | UniProt ID | Ligand Name | Ligand Identifier | Ligand Position | Source |
|---|---|---|---|---|---|---|
| 943 | ynfE | P77374 | Mo(=O)-bis(molybdopterin guanine dinucleotide)(4-); iron cation, molybdenum cation, selenate; tetra-mu3-sulfido-tetrairon | 1(Mo-bis(molybdopterin guanine dinucleotide)); -; 1([4Fe-4S] cluster) | 196; -; 56, 60, 64, 96 | binding & cofactor; Keyword & go term, Keyword & go term, go term; Keyword & binding & cofactor & go term |
| 944 | ydhX | P77375 | iron cation; tetra-mu3-sulfido-tetrairon; tetra-mu3-sulfido-tetrairon; tetra-mu3-sulfido-tetrairon; tetra-mu3-sulfido-tetrairon | -; 1([4Fe-4S] cluster); 2([4Fe-4S] cluster); 3([4Fe-4S] cluster); 4([4Fe-4S] cluster) | -; 46, 49, 52, 171; 56, 151, 154, 167; 92, 95, 100, 134; 104, 124, 127, 130 | Keyword & go term; Keyword & binding & go term; Keyword & binding & go term; Keyword & binding & go term; Keyword & binding & go term |
| 945 | ydhU | P77409 | ferroheme b(2-); ferroheme b(2-); iron cation | 1(heme b); 2(heme b); - | 77, 237; 111, 223; - | binding & cofactor & go term; binding & cofactor & go term; Keyword & go term |
| 946 | hyfH | P77423 | iron cation; tetra-mu3-sulfido-tetrairon; tetra-mu3-sulfido-tetrairon | -; 1([4Fe-4S] cluster); 2([4Fe-4S] cluster) | -; 40, 43, 46, 50; 75, 78, 81, 85 | Keyword & go term; Keyword & binding & go term; Keyword & binding & go term |
| 947 | allC | P77425 | manganese(2+); zinc(2+); zinc(2+) | -; 1(Zn(2+)); 2(Zn(2+)) | -; 81, 92, 190; 92, 127, 382 | Keyword & go term; Keyword & binding & cofactor & go term; Keyword & binding & cofactor & go term |
| 948 | fryA | P77439 | magnesium(2+) | 1(Mg(2+)) | 540, 564 | Keyword & binding & cofactor & go term |
| 949 | yqaB | P77475 | magnesium(2+); manganese(2+), cobalt(2+), zinc(2+) | 1(Mg(2+)); - | 11, 13, 167; - | Keyword & binding & cofactor & go term; Keyword & cofactor & go term, Keyword & cofactor & go term, Keyword & cofactor & go term |
| 950 | cusS | P77485 | copper cation | - | - | Keyword & go term |
| 951 | dxs | P77488 | magnesium(2+) | 1(Mg(2+)) | 152, 181 | Keyword & binding & cofactor & go term |
| 952 | paoC | P77489 | Mo(VI)-molybdopterin cytosine dinucleotide(4-); molybdenum cation, iron cation | 1(Mo-molybdopterin cytosine dinucleotide); - | 241, 242, 468, 469, 470, 511, 512, 615, 616, 617, 618, 619, 620, 621, 625, 688, 689, 690, 691; - | binding & cofactor; Keyword & go term, Keyword & go term |
| 953 | sufB | P77522 | Fe2S2 iron-sulfur cluster, Fe4S4 iron-sulfur cluster | - | - | go term, go term |
| 954 | ykgF | P77536 | iron cation; tetra-mu3-sulfido-tetrairon; tetra-mu3-sulfido-tetrairon | -; 1([4Fe-4S] cluster); 2([4Fe-4S] cluster) | -; 312, 315, 318, 372; 322, 362, 365, 368 | Keyword & go term; Keyword & binding & go term; Keyword & binding & go term |

Supplementary Table 2: E. coli annotated metal-binding proteome *(continued)*

| # | Gene Name | UniProt ID | Ligand Name | Ligand Identifier | Ligand Position | Source |
|---|---|---|---|---|---|---|
| 955 | ydjL | P77539 | zinc(2+); zinc(2+) | 1(Zn(2+)); 2(Zn(2+)) | 39, 61, 157; 92, 95, 98, 106 | Keyword & binding & cofactor & go term; Keyword & binding & cofactor & go term |
| 956 | prpB | P77541 | magnesium(2+) | 1(Mg(2+)) | 85, 87 | Keyword & binding & cofactor & go term |
| 957 | ydeP | P77561 | iron cation, molybdenum cation, Mo(=O)-bis(molybdopterin guanine dinucleotide)(4-); tetra-mu3-sulfido-tetrairon | -; 1([4Fe-4S] cluster) | -; 49, 52 | Keyword & go term, Keyword & go term, cofactor; Keyword & binding & cofactor & go term |
| 958 | ypdE | P77585 | cobalt(2+), nickel(2+), manganese(2+), copper(2+); cobalt(2+), nickel(2+), manganese(2+), copper(2+) | 1(a divalent metal cation); 2(a divalent metal cation) | 62, 166, 221; 166, 199, 308 | Keyword & binding & cofactor & go term, Keyword & binding & cofactor & go term, Keyword & binding & cofactor & go term, Keyword & binding & cofactor & go term; Keyword & binding & cofactor & go term, Keyword & binding & cofactor & go term, Keyword & binding & cofactor & go term, Keyword & binding & cofactor & go term |
| 959 | mhpD | P77608 | manganese(2+) | - | - | cofactor & go term |
| 960 | rsxC | P77611 | iron cation; tetra-mu3-sulfido-tetrairon; tetra-mu3-sulfido-tetrairon | -; 1([4Fe-4S] cluster); 2([4Fe-4S] cluster) | -; 377, 380, 383, 426; 387, 416, 419, 422 | Keyword & go term; Keyword & binding & cofactor & go term; Keyword & binding & cofactor & go term |
| 961 | hxpA | P77625 | cobalt(2+), magnesium(2+), manganese(2+) | 1(a divalent metal cation) | 9, 11, 163 | Keyword & binding & cofactor & go term, Keyword & binding & cofactor & go term, Keyword & binding & cofactor & go term |
| 962 | selO | P77649 | magnesium(2+) | 1(Mg(2+)) | 247, 256 | Keyword & binding & cofactor & go term |
| 963 | hcaD | P77650 | di-mu-sulfido-diiron(2+), di-mu-sulfido-diiron(1+) | - | - | catalytic activity & go term, catalytic activity & go term |
| 964 | sufA | P77667 | di-mu-sulfido-diiron; iron cation; tetra-mu3-sulfido-tetrairon | 1([2Fe-2S] cluster); -; 1([4Fe-4S] cluster) | 50, 114, 116; -; 50, 114, 116 | Keyword & binding & go term; Keyword & go term; Keyword & binding & go term |
| 965 | hyfI | P77668 | iron cation; tetra-mu3-sulfido-tetrairon | -; 1([4Fe-4S] cluster) | -; 41, 47, 111, 141 | Keyword & go term; Keyword & binding & cofactor & go term |

Supplementary Table 2: E. coli annotated metal-binding proteome *(continued)*

| # | Gene Name | UniProt ID | Ligand Name | Ligand Identifier | Ligand Position | Source |
|---|---|---|---|---|---|---|
| 966 | allB | P77671 | nickel(2+), cobalt(2+), manganese(2+); zinc(2+); zinc(2+) | -; 1(Zn(2+)); 2(Zn(2+)) | -; 59, 61, 146, 315; 146, 186, 242 | Keyword & cofactor & go term, Keyword & cofactor & go term, Keyword & cofactor & go term; Keyword & binding & cofactor & go term; Keyword & binding & cofactor & go term |
| 967 | ydjI | P77704 | zinc(2+) | - | - | go term |
| 968 | ydiT | P77714 | iron cation, iron-sulfur cluster | - | - | Keyword & go term, go term |
| 969 | thiI | P77718 | di-mu-sulfido-diiron(2+), di-mu-sulfido-diiron(1+) | - | - | catalytic activity & go term, catalytic activity & go term |
| 970 | allA | P77731 | nickel(2+) | - | - | cofactor |
| 971 | ydiJ | P77748 | iron(3+), iron(2+); tetra-mu3-sulfido-tetrairon | -; 1([4Fe-4S] cluster) | -; 673, 676, 679, 683 | Keyword & go term, Keyword & go term; Keyword & binding & cofactor & go term |
| 972 | queC | P77756 | zinc(2+) | 1(Zn(2+)) | 188, 197, 200, 203 | Keyword & binding & cofactor & go term |
| 973 | rnm | P77766 | manganese(2+); manganese(2+); manganese(2+) | 1(Mn(2+)); 2(Mn(2+)); 3(Mn(2+)) | 13, 15, 72, 255; 20, 45, 257; 72, 83, 198 | Keyword & binding & cofactor & go term; Keyword & binding & cofactor & go term; Keyword & binding & cofactor & go term |
| 974 | rpnB | P77768 | magnesium cation | - | - | Keyword |
| 975 | ynfF | P77783 | Mo(=O)-bis(molybdopterin guanine dinucleotide)(4-); iron cation, molybdenum cation, selenate; tetra-mu3-sulfido-tetrairon | 1(Mo-bis(molybdopterin guanine dinucleotide)); -; 1([4Fe-4S] cluster) | 195; -; 59, 63, 67, 99 | binding & cofactor; Keyword & go term, Keyword & go term, go term; Keyword & binding & cofactor & go term |
| 976 | nudG | P77788 | magnesium(2+), manganese(2+) | - | - | Keyword & cofactor, Keyword & cofactor |
| 977 | ddpX | P77790 | zinc(2+) | 1(Zn(2+)) | 98, 105, 165 | Keyword & binding & cofactor & go term |
| 978 | puuA | P78061 | magnesium(2+), manganese(2+) | - | - | Keyword & cofactor, Keyword & cofactor |
| 979 | ygeR | Q46798 | metal cation | - | - | go term |
| 980 | xdhA | Q46799 | Mo(VI)O2(OH)-molybdopterin cofactor(4-); molybdenum cation, iron cation | 1(Mo-molybdopterin); - | 206, 237, 350, 516; - | binding & cofactor; Keyword & go term, Keyword & go term |
| 981 | xdhC | Q46801 | di-mu-sulfido-diiron; iron cation | 1([2Fe-2S] cluster); - | 44, 49, 52; - | Keyword & binding & cofactor & go term; Keyword & go term |

Supplementary Table 2: E. coli annotated metal-binding proteome *(continued)*

| # | Gene Name | UniProt ID | Ligand Name | Ligand Identifier | Ligand Position | Source |
|---|---|---|---|---|---|---|
| 982 | hyuA | Q46806 | zinc(2+), nickel(2+), cobalt(2+), manganese(2+); zinc(2+), nickel(2+), cobalt(2+), manganese(2+) | 1(a divalent metal cation); 2(a divalent metal cation) | 59, 61, 151, 313; 151, 182, 239 | Keyword & binding & cofactor & go term, Keyword & binding & cofactor & go term, Keyword & binding & cofactor & go term, Keyword & binding & cofactor & go term; Keyword & binding & cofactor & go term, Keyword & binding & cofactor & go term, Keyword & binding & cofactor & go term, Keyword & binding & cofactor & go term |
| 983 | mocA | Q46810 | Mo(VI)O2(OH)-molybdopterin cofactor(4-), Mo(VI)-molybdopterin cytosine dinucleotide(4-), manganese(2+); magnesium(2+) | -; 1(Mg(2+)) | -; 101 | catalytic activity & go term, catalytic activity & go term, Keyword & cofactor & go term; Keyword & binding & cofactor & go term |
| 984 | ygfK | Q46811 | iron cation; tetra-mu3-sulfido-tetrairon | -; 1([4Fe-4S] cluster) | -; 938, 941, 944, 948 | Keyword & go term; Keyword & binding & cofactor & go term |
| 985 | ssnA | Q46812 | zinc(2+) | 1(Zn(2+)) | 62, 64, 227, 312 | Keyword & binding & go term |
| 986 | xdhD | Q46814 | Mo(VI)O2(OH)-molybdopterin cofactor(4-); di-mu-sulfido-diiron, iron cation, molybdenum cation | 1(Mo-molybdopterin); - | 414, 445, 727; - | binding & cofactor; Keyword & cofactor & go term, Keyword & go term, Keyword & go term |
| 987 | ygfS | Q46819 | iron cation; tetra-mu3-sulfido-tetrairon; tetra-mu3-sulfido-tetrairon; tetra-mu3-sulfido-tetrairon; tetra-mu3-sulfido-tetrairon | -; 1([4Fe-4S] cluster); 2([4Fe-4S] cluster); 3([4Fe-4S] cluster); 4([4Fe-4S] cluster) | -; 12, 15, 18, 133; 22, 117, 120, 129; 55, 58, 63, 96; 67, 86, 89, 92 | Keyword & go term; Keyword & binding & go term; Keyword & binding & go term; Keyword & binding & go term; Keyword & binding & go term |
| 988 | uacF | Q46820 | iron cation; tetra-mu3-sulfido-tetrairon; tetra-mu3-sulfido-tetrairon; tetra-mu3-sulfido-tetrairon; tetra-mu3-sulfido-tetrairon; tetra-mu3-sulfido-tetrairon | -; 1([4Fe-4S] cluster); 2([4Fe-4S] cluster); 3([4Fe-4S] cluster); 4([4Fe-4S] cluster); 5([4Fe-4S] cluster) | -; 12, 15, 18, 129; 22, 112, 115, 125; 56, 59, 64, 97; 68, 87, 90, 93; 210, 213, 219, 223 | Keyword & go term; Keyword & binding & cofactor & go term; Keyword & binding & cofactor & go term; Keyword & binding & cofactor & go term; Keyword & binding & cofactor & go term; Keyword & binding & cofactor & go term |

Supplementary Table 2: E. coli annotated metal-binding proteome *(continued)*

| # | Gene Name | UniProt ID | Ligand Name | Ligand Identifier | Ligand Position | Source |
|---|---|---|---|---|---|---|
| 989 | idi | Q46822 | magnesium(2+); manganese(2+); zinc(2+) | 1(Mg(2+)); 1(Mn(2+)); - | 67, 87; 25, 32, 69, 114, 116; - | Keyword & binding & cofactor & go term; Keyword & binding & cofactor & go term; Keyword & go term |
| 990 | yqhD | Q46856 | zinc(2+) | 1(Zn(2+)) | 194, 198, 267, 281 | Keyword & binding & cofactor & go term |
| 991 | ygiQ | Q46861 | iron cation; tetra-mu3-sulfido-tetrairon | -; 1([4Fe-4S] cluster) | -; 386, 390, 393 | Keyword & go term; Keyword & binding & cofactor & go term |
| 992 | mqsA | Q46864 | zinc(2+) | 1(Zn(2+)) | 3, 6, 37, 40 | Keyword & binding & cofactor & go term |
| 993 | yqjH | Q46871 | iron(2+), iron(3+) | - | - | catalytic activity & go term, catalytic activity & go term |
| 994 | norV | Q46877 | iron(2+), iron(3+); iron(2+), iron(3+); iron(2+), iron(3+) | 1(Fe cation); 2(Fe cation); 3(Fe cation) | 79, 81, 147, 166; 83, 166, 227; 428, 431, 461, 464 | Keyword & binding & cofactor & go term, Keyword & binding & cofactor & go term; Keyword & binding & cofactor & go term, Keyword & binding & cofactor & go term; Keyword & binding & cofactor & go term, Keyword & binding & cofactor & go term |
| 995 | otnC | Q46890 | zinc(2+) | 1(Zn(2+)) | 79, 98, 100, 165 | Keyword & binding & cofactor & go term |
| 996 | otnI | Q46891 | magnesium(2+) | 1(Mg(2+)) | 143, 178, 204, 240 | Keyword & binding & go term |
| 997 | ispD | Q46893 | magnesium(2+), manganese(2+), cobalt(2+) | - | - | Keyword & cofactor & go term, Keyword & cofactor, Keyword & cofactor |
| 998 | ygbT | Q46896 | magnesium(2+) | 1(Mg(2+)) | 141, 208, 221 | Keyword & binding & cofactor & go term |
| 999 | casA | Q46901 | zinc(2+) | - | - | go term |
| 1000 | ygcO | Q46905 | iron cation, iron-sulfur cluster | - | - | Keyword & go term, go term |
| 1001 | ygcU | Q46911 | iron(3+), iron(2+) | - | - | go term, go term |
| 1002 | gudX | Q46915 | magnesium(2+) | 1(Mg(2+)) | 234, 265, 288 | Keyword & binding & cofactor & go term |
| 1003 | tcdA | Q46927 | potassium(1+), sodium(1+) | - | - | go term, go term |
| 1004 | kduI | Q46938 | zinc(2+) | 1(Zn(2+)) | 196, 198, 203, 245 | Keyword & binding & cofactor & go term |
| 1005 | rayT | Q47152 | manganese cation | - | - | Keyword |
| 1006 | dinB | Q47155 | magnesium(2+) | 1(Mg(2+)) | 8, 103 | Keyword & binding & cofactor & go term |
| 1007 | tapT | Q47319 | zinc(2+) | 1(Zn(2+)) | 31, 34, 41, 43 | Keyword & binding & go term |
| 1008 | ykfG | Q47685 | zinc(2+) | 1(Zn(2+)) | 107, 109, 120 | Keyword & binding & go term |
| 1009 | ykfC | Q47688 | magnesium(2+) | 1(Mg(2+)) | 166, 284, 285 | Keyword & binding & go term |
| 1010 | mmuM | Q47690 | zinc(2+) | 1(Zn(2+)) | 229, 295, 296 | Keyword & binding & cofactor & go term |

Supplementary Table 2: E. coli annotated metal-binding proteome *(continued)*

| # | Gene Name | UniProt ID | Ligand Name | Ligand Identifier | Ligand Position | Source |
|---|---|---|---|---|---|---|
| 1011 | copA | Q59385 | copper(1+);<br>copper(1+);<br>copper(2+);<br>magnesium(2+) | 1(Cu(+));<br>2(Cu(+));<br>-;<br>1(Mg(2+)) | 14, 17;<br>110, 113;<br>-;<br>720, 724 | Keyword & binding & catalytic activity & go term;<br>Keyword & binding & catalytic activity & go term;<br>Keyword & go term;<br>Keyword & binding & go term |
| 1012 | dgoD | Q6BF17 | magnesium(2+) | 1(Mg(2+)) | 183, 209, 235 | Keyword & binding & cofactor & go term |
| 1013 | nudF | Q93K97 | magnesium(2+);<br>magnesium(2+);<br>magnesium(2+);<br>manganese cation | 1(Mg(2+));<br>2(Mg(2+));<br>3(Mg(2+));<br>- | 96, 116;<br>112;<br>112, 116, 164;<br>- | Keyword & binding & cofactor & go term;<br>Keyword & binding & cofactor & go term;<br>Keyword & binding & cofactor & go term;<br>Keyword & go term |

Supplementary Table 3: E. coli predicted metal-binding proteome

| # | Gene Name | UniProt ID | MoM Prediction | M3D Prediction | GASS Prediction |
|---|---|---|---|---|---|
| 1 | gloB | P0AC84 | TRUE | TRUE | TRUE |
| 2 | basR | P30843 | FALSE | FALSE | TRUE |
| 3 | yliI | P75804 | FALSE | FALSE | FALSE |
| 4 | mhpF | P77580 | FALSE | FALSE | FALSE |
| 5 | lpxM | P24205 | FALSE | FALSE | FALSE |
| 6 | ygeG | Q46787 | FALSE | FALSE | FALSE |
| 7 | gadB | P69910 | FALSE | FALSE | FALSE |
| 8 | fabI | P0AEK4 | FALSE | FALSE | FALSE |
| 9 | rplM | P0AA10 | FALSE | MISSING | FALSE |
| 10 | malP | P00490 | FALSE | FALSE | FALSE |
| 11 | casC | Q46899 | FALSE | FALSE | FALSE |
| 12 | pepB | P37095 | TRUE | TRUE | FALSE |
| 13 | alpA | P33997 | FALSE | FALSE | FALSE |
| 14 | macA | P75830 | FALSE | FALSE | FALSE |
| 15 | waaJ | P27129 | FALSE | TRUE | FALSE |
| 16 | purB | P0AB89 | FALSE | FALSE | FALSE |
| 17 | hemF | P36553 | FALSE | FALSE | TRUE |
| 18 | cynS | P00816 | FALSE | FALSE | FALSE |
| 19 | map | P0AE18 | FALSE | TRUE | FALSE |
| 20 | pyrI | P0A7F3 | TRUE | TRUE | FALSE |
| 21 | murG | P17443 | FALSE | FALSE | FALSE |
| 22 | dinB | Q47155 | FALSE | FALSE | FALSE |
| 23 | prmB | P39199 | FALSE | FALSE | FALSE |
| 24 | folB | P0AC16 | FALSE | FALSE | FALSE |
| 25 | deoR | P0ACK5 | FALSE | FALSE | FALSE |
| 26 | cbl | Q47083 | FALSE | FALSE | FALSE |
| 27 | recR | P0A7H6 | TRUE | TRUE | TRUE |
| 28 | lpdA | P0A9P0 | FALSE | FALSE | FALSE |
| 29 | prpD | P77243 | TRUE | TRUE | FALSE |
| 30 | ykfH | Q9XB42 | FALSE | FALSE | FALSE |
| 31 | nfo | P0A6C1 | TRUE | TRUE | FALSE |
| 32 | cysD | P21156 | FALSE | FALSE | FALSE |
| 33 | edd | P0ADF6 | FALSE | FALSE | TRUE |
| 34 | gutQ | P17115 | FALSE | FALSE | FALSE |
| 35 | cysS | P21888 | TRUE | TRUE | FALSE |
| 36 | narI | P11350 | FALSE | TRUE | FALSE |
| 37 | hisB | P06987 | TRUE | TRUE | FALSE |
| 38 | nagZ | P75949 | FALSE | FALSE | FALSE |
| 39 | efeB | P31545 | FALSE | FALSE | FALSE |
| 40 | dcd | P28248 | FALSE | TRUE | FALSE |
| 41 | yqeA | Q46807 | FALSE | FALSE | FALSE |
| 42 | tatD | P27859 | TRUE | TRUE | FALSE |
| 43 | lipB | P60720 | FALSE | FALSE | FALSE |
| 44 | eutB | P0AEJ6 | TRUE | FALSE | FALSE |
| 45 | suhB | P0ADG4 | FALSE | FALSE | FALSE |
| 46 | ulaG | P39300 | TRUE | TRUE | FALSE |
| 47 | pyrH | P0A7E9 | FALSE | FALSE | TRUE |
| 48 | ilvI | P00893 | FALSE | FALSE | FALSE |
| 49 | stfP | P45581 | FALSE | FALSE | FALSE |
| 50 | yobA | P0AA57 | FALSE | FALSE | TRUE |
| 51 | bamA | P0A940 | TRUE | FALSE | FALSE |
| 52 | glgC | P0A6V1 | FALSE | FALSE | FALSE |
| 53 | pflC | P32675 | FALSE | TRUE | FALSE |
| 54 | yijF | P32668 | FALSE | FALSE | TRUE |
| 55 | betA | P17444 | FALSE | FALSE | TRUE |
| 56 | fis | P0A6R3 | FALSE | MISSING | FALSE |

Supplementary Table 3: E. coli predicted metal-binding proteome *(continued)*

| # | Gene Name | UniProt ID | MoM Prediction | M3D Prediction | GASS Prediction |
|---|---|---|---|---|---|
| 57 | ydeJ | P31131 | FALSE | FALSE | FALSE |
| 58 | lolD | P75957 | FALSE | FALSE | FALSE |
| 59 | glf | P37747 | FALSE | FALSE | FALSE |
| 60 | rpsL | P0A7S3 | FALSE | FALSE | FALSE |
| 61 | uvrA | P0A698 | TRUE | TRUE | TRUE |
| 62 | ispF | P62617 | TRUE | TRUE | FALSE |
| 63 | bcsQ | P37655 | FALSE | FALSE | FALSE |
| 64 | araD | P08203 | TRUE | TRUE | TRUE |
| 65 | nrdI | P0A772 | FALSE | FALSE | TRUE |
| 66 | pldA | P0A921 | FALSE | FALSE | FALSE |
| 67 | mlaC | P0ADV7 | FALSE | MISSING | FALSE |
| 68 | fixA | P60566 | FALSE | MISSING | FALSE |
| 69 | yjhQ | P39368 | FALSE | FALSE | TRUE |
| 70 | yciB | P0A710 | FALSE | FALSE | FALSE |
| 71 | yphC | P77360 | TRUE | TRUE | TRUE |
| 72 | fryB | P69808 | FALSE | FALSE | FALSE |
| 73 | yjdF | P39270 | FALSE | FALSE | FALSE |
| 74 | mnmC | P77182 | FALSE | FALSE | FALSE |
| 75 | queE | P64554 | FALSE | FALSE | TRUE |
| 76 | yfcQ | P76500 | FALSE | FALSE | FALSE |
| 77 | nuoC | P33599 | FALSE | FALSE | FALSE |
| 78 | yqiA | P0A8Z7 | FALSE | FALSE | FALSE |
| 79 | abgR | P77744 | FALSE | FALSE | FALSE |
| 80 | pflB | P09373 | FALSE | FALSE | TRUE |
| 81 | chbF | P17411 | FALSE | FALSE | FALSE |
| 82 | intR | P76056 | FALSE | FALSE | FALSE |
| 83 | thiH | P30140 | TRUE | FALSE | FALSE |
| 84 | yjcO | P0AF56 | FALSE | FALSE | FALSE |
| 85 | fruB | P69811 | FALSE | FALSE | FALSE |
| 86 | atpF | P0ABA0 | FALSE | FALSE | FALSE |
| 87 | leuS | P07813 | TRUE | TRUE | FALSE |
| 88 | gfcE | P0A932 | FALSE | TRUE | FALSE |
| 89 | uspB | P0A8S5 | FALSE | FALSE | FALSE |
| 90 | sapB | P0AGH3 | FALSE | FALSE | FALSE |
| 91 | aspA | P0AC38 | FALSE | FALSE | TRUE |
| 92 | ptrA | P05458 | TRUE | TRUE | TRUE |
| 93 | yqeB | Q46808 | FALSE | FALSE | FALSE |
| 94 | yhhX | P46853 | FALSE | FALSE | FALSE |
| 95 | ynaI | P0AEB5 | FALSE | FALSE | FALSE |
| 96 | ylbH | P77759 | TRUE | TRUE | TRUE |
| 97 | ydeP | P77561 | TRUE | FALSE | TRUE |
| 98 | ygiL | P39834 | FALSE | FALSE | FALSE |
| 99 | citG | P77231 | FALSE | FALSE | FALSE |
| 100 | lplA | P32099 | FALSE | FALSE | FALSE |
| 101 | sad | P76149 | FALSE | FALSE | FALSE |
| 102 | alsE | P32719 | TRUE | TRUE | TRUE |
| 103 | ybiV | P75792 | FALSE | FALSE | FALSE |
| 104 | ydhV | P76192 | TRUE | FALSE | FALSE |
| 105 | yeaX | P76254 | TRUE | TRUE | FALSE |
| 106 | ycfP | P0A8E1 | FALSE | FALSE | FALSE |
| 107 | frwD | P32676 | FALSE | FALSE | FALSE |
| 108 | mutM | P05523 | TRUE | TRUE | TRUE |
| 109 | yjgH | P39332 | FALSE | FALSE | FALSE |
| 110 | sgcA | P39363 | FALSE | FALSE | FALSE |
| 111 | intZ | P76542 | FALSE | FALSE | FALSE |
| 112 | fabH | P0A6R0 | FALSE | FALSE | FALSE |

Supplementary Table 3: E. coli predicted metal-binding proteome *(continued)*

| # | Gene Name | UniProt ID | MoM Prediction | M3D Prediction | GASS Prediction |
|---|---|---|---|---|---|
| 113 | yqiH | P77616 | FALSE | MISSING | FALSE |
| 114 | proA | P07004 | FALSE | FALSE | FALSE |
| 115 | atoA | P76459 | FALSE | FALSE | FALSE |
| 116 | lpxT | P76445 | FALSE | FALSE | TRUE |
| 117 | dnaK | P0A6Y8 | FALSE | FALSE | FALSE |
| 118 | hemW | P52062 | FALSE | FALSE | FALSE |
| 119 | ccmB | P0ABL8 | FALSE | FALSE | FALSE |
| 120 | clpP | P0A6G7 | FALSE | FALSE | FALSE |
| 121 | yigF | P27842 | FALSE | MISSING | FALSE |
| 122 | rutR | P0ACU2 | FALSE | FALSE | FALSE |
| 123 | napA | P33937 | TRUE | FALSE | FALSE |
| 124 | yaeI | P37049 | TRUE | TRUE | FALSE |
| 125 | metE | P25665 | TRUE | TRUE | TRUE |
| 126 | gpsA | P0A6S7 | FALSE | FALSE | FALSE |
| 127 | ldcA | P76008 | FALSE | FALSE | FALSE |
| 128 | thiC | P30136 | FALSE | TRUE | TRUE |
| 129 | yceO | P64442 | FALSE | MISSING | FALSE |
| 130 | frdA | P00363 | FALSE | FALSE | TRUE |
| 131 | rhaA | P32170 | FALSE | TRUE | FALSE |
| 132 | gltK | P0AER5 | FALSE | FALSE | FALSE |
| 133 | yadG | P36879 | FALSE | FALSE | FALSE |
| 134 | alaC | P77434 | FALSE | FALSE | FALSE |
| 135 | btuF | P37028 | FALSE | FALSE | FALSE |
| 136 | zur | P0AC51 | TRUE | TRUE | FALSE |
| 137 | ycjO | P0AFR7 | FALSE | FALSE | FALSE |
| 138 | oppA | P23843 | FALSE | TRUE | FALSE |
| 139 | bisC | P20099 | FALSE | TRUE | FALSE |
| 140 | tpiA | P0A858 | FALSE | FALSE | FALSE |
| 141 | proS | P16659 | FALSE | FALSE | FALSE |
| 142 | mug | P0A9H1 | FALSE | FALSE | FALSE |
| 143 | phoA | P00634 | TRUE | TRUE | FALSE |
| 144 | yggS | P67080 | FALSE | FALSE | FALSE |
| 145 | metG | P00959 | TRUE | TRUE | TRUE |
| 146 | alkB | P05050 | TRUE | TRUE | TRUE |
| 147 | fixB | P31574 | TRUE | TRUE | FALSE |
| 148 | patD | P77674 | FALSE | FALSE | FALSE |
| 149 | pstS | P0AG82 | FALSE | FALSE | FALSE |
| 150 | pabC | P28305 | FALSE | FALSE | FALSE |
| 151 | digH | P64426 | FALSE | FALSE | FALSE |
| 152 | aas | P31119 | FALSE | FALSE | FALSE |
| 153 | ydfG | P39831 | FALSE | FALSE | FALSE |
| 154 | rrrQ | P76159 | FALSE | FALSE | FALSE |
| 155 | cpxR | P0AE88 | FALSE | FALSE | FALSE |
| 156 | fre | P0AEN1 | FALSE | FALSE | FALSE |
| 157 | murC | P17952 | FALSE | FALSE | FALSE |
| 158 | murA | P0A749 | FALSE | FALSE | FALSE |
| 159 | ydhT | P77147 | FALSE | FALSE | FALSE |
| 160 | nuoM | P0AFE8 | FALSE | FALSE | FALSE |
| 161 | ydcZ | P76111 | FALSE | MISSING | FALSE |
| 162 | rsmG | P0A6U5 | FALSE | FALSE | FALSE |
| 163 | ygfA | P0AC28 | FALSE | FALSE | FALSE |
| 164 | ascG | P24242 | FALSE | TRUE | FALSE |
| 165 | ascB | P24240 | FALSE | FALSE | FALSE |
| 166 | yaiO | Q47534 | FALSE | FALSE | FALSE |
| 167 | menF | P38051 | FALSE | FALSE | FALSE |
| 168 | rsmD | P0ADX9 | FALSE | FALSE | FALSE |

Supplementary Table 3: E. coli predicted metal-binding proteome *(continued)*

| # | Gene Name | UniProt ID | MoM Prediction | M3D Prediction | GASS Prediction |
|---|---|---|---|---|---|
| 169 | wecF | P56258 | FALSE | TRUE | TRUE |
| 170 | cysK | P0ABK5 | FALSE | FALSE | FALSE |
| 171 | ypfG | P76559 | FALSE | FALSE | FALSE |
| 172 | ynaK | P76068 | FALSE | FALSE | FALSE |
| 173 | cnoX | P77395 | FALSE | FALSE | FALSE |
| 174 | dppF | P37313 | FALSE | FALSE | FALSE |
| 175 | yhfW | P45549 | TRUE | TRUE | TRUE |
| 176 | acnA | P25516 | FALSE | FALSE | TRUE |
| 177 | carB | P00968 | FALSE | TRUE | FALSE |
| 178 | lpxP | P0ACV2 | FALSE | FALSE | FALSE |
| 179 | hpt | P0A9M2 | FALSE | FALSE | FALSE |
| 180 | rhaD | P32169 | TRUE | TRUE | TRUE |
| 181 | waaY | P27240 | FALSE | FALSE | FALSE |
| 182 | ytfQ | P39325 | FALSE | FALSE | FALSE |
| 183 | fliJ | P52613 | FALSE | FALSE | FALSE |
| 184 | ompL | P76773 | FALSE | FALSE | FALSE |
| 185 | grxC | P0AC62 | FALSE | FALSE | FALSE |
| 186 | ykgJ | P0AAL9 | TRUE | TRUE | FALSE |
| 187 | glvG | P31450 | FALSE | TRUE | FALSE |
| 188 | yceH | P29217 | FALSE | FALSE | FALSE |
| 189 | frlR | P45544 | FALSE | FALSE | FALSE |
| 190 | nanS | P39370 | FALSE | FALSE | FALSE |
| 191 | mppA | P77348 | FALSE | FALSE | FALSE |
| 192 | eutS | P63746 | FALSE | FALSE | TRUE |
| 193 | pphC | P76395 | FALSE | FALSE | FALSE |
| 194 | aer | P50466 | FALSE | FALSE | FALSE |
| 195 | entE | P10378 | FALSE | FALSE | FALSE |
| 196 | yneG | P76148 | FALSE | TRUE | FALSE |
| 197 | mdlA | P77265 | FALSE | TRUE | FALSE |
| 198 | idnK | P39208 | FALSE | FALSE | FALSE |
| 199 | pepP | P15034 | FALSE | TRUE | FALSE |
| 200 | ybhP | P0AAW1 | TRUE | FALSE | FALSE |
| 201 | ybbA | P0A9T8 | FALSE | FALSE | FALSE |
| 202 | yegP | P76402 | FALSE | MISSING | FALSE |
| 203 | gltX | P04805 | TRUE | TRUE | FALSE |
| 204 | malQ | P15977 | TRUE | TRUE | TRUE |
| 205 | ompN | P77747 | FALSE | FALSE | FALSE |
| 206 | lsrF | P76143 | FALSE | FALSE | FALSE |
| 207 | surE | P0A840 | FALSE | FALSE | FALSE |
| 208 | argF | P06960 | FALSE | FALSE | TRUE |
| 209 | fepC | P23878 | FALSE | FALSE | FALSE |
| 210 | dadA | P0A6J5 | FALSE | TRUE | FALSE |
| 211 | ilvG | P0DP89 | FALSE | FALSE | FALSE |
| 212 | nanA | P0A6L4 | FALSE | FALSE | FALSE |
| 213 | ppx | P0AFL6 | TRUE | FALSE | FALSE |
| 214 | aslA | P25549 | FALSE | FALSE | TRUE |
| 215 | focA | P0AC23 | FALSE | FALSE | FALSE |
| 216 | php | P45548 | TRUE | TRUE | TRUE |
| 217 | ydhQ | P77552 | FALSE | FALSE | FALSE |
| 218 | yjiJ | P39381 | FALSE | FALSE | FALSE |
| 219 | ybjI | P75809 | FALSE | FALSE | FALSE |
| 220 | hmp | P24232 | FALSE | FALSE | TRUE |
| 221 | livF | P22731 | FALSE | FALSE | FALSE |
| 222 | ynjE | P78067 | FALSE | FALSE | FALSE |
| 223 | lpxD | P21645 | FALSE | FALSE | FALSE |
| 224 | nhaB | P0AFA7 | TRUE | FALSE | FALSE |

Supplementary Table 3: E. coli predicted metal-binding proteome *(continued)*

| # | Gene Name | UniProt ID | MoM Prediction | M3D Prediction | GASS Prediction |
|---|---|---|---|---|---|
| 225 | cusA | P38054 | FALSE | FALSE | FALSE |
| 226 | yggC | P11664 | FALSE | FALSE | FALSE |
| 227 | kbaZ | P0C8K0 | TRUE | TRUE | FALSE |
| 228 | murQ | P76535 | FALSE | FALSE | FALSE |
| 229 | ynbC | P76092 | TRUE | FALSE | TRUE |
| 230 | ynbD | P76093 | FALSE | TRUE | TRUE |
| 231 | rimO | P0AEI4 | FALSE | FALSE | FALSE |
| 232 | purK | P09029 | FALSE | TRUE | FALSE |
| 233 | yaaA | P0A8I3 | FALSE | FALSE | FALSE |
| 234 | rppH | P0A776 | FALSE | FALSE | FALSE |
| 235 | agaB | P42909 | FALSE | FALSE | FALSE |
| 236 | glvC | P31452 | FALSE | TRUE | FALSE |
| 237 | ybdF | P0AAT2 | FALSE | FALSE | FALSE |
| 238 | nei | P50465 | TRUE | TRUE | FALSE |
| 239 | menH | P37355 | FALSE | FALSE | FALSE |
| 240 | ubiE | P0A887 | FALSE | FALSE | FALSE |
| 241 | mpl | P37773 | FALSE | TRUE | FALSE |
| 242 | thrC | P00934 | FALSE | FALSE | FALSE |
| 243 | lysC | P08660 | FALSE | FALSE | FALSE |
| 244 | yheT | P45524 | FALSE | FALSE | FALSE |
| 245 | dcm | P0AED9 | TRUE | FALSE | TRUE |
| 246 | dsdA | P00926 | FALSE | FALSE | FALSE |
| 247 | yoaB | P0AEB7 | FALSE | FALSE | FALSE |
| 248 | phoU | P0A9K7 | FALSE | FALSE | FALSE |
| 249 | caiT | P31553 | FALSE | TRUE | FALSE |
| 250 | thiF | P30138 | TRUE | TRUE | FALSE |
| 251 | metAS | P07623 | FALSE | FALSE | FALSE |
| 252 | hexR | P46118 | FALSE | FALSE | FALSE |
| 253 | ispB | P0AD57 | TRUE | FALSE | FALSE |
| 254 | gabD | P25526 | FALSE | FALSE | FALSE |
| 255 | galM | P0A9C3 | FALSE | TRUE | FALSE |
| 256 | pnp | P05055 | FALSE | FALSE | FALSE |
| 257 | yfiB | P07021 | FALSE | MISSING | FALSE |
| 258 | hisQ | P52094 | FALSE | MISSING | FALSE |
| 259 | ycaK | P43340 | FALSE | FALSE | FALSE |
| 260 | gsiC | P75798 | FALSE | FALSE | FALSE |
| 261 | guaD | P76641 | TRUE | TRUE | FALSE |
| 262 | thiQ | P31548 | FALSE | FALSE | FALSE |
| 263 | pgm | P36938 | FALSE | TRUE | TRUE |
| 264 | qorA | P28304 | FALSE | FALSE | FALSE |
| 265 | deoC | P0A6L0 | FALSE | FALSE | FALSE |
| 266 | artI | P30859 | FALSE | TRUE | FALSE |
| 267 | yiaL | P37673 | TRUE | TRUE | FALSE |
| 268 | rimP | P0A8A8 | FALSE | FALSE | FALSE |
| 269 | bsmA | P39297 | FALSE | FALSE | FALSE |
| 270 | mpaA | P0ACV6 | TRUE | FALSE | FALSE |
| 271 | gmhB | P63228 | TRUE | TRUE | TRUE |
| 272 | tolQ | P0ABU9 | FALSE | FALSE | FALSE |
| 273 | yccF | P0AB12 | FALSE | FALSE | FALSE |
| 274 | pdeG | P75995 | FALSE | FALSE | FALSE |
| 275 | cysC | P0A6J1 | FALSE | FALSE | FALSE |
| 276 | ddpF | P77622 | FALSE | FALSE | FALSE |
| 277 | mngB | P54746 | FALSE | TRUE | FALSE |
| 278 | lnt | P23930 | FALSE | FALSE | FALSE |
| 279 | fixX | P68646 | FALSE | FALSE | FALSE |
| 280 | ydbL | P76076 | FALSE | MISSING | FALSE |

Supplementary Table 3: E. coli predicted metal-binding proteome *(continued)*

| # | Gene Name | UniProt ID | MoM Prediction | M3D Prediction | GASS Prediction |
|---|---|---|---|---|---|
| 281 | rhaB | P32171 | FALSE | FALSE | FALSE |
| 282 | solA | P40874 | FALSE | TRUE | FALSE |
| 283 | yjdJ | P39274 | FALSE | FALSE | FALSE |
| 284 | yceD | P0AB28 | TRUE | TRUE | FALSE |
| 285 | narU | P37758 | FALSE | FALSE | FALSE |
| 286 | serA | P0A9T0 | FALSE | FALSE | FALSE |
| 287 | flgE | P75937 | FALSE | FALSE | TRUE |
| 288 | mazG | P0AEY3 | FALSE | FALSE | FALSE |
| 289 | tldD | P0AGG8 | TRUE | TRUE | FALSE |
| 290 | satP | P0AC98 | FALSE | FALSE | FALSE |
| 291 | yaiW | P77562 | FALSE | FALSE | FALSE |
| 292 | mak | P23917 | TRUE | TRUE | FALSE |
| 293 | tcdA | Q46927 | FALSE | FALSE | FALSE |
| 294 | phrB | P00914 | FALSE | FALSE | FALSE |
| 295 | waaL | P27243 | FALSE | FALSE | FALSE |
| 296 | thiM | P76423 | FALSE | FALSE | TRUE |
| 297 | gmd | P0AC88 | TRUE | FALSE | FALSE |
| 298 | entA | P15047 | FALSE | FALSE | FALSE |
| 299 | ispH | P62623 | FALSE | FALSE | FALSE |
| 300 | glgX | P15067 | FALSE | FALSE | TRUE |
| 301 | rpsP | P0A7T3 | FALSE | FALSE | FALSE |
| 302 | ytfA | P39309 | FALSE | FALSE | FALSE |
| 303 | aroC | P12008 | FALSE | FALSE | FALSE |
| 304 | bioC | P12999 | FALSE | FALSE | TRUE |
| 305 | exoD | P75717 | FALSE | FALSE | FALSE |
| 306 | hldE | P76658 | FALSE | FALSE | FALSE |
| 307 | yjiE | P39376 | FALSE | FALSE | FALSE |
| 308 | malF | P02916 | FALSE | FALSE | FALSE |
| 309 | rutB | P75897 | FALSE | FALSE | FALSE |
| 310 | appC | P26459 | FALSE | FALSE | FALSE |
| 311 | lpxK | P27300 | FALSE | FALSE | FALSE |
| 312 | sfmC | P77249 | FALSE | FALSE | FALSE |
| 313 | hxpB | P77247 | FALSE | FALSE | FALSE |
| 314 | trpE | P00895 | FALSE | FALSE | FALSE |
| 315 | ykgE | P77252 | TRUE | TRUE | FALSE |
| 316 | dsbE | P0AA86 | FALSE | FALSE | FALSE |
| 317 | yldA | P0DPN4 | FALSE | MISSING | FALSE |
| 318 | gcvP | P33195 | FALSE | FALSE | FALSE |
| 319 | sapF | P0AAH8 | FALSE | FALSE | TRUE |
| 320 | puuC | P23883 | FALSE | FALSE | FALSE |
| 321 | pabB | P05041 | FALSE | FALSE | TRUE |
| 322 | yeiL | P0A9E9 | FALSE | FALSE | FALSE |
| 323 | bioD2 | P0A6E9 | FALSE | FALSE | FALSE |
| 324 | yjgM | P39337 | FALSE | FALSE | FALSE |
| 325 | yhjE | P37643 | FALSE | FALSE | FALSE |
| 326 | potE | P0AAF1 | FALSE | FALSE | FALSE |
| 327 | flgL | P29744 | FALSE | TRUE | FALSE |
| 328 | tnaA | P0A853 | FALSE | FALSE | FALSE |
| 329 | paaA | P76077 | FALSE | TRUE | FALSE |
| 330 | artP | P0AAF6 | FALSE | FALSE | FALSE |
| 331 | ybdN | P77216 | FALSE | FALSE | FALSE |
| 332 | nanY | P39353 | FALSE | TRUE | TRUE |
| 333 | gdx | P69937 | FALSE | FALSE | FALSE |
| 334 | yihX | P0A8Y3 | FALSE | FALSE | FALSE |
| 335 | ydjA | P0ACY1 | FALSE | FALSE | FALSE |
| 336 | shoB | C1P611 | FALSE | MISSING | FALSE |

Supplementary Table 3: E. coli predicted metal-binding proteome *(continued)*

| # | Gene Name | UniProt ID | MoM Prediction | M3D Prediction | GASS Prediction |
|---|---|---|---|---|---|
| 337 | yjjJ | P39410 | FALSE | FALSE | FALSE |
| 338 | ftnB | P0A9A2 | FALSE | FALSE | FALSE |
| 339 | nadD | P0A752 | FALSE | FALSE | FALSE |
| 340 | qseB | P52076 | FALSE | TRUE | FALSE |
| 341 | tktA | P27302 | FALSE | TRUE | FALSE |
| 342 | agaR | P0ACK2 | FALSE | FALSE | FALSE |
| 343 | cysP | P16700 | FALSE | TRUE | FALSE |
| 344 | uxuB | P39160 | TRUE | FALSE | FALSE |
| 345 | wcaA | P77414 | FALSE | FALSE | TRUE |
| 346 | yfhL | P52102 | FALSE | FALSE | FALSE |
| 347 | rlmI | P75876 | FALSE | TRUE | FALSE |
| 348 | yphA | P0AD47 | FALSE | FALSE | FALSE |
| 349 | ytfH | P0ACN2 | FALSE | FALSE | FALSE |
| 350 | fadI | P76503 | FALSE | TRUE | TRUE |
| 351 | miaB | P0AEI1 | FALSE | FALSE | FALSE |
| 352 | gpr | Q46851 | FALSE | TRUE | FALSE |
| 353 | cmoB | P76291 | FALSE | FALSE | FALSE |
| 354 | quuQ | P76161 | TRUE | TRUE | TRUE |
| 355 | mrdA | P0AD65 | FALSE | FALSE | FALSE |
| 356 | ddpX | P77790 | TRUE | TRUE | TRUE |
| 357 | yihQ | P32138 | FALSE | FALSE | FALSE |
| 358 | lptD | P31554 | FALSE | FALSE | FALSE |
| 359 | nupG | P0AFF4 | FALSE | FALSE | FALSE |
| 360 | metI | P31547 | FALSE | MISSING | FALSE |
| 361 | fecB | P15028 | FALSE | FALSE | FALSE |
| 362 | psuK | P30235 | FALSE | FALSE | TRUE |
| 363 | fes | P13039 | TRUE | FALSE | TRUE |
| 364 | yihG | P32129 | FALSE | FALSE | FALSE |
| 365 | aroF | P00888 | FALSE | FALSE | FALSE |
| 366 | insH5 | P76071 | FALSE | FALSE | FALSE |
| 367 | epmA | P0A8N7 | FALSE | TRUE | FALSE |
| 368 | panB | P31057 | FALSE | FALSE | FALSE |
| 369 | yqjF | P42619 | FALSE | FALSE | FALSE |
| 370 | tisB | A5A627 | FALSE | MISSING | FALSE |
| 371 | leuC | P0A6A6 | TRUE | FALSE | TRUE |
| 372 | idnO | P0A9P9 | FALSE | FALSE | FALSE |
| 373 | nudI | P52006 | FALSE | FALSE | FALSE |
| 374 | rbn | P0A8V0 | TRUE | TRUE | FALSE |
| 375 | ydhR | P0ACX3 | FALSE | FALSE | TRUE |
| 376 | eutJ | P77277 | FALSE | TRUE | FALSE |
| 377 | tsaD | P05852 | TRUE | TRUE | TRUE |
| 378 | metB | P00935 | FALSE | FALSE | FALSE |
| 379 | aaaT | P46854 | FALSE | TRUE | FALSE |
| 380 | sgrR | P33595 | TRUE | FALSE | TRUE |
| 381 | fsaB | P32669 | FALSE | FALSE | FALSE |
| 382 | ycjN | P76042 | FALSE | FALSE | FALSE |
| 383 | polB | P21189 | FALSE | FALSE | FALSE |
| 384 | fucR | P0ACK8 | FALSE | FALSE | FALSE |
| 385 | yaiI | P0A8D3 | FALSE | FALSE | FALSE |
| 386 | tdcF | P0AGL2 | FALSE | MISSING | FALSE |
| 387 | yedK | P76318 | FALSE | FALSE | FALSE |
| 388 | kdgT | P0A712 | FALSE | FALSE | FALSE |
| 389 | hcaC | P0ABW0 | TRUE | FALSE | FALSE |
| 390 | rsmB | P36929 | FALSE | TRUE | TRUE |
| 391 | murR | P77245 | FALSE | FALSE | FALSE |
| 392 | rplW | P0ADZ0 | FALSE | FALSE | FALSE |

Supplementary Table 3: E. coli predicted metal-binding proteome *(continued)*

| # | Gene Name | UniProt ID | MoM Prediction | M3D Prediction | GASS Prediction |
|---|---|---|---|---|---|
| 393 | fpr | P28861 | FALSE | FALSE | FALSE |
| 394 | rffG | P27830 | FALSE | FALSE | FALSE |
| 395 | pdxI | P25906 | FALSE | FALSE | FALSE |
| 396 | mqsA | Q46864 | TRUE | TRUE | FALSE |
| 397 | modB | P0AF01 | FALSE | MISSING | FALSE |
| 398 | tcyJ | P0AEM9 | FALSE | FALSE | FALSE |
| 399 | ychJ | P37052 | TRUE | TRUE | FALSE |
| 400 | gfcC | P75883 | FALSE | FALSE | FALSE |
| 401 | sgbE | P37680 | TRUE | TRUE | TRUE |
| 402 | parC | P0AFI2 | FALSE | FALSE | FALSE |
| 403 | ydhJ | P76185 | FALSE | FALSE | FALSE |
| 404 | mrp | P0AF08 | TRUE | TRUE | FALSE |
| 405 | nrdH | P0AC65 | FALSE | MISSING | FALSE |
| 406 | ychF | P0ABU2 | FALSE | FALSE | FALSE |
| 407 | rlmE | P0C0R7 | FALSE | MISSING | FALSE |
| 408 | manB | P24175 | FALSE | TRUE | TRUE |
| 409 | yejO | P33924 | FALSE | TRUE | TRUE |
| 410 | kdgK | P37647 | FALSE | FALSE | FALSE |
| 411 | mdaB | P0AEY5 | FALSE | FALSE | FALSE |
| 412 | cueR | P0A9G4 | FALSE | FALSE | FALSE |
| 413 | tdk | P23331 | TRUE | TRUE | FALSE |
| 414 | ptsG | P69786 | FALSE | FALSE | FALSE |
| 415 | ycjM | P76041 | FALSE | FALSE | FALSE |
| 416 | arnD | P76472 | FALSE | TRUE | FALSE |
| 417 | nupC | P0AFF2 | FALSE | FALSE | FALSE |
| 418 | yoeB | P69348 | FALSE | FALSE | FALSE |
| 419 | thiB | P31550 | FALSE | MISSING | FALSE |
| 420 | wcaI | P32057 | FALSE | FALSE | FALSE |
| 421 | yhaJ | P67660 | FALSE | FALSE | FALSE |
| 422 | nth | P0AB83 | FALSE | FALSE | FALSE |
| 423 | napC | P0ABL5 | TRUE | TRUE | FALSE |
| 424 | mlaF | P63386 | FALSE | FALSE | FALSE |
| 425 | btuD | P06611 | FALSE | FALSE | FALSE |
| 426 | fumB | P14407 | FALSE | FALSE | FALSE |
| 427 | yahE | P77297 | FALSE | FALSE | FALSE |
| 428 | rseB | P0AFX9 | FALSE | FALSE | FALSE |
| 429 | lhgD | P37339 | FALSE | FALSE | FALSE |
| 430 | gntR | P0ACP5 | FALSE | FALSE | FALSE |
| 431 | cybB | P0ABE5 | FALSE | TRUE | FALSE |
| 432 | lsrK | P77432 | FALSE | FALSE | FALSE |
| 433 | selU | P33667 | FALSE | FALSE | FALSE |
| 434 | ylcJ | P0DPN3 | FALSE | MISSING | FALSE |
| 435 | yqhC | Q46855 | FALSE | FALSE | FALSE |
| 436 | lysS | P0A8N3 | FALSE | FALSE | FALSE |
| 437 | hyfE | P0AEW1 | FALSE | FALSE | FALSE |
| 438 | speA | P21170 | FALSE | FALSE | FALSE |
| 439 | dadX | P29012 | FALSE | FALSE | FALSE |
| 440 | rayT | Q47152 | TRUE | TRUE | TRUE |
| 441 | metN | P30750 | FALSE | FALSE | FALSE |
| 442 | dmsB | P18776 | TRUE | FALSE | FALSE |
| 443 | ftsP | P26648 | FALSE | FALSE | FALSE |
| 444 | yfjX | P52139 | FALSE | FALSE | FALSE |
| 445 | gstA | P0A9D2 | FALSE | FALSE | FALSE |
| 446 | yqjC | P42616 | FALSE | FALSE | FALSE |
| 447 | decR | P0ACJ5 | FALSE | FALSE | FALSE |
| 448 | sodC | P0AGD1 | TRUE | TRUE | FALSE |

Supplementary Table 3: E. coli predicted metal-binding proteome *(continued)*

| # | Gene Name | UniProt ID | MoM Prediction | M3D Prediction | GASS Prediction |
|---|---|---|---|---|---|
| 449 | pfkB | P06999 | FALSE | FALSE | FALSE |
| 450 | nrdA | P00452 | FALSE | FALSE | FALSE |
| 451 | narK | P10903 | FALSE | FALSE | FALSE |
| 452 | waaO | P27128 | FALSE | FALSE | FALSE |
| 453 | glpK | P0A6F3 | FALSE | FALSE | FALSE |
| 454 | glcD | P0AEP9 | FALSE | TRUE | FALSE |
| 455 | yphD | P77315 | FALSE | FALSE | FALSE |
| 456 | mltA | P0A935 | FALSE | FALSE | FALSE |
| 457 | ycbJ | P0AB03 | FALSE | FALSE | TRUE |
| 458 | cheB | P07330 | FALSE | TRUE | FALSE |
| 459 | amyA | P26612 | FALSE | FALSE | FALSE |
| 460 | chaC | P39163 | FALSE | FALSE | FALSE |
| 461 | hyfB | P23482 | FALSE | FALSE | FALSE |
| 462 | msrA | P0A744 | FALSE | FALSE | FALSE |
| 463 | tesA | P0ADA1 | FALSE | FALSE | FALSE |
| 464 | fixC | P68644 | FALSE | FALSE | FALSE |
| 465 | accD | P0A9Q5 | TRUE | TRUE | FALSE |
| 466 | bioA | P12995 | FALSE | FALSE | FALSE |
| 467 | agaA | P42906 | TRUE | FALSE | TRUE |
| 468 | astB | P76216 | FALSE | FALSE | TRUE |
| 469 | hisJ | P0AEU0 | FALSE | FALSE | FALSE |
| 470 | ydhF | P76187 | FALSE | FALSE | FALSE |
| 471 | surA | P0ABZ6 | FALSE | FALSE | FALSE |
| 472 | cecR | P0ACU0 | FALSE | TRUE | FALSE |
| 473 | yhdJ | P28638 | FALSE | FALSE | TRUE |
| 474 | ndk | P0A763 | FALSE | FALSE | FALSE |
| 475 | gcl | P0AEP7 | FALSE | TRUE | FALSE |
| 476 | waaF | P37692 | FALSE | FALSE | FALSE |
| 477 | ybeH | P0DP63 | FALSE | FALSE | FALSE |
| 478 | speC | P21169 | FALSE | FALSE | TRUE |
| 479 | chiP | P75733 | FALSE | FALSE | FALSE |
| 480 | dppC | P0AEG1 | FALSE | FALSE | FALSE |
| 481 | cadA | P0A9H3 | FALSE | FALSE | FALSE |
| 482 | holD | P28632 | FALSE | FALSE | FALSE |
| 483 | ydgJ | P77376 | FALSE | FALSE | FALSE |
| 484 | asnS | P0A8M0 | FALSE | FALSE | FALSE |
| 485 | insF4 | P0CF82 | FALSE | FALSE | FALSE |
| 486 | kdpF | P36937 | FALSE | MISSING | FALSE |
| 487 | rusA | P0AG74 | FALSE | FALSE | FALSE |
| 488 | rlmA | P36999 | TRUE | TRUE | FALSE |
| 489 | scpB | P52045 | FALSE | FALSE | FALSE |
| 490 | eptA | P30845 | TRUE | TRUE | FALSE |
| 491 | zntB | P64423 | FALSE | TRUE | FALSE |
| 492 | rsd | P0AFX4 | FALSE | FALSE | FALSE |
| 493 | mraZ | P22186 | FALSE | FALSE | FALSE |
| 494 | ygeY | P65807 | TRUE | TRUE | TRUE |
| 495 | yedE | P31064 | FALSE | FALSE | FALSE |
| 496 | argH | P11447 | FALSE | FALSE | FALSE |
| 497 | ybgA | P24252 | FALSE | FALSE | FALSE |
| 498 | dgcM | P77302 | TRUE | FALSE | FALSE |
| 499 | yibF | P0ACA1 | FALSE | FALSE | FALSE |
| 500 | yraI | P42914 | FALSE | MISSING | FALSE |
| 501 | ycjQ | P76043 | FALSE | TRUE | FALSE |
| 502 | yfcJ | P77549 | FALSE | FALSE | FALSE |
| 503 | ydcO | P76103 | FALSE | FALSE | FALSE |
| 504 | sgcX | P39366 | FALSE | TRUE | TRUE |

Supplementary Table 3: E. coli predicted metal-binding proteome *(continued)*

| # | Gene Name | UniProt ID | MoM Prediction | M3D Prediction | GASS Prediction |
|---|---|---|---|---|---|
| 505 | yqfA | P67153 | TRUE | TRUE | FALSE |
| 506 | rph | P0CG19 | FALSE | FALSE | FALSE |
| 507 | uacF | Q46820 | TRUE | FALSE | TRUE |
| 508 | metF | P0AEZ1 | FALSE | TRUE | FALSE |
| 509 | gltB | P09831 | FALSE | FALSE | FALSE |
| 510 | hycE | P16431 | TRUE | TRUE | FALSE |
| 511 | cdd | P0ABF6 | TRUE | TRUE | FALSE |
| 512 | rpoB | P0A8V2 | FALSE | FALSE | TRUE |
| 513 | pxpB | P0AAV4 | FALSE | MISSING | TRUE |
| 514 | glsA2 | P0A6W0 | FALSE | FALSE | FALSE |
| 515 | yehZ | P33362 | FALSE | MISSING | FALSE |
| 516 | ftsA | P0ABH0 | TRUE | FALSE | FALSE |
| 517 | torT | P38683 | FALSE | FALSE | FALSE |
| 518 | aceE | P0AFG8 | TRUE | TRUE | FALSE |
| 519 | fhuC | P07821 | FALSE | FALSE | FALSE |
| 520 | nudL | P43337 | FALSE | FALSE | FALSE |
| 521 | atpD | P0ABB4 | FALSE | FALSE | FALSE |
| 522 | hcaD | P77650 | FALSE | FALSE | FALSE |
| 523 | sufD | P77689 | FALSE | FALSE | FALSE |
| 524 | gshA | P0A6W9 | FALSE | TRUE | FALSE |
| 525 | glyA | P0A825 | FALSE | FALSE | FALSE |
| 526 | yebC | P0A8A0 | FALSE | FALSE | FALSE |
| 527 | yajQ | P0A8E7 | FALSE | FALSE | FALSE |
| 528 | yfdM | P76509 | FALSE | MISSING | FALSE |
| 529 | nikC | P0AFA9 | TRUE | TRUE | FALSE |
| 530 | phnM | P16689 | TRUE | TRUE | TRUE |
| 531 | pgl | P52697 | TRUE | FALSE | FALSE |
| 532 | ydjJ | P77280 | TRUE | TRUE | FALSE |
| 533 | yigB | P0ADP0 | FALSE | FALSE | FALSE |
| 534 | yagG | P75683 | FALSE | FALSE | FALSE |
| 535 | ygjK | P42592 | FALSE | FALSE | FALSE |
| 536 | yfdR | P76514 | FALSE | TRUE | FALSE |
| 537 | yjfI | P0AF76 | FALSE | FALSE | FALSE |
| 538 | nhoA | P77567 | FALSE | FALSE | FALSE |
| 539 | rarD | P27844 | FALSE | FALSE | FALSE |
| 540 | elaA | P0AEH3 | TRUE | FALSE | FALSE |
| 541 | iscX | P0C0L9 | FALSE | FALSE | FALSE |
| 542 | gltI | P37902 | FALSE | MISSING | FALSE |
| 543 | dctR | P37195 | FALSE | FALSE | FALSE |
| 544 | asnB | P22106 | FALSE | TRUE | FALSE |
| 545 | ydfI | P77260 | TRUE | FALSE | FALSE |
| 546 | aroH | P00887 | FALSE | FALSE | FALSE |
| 547 | lpxL | P0ACV0 | FALSE | FALSE | FALSE |
| 548 | mtgA | P46022 | FALSE | FALSE | TRUE |
| 549 | ugpE | P10906 | FALSE | FALSE | FALSE |
| 550 | ssnA | Q46812 | TRUE | TRUE | TRUE |
| 551 | ybiT | P0A9U3 | FALSE | FALSE | FALSE |
| 552 | glpA | P0A9C0 | TRUE | FALSE | FALSE |
| 553 | yeiP | P0A6N8 | FALSE | FALSE | FALSE |
| 554 | uxaA | P42604 | TRUE | TRUE | TRUE |
| 555 | rseP | P0AEH1 | TRUE | TRUE | FALSE |
| 556 | rluB | P37765 | FALSE | FALSE | FALSE |
| 557 | ydeT | P76137 | FALSE | FALSE | FALSE |
| 558 | ddpB | P77308 | FALSE | FALSE | FALSE |
| 559 | sgbU | P37679 | FALSE | TRUE | FALSE |
| 560 | caiB | P31572 | FALSE | FALSE | FALSE |

Supplementary Table 3: E. coli predicted metal-binding proteome *(continued)*

| # | Gene Name | UniProt ID | MoM Prediction | M3D Prediction | GASS Prediction |
|---|---|---|---|---|---|
| 561 | fepB | P0AEL6 | FALSE | FALSE | FALSE |
| 562 | yhhH | P28911 | FALSE | TRUE | FALSE |
| 563 | livG | P0A9S7 | FALSE | FALSE | FALSE |
| 564 | frlB | P0AC00 | FALSE | FALSE | FALSE |
| 565 | ydiO | P0A9U8 | FALSE | FALSE | FALSE |
| 566 | lplT | P39196 | FALSE | FALSE | FALSE |
| 567 | btuC | P06609 | FALSE | FALSE | FALSE |
| 568 | mepK | P0AB06 | TRUE | TRUE | TRUE |
| 569 | nlpI | P0AFB1 | FALSE | TRUE | FALSE |
| 570 | gcd | P15877 | FALSE | FALSE | FALSE |
| 571 | casE | Q46897 | FALSE | FALSE | FALSE |
| 572 | torC | P33226 | TRUE | TRUE | FALSE |
| 573 | tcyL | P0AFT2 | FALSE | FALSE | FALSE |
| 574 | lepB | P00803 | FALSE | FALSE | FALSE |
| 575 | cspE | P0A972 | FALSE | FALSE | FALSE |
| 576 | ppiB | P23869 | FALSE | FALSE | FALSE |
| 577 | ypeB | P0AD40 | FALSE | MISSING | FALSE |
| 578 | lolA | P61316 | FALSE | FALSE | TRUE |
| 579 | treA | P13482 | FALSE | FALSE | FALSE |
| 580 | dcuD | P45428 | FALSE | FALSE | FALSE |
| 581 | pepE | P0A7C6 | FALSE | FALSE | FALSE |
| 582 | yhdX | P45767 | FALSE | FALSE | FALSE |
| 583 | paaE | P76081 | TRUE | FALSE | FALSE |
| 584 | lpxH | P43341 | TRUE | TRUE | TRUE |
| 585 | yfdK | P77656 | FALSE | MISSING | FALSE |
| 586 | dhaK | P76015 | FALSE | TRUE | FALSE |
| 587 | dppB | P0AEF8 | FALSE | FALSE | FALSE |
| 588 | kefG | P0A756 | FALSE | FALSE | FALSE |
| 589 | intB | P39347 | FALSE | TRUE | FALSE |
| 590 | malK | P68187 | FALSE | FALSE | FALSE |
| 591 | tsaB | P76256 | FALSE | FALSE | FALSE |
| 592 | pbpC | P76577 | FALSE | FALSE | FALSE |
| 593 | ghrB | P37666 | FALSE | FALSE | FALSE |
| 594 | gapA | P0A9B2 | FALSE | TRUE | FALSE |
| 595 | ibsC | C1P615 | FALSE | MISSING | FALSE |
| 596 | hemG | P0ACB4 | FALSE | FALSE | FALSE |
| 597 | ypdE | P77585 | TRUE | TRUE | TRUE |
| 598 | allH | P0AAS5 | FALSE | FALSE | FALSE |
| 599 | pheS | P08312 | FALSE | FALSE | FALSE |
| 600 | glnH | P0AEQ3 | FALSE | FALSE | FALSE |
| 601 | umuC | P04152 | FALSE | FALSE | FALSE |
| 602 | ttdR | P45463 | FALSE | FALSE | FALSE |
| 603 | ybhB | P12994 | FALSE | FALSE | FALSE |
| 604 | yphG | P76585 | FALSE | FALSE | FALSE |
| 605 | yejA | P33913 | FALSE | FALSE | FALSE |
| 606 | paaY | P77181 | FALSE | TRUE | FALSE |
| 607 | glcF | P52074 | TRUE | TRUE | TRUE |
| 608 | yjbH | P32689 | FALSE | FALSE | FALSE |
| 609 | rlmL | P75864 | FALSE | TRUE | FALSE |
| 610 | hcp | P75825 | TRUE | TRUE | FALSE |
| 611 | ansA | P0A962 | FALSE | FALSE | FALSE |
| 612 | ahpF | P35340 | FALSE | FALSE | FALSE |
| 613 | pmbA | P0AFK0 | FALSE | FALSE | TRUE |
| 614 | yagU | P0AAA1 | FALSE | FALSE | FALSE |
| 615 | ybcO | P68661 | TRUE | TRUE | FALSE |
| 616 | fdhE | P13024 | TRUE | TRUE | TRUE |

Supplementary Table 3: E. coli predicted metal-binding proteome *(continued)*

| # | Gene Name | UniProt ID | MoM Prediction | M3D Prediction | GASS Prediction |
|---|---|---|---|---|---|
| 617 | pbpG | P0AFI5 | FALSE | FALSE | FALSE |
| 618 | mtnN | P0AF12 | TRUE | FALSE | FALSE |
| 619 | gnsA | P0AC92 | FALSE | MISSING | FALSE |
| 620 | fucI | P69922 | FALSE | TRUE | FALSE |
| 621 | pgpB | P0A924 | FALSE | FALSE | FALSE |
| 622 | nagB | P0A759 | FALSE | FALSE | FALSE |
| 623 | yhfX | P45550 | FALSE | FALSE | FALSE |
| 624 | ydaF | P0ACW0 | FALSE | MISSING | FALSE |
| 625 | mgtS | A5A616 | FALSE | MISSING | FALSE |
| 626 | mdtM | P39386 | FALSE | FALSE | FALSE |
| 627 | sodB | P0AGD3 | TRUE | TRUE | FALSE |
| 628 | ppdB | P08371 | FALSE | FALSE | FALSE |
| 629 | fldB | P0ABY4 | FALSE | FALSE | FALSE |
| 630 | allR | P0ACN4 | FALSE | FALSE | FALSE |
| 631 | yhfS | P45545 | FALSE | FALSE | FALSE |
| 632 | alaS | P00957 | TRUE | TRUE | TRUE |
| 633 | fmt | P23882 | FALSE | FALSE | FALSE |
| 634 | tyrS | P0AGJ9 | FALSE | FALSE | FALSE |
| 635 | trmL | P0AGJ7 | FALSE | TRUE | FALSE |
| 636 | gspK | P45762 | FALSE | FALSE | FALSE |
| 637 | torA | P33225 | FALSE | TRUE | TRUE |
| 638 | ytfE | P69506 | FALSE | TRUE | FALSE |
| 639 | hisS | P60906 | FALSE | FALSE | FALSE |
| 640 | nudE | P45799 | FALSE | FALSE | FALSE |
| 641 | rna | P21338 | FALSE | FALSE | FALSE |
| 642 | proB | P0A7B5 | FALSE | FALSE | TRUE |
| 643 | glyQ | P00960 | FALSE | FALSE | FALSE |
| 644 | ypdF | P76524 | FALSE | TRUE | FALSE |
| 645 | caiD | P31551 | FALSE | FALSE | FALSE |
| 646 | cutC | P67826 | FALSE | TRUE | TRUE |
| 647 | ybiY | P75794 | TRUE | FALSE | FALSE |
| 648 | rpmD | P0AG51 | FALSE | MISSING | FALSE |
| 649 | aqpZ | P60844 | FALSE | FALSE | FALSE |
| 650 | gltL | P0AAG3 | FALSE | FALSE | FALSE |
| 651 | nemA | P77258 | FALSE | FALSE | FALSE |
| 652 | mhpE | P51020 | TRUE | TRUE | FALSE |
| 653 | ppsR | P0A8A4 | FALSE | FALSE | FALSE |
| 654 | nuoE | P0AFD1 | TRUE | FALSE | FALSE |
| 655 | gatZ | P0C8J8 | TRUE | TRUE | TRUE |
| 656 | paaI | P76084 | FALSE | FALSE | TRUE |
| 657 | fadH | P42593 | FALSE | FALSE | FALSE |
| 658 | cysG | P0AEA8 | FALSE | FALSE | FALSE |
| 659 | hemC | P06983 | FALSE | FALSE | FALSE |
| 660 | yidZ | P31463 | FALSE | FALSE | FALSE |
| 661 | exbB | P0ABU7 | FALSE | FALSE | FALSE |
| 662 | bluR | P75989 | FALSE | FALSE | FALSE |
| 663 | perR | Q57083 | FALSE | FALSE | FALSE |
| 664 | yieP | P31475 | FALSE | TRUE | FALSE |
| 665 | yaeP | P0A8K5 | FALSE | MISSING | FALSE |
| 666 | sdhA | P0AC41 | FALSE | TRUE | TRUE |
| 667 | miaA | P16384 | FALSE | FALSE | FALSE |
| 668 | osmC | P0C0L2 | FALSE | FALSE | FALSE |
| 669 | kduD | P37769 | FALSE | FALSE | FALSE |
| 670 | ccmA | P33931 | FALSE | FALSE | FALSE |
| 671 | ydcT | P77795 | FALSE | FALSE | FALSE |
| 672 | gfcD | P75882 | FALSE | FALSE | FALSE |

Supplementary Table 3: E. coli predicted metal-binding proteome *(continued)*

| # | Gene Name | UniProt ID | MoM Prediction | M3D Prediction | GASS Prediction |
|---|---|---|---|---|---|
| 673 | xapR | P23841 | FALSE | TRUE | FALSE |
| 674 | gatY | P0C8J6 | TRUE | TRUE | FALSE |
| 675 | nuoL | P33607 | TRUE | FALSE | TRUE |
| 676 | iscS | P0A6B7 | FALSE | FALSE | FALSE |
| 677 | wbbL | P36667 | FALSE | FALSE | FALSE |
| 678 | atpG | P0ABA6 | FALSE | FALSE | FALSE |
| 679 | ilvA | P04968 | FALSE | FALSE | FALSE |
| 680 | pdxA | P19624 | FALSE | TRUE | FALSE |
| 681 | rpmI | P0A7Q1 | FALSE | FALSE | FALSE |
| 682 | panC | P31663 | FALSE | FALSE | FALSE |
| 683 | galF | P0AAB6 | FALSE | FALSE | FALSE |
| 684 | ybfQ | Q2EEQ8 | FALSE | FALSE | FALSE |
| 685 | ygaC | P0AD53 | FALSE | FALSE | FALSE |
| 686 | nuoI | P0AFD6 | FALSE | FALSE | FALSE |
| 687 | bglB | P11988 | FALSE | TRUE | FALSE |
| 688 | yjiL | P39383 | FALSE | TRUE | FALSE |
| 689 | araA | P08202 | FALSE | TRUE | FALSE |
| 690 | ybhA | P21829 | TRUE | FALSE | FALSE |
| 691 | ydiT | P77714 | FALSE | FALSE | FALSE |
| 692 | agaS | P42907 | FALSE | FALSE | TRUE |
| 693 | phnG | P16685 | FALSE | FALSE | FALSE |
| 694 | ydcK | P76100 | FALSE | FALSE | TRUE |
| 695 | yefM | P69346 | FALSE | FALSE | FALSE |
| 696 | iroK | U3PVA8 | FALSE | MISSING | FALSE |
| 697 | araJ | P23910 | FALSE | FALSE | FALSE |
| 698 | yjeJ | P39279 | FALSE | FALSE | TRUE |
| 699 | trxA | P0AA25 | FALSE | FALSE | FALSE |
| 700 | ansB | P00805 | FALSE | FALSE | FALSE |
| 701 | trpC | P00909 | FALSE | FALSE | FALSE |
| 702 | yrbL | P64610 | FALSE | FALSE | TRUE |
| 703 | cpdB | P08331 | TRUE | FALSE | FALSE |
| 704 | paaK | P76085 | FALSE | FALSE | FALSE |
| 705 | recA | P0A7G6 | FALSE | FALSE | FALSE |
| 706 | hicA | P76106 | FALSE | FALSE | FALSE |
| 707 | cspC | P0A9Y6 | FALSE | MISSING | FALSE |
| 708 | ybaQ | P0A9T6 | FALSE | FALSE | FALSE |
| 709 | yihT | P32141 | FALSE | FALSE | FALSE |
| 710 | ibsD | C1P616 | FALSE | MISSING | FALSE |
| 711 | rplR | P0C018 | FALSE | MISSING | FALSE |
| 712 | gmk | P60546 | FALSE | FALSE | FALSE |
| 713 | yidA | P0A8Y5 | FALSE | FALSE | TRUE |
| 714 | malE | P0AEX9 | FALSE | FALSE | FALSE |
| 715 | yceJ | P75925 | FALSE | TRUE | FALSE |
| 716 | narY | P19318 | TRUE | FALSE | FALSE |
| 717 | ahpC | P0AE08 | FALSE | FALSE | FALSE |
| 718 | evgA | P0ACZ4 | FALSE | TRUE | FALSE |
| 719 | paaX | P76086 | FALSE | FALSE | FALSE |
| 720 | rbsD | P04982 | FALSE | FALSE | FALSE |
| 721 | rlmC | P75817 | FALSE | FALSE | FALSE |
| 722 | yfeN | P45564 | FALSE | FALSE | FALSE |
| 723 | yncL | A5A615 | FALSE | MISSING | FALSE |
| 724 | soxS | P0A9E2 | FALSE | FALSE | FALSE |
| 725 | dacD | P33013 | FALSE | FALSE | FALSE |
| 726 | alsB | P39265 | FALSE | MISSING | FALSE |
| 727 | yiaO | P37676 | FALSE | FALSE | FALSE |
| 728 | yghO | Q46840 | FALSE | FALSE | FALSE |

Supplementary Table 3: E. coli predicted metal-binding proteome *(continued)*

| # | Gene Name | UniProt ID | MoM Prediction | M3D Prediction | GASS Prediction |
|---|---|---|---|---|---|
| 729 | yrdB | P45795 | FALSE | FALSE | FALSE |
| 730 | add | P22333 | TRUE | TRUE | FALSE |
| 731 | mazF | P0AE70 | FALSE | FALSE | FALSE |
| 732 | ddpA | P76128 | FALSE | FALSE | FALSE |
| 733 | cadB | P0AAE8 | FALSE | FALSE | FALSE |
| 734 | aphA | P0AE22 | FALSE | TRUE | FALSE |
| 735 | entD | P19925 | FALSE | FALSE | FALSE |
| 736 | yehA | P33340 | FALSE | FALSE | FALSE |
| 737 | yciY | A5A613 | FALSE | FALSE | FALSE |
| 738 | srlR | P15082 | FALSE | FALSE | FALSE |
| 739 | eno | P0A6P9 | FALSE | FALSE | FALSE |
| 740 | waaS | P27126 | FALSE | FALSE | TRUE |
| 741 | argB | P0A6C8 | FALSE | FALSE | FALSE |
| 742 | fliI | P52612 | FALSE | FALSE | FALSE |
| 743 | yahD | P77736 | FALSE | FALSE | FALSE |
| 744 | ugpB | P0AG80 | FALSE | FALSE | FALSE |
| 745 | galS | P25748 | FALSE | FALSE | FALSE |
| 746 | nuoG | P33602 | FALSE | FALSE | TRUE |
| 747 | agp | P19926 | FALSE | FALSE | FALSE |
| 748 | yecD | P0ADI7 | FALSE | FALSE | FALSE |
| 749 | kefF | P0A754 | FALSE | TRUE | FALSE |
| 750 | gpmI | P37689 | TRUE | TRUE | FALSE |
| 751 | yidJ | P31447 | FALSE | TRUE | FALSE |
| 752 | elyC | P0AB01 | FALSE | FALSE | FALSE |
| 753 | msrQ | P76343 | FALSE | TRUE | FALSE |
| 754 | mocA | Q46810 | FALSE | TRUE | FALSE |
| 755 | ynfF | P77783 | TRUE | TRUE | TRUE |
| 756 | ssuA | P75853 | FALSE | FALSE | FALSE |
| 757 | queF | Q46920 | FALSE | FALSE | FALSE |
| 758 | pnuC | P0AFK2 | FALSE | FALSE | FALSE |
| 759 | yjfC | P33222 | FALSE | FALSE | TRUE |
| 760 | yagF | P77596 | TRUE | TRUE | TRUE |
| 761 | rluA | P0AA37 | TRUE | FALSE | FALSE |
| 762 | hisD | P06988 | FALSE | FALSE | FALSE |
| 763 | rlmH | P0A8I8 | FALSE | FALSE | FALSE |
| 764 | pdhR | P0ACL9 | TRUE | FALSE | FALSE |
| 765 | rhaT | P27125 | FALSE | FALSE | FALSE |
| 766 | pdxK | P40191 | FALSE | FALSE | FALSE |
| 767 | yneE | P76146 | FALSE | TRUE | FALSE |
| 768 | yahG | P77221 | FALSE | FALSE | FALSE |
| 769 | hybG | P0AAM7 | FALSE | FALSE | FALSE |
| 770 | nmpC | P21420 | FALSE | FALSE | FALSE |
| 771 | prkB | P0AEX5 | FALSE | FALSE | FALSE |
| 772 | rhmR | P77732 | FALSE | FALSE | FALSE |
| 773 | dinI | P0ABR1 | FALSE | FALSE | FALSE |
| 774 | hha | P0ACE3 | FALSE | MISSING | FALSE |
| 775 | accA | P0ABD5 | FALSE | TRUE | FALSE |
| 776 | bglG | P11989 | TRUE | FALSE | FALSE |
| 777 | yaeQ | P0AA97 | FALSE | FALSE | FALSE |
| 778 | hda | P69931 | FALSE | FALSE | FALSE |
| 779 | gudX | Q46915 | FALSE | TRUE | FALSE |
| 780 | dcyD | P76316 | FALSE | FALSE | FALSE |
| 781 | yceF | P0A729 | FALSE | FALSE | FALSE |
| 782 | pptA | P31992 | FALSE | MISSING | FALSE |
| 783 | ampH | P0AD70 | FALSE | FALSE | FALSE |
| 784 | rlmD | P55135 | TRUE | FALSE | FALSE |

Supplementary Table 3: E. coli predicted metal-binding proteome *(continued)*

| # | Gene Name | UniProt ID | MoM Prediction | M3D Prediction | GASS Prediction |
|---|---|---|---|---|---|
| 785 | nrfE | P32710 | FALSE | TRUE | FALSE |
| 786 | alaA | P0A959 | FALSE | FALSE | FALSE |
| 787 | gph | P32662 | FALSE | TRUE | FALSE |
| 788 | dbpA | P21693 | TRUE | FALSE | FALSE |
| 789 | yicI | P31434 | TRUE | FALSE | FALSE |
| 790 | arnB | P77690 | TRUE | FALSE | FALSE |
| 791 | prpB | P77541 | FALSE | FALSE | FALSE |
| 792 | atoD | P76458 | FALSE | FALSE | FALSE |
| 793 | arsB | P0AB93 | FALSE | FALSE | FALSE |
| 794 | hemN | P32131 | FALSE | FALSE | TRUE |
| 795 | prpC | P31660 | FALSE | FALSE | FALSE |
| 796 | zapC | P75862 | FALSE | FALSE | FALSE |
| 797 | ygjQ | P42598 | FALSE | FALSE | FALSE |
| 798 | rarA | P0AAZ4 | FALSE | TRUE | FALSE |
| 799 | aegA | P37127 | TRUE | TRUE | TRUE |
| 800 | yjdI | P0AF59 | FALSE | FALSE | FALSE |
| 801 | hydN | P0AAK4 | FALSE | FALSE | TRUE |
| 802 | pntA | P07001 | FALSE | FALSE | FALSE |
| 803 | aspS | P21889 | FALSE | TRUE | FALSE |
| 804 | fucK | P11553 | FALSE | FALSE | FALSE |
| 805 | ycjU | P77366 | FALSE | FALSE | FALSE |
| 806 | hisC | P06986 | FALSE | MISSING | FALSE |
| 807 | torD | P36662 | FALSE | FALSE | FALSE |
| 808 | sdhC | P69054 | FALSE | FALSE | FALSE |
| 809 | waaB | P27127 | FALSE | FALSE | FALSE |
| 810 | hypE | P24193 | FALSE | FALSE | FALSE |
| 811 | cmoM | P36566 | FALSE | FALSE | FALSE |
| 812 | yghX | Q7DFU6 | FALSE | MISSING | FALSE |
| 813 | zapB | P0AF36 | FALSE | FALSE | FALSE |
| 814 | serB | P0AGB0 | FALSE | FALSE | FALSE |
| 815 | dicA | P06966 | FALSE | FALSE | FALSE |
| 816 | ilvD | P05791 | FALSE | FALSE | TRUE |
| 817 | paaB | P76078 | FALSE | FALSE | FALSE |
| 818 | adk | P69441 | FALSE | FALSE | FALSE |
| 819 | mhpB | P0ABR9 | FALSE | TRUE | FALSE |
| 820 | bfd | P0AE56 | TRUE | FALSE | TRUE |
| 821 | dtpB | P36837 | FALSE | FALSE | FALSE |
| 822 | rnlB | P52130 | FALSE | FALSE | FALSE |
| 823 | yfcS | P77599 | FALSE | FALSE | FALSE |
| 824 | mlaB | P64602 | FALSE | FALSE | FALSE |
| 825 | yahK | P75691 | TRUE | TRUE | TRUE |
| 826 | yjhG | P39358 | FALSE | FALSE | TRUE |
| 827 | yhjX | P37662 | FALSE | FALSE | FALSE |
| 828 | dgcP | P76245 | FALSE | FALSE | TRUE |
| 829 | yedD | P31063 | FALSE | FALSE | FALSE |
| 830 | otnI | Q46891 | FALSE | FALSE | FALSE |
| 831 | astE | P76215 | TRUE | TRUE | FALSE |
| 832 | yfcH | P77775 | FALSE | FALSE | FALSE |
| 833 | gcvH | P0A6T9 | FALSE | TRUE | FALSE |
| 834 | tfaQ | P76155 | FALSE | TRUE | FALSE |
| 835 | wcaC | P71237 | FALSE | FALSE | TRUE |
| 836 | nrdD | P28903 | TRUE | TRUE | FALSE |
| 837 | tusD | P45532 | FALSE | FALSE | FALSE |
| 838 | thiG | P30139 | FALSE | FALSE | FALSE |
| 839 | yoaC | P64490 | FALSE | MISSING | FALSE |
| 840 | puuE | P50457 | FALSE | FALSE | FALSE |

Supplementary Table 3: E. coli predicted metal-binding proteome *(continued)*

| # | Gene Name | UniProt ID | MoM Prediction | M3D Prediction | GASS Prediction |
|---|---|---|---|---|---|
| 841 | ybhC | P46130 | FALSE | FALSE | FALSE |
| 842 | umuD | P0AG11 | FALSE | MISSING | FALSE |
| 843 | ttdT | P39414 | FALSE | FALSE | TRUE |
| 844 | argI | P04391 | FALSE | TRUE | TRUE |
| 845 | nirD | P0A9I8 | FALSE | FALSE | FALSE |
| 846 | yfeW | P77619 | FALSE | TRUE | TRUE |
| 847 | yobB | P76280 | FALSE | FALSE | FALSE |
| 848 | fecD | P15029 | FALSE | FALSE | FALSE |
| 849 | yeiG | P33018 | FALSE | FALSE | FALSE |
| 850 | yibN | P0AG27 | FALSE | FALSE | FALSE |
| 851 | rsmI | P67087 | FALSE | FALSE | FALSE |
| 852 | mdtG | P25744 | FALSE | FALSE | FALSE |
| 853 | napB | P0ABL3 | FALSE | TRUE | FALSE |
| 854 | yfbR | P76491 | FALSE | TRUE | FALSE |
| 855 | feaB | P80668 | FALSE | FALSE | TRUE |
| 856 | hemD | P09126 | FALSE | FALSE | FALSE |
| 857 | cynT | P0ABE9 | TRUE | TRUE | FALSE |
| 858 | tauB | Q47538 | FALSE | FALSE | FALSE |
| 859 | yeeN | P0A8A2 | FALSE | FALSE | FALSE |
| 860 | fadE | Q47146 | FALSE | TRUE | TRUE |
| 861 | pabA | P00903 | FALSE | FALSE | FALSE |
| 862 | ispA | P22939 | FALSE | FALSE | FALSE |
| 863 | ilvE | P0AB80 | FALSE | FALSE | FALSE |
| 864 | glcE | P52073 | FALSE | FALSE | FALSE |
| 865 | yceM | P75931 | FALSE | FALSE | FALSE |
| 866 | ahr | P27250 | TRUE | TRUE | TRUE |
| 867 | kduI | Q46938 | FALSE | TRUE | TRUE |
| 868 | rplQ | P0AG44 | FALSE | FALSE | FALSE |
| 869 | ygbT | Q46896 | FALSE | FALSE | FALSE |
| 870 | glmM | P31120 | FALSE | FALSE | TRUE |
| 871 | dinF | P28303 | FALSE | FALSE | FALSE |
| 872 | bcsC | P37650 | FALSE | TRUE | FALSE |
| 873 | ydiB | P0A6D5 | FALSE | FALSE | FALSE |
| 874 | tamA | P0ADE4 | FALSE | FALSE | FALSE |
| 875 | ycaI | P37443 | TRUE | TRUE | FALSE |
| 876 | yceB | P0AB26 | FALSE | FALSE | FALSE |
| 877 | thiE | P30137 | FALSE | FALSE | FALSE |
| 878 | livJ | P0AD96 | FALSE | FALSE | FALSE |
| 879 | truA | P07649 | FALSE | TRUE | FALSE |
| 880 | dgkA | P0ABN1 | FALSE | FALSE | FALSE |
| 881 | hemL | P23893 | FALSE | FALSE | FALSE |
| 882 | sdhD | P0AC44 | FALSE | FALSE | FALSE |
| 883 | ackA | P0A6A3 | FALSE | FALSE | FALSE |
| 884 | ygcR | Q46908 | FALSE | FALSE | FALSE |
| 885 | ariR | P75993 | FALSE | FALSE | FALSE |
| 886 | tauC | Q47539 | FALSE | FALSE | FALSE |
| 887 | ynbB | P76091 | FALSE | FALSE | TRUE |
| 888 | potF | P31133 | FALSE | FALSE | FALSE |
| 889 | yfbM | P76483 | FALSE | FALSE | FALSE |
| 890 | nudJ | P0AEI6 | FALSE | TRUE | FALSE |
| 891 | htrL | P25666 | FALSE | FALSE | FALSE |
| 892 | lpxA | P0A722 | FALSE | FALSE | FALSE |
| 893 | lacZ | P00722 | FALSE | TRUE | FALSE |
| 894 | cobT | P36562 | FALSE | FALSE | FALSE |
| 895 | yebV | P64503 | FALSE | FALSE | FALSE |
| 896 | yiaT | P37681 | FALSE | FALSE | FALSE |

Supplementary Table 3: E. coli predicted metal-binding proteome *(continued)*

| # | Gene Name | UniProt ID | MoM Prediction | M3D Prediction | GASS Prediction |
|---|---|---|---|---|---|
| 897 | allD | P77555 | FALSE | FALSE | FALSE |
| 898 | ybiS | P0AAX8 | FALSE | FALSE | FALSE |
| 899 | entC | P0AEJ2 | FALSE | FALSE | FALSE |
| 900 | sgcC | P39365 | FALSE | FALSE | FALSE |
| 901 | dppA | P23847 | FALSE | FALSE | FALSE |
| 902 | etp | P0ACZ2 | FALSE | FALSE | FALSE |
| 903 | dinJ | Q47150 | FALSE | FALSE | FALSE |
| 904 | yecN | P64515 | FALSE | FALSE | FALSE |
| 905 | tas | P0A9T4 | TRUE | FALSE | TRUE |
| 906 | csqR | P32144 | FALSE | FALSE | TRUE |
| 907 | lysR | P03030 | FALSE | TRUE | FALSE |
| 908 | ygfF | P52037 | FALSE | FALSE | FALSE |
| 909 | greA | P0A6W5 | FALSE | FALSE | FALSE |
| 910 | phnH | P16686 | FALSE | FALSE | FALSE |
| 911 | selO | P77649 | FALSE | FALSE | FALSE |
| 912 | ycfS | P75954 | FALSE | FALSE | FALSE |
| 913 | aldA | P25553 | FALSE | TRUE | FALSE |
| 914 | yhiD | P0AFV2 | FALSE | FALSE | FALSE |
| 915 | ybgO | P75748 | FALSE | FALSE | FALSE |
| 916 | ycjS | P77503 | TRUE | FALSE | FALSE |
| 917 | idi | Q46822 | FALSE | TRUE | FALSE |
| 918 | mntR | P0A9F1 | FALSE | FALSE | TRUE |
| 919 | manC | P24174 | FALSE | TRUE | TRUE |
| 920 | orn | P0A784 | FALSE | FALSE | FALSE |
| 921 | proC | P0A9L8 | FALSE | FALSE | FALSE |
| 922 | hyuA | Q46806 | TRUE | TRUE | TRUE |
| 923 | sucC | P0A836 | FALSE | FALSE | FALSE |
| 924 | galT | P09148 | TRUE | TRUE | FALSE |
| 925 | nikR | P0A6Z6 | TRUE | TRUE | FALSE |
| 926 | glnD | P27249 | FALSE | TRUE | FALSE |
| 927 | talB | P0A870 | FALSE | FALSE | FALSE |
| 928 | lacA | P07464 | FALSE | FALSE | FALSE |
| 929 | lldP | P33231 | FALSE | FALSE | TRUE |
| 930 | rplP | P0ADY7 | FALSE | MISSING | FALSE |
| 931 | tusC | P45531 | FALSE | MISSING | FALSE |
| 932 | leuA | P09151 | TRUE | TRUE | FALSE |
| 933 | udp | P12758 | FALSE | FALSE | FALSE |
| 934 | yddE | P37757 | FALSE | FALSE | FALSE |
| 935 | fumD | P0ACX5 | FALSE | FALSE | FALSE |
| 936 | thiS | O32583 | FALSE | FALSE | FALSE |
| 937 | pflA | P0A9N4 | FALSE | FALSE | FALSE |
| 938 | glnQ | P10346 | FALSE | MISSING | FALSE |
| 939 | cbpM | P63264 | FALSE | FALSE | FALSE |
| 940 | eco | P23827 | FALSE | FALSE | FALSE |
| 941 | yfjP | P52131 | FALSE | FALSE | FALSE |
| 942 | yfgD | P76569 | FALSE | MISSING | TRUE |
| 943 | yfbL | P76482 | FALSE | FALSE | FALSE |
| 944 | yehX | P33360 | FALSE | FALSE | FALSE |
| 945 | ybiA | P30176 | FALSE | FALSE | FALSE |
| 946 | ybcH | P37325 | FALSE | FALSE | FALSE |
| 947 | yaiX | P75697 | FALSE | FALSE | FALSE |
| 948 | mnmA | P25745 | FALSE | FALSE | FALSE |
| 949 | hslR | P0ACG8 | FALSE | FALSE | FALSE |
| 950 | rdgB | P52061 | FALSE | FALSE | FALSE |
| 951 | oxyR | P0ACQ4 | FALSE | FALSE | TRUE |
| 952 | yfeK | Q47702 | FALSE | FALSE | FALSE |

Supplementary Table 3: E. coli predicted metal-binding proteome *(continued)*

| # | Gene Name | UniProt ID | MoM Prediction | M3D Prediction | GASS Prediction |
|---|---|---|---|---|---|
| 953 | tusE | P0AB18 | FALSE | MISSING | FALSE |
| 954 | yejF | P33916 | FALSE | FALSE | FALSE |
| 955 | ttdA | P05847 | TRUE | FALSE | FALSE |
| 956 | truD | Q57261 | FALSE | FALSE | FALSE |
| 957 | chpS | P08365 | FALSE | MISSING | FALSE |
| 958 | pgaA | P69434 | TRUE | FALSE | FALSE |
| 959 | yjiK | P39382 | FALSE | FALSE | FALSE |
| 960 | rpsK | P0A7R9 | FALSE | MISSING | FALSE |
| 961 | yhfU | P64631 | FALSE | FALSE | FALSE |
| 962 | alr | P0A6B4 | FALSE | TRUE | FALSE |
| 963 | dps | P0ABT2 | TRUE | FALSE | TRUE |
| 964 | ymjA | P0ACV8 | FALSE | FALSE | FALSE |
| 965 | yadE | P31666 | TRUE | TRUE | FALSE |
| 966 | ygiC | P0ADT5 | FALSE | FALSE | FALSE |
| 967 | casD | Q46898 | FALSE | FALSE | FALSE |
| 968 | fadA | P21151 | TRUE | TRUE | TRUE |
| 969 | elbB | P0ABU5 | FALSE | FALSE | FALSE |
| 970 | bolA | P0ABE2 | FALSE | FALSE | FALSE |
| 971 | topB | P14294 | FALSE | FALSE | FALSE |
| 972 | rspA | P38104 | FALSE | FALSE | TRUE |
| 973 | abgA | P77357 | TRUE | TRUE | TRUE |
| 974 | hslO | P0A6Y5 | TRUE | TRUE | TRUE |
| 975 | wzzE | P0AG00 | FALSE | FALSE | FALSE |
| 976 | rpmH | P0A7P5 | FALSE | FALSE | FALSE |
| 977 | scpA | P27253 | FALSE | FALSE | TRUE |
| 978 | rspR | P0ACM2 | TRUE | TRUE | FALSE |
| 979 | fbaB | P0A991 | FALSE | FALSE | FALSE |
| 980 | cysQ | P22255 | FALSE | FALSE | FALSE |
| 981 | gcvT | P27248 | FALSE | FALSE | FALSE |
| 982 | nupX | P33021 | FALSE | FALSE | FALSE |
| 983 | pstC | P0AGH8 | FALSE | FALSE | FALSE |
| 984 | yciE | P21363 | FALSE | TRUE | FALSE |
| 985 | metJ | P0A8U6 | FALSE | FALSE | FALSE |
| 986 | yfbN | P76484 | FALSE | FALSE | FALSE |
| 987 | tag | P05100 | TRUE | TRUE | FALSE |
| 988 | insH7 | P0CE54 | FALSE | FALSE | FALSE |
| 989 | folM | P0AFS3 | FALSE | FALSE | FALSE |
| 990 | tpx | P0A862 | FALSE | FALSE | FALSE |
| 991 | azoR | P41407 | FALSE | FALSE | FALSE |
| 992 | rihA | P41409 | FALSE | FALSE | FALSE |
| 993 | tdcB | P0AGF6 | FALSE | FALSE | FALSE |
| 994 | yqhH | P65298 | FALSE | MISSING | FALSE |
| 995 | rsfS | P0AAT6 | FALSE | FALSE | FALSE |
| 996 | ebgR | P06846 | FALSE | FALSE | FALSE |
| 997 | ycbZ | P75867 | FALSE | FALSE | FALSE |
| 998 | pncB | P18133 | FALSE | FALSE | FALSE |
| 999 | higA | P67701 | FALSE | FALSE | FALSE |
| 1000 | metK | P0A817 | FALSE | FALSE | TRUE |
| 1001 | sdaA | P16095 | FALSE | TRUE | FALSE |
| 1002 | hxpA | P77625 | TRUE | FALSE | FALSE |
| 1003 | yegT | P76417 | FALSE | FALSE | FALSE |
| 1004 | yjaG | P32680 | FALSE | FALSE | FALSE |
| 1005 | sdiA | P07026 | FALSE | FALSE | FALSE |
| 1006 | csdA | Q46925 | TRUE | FALSE | FALSE |
| 1007 | ribC | P0AFU8 | FALSE | TRUE | FALSE |
| 1008 | crp | P0ACJ8 | FALSE | FALSE | FALSE |

Supplementary Table 3: E. coli predicted metal-binding proteome *(continued)*

| # | Gene Name | UniProt ID | MoM Prediction | M3D Prediction | GASS Prediction |
|---|---|---|---|---|---|
| 1009 | modA | P37329 | FALSE | FALSE | FALSE |
| 1010 | punR | P0ACR2 | FALSE | FALSE | FALSE |
| 1011 | uxaC | P0A8G3 | TRUE | TRUE | FALSE |
| 1012 | ppnP | P0C037 | FALSE | FALSE | FALSE |
| 1013 | tfaE | P09153 | FALSE | FALSE | FALSE |
| 1014 | emrA | P27303 | FALSE | FALSE | FALSE |
| 1015 | glcB | P37330 | FALSE | FALSE | FALSE |
| 1016 | mutH | P06722 | FALSE | FALSE | FALSE |
| 1017 | moaA | P30745 | FALSE | FALSE | FALSE |
| 1018 | gmhA | P63224 | FALSE | FALSE | FALSE |
| 1019 | mglC | P23200 | FALSE | MISSING | FALSE |
| 1020 | menA | P32166 | FALSE | TRUE | FALSE |
| 1021 | ynjD | P76909 | FALSE | FALSE | FALSE |
| 1022 | fhuF | P39405 | TRUE | FALSE | FALSE |
| 1023 | fabF | P0AAI5 | FALSE | TRUE | FALSE |
| 1024 | ygfB | P0A8C4 | FALSE | FALSE | FALSE |
| 1025 | alkA | P04395 | FALSE | FALSE | FALSE |
| 1026 | yaiS | P71311 | TRUE | TRUE | TRUE |
| 1027 | sixA | P76502 | FALSE | FALSE | FALSE |
| 1028 | ribD | P25539 | TRUE | TRUE | TRUE |
| 1029 | sbcD | P0AG76 | TRUE | TRUE | FALSE |
| 1030 | yjjP | P0ADD5 | FALSE | FALSE | FALSE |
| 1031 | yggM | P46142 | FALSE | FALSE | FALSE |
| 1032 | yagH | P77713 | FALSE | FALSE | TRUE |
| 1033 | yahI | P77624 | FALSE | FALSE | FALSE |
| 1034 | cmk | P0A6I0 | FALSE | FALSE | FALSE |
| 1035 | metQ | P28635 | FALSE | FALSE | FALSE |
| 1036 | ybeM | P0DP64 | FALSE | FALSE | FALSE |
| 1037 | pxpC | P75745 | FALSE | FALSE | FALSE |
| 1038 | dinQ | A5A624 | FALSE | MISSING | FALSE |
| 1039 | fimE | P0ADH7 | FALSE | FALSE | FALSE |
| 1040 | yafO | Q47157 | FALSE | TRUE | FALSE |
| 1041 | tsgA | P60778 | FALSE | FALSE | FALSE |
| 1042 | exoX | P0AEK0 | FALSE | FALSE | FALSE |
| 1043 | yddH | P76121 | FALSE | FALSE | FALSE |
| 1044 | ydhS | P77148 | FALSE | FALSE | FALSE |
| 1045 | fimC | P31697 | FALSE | MISSING | FALSE |
| 1046 | btuB | P06129 | FALSE | FALSE | TRUE |
| 1047 | ykgG | P77433 | FALSE | MISSING | FALSE |
| 1048 | lspA | P00804 | FALSE | FALSE | FALSE |
| 1049 | matP | P0A8N0 | FALSE | FALSE | FALSE |
| 1050 | ulaE | P39305 | FALSE | TRUE | FALSE |
| 1051 | tktB | P33570 | TRUE | FALSE | FALSE |
| 1052 | hybD | P37182 | FALSE | TRUE | FALSE |
| 1053 | mutS | P23909 | FALSE | FALSE | FALSE |
| 1054 | pepT | P29745 | FALSE | TRUE | FALSE |
| 1055 | epd | P0A9B6 | TRUE | TRUE | FALSE |
| 1056 | phnP | P16692 | TRUE | TRUE | FALSE |
| 1057 | codA | P25524 | TRUE | TRUE | TRUE |
| 1058 | sotB | P31122 | FALSE | FALSE | FALSE |
| 1059 | pcm | P0A7A5 | FALSE | FALSE | FALSE |
| 1060 | citX | P0A6G5 | TRUE | TRUE | FALSE |
| 1061 | citC | P77390 | FALSE | TRUE | FALSE |
| 1062 | glsA1 | P77454 | FALSE | FALSE | FALSE |
| 1063 | opgE | P75785 | FALSE | TRUE | FALSE |
| 1064 | can | P61517 | TRUE | TRUE | TRUE |

Supplementary Table 3: E. coli predicted metal-binding proteome *(continued)*

| # | Gene Name | UniProt ID | MoM Prediction | M3D Prediction | GASS Prediction |
|---|---|---|---|---|---|
| 1065 | tauA | Q47537 | FALSE | FALSE | FALSE |
| 1066 | slyA | P0A8W2 | FALSE | FALSE | FALSE |
| 1067 | pepQ | P21165 | FALSE | TRUE | FALSE |
| 1068 | hisP | P07109 | FALSE | FALSE | FALSE |
| 1069 | pdxH | P0AFI7 | FALSE | FALSE | FALSE |
| 1070 | cra | P0ACP1 | FALSE | FALSE | FALSE |
| 1071 | ygfZ | P0ADE8 | FALSE | FALSE | FALSE |
| 1072 | hcxA | P45579 | FALSE | TRUE | TRUE |
| 1073 | yneJ | P77309 | FALSE | FALSE | FALSE |
| 1074 | nusB | P0A780 | FALSE | FALSE | FALSE |
| 1075 | ravA | P31473 | FALSE | FALSE | FALSE |
| 1076 | ydjH | P77493 | FALSE | FALSE | FALSE |
| 1077 | yehS | P33355 | FALSE | FALSE | FALSE |
| 1078 | yecE | P37348 | FALSE | FALSE | FALSE |
| 1079 | dcuA | P0ABN5 | FALSE | FALSE | FALSE |
| 1080 | nrdG | P0A9N8 | FALSE | FALSE | TRUE |
| 1081 | ssuD | P80645 | FALSE | TRUE | FALSE |
| 1082 | glyS | P00961 | FALSE | FALSE | TRUE |
| 1083 | ubiF | P75728 | FALSE | FALSE | FALSE |
| 1084 | yjjV | P39408 | TRUE | TRUE | FALSE |
| 1085 | rutF | P75893 | FALSE | FALSE | TRUE |
| 1086 | sgcQ | P39364 | FALSE | FALSE | FALSE |
| 1087 | paaJ | P0C7L2 | FALSE | TRUE | FALSE |
| 1088 | epmB | P39280 | FALSE | FALSE | FALSE |
| 1089 | ypaA | V9HVX0 | FALSE | FALSE | FALSE |
| 1090 | ybhH | P0AAV8 | FALSE | FALSE | FALSE |
| 1091 | adiC | P60061 | FALSE | FALSE | FALSE |
| 1092 | ynjA | P76222 | TRUE | MISSING | FALSE |
| 1093 | rhaS | P09377 | TRUE | TRUE | TRUE |
| 1094 | lyx | P37677 | FALSE | FALSE | TRUE |
| 1095 | yeiQ | P33029 | TRUE | FALSE | FALSE |
| 1096 | tesB | P0AGG2 | FALSE | FALSE | FALSE |
| 1097 | yhbQ | P45472 | FALSE | MISSING | FALSE |
| 1098 | ulaF | P39306 | TRUE | TRUE | TRUE |
| 1099 | flhE | P76297 | FALSE | FALSE | TRUE |
| 1100 | rbsK | P0A9J6 | FALSE | FALSE | FALSE |
| 1101 | ucpA | P37440 | FALSE | FALSE | FALSE |
| 1102 | hemA | P0A6X1 | FALSE | FALSE | FALSE |
| 1103 | hisM | P0AEU3 | FALSE | FALSE | TRUE |
| 1104 | hyfD | P77416 | FALSE | FALSE | FALSE |
| 1105 | dmsA | P18775 | FALSE | FALSE | FALSE |
| 1106 | rpmA | P0A7L8 | FALSE | FALSE | FALSE |
| 1107 | ygiF | P30871 | FALSE | FALSE | FALSE |
| 1108 | arnT | P76473 | FALSE | FALSE | FALSE |
| 1109 | atpA | P0ABB0 | FALSE | TRUE | FALSE |
| 1110 | ylbG | P77688 | FALSE | FALSE | FALSE |
| 1111 | ycdU | P75910 | FALSE | FALSE | FALSE |
| 1112 | yqaB | P77475 | FALSE | FALSE | TRUE |
| 1113 | amtB | P69681 | FALSE | FALSE | FALSE |
| 1114 | btuR | P0A9H5 | FALSE | FALSE | FALSE |
| 1115 | yjaH | P32681 | FALSE | FALSE | FALSE |
| 1116 | efeO | P0AB24 | FALSE | FALSE | FALSE |
| 1117 | kbl | P0AB77 | FALSE | FALSE | FALSE |
| 1118 | srkA | P0C0K3 | FALSE | FALSE | FALSE |
| 1119 | zwf | P0AC53 | FALSE | FALSE | FALSE |
| 1120 | wecH | P37669 | FALSE | FALSE | FALSE |

Supplementary Table 3: E. coli predicted metal-binding proteome *(continued)*

| # | Gene Name | UniProt ID | MoM Prediction | M3D Prediction | GASS Prediction |
|---|---|---|---|---|---|
| 1121 | ykgR | C1P5Z8 | FALSE | MISSING | FALSE |
| 1122 | cysB | P0A9F3 | FALSE | FALSE | FALSE |
| 1123 | fepA | P05825 | FALSE | FALSE | FALSE |
| 1124 | cysA | P16676 | FALSE | FALSE | FALSE |
| 1125 | yihU | P0A9V8 | FALSE | FALSE | FALSE |
| 1126 | mobB | P32125 | FALSE | FALSE | FALSE |
| 1127 | yhbO | P45470 | FALSE | TRUE | FALSE |
| 1128 | argS | P11875 | FALSE | FALSE | FALSE |
| 1129 | ugd | P76373 | FALSE | FALSE | FALSE |
| 1130 | cysE | P0A9D4 | FALSE | FALSE | FALSE |
| 1131 | rplI | P0A7R1 | FALSE | FALSE | FALSE |
| 1132 | rnhA | P0A7Y4 | FALSE | FALSE | FALSE |
| 1133 | murD | P14900 | FALSE | FALSE | FALSE |
| 1134 | narG | P09152 | FALSE | FALSE | TRUE |
| 1135 | acrR | P0ACS9 | FALSE | FALSE | FALSE |
| 1136 | hybC | P0ACE0 | TRUE | TRUE | FALSE |
| 1137 | purD | P15640 | FALSE | FALSE | FALSE |
| 1138 | yhbW | P0ADV5 | FALSE | FALSE | FALSE |
| 1139 | ybdR | P77316 | TRUE | TRUE | TRUE |
| 1140 | arnA | P77398 | TRUE | FALSE | FALSE |
| 1141 | yhbP | P67762 | FALSE | FALSE | FALSE |
| 1142 | yhfT | P45546 | FALSE | FALSE | FALSE |
| 1143 | yahV | P0DPN0 | FALSE | MISSING | FALSE |
| 1144 | yedJ | P46144 | FALSE | TRUE | FALSE |
| 1145 | rclR | P77379 | FALSE | FALSE | FALSE |
| 1146 | glpX | P0A9C9 | FALSE | TRUE | FALSE |
| 1147 | yadC | P31058 | FALSE | FALSE | FALSE |
| 1148 | yfjY | P52140 | TRUE | TRUE | TRUE |
| 1149 | adhE | P0A9Q7 | TRUE | TRUE | FALSE |
| 1150 | ydiL | P76196 | FALSE | FALSE | FALSE |
| 1151 | rpmB | P0A7M2 | FALSE | FALSE | FALSE |
| 1152 | yoaI | P76239 | FALSE | MISSING | FALSE |
| 1153 | yiiM | P32157 | FALSE | FALSE | FALSE |
| 1154 | bamD | P0AC02 | FALSE | FALSE | FALSE |
| 1155 | gudD | P0AES2 | TRUE | FALSE | TRUE |
| 1156 | ynfH | P76173 | FALSE | FALSE | FALSE |
| 1157 | sbmA | P0AFY6 | FALSE | FALSE | FALSE |
| 1158 | rzpR | P77551 | FALSE | MISSING | FALSE |
| 1159 | yrdA | P0A9W9 | TRUE | TRUE | TRUE |
| 1160 | rnc | P0A7Y0 | FALSE | FALSE | FALSE |
| 1161 | yqjH | Q46871 | FALSE | FALSE | FALSE |
| 1162 | cbrA | P31456 | FALSE | FALSE | TRUE |
| 1163 | speG | P0A951 | FALSE | FALSE | FALSE |
| 1164 | allG | P77129 | FALSE | FALSE | FALSE |
| 1165 | ycjD | P45736 | FALSE | TRUE | FALSE |
| 1166 | dmsD | P69853 | FALSE | FALSE | FALSE |
| 1167 | yagE | P75682 | FALSE | FALSE | FALSE |
| 1168 | ybbO | P0AFP4 | FALSE | FALSE | FALSE |
| 1169 | ybcY | P77460 | FALSE | FALSE | FALSE |
| 1170 | talA | P0A867 | FALSE | FALSE | FALSE |
| 1171 | wzb | P0AAB2 | FALSE | FALSE | FALSE |
| 1172 | iaaA | P37595 | FALSE | FALSE | FALSE |
| 1173 | xdhD | Q46814 | TRUE | FALSE | FALSE |
| 1174 | lamB | P02943 | FALSE | TRUE | FALSE |
| 1175 | citT | P0AE74 | FALSE | FALSE | FALSE |
| 1176 | ygcO | Q46905 | FALSE | FALSE | FALSE |

Supplementary Table 3: E. coli predicted metal-binding proteome *(continued)*

| # | Gene Name | UniProt ID | MoM Prediction | M3D Prediction | GASS Prediction |
|---|---|---|---|---|---|
| 1177 | rutE | P75894 | FALSE | FALSE | FALSE |
| 1178 | ycjY | P76049 | FALSE | FALSE | FALSE |
| 1179 | rsmA | P06992 | FALSE | FALSE | FALSE |
| 1180 | yraH | P42913 | FALSE | FALSE | FALSE |
| 1181 | yrhB | P46857 | FALSE | FALSE | FALSE |
| 1182 | mreB | P0A9X4 | FALSE | FALSE | FALSE |
| 1183 | ispG | P62620 | FALSE | FALSE | TRUE |
| 1184 | glpF | P0AER0 | FALSE | FALSE | FALSE |
| 1185 | ybhF | P0A9U1 | FALSE | FALSE | FALSE |
| 1186 | folK | P26281 | FALSE | FALSE | FALSE |
| 1187 | eutH | P76552 | FALSE | FALSE | FALSE |
| 1188 | hisH | P60595 | FALSE | FALSE | FALSE |
| 1189 | oppD | P76027 | FALSE | FALSE | FALSE |
| 1190 | yafJ | Q47147 | FALSE | FALSE | TRUE |
| 1191 | ybhJ | P75764 | TRUE | FALSE | FALSE |
| 1192 | thiI | P77718 | FALSE | FALSE | FALSE |
| 1193 | argC | P11446 | FALSE | FALSE | FALSE |
| 1194 | tadA | P68398 | TRUE | TRUE | FALSE |
| 1195 | intE | P75969 | FALSE | FALSE | FALSE |
| 1196 | pstA | P07654 | FALSE | MISSING | FALSE |
| 1197 | hcaT | Q47142 | FALSE | FALSE | FALSE |
| 1198 | ompW | P0A915 | TRUE | FALSE | FALSE |
| 1199 | coaD | P0A6I6 | FALSE | FALSE | FALSE |
| 1200 | coaBC | P0ABQ0 | FALSE | TRUE | FALSE |
| 1201 | avtA | P09053 | FALSE | FALSE | FALSE |
| 1202 | nrdF | P37146 | FALSE | TRUE | FALSE |
| 1203 | alsC | P32720 | FALSE | MISSING | FALSE |
| 1204 | hisI | P06989 | FALSE | TRUE | TRUE |
| 1205 | nuoK | P0AFE4 | FALSE | FALSE | FALSE |
| 1206 | sutR | P77626 | FALSE | TRUE | FALSE |
| 1207 | gnsB | P77695 | FALSE | MISSING | FALSE |
| 1208 | appB | P26458 | FALSE | FALSE | FALSE |
| 1209 | thrS | P0A8M3 | TRUE | TRUE | FALSE |
| 1210 | garR | P0ABQ2 | FALSE | FALSE | TRUE |
| 1211 | skp | P0AEU7 | FALSE | MISSING | FALSE |
| 1212 | adhP | P39451 | TRUE | TRUE | TRUE |
| 1213 | malS | P25718 | FALSE | FALSE | FALSE |
| 1214 | lsrG | P64461 | FALSE | FALSE | TRUE |
| 1215 | metH | P13009 | TRUE | TRUE | TRUE |
| 1216 | yeeZ | P0AD12 | FALSE | FALSE | FALSE |
| 1217 | amiD | P75820 | TRUE | TRUE | FALSE |
| 1218 | yecF | P0AD07 | FALSE | MISSING | FALSE |
| 1219 | aceA | P0A9G6 | FALSE | TRUE | TRUE |
| 1220 | pphA | P55798 | TRUE | FALSE | FALSE |
| 1221 | ygdD | P0ADR2 | FALSE | FALSE | FALSE |
| 1222 | yejK | P33920 | FALSE | FALSE | TRUE |
| 1223 | nrdE | P39452 | FALSE | FALSE | FALSE |
| 1224 | yafP | Q47158 | FALSE | FALSE | FALSE |
| 1225 | ung | P12295 | FALSE | FALSE | FALSE |
| 1226 | ldrD | Q6BF25 | FALSE | FALSE | FALSE |
| 1227 | rutA | P75898 | FALSE | FALSE | FALSE |
| 1228 | cstA | P15078 | FALSE | TRUE | FALSE |
| 1229 | ybdK | P77213 | FALSE | TRUE | FALSE |
| 1230 | yagJ | P77169 | FALSE | TRUE | FALSE |
| 1231 | rfbC | P37745 | FALSE | FALSE | FALSE |
| 1232 | ompC | P06996 | FALSE | FALSE | FALSE |

Supplementary Table 3: E. coli predicted metal-binding proteome *(continued)*

| # | Gene Name | UniProt ID | MoM Prediction | M3D Prediction | GASS Prediction |
|---|---|---|---|---|---|
| 1233 | yfeO | P67729 | FALSE | TRUE | FALSE |
| 1234 | abgB | P76052 | FALSE | TRUE | TRUE |
| 1235 | phoP | P23836 | FALSE | FALSE | FALSE |
| 1236 | ydeQ | P77588 | FALSE | FALSE | FALSE |
| 1237 | cfa | P0A9H7 | TRUE | FALSE | FALSE |
| 1238 | fucA | P0AB87 | TRUE | TRUE | FALSE |
| 1239 | flgC | P0ABX2 | FALSE | MISSING | FALSE |
| 1240 | yieH | P31467 | FALSE | FALSE | FALSE |
| 1241 | fadB | P21177 | FALSE | FALSE | FALSE |
| 1242 | wecC | P27829 | FALSE | FALSE | FALSE |
| 1243 | ybaA | P0AAQ6 | FALSE | MISSING | FALSE |
| 1244 | eptB | P37661 | FALSE | TRUE | FALSE |
| 1245 | yncE | P76116 | FALSE | FALSE | FALSE |
| 1246 | yeaE | P76234 | FALSE | FALSE | FALSE |
| 1247 | glnK | P0AC55 | FALSE | FALSE | FALSE |
| 1248 | yehY | P33361 | FALSE | FALSE | FALSE |
| 1249 | yhdZ | P45769 | FALSE | FALSE | FALSE |
| 1250 | hyfF | P77437 | FALSE | FALSE | FALSE |
| 1251 | allF | Q47208 | FALSE | FALSE | FALSE |
| 1252 | dolP | P64596 | FALSE | MISSING | FALSE |
| 1253 | csgG | P0AEA2 | FALSE | FALSE | FALSE |
| 1254 | lpxB | P10441 | FALSE | FALSE | FALSE |
| 1255 | hchA | P31658 | FALSE | TRUE | FALSE |
| 1256 | dcp | P24171 | TRUE | TRUE | FALSE |
| 1257 | sbcB | P04995 | FALSE | FALSE | TRUE |
| 1258 | fimB | P0ADH5 | FALSE | FALSE | FALSE |
| 1259 | psuG | P33025 | FALSE | FALSE | FALSE |
| 1260 | yagK | P77657 | TRUE | TRUE | TRUE |
| 1261 | nadA | P11458 | FALSE | FALSE | FALSE |
| 1262 | nagC | P0AF20 | TRUE | TRUE | FALSE |
| 1263 | bioH | P13001 | FALSE | TRUE | FALSE |
| 1264 | nudB | P0AFC0 | FALSE | FALSE | FALSE |
| 1265 | sbmC | P33012 | FALSE | FALSE | FALSE |
| 1266 | tmcA | P76562 | FALSE | FALSE | FALSE |
| 1267 | treR | P36673 | FALSE | FALSE | FALSE |
| 1268 | yphB | P76584 | TRUE | TRUE | TRUE |
| 1269 | cynR | P27111 | FALSE | FALSE | FALSE |
| 1270 | fecI | P23484 | FALSE | FALSE | FALSE |
| 1271 | ymfE | P75968 | FALSE | FALSE | FALSE |
| 1272 | bglX | P33363 | FALSE | FALSE | FALSE |
| 1273 | dtpA | P77304 | FALSE | FALSE | FALSE |
| 1274 | astD | P76217 | FALSE | FALSE | FALSE |
| 1275 | pth | P0A7D1 | FALSE | FALSE | FALSE |
| 1276 | yihV | P32143 | FALSE | FALSE | FALSE |
| 1277 | ykiD | P0DPN2 | FALSE | MISSING | FALSE |
| 1278 | priB | P07013 | FALSE | FALSE | FALSE |
| 1279 | fklB | P0A9L3 | FALSE | MISSING | FALSE |
| 1280 | pdxY | P77150 | FALSE | FALSE | FALSE |
| 1281 | uspF | P37903 | FALSE | FALSE | FALSE |
| 1282 | uxaB | P0A6L7 | FALSE | FALSE | FALSE |
| 1283 | fbp | P0A993 | FALSE | FALSE | FALSE |
| 1284 | yibH | P0AFV0 | FALSE | TRUE | TRUE |
| 1285 | menI | P77781 | FALSE | FALSE | FALSE |
| 1286 | lgoD | P39400 | TRUE | TRUE | TRUE |
| 1287 | diaA | P66817 | FALSE | FALSE | FALSE |
| 1288 | yddG | P46136 | FALSE | FALSE | FALSE |

Supplementary Table 3: E. coli predicted metal-binding proteome *(continued)*

| # | Gene Name | UniProt ID | MoM Prediction | M3D Prediction | GASS Prediction |
|---|---|---|---|---|---|
| 1289 | holC | P28905 | FALSE | MISSING | FALSE |
| 1290 | lgt | P60955 | FALSE | FALSE | FALSE |
| 1291 | yihL | P0ACM9 | FALSE | FALSE | FALSE |
| 1292 | adiA | P28629 | FALSE | FALSE | TRUE |
| 1293 | purR | P0ACP7 | FALSE | FALSE | FALSE |
| 1294 | mrcA | P02918 | FALSE | FALSE | FALSE |
| 1295 | insQ | P76102 | TRUE | TRUE | TRUE |
| 1296 | ytiA | P0DN74 | FALSE | FALSE | FALSE |
| 1297 | mlc | P50456 | TRUE | TRUE | FALSE |
| 1298 | nikD | P33593 | FALSE | FALSE | TRUE |
| 1299 | ydfZ | P64463 | FALSE | FALSE | FALSE |
| 1300 | ybcI | P45570 | FALSE | TRUE | FALSE |
| 1301 | moaD | P30748 | FALSE | FALSE | FALSE |
| 1302 | paoC | P77489 | FALSE | FALSE | FALSE |
| 1303 | aceK | P11071 | TRUE | FALSE | FALSE |
| 1304 | lysU | P0A8N5 | FALSE | FALSE | FALSE |
| 1305 | rlmJ | P37634 | FALSE | FALSE | FALSE |
| 1306 | hcaB | P0CI31 | FALSE | FALSE | FALSE |
| 1307 | ygeA | P03813 | FALSE | FALSE | FALSE |
| 1308 | allB | P77671 | TRUE | TRUE | TRUE |
| 1309 | caiC | P31552 | FALSE | FALSE | TRUE |
| 1310 | ihfA | P0A6X7 | FALSE | MISSING | FALSE |
| 1311 | pgaB | P75906 | TRUE | TRUE | FALSE |
| 1312 | rplB | P60422 | FALSE | FALSE | FALSE |
| 1313 | ddlB | P07862 | FALSE | FALSE | FALSE |
| 1314 | rraA | P0A8R0 | FALSE | FALSE | FALSE |
| 1315 | frwB | P69816 | FALSE | TRUE | FALSE |
| 1316 | corA | P0ABI4 | FALSE | FALSE | FALSE |
| 1317 | yjfF | P37772 | FALSE | MISSING | FALSE |
| 1318 | uhpA | P0AGA6 | FALSE | TRUE | FALSE |
| 1319 | ydcJ | P76097 | FALSE | TRUE | FALSE |
| 1320 | garD | P39829 | FALSE | TRUE | FALSE |
| 1321 | yfiM | P46126 | FALSE | FALSE | FALSE |
| 1322 | yebY | P64506 | FALSE | MISSING | FALSE |
| 1323 | yfgM | P76576 | FALSE | FALSE | FALSE |
| 1324 | yihM | P32134 | FALSE | TRUE | FALSE |
| 1325 | sfmH | P75715 | FALSE | FALSE | TRUE |
| 1326 | yaeR | P52096 | FALSE | TRUE | FALSE |
| 1327 | yjcZ | P39267 | FALSE | FALSE | TRUE |
| 1328 | yhaM | P42626 | FALSE | FALSE | FALSE |
| 1329 | mcbR | P76114 | FALSE | FALSE | FALSE |
| 1330 | ampC | P00811 | FALSE | FALSE | FALSE |
| 1331 | rhaR | P09378 | TRUE | TRUE | TRUE |
| 1332 | ftsE | P0A9R7 | FALSE | FALSE | FALSE |
| 1333 | waaZ | P27241 | FALSE | FALSE | FALSE |
| 1334 | rimJ | P0A948 | FALSE | FALSE | TRUE |
| 1335 | queA | P0A7F9 | FALSE | FALSE | TRUE |
| 1336 | helD | P15038 | FALSE | FALSE | FALSE |
| 1337 | wecB | P27828 | FALSE | FALSE | FALSE |
| 1338 | garL | P23522 | FALSE | FALSE | FALSE |
| 1339 | ybeR | P77627 | FALSE | FALSE | FALSE |
| 1340 | hisA | P10371 | FALSE | FALSE | FALSE |
| 1341 | ruvA | P0A809 | FALSE | FALSE | FALSE |
| 1342 | yfcG | P77526 | FALSE | FALSE | FALSE |
| 1343 | yfdG | P77682 | FALSE | FALSE | FALSE |
| 1344 | dgoA | Q6BF16 | FALSE | FALSE | FALSE |

Supplementary Table 3: E. coli predicted metal-binding proteome *(continued)*

| # | Gene Name | UniProt ID | MoM Prediction | M3D Prediction | GASS Prediction |
|---|---|---|---|---|---|
| 1345 | ppdA | P33554 | TRUE | FALSE | FALSE |
| 1346 | ybjN | P0AAY6 | TRUE | TRUE | FALSE |
| 1347 | nudK | P37128 | FALSE | FALSE | FALSE |
| 1348 | yhcO | P64616 | FALSE | FALSE | FALSE |
| 1349 | ecpB | P77188 | FALSE | FALSE | TRUE |
| 1350 | glnP | P0AEQ6 | FALSE | FALSE | FALSE |
| 1351 | ldcC | P52095 | FALSE | FALSE | FALSE |
| 1352 | yceI | P0A8X2 | FALSE | FALSE | FALSE |
| 1353 | fabD | P0AAI9 | FALSE | FALSE | FALSE |
| 1354 | yojI | P33941 | FALSE | FALSE | TRUE |
| 1355 | yihN | P32135 | FALSE | FALSE | FALSE |
| 1356 | frwC | P32672 | FALSE | FALSE | FALSE |
| 1357 | rpsD | P0A7V8 | FALSE | FALSE | FALSE |
| 1358 | intA | P32053 | FALSE | FALSE | FALSE |
| 1359 | rplS | P0A7K6 | FALSE | FALSE | FALSE |
| 1360 | carA | P0A6F1 | TRUE | TRUE | TRUE |
| 1361 | ulaB | P69822 | FALSE | FALSE | FALSE |
| 1362 | rfbA | P37744 | FALSE | FALSE | FALSE |
| 1363 | ade | P31441 | TRUE | TRUE | TRUE |
| 1364 | ygdG | P38506 | FALSE | FALSE | FALSE |
| 1365 | ttdB | P0AC35 | FALSE | TRUE | TRUE |
| 1366 | recC | P07648 | FALSE | TRUE | FALSE |
| 1367 | nrdR | P0A8D0 | TRUE | TRUE | TRUE |
| 1368 | yidK | P31448 | FALSE | FALSE | FALSE |
| 1369 | lacI | P03023 | FALSE | FALSE | FALSE |
| 1370 | rpiR | P0ACS7 | FALSE | FALSE | FALSE |
| 1371 | scpC | P52043 | FALSE | FALSE | FALSE |
| 1372 | fadM | P77712 | FALSE | FALSE | TRUE |
| 1373 | ulaC | P69820 | FALSE | FALSE | FALSE |
| 1374 | asnA | P00963 | FALSE | FALSE | TRUE |
| 1375 | thyA | P0A884 | FALSE | FALSE | FALSE |
| 1376 | rpoZ | P0A800 | FALSE | MISSING | FALSE |
| 1377 | yjdC | P0ACU7 | FALSE | TRUE | FALSE |
| 1378 | chpB | P33647 | FALSE | FALSE | FALSE |
| 1379 | nirC | P0AC26 | FALSE | FALSE | TRUE |
| 1380 | roxA | P27431 | TRUE | TRUE | FALSE |
| 1381 | folX | P0AC19 | FALSE | FALSE | FALSE |
| 1382 | ypfM | A5A621 | FALSE | MISSING | FALSE |
| 1383 | cydA | P0ABJ9 | FALSE | FALSE | FALSE |
| 1384 | pyrC | P05020 | TRUE | TRUE | FALSE |
| 1385 | cdh | P06282 | FALSE | FALSE | FALSE |
| 1386 | sucD | P0AGE9 | FALSE | FALSE | FALSE |
| 1387 | rpsQ | P0AG63 | FALSE | FALSE | FALSE |
| 1388 | slmA | P0C093 | FALSE | FALSE | FALSE |
| 1389 | nemR | P67430 | FALSE | FALSE | FALSE |
| 1390 | maa | P77791 | FALSE | FALSE | FALSE |
| 1391 | dld | P06149 | TRUE | TRUE | FALSE |
| 1392 | glk | P0A6V8 | FALSE | FALSE | FALSE |
| 1393 | rplX | P60624 | FALSE | FALSE | FALSE |
| 1394 | yeeT | P64521 | FALSE | FALSE | FALSE |
| 1395 | hyfC | P77858 | FALSE | FALSE | FALSE |
| 1396 | gpt | P0A9M5 | FALSE | FALSE | FALSE |
| 1397 | caiA | P60584 | FALSE | FALSE | FALSE |
| 1398 | wecD | P27832 | FALSE | FALSE | TRUE |
| 1399 | insF2 | P0CF80 | FALSE | FALSE | FALSE |
| 1400 | aroG | P0AB91 | TRUE | TRUE | FALSE |

Supplementary Table 3: E. coli predicted metal-binding proteome *(continued)*

| # | Gene Name | UniProt ID | MoM Prediction | M3D Prediction | GASS Prediction |
|---|---|---|---|---|---|
| 1401 | ykiA | P75704 | FALSE | FALSE | FALSE |
| 1402 | fabA | P0A6Q3 | FALSE | FALSE | FALSE |
| 1403 | rspB | P38105 | TRUE | TRUE | FALSE |
| 1404 | yhbX | P42640 | FALSE | TRUE | FALSE |
| 1405 | yajG | P0ADA5 | FALSE | FALSE | FALSE |
| 1406 | hslV | P0A7B8 | FALSE | FALSE | FALSE |
| 1407 | mdfA | P0AEY8 | FALSE | FALSE | FALSE |
| 1408 | yciA | P0A8Z0 | FALSE | FALSE | FALSE |
| 1409 | tdcE | P42632 | FALSE | TRUE | TRUE |
| 1410 | xdhB | Q46800 | FALSE | FALSE | FALSE |
| 1411 | frdC | P0A8Q0 | FALSE | MISSING | FALSE |
| 1412 | nanQ | P45424 | FALSE | TRUE | FALSE |
| 1413 | cpdA | P0AEW4 | TRUE | TRUE | TRUE |
| 1414 | ggt | P18956 | FALSE | FALSE | FALSE |
| 1415 | marR | P27245 | FALSE | FALSE | FALSE |
| 1416 | yegU | P76418 | FALSE | FALSE | TRUE |
| 1417 | tmaR | P0A8M6 | FALSE | MISSING | FALSE |
| 1418 | casA | Q46901 | TRUE | TRUE | TRUE |
| 1419 | ppa | P0A7A9 | FALSE | FALSE | FALSE |
| 1420 | ibsA | C1P607 | FALSE | MISSING | FALSE |
| 1421 | yhdE | P25536 | FALSE | FALSE | FALSE |
| 1422 | yjaZ | P27375 | FALSE | FALSE | FALSE |
| 1423 | aspC | P00509 | FALSE | FALSE | FALSE |
| 1424 | ogt | P0AFH0 | FALSE | FALSE | FALSE |
| 1425 | fadJ | P77399 | FALSE | FALSE | TRUE |
| 1426 | lldD | P33232 | FALSE | FALSE | FALSE |
| 1427 | pykA | P21599 | FALSE | FALSE | FALSE |
| 1428 | yfdL | P76508 | FALSE | MISSING | FALSE |
| 1429 | ubiH | P25534 | TRUE | TRUE | FALSE |
| 1430 | purT | P33221 | FALSE | FALSE | FALSE |
| 1431 | ycdY | P75915 | FALSE | FALSE | FALSE |
| 1432 | rsuA | P0AA43 | FALSE | FALSE | FALSE |
| 1433 | slyX | P0A8R4 | FALSE | FALSE | FALSE |
| 1434 | rnt | P30014 | FALSE | FALSE | TRUE |
| 1435 | rhmA | P76469 | FALSE | FALSE | FALSE |
| 1436 | ymfS | P09154 | FALSE | FALSE | FALSE |
| 1437 | yahB | P77700 | FALSE | TRUE | FALSE |
| 1438 | pspE | P23857 | FALSE | FALSE | FALSE |
| 1439 | glpD | P13035 | FALSE | FALSE | FALSE |
| 1440 | yeaW | P0ABR7 | TRUE | TRUE | TRUE |
| 1441 | clcB | P76175 | FALSE | FALSE | FALSE |
| 1442 | fadL | P10384 | FALSE | FALSE | FALSE |
| 1443 | ycaQ | P75843 | FALSE | FALSE | FALSE |
| 1444 | tyrP | P0AAD4 | FALSE | FALSE | TRUE |
| 1445 | ycaC | P21367 | FALSE | FALSE | FALSE |
| 1446 | hybF | P0A703 | TRUE | TRUE | FALSE |
| 1447 | iprA | P0AAP5 | FALSE | FALSE | FALSE |
| 1448 | purE | P0AG18 | FALSE | FALSE | FALSE |
| 1449 | fliQ | P0AC07 | FALSE | MISSING | FALSE |
| 1450 | cysI | P17846 | FALSE | FALSE | FALSE |
| 1451 | ssuB | P0AAI1 | FALSE | FALSE | FALSE |
| 1452 | zntR | P0ACS5 | FALSE | FALSE | FALSE |
| 1453 | purU | P37051 | FALSE | FALSE | FALSE |
| 1454 | recO | P0A7H3 | FALSE | FALSE | FALSE |
| 1455 | kdgR | P76268 | FALSE | FALSE | FALSE |
| 1456 | gor | P06715 | FALSE | TRUE | FALSE |

Supplementary Table 3: E. coli predicted metal-binding proteome *(continued)*

| # | Gene Name | UniProt ID | MoM Prediction | M3D Prediction | GASS Prediction |
|---|---|---|---|---|---|
| 1457 | phnK | P16678 | FALSE | FALSE | FALSE |
| 1458 | yfjZ | P52141 | FALSE | FALSE | FALSE |
| 1459 | sucA | P0AFG3 | FALSE | TRUE | TRUE |
| 1460 | mobA | P32173 | FALSE | FALSE | TRUE |
| 1461 | yqeC | Q46809 | FALSE | FALSE | FALSE |
| 1462 | fabZ | P0A6Q6 | FALSE | FALSE | FALSE |
| 1463 | ydeU | P77286 | TRUE | FALSE | FALSE |
| 1464 | tusA | P0A890 | FALSE | FALSE | FALSE |
| 1465 | fbpC | P37009 | FALSE | FALSE | FALSE |
| 1466 | paaH | P76083 | FALSE | FALSE | TRUE |
| 1467 | acuI | P26646 | FALSE | FALSE | FALSE |
| 1468 | ydfO | P76156 | FALSE | FALSE | FALSE |
| 1469 | ghrA | P75913 | FALSE | FALSE | FALSE |
| 1470 | yifN | P56259 | FALSE | FALSE | FALSE |
| 1471 | purH | P15639 | FALSE | FALSE | FALSE |
| 1472 | ybaK | P0AAR3 | FALSE | FALSE | FALSE |
| 1473 | ypdK | C1P610 | FALSE | MISSING | FALSE |
| 1474 | yajO | P77735 | FALSE | FALSE | TRUE |
| 1475 | rpsI | P0A7X3 | FALSE | MISSING | FALSE |
| 1476 | hupB | P0ACF4 | FALSE | MISSING | FALSE |
| 1477 | holB | P28631 | TRUE | TRUE | FALSE |
| 1478 | nfsA | P17117 | FALSE | FALSE | FALSE |
| 1479 | yjjG | P0A8Y1 | FALSE | FALSE | FALSE |
| 1480 | dxs | P77488 | FALSE | FALSE | TRUE |
| 1481 | betB | P17445 | FALSE | FALSE | FALSE |
| 1482 | prlC | P27298 | TRUE | TRUE | FALSE |
| 1483 | menD | P17109 | FALSE | FALSE | TRUE |
| 1484 | sthA | P27306 | FALSE | FALSE | FALSE |
| 1485 | ytfJ | P39187 | FALSE | MISSING | FALSE |
| 1486 | lolB | P61320 | FALSE | FALSE | FALSE |
| 1487 | ydeE | P31126 | FALSE | MISSING | FALSE |
| 1488 | pgpC | P0AD42 | FALSE | FALSE | FALSE |
| 1489 | ubiD | P0AAB4 | FALSE | FALSE | TRUE |
| 1490 | caiE | P39206 | FALSE | FALSE | FALSE |
| 1491 | xapB | P45562 | FALSE | FALSE | FALSE |
| 1492 | proXp-y | P64483 | FALSE | FALSE | FALSE |
| 1493 | yqeF | Q46939 | FALSE | FALSE | FALSE |
| 1494 | rihC | P22564 | FALSE | FALSE | FALSE |
| 1495 | tdh | P07913 | TRUE | TRUE | TRUE |
| 1496 | argE | P23908 | FALSE | TRUE | FALSE |
| 1497 | wcaM | P71244 | FALSE | FALSE | FALSE |
| 1498 | norW | P37596 | FALSE | FALSE | FALSE |
| 1499 | cueO | P36649 | TRUE | TRUE | TRUE |
| 1500 | folC | P08192 | TRUE | FALSE | FALSE |
| 1501 | rplU | P0AG48 | FALSE | FALSE | FALSE |
| 1502 | eda | P0A955 | FALSE | FALSE | FALSE |
| 1503 | yqgA | Q46831 | FALSE | FALSE | FALSE |
| 1504 | yohK | P0AD19 | FALSE | FALSE | FALSE |
| 1505 | moaC | P0A738 | FALSE | FALSE | FALSE |
| 1506 | nudF | Q93K97 | FALSE | FALSE | FALSE |
| 1507 | greB | P30128 | FALSE | MISSING | FALSE |
| 1508 | rnd | P09155 | FALSE | FALSE | FALSE |
| 1509 | rplO | P02413 | FALSE | FALSE | FALSE |
| 1510 | yegV | P76419 | FALSE | FALSE | FALSE |
| 1511 | yehL | P33348 | FALSE | FALSE | TRUE |
| 1512 | ydcS | P76108 | FALSE | TRUE | FALSE |

Supplementary Table 3: E. coli predicted metal-binding proteome *(continued)*

| # | Gene Name | UniProt ID | MoM Prediction | M3D Prediction | GASS Prediction |
|---|---|---|---|---|---|
| 1513 | glnE | P30870 | FALSE | TRUE | FALSE |
| 1514 | pfkA | P0A796 | FALSE | FALSE | FALSE |
| 1515 | mhpC | P77044 | FALSE | FALSE | FALSE |
| 1516 | rsmF | P76273 | FALSE | FALSE | TRUE |
| 1517 | fbaA | P0AB71 | TRUE | FALSE | FALSE |
| 1518 | ycgM | P76004 | FALSE | FALSE | FALSE |
| 1519 | flgA | P75933 | FALSE | FALSE | FALSE |
| 1520 | insH11 | P0CE58 | FALSE | FALSE | FALSE |
| 1521 | ybcM | P77634 | FALSE | FALSE | TRUE |
| 1522 | argT | P09551 | FALSE | FALSE | FALSE |
| 1523 | rplN | P0ADY3 | FALSE | MISSING | FALSE |
| 1524 | wrbA | P0A8G6 | FALSE | FALSE | FALSE |
| 1525 | rpsO | P0ADZ4 | FALSE | FALSE | FALSE |
| 1526 | ushA | P07024 | TRUE | TRUE | TRUE |
| 1527 | guaA | P04079 | FALSE | FALSE | FALSE |
| 1528 | sdhB | P07014 | FALSE | FALSE | TRUE |
| 1529 | gmm | P32056 | FALSE | FALSE | FALSE |
| 1530 | gldA | P0A9S5 | FALSE | TRUE | TRUE |
| 1531 | uvrY | P0AED5 | FALSE | TRUE | FALSE |
| 1532 | ycdZ | P75916 | FALSE | FALSE | FALSE |
| 1533 | katG | P13029 | FALSE | FALSE | FALSE |
| 1534 | rfbB | P37759 | FALSE | FALSE | FALSE |
| 1535 | hypF | P30131 | TRUE | TRUE | TRUE |
| 1536 | gatA | P69828 | FALSE | FALSE | FALSE |
| 1537 | yahC | P77219 | FALSE | MISSING | FALSE |
| 1538 | emtA | P0C960 | FALSE | FALSE | FALSE |
| 1539 | ybcV | P77598 | FALSE | FALSE | FALSE |
| 1540 | torY | P52005 | TRUE | TRUE | TRUE |
| 1541 | rnm | P77766 | TRUE | TRUE | TRUE |
| 1542 | purC | P0A7D7 | FALSE | FALSE | FALSE |
| 1543 | yddA | P31826 | FALSE | TRUE | FALSE |
| 1544 | hypD | P24192 | FALSE | FALSE | FALSE |
| 1545 | tdcG | P42630 | FALSE | TRUE | FALSE |
| 1546 | yjjB | P0ADD2 | FALSE | MISSING | FALSE |
| 1547 | xylB | P09099 | FALSE | FALSE | FALSE |
| 1548 | cirA | P17315 | TRUE | TRUE | TRUE |
| 1549 | dhaL | P76014 | FALSE | FALSE | FALSE |
| 1550 | ydiR | P77378 | FALSE | FALSE | FALSE |
| 1551 | pheA | P0A9J8 | FALSE | FALSE | FALSE |
| 1552 | patA | P42588 | FALSE | FALSE | FALSE |
| 1553 | ynbA | P76090 | FALSE | FALSE | FALSE |
| 1554 | fruK | P0AEW9 | FALSE | FALSE | FALSE |
| 1555 | clpS | P0A8Q6 | FALSE | MISSING | FALSE |
| 1556 | dkgB | P30863 | FALSE | FALSE | FALSE |
| 1557 | yfcF | P77544 | FALSE | FALSE | FALSE |
| 1558 | psd | P0A8K1 | FALSE | FALSE | FALSE |
| 1559 | bamE | P0A937 | FALSE | MISSING | FALSE |
| 1560 | agaI | P42912 | FALSE | FALSE | FALSE |
| 1561 | mdoG | P33136 | FALSE | FALSE | FALSE |
| 1562 | hycD | P16430 | FALSE | FALSE | FALSE |
| 1563 | ribA | P0A7I7 | TRUE | TRUE | TRUE |
| 1564 | trmB | P0A8I5 | FALSE | FALSE | FALSE |
| 1565 | aaeX | P46478 | FALSE | MISSING | FALSE |
| 1566 | yebZ | P76278 | FALSE | FALSE | FALSE |
| 1567 | frmA | P25437 | TRUE | TRUE | TRUE |
| 1568 | rtcB | P46850 | TRUE | TRUE | TRUE |

Supplementary Table 3: E. coli predicted metal-binding proteome *(continued)*

| # | Gene Name | UniProt ID | MoM Prediction | M3D Prediction | GASS Prediction |
|---|---|---|---|---|---|
| 1569 | yafQ | Q47149 | TRUE | FALSE | FALSE |
| 1570 | fiu | P75780 | FALSE | FALSE | FALSE |
| 1571 | allS | P0ACR0 | FALSE | MISSING | FALSE |
| 1572 | tehA | P25396 | FALSE | FALSE | FALSE |
| 1573 | guaC | P60560 | FALSE | FALSE | FALSE |
| 1574 | yecA | P0AD05 | TRUE | TRUE | FALSE |
| 1575 | ydbK | P52647 | FALSE | FALSE | FALSE |
| 1576 | tfaR | P77163 | FALSE | TRUE | FALSE |
| 1577 | glpR | P0ACL0 | FALSE | FALSE | FALSE |
| 1578 | rfbD | P37760 | FALSE | FALSE | FALSE |
| 1579 | yfaQ | P76463 | TRUE | TRUE | FALSE |
| 1580 | upp | P0A8F0 | FALSE | FALSE | FALSE |
| 1581 | lapB | P0AB58 | TRUE | TRUE | FALSE |
| 1582 | coaA | P0A6I3 | TRUE | FALSE | FALSE |
| 1583 | murI | P22634 | FALSE | TRUE | FALSE |
| 1584 | yihR | P32139 | FALSE | FALSE | TRUE |
| 1585 | hdeD | P0AET5 | FALSE | MISSING | FALSE |
| 1586 | yicH | P31433 | FALSE | FALSE | FALSE |
| 1587 | poxB | P07003 | FALSE | FALSE | TRUE |
| 1588 | yieK | P31470 | FALSE | FALSE | FALSE |
| 1589 | araB | P08204 | TRUE | TRUE | FALSE |
| 1590 | yciT | P76034 | FALSE | FALSE | FALSE |
| 1591 | otnK | Q46889 | FALSE | FALSE | FALSE |
| 1592 | moeB | P12282 | TRUE | TRUE | FALSE |
| 1593 | aidB | P33224 | FALSE | FALSE | FALSE |
| 1594 | ygeX | P66899 | FALSE | FALSE | FALSE |
| 1595 | yhiM | P37630 | TRUE | FALSE | FALSE |
| 1596 | iadA | P39377 | TRUE | TRUE | FALSE |
| 1597 | rcsA | P0DMC9 | FALSE | FALSE | FALSE |
| 1598 | truC | P0AA41 | FALSE | TRUE | FALSE |
| 1599 | tufB | P0CE48 | FALSE | FALSE | FALSE |
| 1600 | yibL | P0ADK8 | FALSE | FALSE | FALSE |
| 1601 | wcaL | P71243 | FALSE | FALSE | FALSE |
| 1602 | yeaD | P39173 | FALSE | FALSE | TRUE |
| 1603 | nsrR | P0AF63 | FALSE | MISSING | FALSE |
| 1604 | pheT | P07395 | FALSE | FALSE | FALSE |
| 1605 | glpB | P13033 | FALSE | TRUE | FALSE |
| 1606 | fdnI | P0AEK7 | FALSE | TRUE | FALSE |
| 1607 | yhbY | P0AGK4 | FALSE | FALSE | FALSE |
| 1608 | yhcC | P0ADW6 | TRUE | FALSE | FALSE |
| 1609 | leuO | P10151 | FALSE | TRUE | FALSE |
| 1610 | ygfK | Q46811 | FALSE | TRUE | FALSE |
| 1611 | rsmH | P60390 | FALSE | FALSE | FALSE |
| 1612 | cof | P46891 | TRUE | TRUE | TRUE |
| 1613 | arcA | P0A9Q1 | FALSE | TRUE | FALSE |
| 1614 | yjhR | P39369 | FALSE | FALSE | FALSE |
| 1615 | ycgX | P75988 | FALSE | FALSE | FALSE |
| 1616 | yfcE | P67095 | TRUE | TRUE | FALSE |
| 1617 | betI | P17446 | FALSE | FALSE | FALSE |
| 1618 | deoD | P0ABP8 | FALSE | MISSING | FALSE |
| 1619 | yhjJ | P37648 | FALSE | FALSE | FALSE |
| 1620 | ribB | P0A7J0 | FALSE | FALSE | FALSE |
| 1621 | ybdL | P77806 | FALSE | FALSE | FALSE |
| 1622 | nrfG | P32712 | FALSE | FALSE | FALSE |
| 1623 | pyrD | P0A7E1 | FALSE | FALSE | FALSE |
| 1624 | ybjS | P75821 | FALSE | FALSE | TRUE |

Supplementary Table 3: E. coli predicted metal-binding proteome *(continued)*

| # | Gene Name | UniProt ID | MoM Prediction | M3D Prediction | GASS Prediction |
|---|---|---|---|---|---|
| 1625 | ydeN | P77318 | FALSE | FALSE | FALSE |
| 1626 | wbbI | P37749 | FALSE | FALSE | FALSE |
| 1627 | cbrC | P31469 | TRUE | TRUE | TRUE |
| 1628 | dgt | P15723 | FALSE | TRUE | TRUE |
| 1629 | clsA | P0A6H8 | FALSE | FALSE | FALSE |
| 1630 | speB | P60651 | TRUE | TRUE | FALSE |
| 1631 | atoE | P76460 | FALSE | FALSE | FALSE |
| 1632 | mcrA | P24200 | TRUE | TRUE | TRUE |
| 1633 | glpQ | P09394 | FALSE | TRUE | FALSE |
| 1634 | dacC | P08506 | FALSE | FALSE | FALSE |
| 1635 | ybcJ | P0AAS7 | FALSE | FALSE | FALSE |
| 1636 | rpsN | P0AG59 | FALSE | MISSING | FALSE |
| 1637 | ynhF | A5A618 | FALSE | MISSING | FALSE |
| 1638 | xylE | P0AGF4 | FALSE | FALSE | FALSE |
| 1639 | gluQ | P27305 | TRUE | TRUE | TRUE |
| 1640 | bdcR | P39334 | FALSE | FALSE | FALSE |
| 1641 | ycjT | P77154 | FALSE | FALSE | FALSE |
| 1642 | waaG | P25740 | FALSE | FALSE | FALSE |
| 1643 | trmA | P23003 | FALSE | FALSE | FALSE |
| 1644 | phnJ | P16688 | TRUE | TRUE | FALSE |
| 1645 | cysH | P17854 | FALSE | FALSE | FALSE |
| 1646 | recF | P0A7H0 | FALSE | FALSE | FALSE |
| 1647 | intS | P37326 | FALSE | FALSE | FALSE |
| 1648 | yicL | P31437 | FALSE | FALSE | FALSE |
| 1649 | dsrB | P0AEG8 | FALSE | FALSE | FALSE |
| 1650 | ypdB | P0AE39 | FALSE | FALSE | FALSE |
| 1651 | ybgC | P0A8Z3 | FALSE | FALSE | FALSE |
| 1652 | tcyN | P37774 | FALSE | FALSE | FALSE |
| 1653 | ybeY | P0A898 | TRUE | TRUE | FALSE |
| 1654 | ltnD | Q46888 | FALSE | FALSE | FALSE |
| 1655 | argP | P0A8S1 | FALSE | FALSE | FALSE |
| 1656 | yggL | P38521 | FALSE | FALSE | FALSE |
| 1657 | glxR | P77161 | FALSE | FALSE | FALSE |
| 1658 | lptB | P0A9V1 | FALSE | FALSE | FALSE |
| 1659 | astA | P0AE37 | FALSE | FALSE | FALSE |
| 1660 | yfiE | P33634 | FALSE | TRUE | TRUE |
| 1661 | ybhI | P75763 | FALSE | FALSE | FALSE |
| 1662 | yddB | P31827 | TRUE | FALSE | FALSE |
| 1663 | dapB | P04036 | FALSE | FALSE | FALSE |
| 1664 | eutN | P0AEJ8 | FALSE | FALSE | FALSE |
| 1665 | trpGD | P00904 | FALSE | FALSE | FALSE |
| 1666 | mepA | P0C0T5 | TRUE | TRUE | TRUE |
| 1667 | ydcF | P34209 | FALSE | FALSE | FALSE |
| 1668 | kdsA | P0A715 | FALSE | FALSE | FALSE |
| 1669 | pssA | P23830 | FALSE | FALSE | FALSE |
| 1670 | flgK | P33235 | FALSE | MISSING | FALSE |
| 1671 | waaC | P24173 | FALSE | TRUE | FALSE |
| 1672 | fdoH | P0AAJ5 | TRUE | FALSE | FALSE |
| 1673 | kdsB | P04951 | FALSE | FALSE | FALSE |
| 1674 | yfjI | P52124 | FALSE | FALSE | FALSE |
| 1675 | rbsB | P02925 | FALSE | FALSE | FALSE |
| 1676 | otnC | Q46890 | TRUE | TRUE | TRUE |
| 1677 | argA | P0A6C5 | FALSE | FALSE | FALSE |
| 1678 | rtcR | P38035 | FALSE | FALSE | TRUE |
| 1679 | rpmF | P0A7N4 | FALSE | FALSE | FALSE |
| 1680 | frvA | P32155 | FALSE | FALSE | FALSE |

Supplementary Table 3: E. coli predicted metal-binding proteome *(continued)*

| # | Gene Name | UniProt ID | MoM Prediction | M3D Prediction | GASS Prediction |
|---|---|---|---|---|---|
| 1681 | glpC | P0A996 | TRUE | TRUE | TRUE |
| 1682 | hyi | P30147 | FALSE | FALSE | FALSE |
| 1683 | rimL | P13857 | FALSE | FALSE | FALSE |
| 1684 | nadR | P27278 | FALSE | FALSE | FALSE |
| 1685 | ygcQ | Q46907 | FALSE | FALSE | FALSE |
| 1686 | yccU | P75874 | FALSE | FALSE | FALSE |
| 1687 | ysaA | P56256 | FALSE | FALSE | FALSE |
| 1688 | moeA | P12281 | FALSE | FALSE | FALSE |
| 1689 | ycfH | P0AFQ7 | TRUE | TRUE | FALSE |
| 1690 | waaU | P27242 | TRUE | TRUE | TRUE |
| 1691 | rutC | P0AFQ5 | FALSE | FALSE | FALSE |
| 1692 | atpE | P68699 | FALSE | MISSING | FALSE |
| 1693 | nac | Q47005 | FALSE | FALSE | FALSE |
| 1694 | rlmG | P42596 | FALSE | FALSE | FALSE |
| 1695 | zapD | P36680 | FALSE | FALSE | FALSE |
| 1696 | yphH | P76586 | TRUE | TRUE | FALSE |
| 1697 | modC | P09833 | FALSE | FALSE | FALSE |
| 1698 | metC | P06721 | FALSE | FALSE | FALSE |
| 1699 | prc | P23865 | FALSE | FALSE | FALSE |
| 1700 | gcvA | P0A9F6 | FALSE | FALSE | FALSE |
| 1701 | ycdX | P75914 | TRUE | TRUE | TRUE |
| 1702 | treF | P62601 | FALSE | FALSE | FALSE |
| 1703 | nrdB | P69924 | FALSE | TRUE | TRUE |
| 1704 | acpT | P37623 | FALSE | FALSE | FALSE |
| 1705 | ycbX | P75863 | TRUE | FALSE | FALSE |
| 1706 | ebgA | P06864 | FALSE | TRUE | TRUE |
| 1707 | yigZ | P27862 | FALSE | FALSE | FALSE |
| 1708 | hpf | P0AFX0 | FALSE | FALSE | FALSE |
| 1709 | mgsA | P0A731 | FALSE | FALSE | TRUE |
| 1710 | ychQ | Q46755 | FALSE | FALSE | FALSE |
| 1711 | cutA | P69488 | FALSE | MISSING | FALSE |
| 1712 | rdsA | P37631 | FALSE | FALSE | FALSE |
| 1713 | ilvH | P00894 | FALSE | FALSE | FALSE |
| 1714 | hisG | P60757 | FALSE | FALSE | FALSE |
| 1715 | sspA | P0ACA3 | FALSE | FALSE | FALSE |
| 1716 | ygbI | P52598 | FALSE | FALSE | FALSE |
| 1717 | yadS | P0AFP0 | FALSE | FALSE | FALSE |
| 1718 | paoD | P77183 | FALSE | FALSE | FALSE |
| 1719 | pflD | P32674 | FALSE | TRUE | TRUE |
| 1720 | yahF | P77187 | FALSE | FALSE | FALSE |
| 1721 | wcaK | P71242 | FALSE | FALSE | FALSE |
| 1722 | potD | P0AFK9 | FALSE | FALSE | FALSE |
| 1723 | dsbA | P0AEG4 | FALSE | FALSE | FALSE |
| 1724 | folP | P0AC13 | FALSE | FALSE | FALSE |
| 1725 | rpsE | P0A7W1 | FALSE | FALSE | FALSE |
| 1726 | deoB | P0A6K6 | TRUE | TRUE | TRUE |
| 1727 | kdsD | P45395 | FALSE | FALSE | FALSE |
| 1728 | metL | P00562 | FALSE | FALSE | FALSE |
| 1729 | ybeQ | P77234 | FALSE | FALSE | FALSE |
| 1730 | yfhR | P77538 | TRUE | FALSE | TRUE |
| 1731 | eptC | P0CB39 | FALSE | TRUE | FALSE |
| 1732 | kdsC | P0ABZ4 | FALSE | FALSE | FALSE |
| 1733 | pliG | P76002 | FALSE | FALSE | FALSE |
| 1734 | narZ | P19319 | FALSE | FALSE | FALSE |
| 1735 | ygiS | Q46863 | FALSE | FALSE | TRUE |
| 1736 | melA | P06720 | FALSE | TRUE | FALSE |

Supplementary Table 3: E. coli predicted metal-binding proteome *(continued)*

| # | Gene Name | UniProt ID | MoM Prediction | M3D Prediction | GASS Prediction |
|---|---|---|---|---|---|
| 1737 | proV | P14175 | FALSE | FALSE | FALSE |
| 1738 | galK | P0A6T3 | FALSE | FALSE | FALSE |
| 1739 | yjaA | P09162 | FALSE | FALSE | TRUE |
| 1740 | ychA | P0AGM5 | FALSE | FALSE | FALSE |
| 1741 | ygjH | P42589 | FALSE | MISSING | TRUE |
| 1742 | ccmC | P0ABM1 | FALSE | FALSE | FALSE |
| 1743 | ybeU | P77427 | FALSE | FALSE | FALSE |
| 1744 | ccmF | P33927 | FALSE | FALSE | FALSE |
| 1745 | cca | P06961 | TRUE | TRUE | FALSE |
| 1746 | phnN | P16690 | TRUE | FALSE | FALSE |
| 1747 | ubiX | P0AG03 | FALSE | FALSE | FALSE |
| 1748 | mnmG | P0A6U3 | FALSE | FALSE | FALSE |
| 1749 | frc | P69902 | FALSE | FALSE | FALSE |
| 1750 | yeeS | P76362 | TRUE | TRUE | FALSE |
| 1751 | marA | P0ACH5 | FALSE | MISSING | FALSE |
| 1752 | ycbF | P40876 | FALSE | FALSE | FALSE |
| 1753 | puuD | P76038 | FALSE | TRUE | FALSE |
| 1754 | aceB | P08997 | FALSE | FALSE | FALSE |
| 1755 | adiY | P33234 | FALSE | FALSE | FALSE |
| 1756 | maeA | P26616 | FALSE | TRUE | FALSE |
| 1757 | yfbU | P0A8W8 | FALSE | FALSE | FALSE |
| 1758 | aslB | P25550 | TRUE | FALSE | TRUE |
| 1759 | cysJ | P38038 | FALSE | FALSE | FALSE |
| 1760 | yhhL | P37614 | FALSE | FALSE | FALSE |
| 1761 | ldhA | P52643 | FALSE | FALSE | FALSE |
| 1762 | leuB | P30125 | FALSE | FALSE | FALSE |
| 1763 | hcaR | Q47141 | FALSE | FALSE | FALSE |
| 1764 | rob | P0ACI0 | FALSE | FALSE | FALSE |
| 1765 | relE | P0C077 | FALSE | FALSE | FALSE |
| 1766 | chbC | P17334 | FALSE | FALSE | FALSE |
| 1767 | yedP | P76329 | FALSE | FALSE | TRUE |
| 1768 | curA | P76113 | FALSE | FALSE | FALSE |
| 1769 | coaE | P0A6I9 | FALSE | TRUE | FALSE |
| 1770 | rhmD | P77215 | FALSE | FALSE | TRUE |
| 1771 | speE | P09158 | TRUE | TRUE | FALSE |
| 1772 | paoB | P77324 | FALSE | TRUE | FALSE |
| 1773 | nuoF | P31979 | TRUE | FALSE | FALSE |
| 1774 | aroK | P0A6D7 | FALSE | FALSE | FALSE |
| 1775 | queG | P39288 | FALSE | FALSE | FALSE |
| 1776 | jayE | P75981 | FALSE | FALSE | FALSE |
| 1777 | rho | P0AG30 | FALSE | FALSE | FALSE |
| 1778 | dusA | P32695 | FALSE | FALSE | TRUE |
| 1779 | cobS | P36561 | FALSE | FALSE | FALSE |
| 1780 | phnO | P16691 | FALSE | TRUE | FALSE |
| 1781 | yohF | P33368 | FALSE | FALSE | FALSE |
| 1782 | chrR | P0AGE6 | FALSE | FALSE | FALSE |
| 1783 | fecC | P15030 | FALSE | FALSE | FALSE |
| 1784 | amn | P0AE12 | FALSE | FALSE | FALSE |
| 1785 | pncC | P0A6G3 | FALSE | MISSING | FALSE |
| 1786 | allA | P77731 | FALSE | TRUE | FALSE |
| 1787 | serC | P23721 | FALSE | FALSE | FALSE |
| 1788 | vsr | P09184 | TRUE | TRUE | TRUE |
| 1789 | yciO | P0AFR4 | FALSE | FALSE | FALSE |
| 1790 | yehC | P33342 | FALSE | FALSE | FALSE |
| 1791 | queD | P65870 | TRUE | TRUE | TRUE |
| 1792 | hlyE | P77335 | FALSE | FALSE | FALSE |

Supplementary Table 3: E. coli predicted metal-binding proteome *(continued)*

| # | Gene Name | UniProt ID | MoM Prediction | M3D Prediction | GASS Prediction |
|---|---|---|---|---|---|
| 1793 | fldA | P61949 | FALSE | FALSE | FALSE |
| 1794 | yidR | P31455 | FALSE | TRUE | FALSE |
| 1795 | qorB | P39315 | FALSE | FALSE | FALSE |
| 1796 | hcxB | P30178 | FALSE | FALSE | FALSE |
| 1797 | katE | P21179 | TRUE | FALSE | FALSE |
| 1798 | potB | P0AFK4 | FALSE | FALSE | FALSE |
| 1799 | dtd | P0A6M4 | FALSE | FALSE | FALSE |
| 1800 | gstB | P0ACA7 | FALSE | MISSING | FALSE |
| 1801 | psuT | P33024 | FALSE | FALSE | FALSE |
| 1802 | ugpC | P10907 | FALSE | FALSE | FALSE |
| 1803 | ispE | P62615 | FALSE | FALSE | FALSE |
| 1804 | yicN | P0ADL3 | FALSE | FALSE | FALSE |
| 1805 | puuA | P78061 | TRUE | TRUE | FALSE |
| 1806 | yehW | P33359 | FALSE | MISSING | FALSE |
| 1807 | yfaT | P76466 | FALSE | FALSE | FALSE |
| 1808 | dlgD | P37672 | FALSE | FALSE | FALSE |
| 1809 | cobC | P52086 | FALSE | FALSE | FALSE |
| 1810 | nanM | P39371 | FALSE | FALSE | FALSE |
| 1811 | dsbC | P0AEG6 | TRUE | FALSE | FALSE |
| 1812 | dxr | P45568 | FALSE | FALSE | FALSE |
| 1813 | yigE | P27840 | FALSE | FALSE | FALSE |
| 1814 | nnr | P31806 | FALSE | FALSE | TRUE |
| 1815 | yeiI | P33020 | FALSE | TRUE | FALSE |
| 1816 | xapA | P45563 | FALSE | FALSE | TRUE |
| 1817 | ymfD | P75967 | FALSE | FALSE | FALSE |
| 1818 | ygeW | Q46803 | FALSE | FALSE | FALSE |
| 1819 | rplC | P60438 | FALSE | FALSE | FALSE |
| 1820 | purN | P08179 | FALSE | FALSE | FALSE |
| 1821 | dapD | P0A9D8 | FALSE | FALSE | FALSE |
| 1822 | valS | P07118 | FALSE | FALSE | TRUE |
| 1823 | fdnH | P0AAJ3 | TRUE | FALSE | FALSE |
| 1824 | mglA | P0AAG8 | FALSE | FALSE | TRUE |
| 1825 | kdpE | P21866 | FALSE | TRUE | FALSE |
| 1826 | btsR | P0AFT5 | FALSE | FALSE | FALSE |
| 1827 | yciI | P0AB55 | FALSE | FALSE | FALSE |
| 1828 | ypjA | P52143 | FALSE | FALSE | FALSE |
| 1829 | fnr | P0A9E5 | FALSE | FALSE | FALSE |
| 1830 | ypfH | P76561 | FALSE | FALSE | FALSE |
| 1831 | sufS | P77444 | TRUE | TRUE | FALSE |
| 1832 | ybaP | P77301 | FALSE | FALSE | FALSE |
| 1833 | ibsB | C1P608 | FALSE | MISSING | FALSE |
| 1834 | ppiA | P0AFL3 | FALSE | FALSE | FALSE |
| 1835 | ykfF | P75677 | FALSE | FALSE | FALSE |
| 1836 | pgi | P0A6T1 | FALSE | FALSE | TRUE |
| 1837 | luxS | P45578 | TRUE | TRUE | FALSE |
| 1838 | eutD | P77218 | FALSE | FALSE | FALSE |
| 1839 | tilS | P52097 | TRUE | TRUE | FALSE |
| 1840 | udk | P0A8F4 | FALSE | FALSE | TRUE |
| 1841 | yhiI | P37626 | FALSE | MISSING | FALSE |
| 1842 | fimH | P08191 | FALSE | FALSE | FALSE |
| 1843 | nanC | P69856 | FALSE | FALSE | FALSE |
| 1844 | hisF | P60664 | FALSE | FALSE | FALSE |
| 1845 | lrp | P0ACJ0 | FALSE | FALSE | FALSE |
| 1846 | eutG | P76553 | TRUE | TRUE | FALSE |
| 1847 | oxc | P0AFI0 | FALSE | FALSE | FALSE |
| 1848 | tsaC | P45748 | FALSE | MISSING | FALSE |

Supplementary Table 3: E. coli predicted metal-binding proteome *(continued)*

| # | Gene Name | UniProt ID | MoM Prediction | M3D Prediction | GASS Prediction |
|---|---|---|---|---|---|
| 1849 | tmk | P0A720 | FALSE | TRUE | FALSE |
| 1850 | sufE | P76194 | FALSE | FALSE | FALSE |
| 1851 | glcA | Q46839 | FALSE | FALSE | FALSE |
| 1852 | dinG | P27296 | FALSE | TRUE | FALSE |
| 1853 | anmK | P77570 | FALSE | FALSE | TRUE |
| 1854 | hemH | P23871 | FALSE | FALSE | FALSE |
| 1855 | polA | P00582 | FALSE | FALSE | FALSE |
| 1856 | guaB | P0ADG7 | FALSE | FALSE | FALSE |
| 1857 | ileS | P00956 | TRUE | TRUE | TRUE |
| 1858 | allK | P37306 | FALSE | FALSE | FALSE |
| 1859 | apaG | P62672 | FALSE | FALSE | FALSE |
| 1860 | kilR | P38393 | FALSE | FALSE | FALSE |
| 1861 | fhuA | P06971 | FALSE | FALSE | FALSE |
| 1862 | ygdB | P08370 | FALSE | FALSE | FALSE |
| 1863 | acpS | P24224 | FALSE | FALSE | FALSE |
| 1864 | pncA | P21369 | TRUE | TRUE | TRUE |
| 1865 | yggT | P64564 | FALSE | MISSING | FALSE |
| 1866 | ydfX | P76165 | FALSE | MISSING | FALSE |
| 1867 | ygcN | Q46904 | FALSE | FALSE | FALSE |
| 1868 | hsdS | P05719 | FALSE | TRUE | FALSE |
| 1869 | yegX | P76421 | FALSE | FALSE | TRUE |
| 1870 | sodA | P00448 | TRUE | TRUE | FALSE |
| 1871 | yebB | P24238 | FALSE | FALSE | FALSE |
| 1872 | folD | P24186 | FALSE | TRUE | FALSE |
| 1873 | ycjR | P76044 | FALSE | TRUE | FALSE |
| 1874 | ulaA | P39301 | FALSE | FALSE | TRUE |
| 1875 | srlD | P05707 | FALSE | FALSE | FALSE |
| 1876 | yieL | P31471 | FALSE | FALSE | FALSE |
| 1877 | rpsB | P0A7V0 | FALSE | FALSE | FALSE |
| 1878 | paaC | P76079 | FALSE | FALSE | FALSE |
| 1879 | otsB | P31678 | FALSE | FALSE | FALSE |
| 1880 | ybjG | P75806 | FALSE | FALSE | FALSE |
| 1881 | ygiN | P0ADU2 | TRUE | FALSE | TRUE |
| 1882 | sdhE | P64559 | FALSE | FALSE | FALSE |
| 1883 | arsC | P0AB96 | FALSE | FALSE | TRUE |
| 1884 | yajL | Q46948 | FALSE | FALSE | FALSE |
| 1885 | ydiQ | P76201 | FALSE | FALSE | FALSE |
| 1886 | pgk | P0A799 | FALSE | FALSE | FALSE |
| 1887 | ptrB | P24555 | FALSE | FALSE | TRUE |
| 1888 | yfeS | P78271 | FALSE | FALSE | FALSE |
| 1889 | nadC | P30011 | FALSE | TRUE | FALSE |
| 1890 | purM | P08178 | FALSE | TRUE | FALSE |
| 1891 | frmB | P51025 | FALSE | FALSE | FALSE |
| 1892 | dut | P06968 | FALSE | FALSE | FALSE |
| 1893 | pxpA | P75746 | TRUE | TRUE | TRUE |
| 1894 | gloC | P75849 | TRUE | TRUE | TRUE |
| 1895 | ggaR | P0ACM5 | FALSE | FALSE | FALSE |
| 1896 | pepN | P04825 | TRUE | TRUE | TRUE |
| 1897 | yiaY | P37686 | TRUE | TRUE | TRUE |
| 1898 | xylF | P37387 | FALSE | FALSE | FALSE |
| 1899 | murE | P22188 | FALSE | TRUE | FALSE |
| 1900 | proX | P0AFM2 | FALSE | FALSE | FALSE |
| 1901 | rpmJ | P0A7Q6 | TRUE | TRUE | TRUE |
| 1902 | trmH | P0AGJ2 | TRUE | FALSE | FALSE |
| 1903 | fsr | P52067 | FALSE | TRUE | FALSE |
| 1904 | fetA | P77279 | FALSE | FALSE | FALSE |

Supplementary Table 3: E. coli predicted metal-binding proteome *(continued)*

| # | Gene Name | UniProt ID | MoM Prediction | M3D Prediction | GASS Prediction |
|---|---|---|---|---|---|
| 1905 | xdhA | Q46799 | TRUE | TRUE | FALSE |
| 1906 | ynfG | P0AAJ1 | TRUE | FALSE | FALSE |
| 1907 | pldB | P07000 | FALSE | TRUE | FALSE |
| 1908 | hyaA | P69739 | TRUE | TRUE | FALSE |
| 1909 | ybhK | P75767 | FALSE | FALSE | FALSE |
| 1910 | ybjJ | P75810 | FALSE | FALSE | FALSE |
| 1911 | trpA | P0A877 | FALSE | FALSE | FALSE |
| 1912 | mtlD | P09424 | FALSE | FALSE | FALSE |
| 1913 | moaE | P30749 | FALSE | FALSE | FALSE |
| 1914 | ruvB | P0A812 | FALSE | FALSE | FALSE |
| 1915 | trpS | P00954 | FALSE | TRUE | FALSE |
| 1916 | ispU | P60472 | FALSE | FALSE | FALSE |
| 1917 | hemB | P0ACB2 | TRUE | TRUE | FALSE |
| 1918 | gadA | P69908 | FALSE | FALSE | FALSE |
| 1919 | yfdS | P76515 | FALSE | MISSING | FALSE |
| 1920 | wecA | P0AC78 | TRUE | FALSE | FALSE |
| 1921 | rfbX | P37746 | FALSE | FALSE | FALSE |
| 1922 | ygcP | Q46906 | FALSE | FALSE | FALSE |
| 1923 | purA | P0A7D4 | FALSE | FALSE | FALSE |
| 1924 | rsmE | P0AGL7 | FALSE | FALSE | TRUE |
| 1925 | yjjI | P37342 | FALSE | FALSE | FALSE |
| 1926 | ygcE | P55138 | FALSE | FALSE | FALSE |
| 1927 | bioD1 | P13000 | FALSE | TRUE | FALSE |
| 1928 | murB | P08373 | FALSE | FALSE | FALSE |
| 1929 | ydeR | P77294 | FALSE | FALSE | FALSE |
| 1930 | fhuD | P07822 | FALSE | FALSE | FALSE |
| 1931 | ymdB | P0A8D6 | FALSE | FALSE | FALSE |
| 1932 | hcr | P75824 | TRUE | FALSE | FALSE |
| 1933 | fucU | P0AEN8 | FALSE | FALSE | FALSE |
| 1934 | yjbQ | P0AF48 | TRUE | TRUE | TRUE |
| 1935 | gabT | P22256 | FALSE | TRUE | FALSE |
| 1936 | bioB | P12996 | FALSE | FALSE | FALSE |
| 1937 | asd | P0A9Q9 | FALSE | TRUE | FALSE |
| 1938 | tam | P76145 | FALSE | FALSE | FALSE |
| 1939 | prmC | P0ACC1 | FALSE | FALSE | FALSE |
| 1940 | yjhP | P39367 | FALSE | FALSE | FALSE |
| 1941 | htpG | P0A6Z3 | FALSE | FALSE | FALSE |
| 1942 | yodB | P76345 | FALSE | TRUE | FALSE |
| 1943 | ribE | P61714 | FALSE | MISSING | FALSE |
| 1944 | ppnN | P0ADR8 | FALSE | FALSE | FALSE |
| 1945 | def | P0A6K3 | TRUE | TRUE | FALSE |
| 1946 | acs | P27550 | TRUE | FALSE | FALSE |
| 1947 | ygbF | P45956 | FALSE | MISSING | FALSE |
| 1948 | pepA | P68767 | FALSE | TRUE | FALSE |
| 1949 | rsmJ | P68567 | FALSE | TRUE | FALSE |
| 1950 | yfaY | P77808 | FALSE | TRUE | FALSE |
| 1951 | ydeM | P76134 | FALSE | FALSE | TRUE |
| 1952 | yafV | Q47679 | FALSE | FALSE | FALSE |
| 1953 | chbG | P37794 | TRUE | TRUE | TRUE |
| 1954 | narL | P0AF28 | FALSE | TRUE | FALSE |
| 1955 | yfdE | P76518 | FALSE | FALSE | TRUE |
| 1956 | malY | P23256 | FALSE | FALSE | FALSE |
| 1957 | ampD | P13016 | TRUE | TRUE | FALSE |
| 1958 | motA | P09348 | FALSE | FALSE | FALSE |
| 1959 | mmuM | Q47690 | TRUE | TRUE | TRUE |
| 1960 | malG | P68183 | FALSE | FALSE | FALSE |

Supplementary Table 3: E. coli predicted metal-binding proteome *(continued)*

| # | Gene Name | UniProt ID | MoM Prediction | M3D Prediction | GASS Prediction |
|---|---|---|---|---|---|
| 1961 | bfr | P0ABD3 | FALSE | TRUE | FALSE |
| 1962 | aroM | P0AE28 | FALSE | FALSE | FALSE |
| 1963 | yfbP | P76486 | FALSE | FALSE | FALSE |
| 1964 | waaQ | P25742 | FALSE | FALSE | TRUE |
| 1965 | wecE | P27833 | FALSE | FALSE | FALSE |
| 1966 | trhO | P24188 | TRUE | TRUE | TRUE |
| 1967 | rtcA | P46849 | FALSE | FALSE | FALSE |
| 1968 | pyrG | P0A7E5 | FALSE | FALSE | FALSE |
| 1969 | crr | P69783 | FALSE | TRUE | FALSE |
| 1970 | hycC | P16429 | FALSE | FALSE | FALSE |
| 1971 | rimM | P0A7X6 | FALSE | MISSING | FALSE |
| 1972 | nikB | P33591 | TRUE | FALSE | FALSE |
| 1973 | ynfC | P67553 | FALSE | FALSE | FALSE |
| 1974 | insN2 | P39212 | FALSE | MISSING | FALSE |
| 1975 | fdoI | P0AEL0 | FALSE | TRUE | TRUE |
| 1976 | frsA | P04335 | FALSE | FALSE | TRUE |
| 1977 | ihfB | P0A6Y1 | FALSE | FALSE | FALSE |
| 1978 | grxB | P0AC59 | FALSE | FALSE | FALSE |
| 1979 | ygfM | P64557 | FALSE | FALSE | FALSE |
| 1980 | yhfG | P0ADX5 | FALSE | MISSING | FALSE |
| 1981 | yiiX | P32167 | FALSE | FALSE | FALSE |
| 1982 | manA | P00946 | FALSE | TRUE | FALSE |
| 1983 | yihS | P32140 | TRUE | FALSE | TRUE |
| 1984 | ybgI | P0AFP6 | FALSE | TRUE | FALSE |
| 1985 | ycgR | P76010 | FALSE | FALSE | FALSE |
| 1986 | mdoD | P40120 | FALSE | TRUE | TRUE |
| 1987 | yibD | P11290 | FALSE | FALSE | FALSE |
| 1988 | dmsC | P18777 | FALSE | FALSE | FALSE |
| 1989 | tdcD | P11868 | TRUE | TRUE | FALSE |
| 1990 | panD | P0A790 | FALSE | FALSE | FALSE |
| 1991 | yhaI | P64592 | FALSE | MISSING | FALSE |
| 1992 | hicB | P67697 | FALSE | FALSE | FALSE |
| 1993 | ilvY | P05827 | FALSE | FALSE | FALSE |
| 1994 | ysgA | P56262 | FALSE | TRUE | FALSE |
| 1995 | gdhA | P00370 | FALSE | FALSE | FALSE |
| 1996 | mscS | P0C0S1 | FALSE | MISSING | FALSE |
| 1997 | mutY | P17802 | FALSE | FALSE | FALSE |
| 1998 | yiaC | P37664 | FALSE | FALSE | FALSE |
| 1999 | rpiB | P37351 | FALSE | TRUE | FALSE |
| 2000 | rrrD | P78285 | TRUE | FALSE | TRUE |
| 2001 | abpB | P52126 | FALSE | FALSE | FALSE |
| 2002 | ubiG | P17993 | FALSE | FALSE | FALSE |
| 2003 | rhaM | P32156 | FALSE | FALSE | FALSE |
| 2004 | dacA | P0AEB2 | FALSE | FALSE | FALSE |
| 2005 | ppc | P00864 | FALSE | FALSE | FALSE |
| 2006 | ydgC | P0ACX0 | FALSE | MISSING | FALSE |
| 2007 | yciF | P21362 | FALSE | FALSE | FALSE |
| 2008 | ygjR | P42599 | FALSE | FALSE | TRUE |
| 2009 | ybiX | P75779 | TRUE | TRUE | TRUE |
| 2010 | sgbH | P37678 | FALSE | FALSE | FALSE |
| 2011 | pagP | P37001 | FALSE | FALSE | FALSE |
| 2012 | arsR | P37309 | TRUE | TRUE | FALSE |
| 2013 | yhaH | P64590 | FALSE | FALSE | TRUE |
| 2014 | insE3 | P0CF68 | FALSE | MISSING | FALSE |
| 2015 | rplF | P0AG55 | FALSE | MISSING | FALSE |
| 2016 | kdpA | P03959 | FALSE | FALSE | FALSE |

Supplementary Table 3: E. coli predicted metal-binding proteome *(continued)*

| # | Gene Name | UniProt ID | MoM Prediction | M3D Prediction | GASS Prediction |
|---|---|---|---|---|---|
| 2017 | citE | P0A9I1 | FALSE | FALSE | FALSE |
| 2018 | ves | P76214 | FALSE | FALSE | FALSE |
| 2019 | tolB | P0A855 | FALSE | FALSE | FALSE |
| 2020 | ycjV | P77481 | TRUE | FALSE | FALSE |
| 2021 | dgoD | Q6BF17 | FALSE | TRUE | FALSE |
| 2022 | ydhP | P77389 | FALSE | FALSE | FALSE |
| 2023 | ybiB | P30177 | FALSE | FALSE | FALSE |
| 2024 | phnD | P16682 | FALSE | FALSE | FALSE |
| 2025 | nfi | P68739 | FALSE | FALSE | FALSE |
| 2026 | hdhA | P0AET8 | FALSE | FALSE | FALSE |
| 2027 | fecE | P15031 | FALSE | FALSE | FALSE |
| 2028 | wcaB | P0ACC9 | FALSE | FALSE | FALSE |
| 2029 | pntB | P0AB67 | FALSE | FALSE | FALSE |
| 2030 | ygjI | P42590 | FALSE | FALSE | FALSE |
| 2031 | mnaT | P76112 | FALSE | FALSE | FALSE |
| 2032 | yqcA | P65367 | FALSE | MISSING | TRUE |
| 2033 | yeiB | P25747 | FALSE | FALSE | FALSE |
| 2034 | modE | P0A9G8 | FALSE | FALSE | FALSE |
| 2035 | frdB | P0AC47 | FALSE | FALSE | TRUE |
| 2036 | ydjI | P77704 | TRUE | TRUE | TRUE |
| 2037 | puuB | P37906 | FALSE | FALSE | FALSE |
| 2038 | ydjY | P76220 | TRUE | TRUE | FALSE |
| 2039 | ppiD | P0ADY1 | FALSE | FALSE | FALSE |
| 2040 | gsiB | P75797 | FALSE | FALSE | FALSE |
| 2041 | zapA | P0ADS2 | FALSE | MISSING | FALSE |
| 2042 | cyaY | P27838 | FALSE | TRUE | FALSE |
| 2043 | asnC | P0ACI6 | FALSE | FALSE | FALSE |
| 2044 | yeeL | P76349 | FALSE | FALSE | FALSE |
| 2045 | nanK | P45425 | TRUE | TRUE | FALSE |
| 2046 | yahJ | P77554 | TRUE | TRUE | TRUE |
| 2047 | yjeM | P39282 | FALSE | FALSE | FALSE |
| 2048 | fliD | P24216 | FALSE | FALSE | FALSE |
| 2049 | hofM | P45753 | FALSE | FALSE | FALSE |
| 2050 | bdcA | P39333 | FALSE | FALSE | FALSE |
| 2051 | sufA | P77667 | FALSE | FALSE | FALSE |
| 2052 | ydjL | P77539 | TRUE | TRUE | FALSE |
| 2053 | trkA | P0AGI8 | FALSE | FALSE | FALSE |
| 2054 | mog | P0AF03 | FALSE | FALSE | FALSE |
| 2055 | rpsF | P02358 | FALSE | FALSE | FALSE |
| 2056 | minD | P0AEZ3 | FALSE | MISSING | FALSE |
| 2057 | gloA | P0AC81 | FALSE | FALSE | FALSE |
| 2058 | trmD | P0A873 | FALSE | TRUE | FALSE |
| 2059 | wbbK | P37751 | FALSE | FALSE | TRUE |
| 2060 | purL | P15254 | FALSE | TRUE | FALSE |
| 2061 | nadK | P0A7B3 | TRUE | FALSE | FALSE |
| 2062 | yccJ | P0AB14 | FALSE | FALSE | FALSE |
| 2063 | acpH | P21515 | FALSE | TRUE | FALSE |
| 2064 | tyrA | P07023 | FALSE | FALSE | FALSE |
| 2065 | yeiW | P0AFT8 | TRUE | FALSE | FALSE |
| 2066 | cyoE | P0AEA5 | FALSE | FALSE | FALSE |
| 2067 | panE | P0A9J4 | FALSE | TRUE | FALSE |
| 2068 | epmC | P76938 | TRUE | TRUE | FALSE |
| 2069 | gpmB | P0A7A2 | FALSE | FALSE | FALSE |
| 2070 | aat | P0A8P1 | FALSE | FALSE | TRUE |
| 2071 | ybfF | P75736 | TRUE | FALSE | FALSE |
| 2072 | dacB | P24228 | FALSE | FALSE | FALSE |

Supplementary Table 3: E. coli predicted metal-binding proteome *(continued)*

| # | Gene Name | UniProt ID | MoM Prediction | M3D Prediction | GASS Prediction |
|---|---|---|---|---|---|
| 2073 | truB | P60340 | FALSE | TRUE | FALSE |
| 2074 | lasT | P37005 | FALSE | FALSE | FALSE |
| 2075 | kdpC | P03961 | FALSE | FALSE | FALSE |
| 2076 | rpsR | P0A7T7 | FALSE | MISSING | FALSE |
| 2077 | apaH | P05637 | TRUE | FALSE | FALSE |
| 2078 | fepD | P23876 | FALSE | FALSE | FALSE |
| 2079 | narH | P11349 | TRUE | FALSE | FALSE |
| 2080 | lipA | P60716 | FALSE | FALSE | TRUE |
| 2081 | yfeR | P0ACR7 | FALSE | FALSE | FALSE |
| 2082 | fabG | P0AEK2 | FALSE | FALSE | FALSE |
| 2083 | aldB | P37685 | FALSE | FALSE | FALSE |
| 2084 | selA | P0A821 | FALSE | FALSE | FALSE |
| 2085 | pykF | P0AD61 | TRUE | FALSE | FALSE |
| 2086 | glmU | P0ACC7 | FALSE | FALSE | FALSE |
| 2087 | rclA | P77212 | FALSE | TRUE | FALSE |
| 2088 | xynR | P77300 | FALSE | FALSE | FALSE |
| 2089 | dapF | P0A6K1 | FALSE | FALSE | FALSE |
| 2090 | yeaR | P64488 | TRUE | TRUE | TRUE |
| 2091 | pqqL | P31828 | FALSE | TRUE | TRUE |
| 2092 | ais | P45565 | FALSE | TRUE | FALSE |
| 2093 | ychN | P0AB52 | FALSE | FALSE | FALSE |
| 2094 | ftsI | P0AD68 | FALSE | FALSE | FALSE |
| 2095 | yncG | P76117 | FALSE | FALSE | FALSE |
| 2096 | pqqU | P76115 | FALSE | FALSE | FALSE |
| 2097 | ydiH | P64476 | FALSE | MISSING | FALSE |
| 2098 | rimK | P0C0U4 | FALSE | FALSE | FALSE |
| 2099 | ybhL | P0AAC4 | FALSE | FALSE | FALSE |
| 2100 | nrfF | P32711 | FALSE | FALSE | FALSE |
| 2101 | trxC | P0AGG4 | TRUE | TRUE | TRUE |
| 2102 | mhpD | P77608 | FALSE | TRUE | FALSE |
| 2103 | cdaR | P37047 | FALSE | FALSE | FALSE |
| 2104 | ppk | P0A7B1 | FALSE | FALSE | TRUE |
| 2105 | yfhH | P37767 | FALSE | FALSE | FALSE |
| 2106 | ulaR | P0A9W0 | FALSE | FALSE | FALSE |
| 2107 | glnA | P0A9C5 | FALSE | TRUE | TRUE |
| 2108 | dcuB | P0ABN9 | FALSE | FALSE | FALSE |
| 2109 | yejM | P0AD27 | FALSE | FALSE | FALSE |
| 2110 | mdtI | P69210 | FALSE | FALSE | FALSE |
| 2111 | mraY | P0A6W3 | FALSE | FALSE | FALSE |
| 2112 | cobU | P0AE76 | FALSE | FALSE | FALSE |
| 2113 | ydiJ | P77748 | TRUE | TRUE | TRUE |
| 2114 | nadB | P10902 | TRUE | FALSE | TRUE |
| 2115 | eutE | P77445 | FALSE | TRUE | FALSE |
| 2116 | ugpQ | P10908 | FALSE | FALSE | FALSE |
| 2117 | rpe | P0AG07 | TRUE | TRUE | TRUE |
| 2118 | yeiS | P64536 | FALSE | MISSING | FALSE |
| 2119 | hipA | P23874 | FALSE | FALSE | FALSE |
| 2120 | glcG | P0AEQ1 | FALSE | FALSE | FALSE |
| 2121 | pphB | P55799 | TRUE | FALSE | TRUE |
| 2122 | sgcR | P39361 | FALSE | FALSE | FALSE |
| 2123 | ratA | P0AGL5 | FALSE | FALSE | FALSE |
| 2124 | rlmM | P0ADR6 | FALSE | FALSE | TRUE |
| 2125 | ebgC | P0AC73 | FALSE | FALSE | FALSE |
| 2126 | phoE | P02932 | FALSE | FALSE | FALSE |
| 2127 | hupA | P0ACF0 | FALSE | MISSING | FALSE |
| 2128 | yjhH | P39359 | FALSE | TRUE | FALSE |

Supplementary Table 3: E. coli predicted metal-binding proteome *(continued)*

| # | Gene Name | UniProt ID | MoM Prediction | M3D Prediction | GASS Prediction |
|---|---|---|---|---|---|
| 2129 | ygaV | P77295 | FALSE | FALSE | FALSE |
| 2130 | yhjC | P37641 | FALSE | FALSE | FALSE |
| 2131 | yfcL | P64540 | FALSE | FALSE | FALSE |
| 2132 | selD | P16456 | FALSE | FALSE | FALSE |
| 2133 | fumE | P11663 | FALSE | FALSE | FALSE |
| 2134 | hscB | P0A6L9 | FALSE | FALSE | FALSE |
| 2135 | ydcH | P0ACW6 | FALSE | MISSING | FALSE |
| 2136 | thiL | P0AGG0 | FALSE | FALSE | FALSE |
| 2137 | trmJ | P0AE01 | FALSE | FALSE | FALSE |
| 2138 | gltJ | P0AER3 | FALSE | MISSING | FALSE |
| 2139 | mdoB | P39401 | TRUE | TRUE | FALSE |
| 2140 | trpB | P0A879 | FALSE | FALSE | TRUE |
| 2141 | yeaO | P76243 | FALSE | FALSE | FALSE |
| 2142 | yigI | P0ADP2 | FALSE | MISSING | FALSE |
| 2143 | otsA | P31677 | FALSE | FALSE | FALSE |
| 2144 | yhaV | P64594 | FALSE | FALSE | FALSE |
| 2145 | pmrD | P37590 | FALSE | FALSE | FALSE |
| 2146 | rplT | P0A7L3 | FALSE | MISSING | FALSE |
| 2147 | rihB | P33022 | FALSE | FALSE | TRUE |
| 2148 | fabB | P0A953 | FALSE | FALSE | FALSE |
| 2149 | fluC | P37002 | FALSE | FALSE | FALSE |
| 2150 | gltA | P0ABH7 | FALSE | FALSE | FALSE |
| 2151 | cdsA | P0ABG1 | FALSE | FALSE | TRUE |
| 2152 | livK | P04816 | FALSE | FALSE | FALSE |
| 2153 | higB | P64578 | FALSE | FALSE | FALSE |
| 2154 | yadI | P36881 | FALSE | TRUE | FALSE |
| 2155 | dusC | P33371 | FALSE | FALSE | FALSE |
| 2156 | astC | P77581 | FALSE | FALSE | FALSE |
| 2157 | yjaB | P09163 | FALSE | TRUE | FALSE |
| 2158 | birA | P06709 | FALSE | MISSING | FALSE |
| 2159 | ykfA | P75678 | FALSE | FALSE | FALSE |
| 2160 | efp | P0A6N4 | FALSE | MISSING | FALSE |
| 2161 | pyrE | P0A7E3 | FALSE | FALSE | FALSE |
| 2162 | speF | P24169 | FALSE | TRUE | TRUE |
| 2163 | kbaY | P0AB74 | TRUE | TRUE | TRUE |
| 2164 | yjeT | P0AF73 | FALSE | MISSING | FALSE |
| 2165 | ydiF | P37766 | FALSE | TRUE | FALSE |
| 2166 | ldrA | P0DPD0 | FALSE | FALSE | FALSE |
| 2167 | ltaE | P75823 | FALSE | FALSE | FALSE |
| 2168 | mltC | P0C066 | FALSE | FALSE | FALSE |
| 2169 | nudG | P77788 | TRUE | FALSE | FALSE |
| 2170 | yobI | C1P604 | FALSE | FALSE | FALSE |
| 2171 | csgC | P52107 | FALSE | MISSING | FALSE |
| 2172 | tsaE | P0AF67 | FALSE | FALSE | FALSE |
| 2173 | artM | P0AE30 | FALSE | FALSE | FALSE |
| 2174 | msrB | P0A746 | TRUE | TRUE | FALSE |
| 2175 | sprT | P39902 | TRUE | TRUE | TRUE |
| 2176 | preA | P25889 | FALSE | FALSE | FALSE |
| 2177 | glgA | P0A6U8 | TRUE | FALSE | FALSE |
| 2178 | yjiG | P0AEH8 | FALSE | FALSE | TRUE |
| 2179 | nlpA | P04846 | FALSE | FALSE | FALSE |
| 2180 | thrA | P00561 | FALSE | FALSE | FALSE |
| 2181 | menC | P29208 | FALSE | FALSE | FALSE |
| 2182 | trpR | P0A881 | FALSE | MISSING | FALSE |
| 2183 | rnhB | P10442 | FALSE | TRUE | FALSE |
| 2184 | glaH | P76621 | TRUE | TRUE | FALSE |

Supplementary Table 3: E. coli predicted metal-binding proteome *(continued)*

| # | Gene Name | UniProt ID | MoM Prediction | M3D Prediction | GASS Prediction |
|---|---|---|---|---|---|
| 2185 | artQ | P0AE34 | FALSE | FALSE | FALSE |
| 2186 | pstB | P0AAH0 | FALSE | FALSE | FALSE |
| 2187 | erfK | P39176 | FALSE | FALSE | FALSE |
| 2188 | yqjG | P42620 | FALSE | FALSE | FALSE |
| 2189 | hldD | P67910 | FALSE | FALSE | FALSE |
| 2190 | yafX | P75676 | FALSE | TRUE | FALSE |
| 2191 | ydeK | P32051 | TRUE | FALSE | FALSE |
| 2192 | ybiW | P75793 | FALSE | TRUE | TRUE |
| 2193 | yeiE | P0ACR4 | FALSE | FALSE | FALSE |
| 2194 | lysA | P00861 | TRUE | FALSE | FALSE |
| 2195 | menB | P0ABU0 | FALSE | FALSE | TRUE |
| 2196 | yigL | P27848 | FALSE | TRUE | FALSE |
| 2197 | atoB | P76461 | FALSE | TRUE | FALSE |
| 2198 | ydiZ | P64479 | FALSE | FALSE | FALSE |
| 2199 | mprA | P0ACR9 | FALSE | TRUE | FALSE |
| 2200 | rclC | P75685 | FALSE | FALSE | FALSE |
| 2201 | rzpD | P75719 | FALSE | MISSING | FALSE |
| 2202 | moaB | P0AEZ9 | FALSE | FALSE | FALSE |
| 2203 | dauA | P0AFR2 | FALSE | FALSE | FALSE |
| 2204 | ycgN | P0A8L5 | FALSE | FALSE | FALSE |
| 2205 | rsxG | P77285 | FALSE | FALSE | FALSE |
| 2206 | elfD | P75856 | FALSE | FALSE | FALSE |
| 2207 | glgP | P0AC86 | FALSE | FALSE | FALSE |
| 2208 | hemE | P29680 | FALSE | FALSE | TRUE |
| 2209 | plsB | P0A7A7 | FALSE | TRUE | FALSE |
| 2210 | lpxC | P0A725 | TRUE | TRUE | FALSE |
| 2211 | apt | P69503 | FALSE | FALSE | FALSE |
| 2212 | frlC | P45541 | FALSE | TRUE | FALSE |
| 2213 | murF | P11880 | FALSE | FALSE | TRUE |
| 2214 | glnS | P00962 | FALSE | FALSE | FALSE |
| 2215 | tsx | P0A927 | FALSE | FALSE | FALSE |
| 2216 | yniA | P77739 | FALSE | FALSE | FALSE |
| 2217 | nagA | P0AF18 | FALSE | TRUE | TRUE |
| 2218 | yqaE | P0AE42 | FALSE | MISSING | FALSE |
| 2219 | yhhW | P46852 | TRUE | TRUE | TRUE |
| 2220 | tus | P16525 | FALSE | TRUE | FALSE |
| 2221 | frvX | P32153 | FALSE | TRUE | FALSE |
| 2222 | pyrF | P08244 | FALSE | FALSE | FALSE |
| 2223 | dsbG | P77202 | FALSE | FALSE | FALSE |
| 2224 | ykgF | P77536 | FALSE | FALSE | TRUE |
| 2225 | yjjX | P39411 | FALSE | FALSE | FALSE |
| 2226 | nagK | P75959 | TRUE | TRUE | FALSE |
| 2227 | phnI | P16687 | FALSE | TRUE | FALSE |
| 2228 | endA | P25736 | TRUE | FALSE | FALSE |
| 2229 | aes | P23872 | FALSE | TRUE | FALSE |
| 2230 | rpiA | P0A7Z0 | FALSE | FALSE | FALSE |
| 2231 | fadR | P0A8V6 | FALSE | FALSE | FALSE |
| 2232 | paaZ | P77455 | FALSE | TRUE | FALSE |
| 2233 | frlA | P45539 | FALSE | MISSING | FALSE |
| 2234 | yhfZ | P45552 | FALSE | FALSE | FALSE |
| 2235 | ybgP | P75749 | FALSE | MISSING | FALSE |
| 2236 | acpP | P0A6A8 | FALSE | FALSE | FALSE |
| 2237 | paaD | P76080 | TRUE | TRUE | TRUE |
| 2238 | tabA | P0AF96 | FALSE | TRUE | FALSE |
| 2239 | ymdE | Q7DFV4 | FALSE | FALSE | FALSE |
| 2240 | rlmN | P36979 | FALSE | FALSE | FALSE |

Supplementary Table 3: E. coli predicted metal-binding proteome *(continued)*

| # | Gene Name | UniProt ID | MoM Prediction | M3D Prediction | GASS Prediction |
|---|---|---|---|---|---|
| 2241 | ydjF | P77721 | FALSE | FALSE | FALSE |
| 2242 | yhfA | P0ADX1 | FALSE | FALSE | FALSE |
| 2243 | mdh | P61889 | FALSE | FALSE | FALSE |
| 2244 | icd | P08200 | FALSE | FALSE | FALSE |
| 2245 | tyrB | P04693 | FALSE | FALSE | FALSE |
| 2246 | paaF | P76082 | FALSE | FALSE | FALSE |
| 2247 | fic | P20605 | FALSE | FALSE | FALSE |
| 2248 | rcnR | P64530 | FALSE | TRUE | FALSE |
| 2249 | mltG | P28306 | FALSE | FALSE | FALSE |
| 2250 | yghU | Q46845 | TRUE | FALSE | FALSE |
| 2251 | hrpB | P37024 | FALSE | FALSE | FALSE |
| 2252 | tynA | P46883 | TRUE | TRUE | TRUE |
| 2253 | idnR | P39343 | FALSE | FALSE | FALSE |
| 2254 | nagD | P0AF24 | FALSE | FALSE | FALSE |
| 2255 | yjeN | P39283 | FALSE | MISSING | FALSE |
| 2256 | yibA | P0ADK6 | FALSE | FALSE | TRUE |
| 2257 | yaaJ | P30143 | FALSE | FALSE | FALSE |
| 2258 | dapE | P0AED7 | FALSE | TRUE | FALSE |
| 2259 | yjiR | P39389 | FALSE | FALSE | FALSE |
| 2260 | ygaP | P55734 | FALSE | FALSE | FALSE |
| 2261 | oppF | P77737 | FALSE | FALSE | FALSE |
| 2262 | slt | P0AGC3 | FALSE | FALSE | FALSE |
| 2263 | fcl | P32055 | FALSE | TRUE | FALSE |
| 2264 | glpE | P0A6V5 | FALSE | FALSE | FALSE |
| 2265 | hcaF | Q47140 | FALSE | FALSE | FALSE |
| 2266 | eutL | P76541 | FALSE | FALSE | FALSE |
| 2267 | leuD | P30126 | FALSE | FALSE | FALSE |
| 2268 | glxK | P77364 | FALSE | FALSE | FALSE |
| 2269 | xthA | P09030 | FALSE | FALSE | FALSE |
| 2270 | treC | P28904 | FALSE | TRUE | FALSE |
| 2271 | wcaE | P71239 | FALSE | FALSE | FALSE |
| 2272 | ydcI | P77171 | FALSE | FALSE | FALSE |
| 2273 | frlD | P45543 | FALSE | FALSE | FALSE |
| 2274 | rcdA | P75811 | FALSE | FALSE | FALSE |
| 2275 | nfsB | P38489 | FALSE | FALSE | FALSE |
| 2276 | allC | P77425 | FALSE | TRUE | FALSE |
| 2277 | uidA | P05804 | TRUE | FALSE | FALSE |
| 2278 | blc | P0A901 | FALSE | FALSE | FALSE |
| 2279 | ppdD | P36647 | TRUE | MISSING | FALSE |
| 2280 | yfiH | P33644 | FALSE | TRUE | FALSE |
| 2281 | bioF | P12998 | FALSE | FALSE | FALSE |
| 2282 | yciK | P31808 | FALSE | FALSE | FALSE |
| 2283 | aroD | P05194 | FALSE | FALSE | FALSE |
| 2284 | eutC | P19636 | FALSE | FALSE | FALSE |
| 2285 | hinT | P0ACE7 | FALSE | FALSE | FALSE |
| 2286 | lexA | P0A7C2 | FALSE | FALSE | FALSE |
| 2287 | glaR | P37338 | TRUE | TRUE | TRUE |
| 2288 | yicG | P0AGM2 | FALSE | FALSE | FALSE |
| 2289 | rsmC | P39406 | FALSE | TRUE | FALSE |
| 2290 | yhbS | P63417 | TRUE | FALSE | FALSE |
| 2291 | msrC | P76270 | FALSE | FALSE | FALSE |
| 2292 | plsY | P60782 | FALSE | FALSE | FALSE |
| 2293 | dgcZ | P31129 | TRUE | TRUE | FALSE |
| 2294 | nadE | P18843 | FALSE | FALSE | FALSE |
| 2295 | hyfH | P77423 | TRUE | TRUE | FALSE |
| 2296 | ecnB | P0ADB7 | FALSE | MISSING | FALSE |

Supplementary Table 3: E. coli predicted metal-binding proteome *(continued)*

| # | Gene Name | UniProt ID | MoM Prediction | M3D Prediction | GASS Prediction |
|---|---|---|---|---|---|
| 2297 | yrfG | P64636 | TRUE | TRUE | FALSE |
| 2298 | sfsA | P0A823 | FALSE | FALSE | FALSE |
| 2299 | ynfL | P77559 | FALSE | FALSE | FALSE |
| 2300 | recJ | P21893 | FALSE | TRUE | FALSE |
| 2301 | rdgC | P36767 | FALSE | FALSE | FALSE |
| 2302 | ubiV | P45475 | FALSE | FALSE | FALSE |
| 2303 | ssuE | P80644 | FALSE | FALSE | FALSE |
| 2304 | nudC | P32664 | TRUE | TRUE | TRUE |
| 2305 | sapD | P0AAH4 | FALSE | FALSE | FALSE |
| 2306 | ycaO | P75838 | FALSE | FALSE | FALSE |
| 2307 | ubiC | P26602 | FALSE | MISSING | FALSE |
| 2308 | fumA | P0AC33 | FALSE | FALSE | FALSE |
| 2309 | yjfK | P39293 | FALSE | FALSE | FALSE |
| 2310 | uraA | P0AGM7 | FALSE | FALSE | FALSE |
| 2311 | thiK | P75948 | FALSE | FALSE | FALSE |
| 2312 | ydiV | P76204 | TRUE | TRUE | TRUE |
| 2313 | glmS | P17169 | FALSE | FALSE | TRUE |
| 2314 | prpE | P77495 | FALSE | FALSE | FALSE |
| 2315 | sgcE | P39362 | TRUE | TRUE | TRUE |
| 2316 | ybbW | P75712 | FALSE | FALSE | FALSE |
| 2317 | ftnA | P0A998 | FALSE | FALSE | FALSE |
| 2318 | agaV | P42904 | FALSE | FALSE | FALSE |
| 2319 | tgt | P0A847 | TRUE | TRUE | FALSE |
| 2320 | fur | P0A9A9 | TRUE | TRUE | TRUE |
| 2321 | ridA | P0AF93 | FALSE | MISSING | FALSE |
| 2322 | ydgV | P0DSF5 | FALSE | FALSE | FALSE |
| 2323 | dppD | P0AAG0 | TRUE | FALSE | FALSE |
| 2324 | glnB | P0A9Z1 | FALSE | FALSE | FALSE |
| 2325 | yjiM | P39384 | TRUE | FALSE | FALSE |
| 2326 | yohP | C1P609 | FALSE | MISSING | FALSE |
| 2327 | phnL | P16679 | FALSE | FALSE | FALSE |
| 2328 | torZ | P46923 | FALSE | TRUE | TRUE |
| 2329 | nikA | P33590 | FALSE | FALSE | FALSE |
| 2330 | xdhC | Q46801 | TRUE | FALSE | FALSE |
| 2331 | topA | P06612 | TRUE | TRUE | TRUE |
| 2332 | yadV | P33128 | FALSE | FALSE | FALSE |
| 2333 | ilvB | P08142 | FALSE | FALSE | FALSE |
| 2334 | gss | P0AES0 | TRUE | TRUE | FALSE |
| 2335 | ygcU | Q46911 | FALSE | FALSE | FALSE |
| 2336 | potG | P31134 | FALSE | FALSE | FALSE |
| 2337 | wbbJ | P37750 | FALSE | FALSE | FALSE |
| 2338 | bcr | P28246 | FALSE | MISSING | FALSE |
| 2339 | yccX | P0AB65 | FALSE | FALSE | FALSE |
| 2340 | entH | P0A8Y8 | FALSE | FALSE | FALSE |
| 2341 | yeiH | P62723 | FALSE | FALSE | FALSE |
| 2342 | sdaB | P30744 | FALSE | TRUE | FALSE |
| 2343 | pdxB | P05459 | FALSE | FALSE | FALSE |
| 2344 | yghA | P0AG84 | FALSE | FALSE | FALSE |
| 2345 | rpsT | P0A7U7 | FALSE | FALSE | FALSE |
| 2346 | dkgA | Q46857 | FALSE | TRUE | TRUE |
| 2347 | rluC | P0AA39 | FALSE | FALSE | FALSE |
| 2348 | aroB | P07639 | FALSE | TRUE | FALSE |
| 2349 | rpsM | P0A7S9 | FALSE | FALSE | FALSE |
| 2350 | mlrA | P33358 | FALSE | FALSE | TRUE |
| 2351 | ecpD | P77694 | FALSE | FALSE | FALSE |
| 2352 | entB | P0ADI4 | FALSE | FALSE | FALSE |

Supplementary Table 3: E. coli predicted metal-binding proteome *(continued)*

| # | Gene Name | UniProt ID | MoM Prediction | M3D Prediction | GASS Prediction |
|---|---|---|---|---|---|
| 2353 | ubiU | P45527 | TRUE | FALSE | FALSE |
| 2354 | ydiS | P77337 | FALSE | TRUE | FALSE |
| 2355 | aroE | P15770 | FALSE | FALSE | FALSE |
| 2356 | rutD | P75895 | FALSE | FALSE | TRUE |
| 2357 | yjdN | P16681 | FALSE | FALSE | FALSE |
| 2358 | ygiV | Q46866 | FALSE | FALSE | FALSE |
| 2359 | galU | P0AEP3 | FALSE | FALSE | FALSE |
| 2360 | purF | P0AG16 | FALSE | FALSE | FALSE |
| 2361 | ubiA | P0AGK1 | FALSE | FALSE | FALSE |
| 2362 | ulaD | P39304 | FALSE | FALSE | FALSE |
| 2363 | btuE | P06610 | FALSE | MISSING | TRUE |
| 2364 | norV | Q46877 | TRUE | TRUE | FALSE |
| 2365 | paaG | P77467 | FALSE | FALSE | FALSE |
| 2366 | cusC | P77211 | FALSE | FALSE | FALSE |
| 2367 | uidR | P0ACT6 | FALSE | FALSE | FALSE |
| 2368 | narV | P0AF32 | FALSE | TRUE | FALSE |
| 2369 | sapC | P0AGH5 | FALSE | FALSE | FALSE |
| 2370 | alsK | P32718 | TRUE | TRUE | FALSE |
| 2371 | panZ | P37613 | FALSE | FALSE | FALSE |
| 2372 | ldrB | Q6BF87 | FALSE | FALSE | FALSE |
| 2373 | gshB | P04425 | FALSE | FALSE | TRUE |
| 2374 | ybiU | P75791 | TRUE | TRUE | FALSE |
| 2375 | groEL | P0A6F5 | FALSE | FALSE | FALSE |
| 2376 | fdx | P0A9R4 | TRUE | FALSE | FALSE |
| 2377 | hycF | P16432 | TRUE | TRUE | TRUE |
| 2378 | tauD | P37610 | TRUE | TRUE | TRUE |
| 2379 | pyrB | P0A786 | FALSE | FALSE | FALSE |
| 2380 | prfH | P28369 | FALSE | FALSE | FALSE |
| 2381 | ygcW | P76633 | FALSE | FALSE | FALSE |
| 2382 | waaP | P25741 | FALSE | FALSE | FALSE |
| 2383 | yrbN | C1P618 | FALSE | FALSE | FALSE |
| 2384 | yhfY | P45551 | FALSE | FALSE | FALSE |
| 2385 | cyoB | P0ABI8 | FALSE | TRUE | TRUE |
| 2386 | yjcS | P32717 | TRUE | TRUE | TRUE |
| 2387 | nirB | P08201 | TRUE | TRUE | FALSE |
| 2388 | ompF | P02931 | FALSE | MISSING | FALSE |
| 2389 | trxB | P0A9P4 | FALSE | TRUE | FALSE |
| 2390 | pepD | P15288 | FALSE | TRUE | TRUE |
| 2391 | grcA | P68066 | FALSE | TRUE | FALSE |
| 2392 | yiaW | P0ADK4 | FALSE | FALSE | FALSE |
| 2393 | rlmB | P63177 | FALSE | FALSE | FALSE |
| 2394 | gsiD | P75799 | FALSE | FALSE | FALSE |
| 2395 | pdxJ | P0A794 | FALSE | FALSE | TRUE |
| 2396 | aroL | P0A6E1 | FALSE | FALSE | FALSE |
| 2397 | hcaE | P0ABR5 | TRUE | TRUE | FALSE |
| 2398 | ptsH | P0AA04 | FALSE | FALSE | FALSE |
| 2399 | ybbD | P33669 | FALSE | FALSE | FALSE |
| 2400 | ypeA | P76539 | FALSE | FALSE | FALSE |
| 2401 | folA | P0ABQ4 | FALSE | FALSE | FALSE |
| 2402 | ynaJ | P64445 | FALSE | MISSING | FALSE |
| 2403 | acnB | P36683 | FALSE | FALSE | FALSE |
| 2404 | artJ | P30860 | FALSE | FALSE | FALSE |
| 2405 | glgB | P07762 | FALSE | FALSE | TRUE |
| 2406 | cmoA | P76290 | FALSE | FALSE | FALSE |
| 2407 | agaD | P42911 | FALSE | MISSING | FALSE |
| 2408 | zinT | P76344 | TRUE | TRUE | FALSE |

Supplementary Table 3: E. coli predicted metal-binding proteome *(continued)*

| # | Gene Name | UniProt ID | MoM Prediction | M3D Prediction | GASS Prediction |
|---|---|---|---|---|---|
| 2409 | argD | P18335 | TRUE | FALSE | FALSE |
| 2410 | lysO | P75826 | FALSE | MISSING | FALSE |
| 2411 | argG | P0A6E4 | FALSE | TRUE | FALSE |
| 2412 | hiuH | P76341 | FALSE | FALSE | FALSE |
| 2413 | sufC | P77499 | FALSE | FALSE | FALSE |
| 2414 | queC | P77756 | TRUE | TRUE | FALSE |
| 2415 | yqhD | Q46856 | TRUE | TRUE | TRUE |
| 2416 | ydcR | P77730 | FALSE | FALSE | TRUE |
| 2417 | uxuA | P24215 | TRUE | TRUE | TRUE |
| 2418 | lsrR | P76141 | FALSE | FALSE | FALSE |
| 2419 | thrB | P00547 | FALSE | FALSE | FALSE |
| 2420 | yphF | P77269 | FALSE | FALSE | FALSE |
| 2421 | recX | P33596 | FALSE | FALSE | FALSE |
| 2422 | serS | P0A8L1 | FALSE | FALSE | FALSE |
| 2423 | btsT | P39396 | FALSE | TRUE | FALSE |
| 2424 | cheY | P0AE67 | FALSE | FALSE | FALSE |
| 2425 | mglB | P0AEE5 | FALSE | FALSE | FALSE |
| 2426 | preT | P76440 | FALSE | FALSE | FALSE |
| 2427 | rluD | P33643 | FALSE | FALSE | FALSE |
| 2428 | ycjW | P77615 | FALSE | FALSE | FALSE |
| 2429 | frdD | P0A8Q3 | TRUE | FALSE | TRUE |
| 2430 | hemY | P0ACB7 | FALSE | FALSE | FALSE |
| 2431 | yicC | P23839 | FALSE | FALSE | FALSE |
| 2432 | putP | P07117 | FALSE | FALSE | FALSE |
| 2433 | pckA | P22259 | FALSE | TRUE | FALSE |
| 2434 | tufA | P0CE47 | FALSE | FALSE | FALSE |
| 2435 | cysM | P16703 | FALSE | FALSE | FALSE |
| 2436 | idnD | P39346 | TRUE | TRUE | FALSE |
| 2437 | ribF | P0AG40 | FALSE | FALSE | FALSE |
| 2438 | appA | P07102 | FALSE | FALSE | FALSE |
| 2439 | eutT | P65643 | FALSE | FALSE | FALSE |
| 2440 | chbB | P69795 | FALSE | FALSE | FALSE |
| 2441 | prs | P0A717 | FALSE | FALSE | FALSE |
| 2442 | hyaE | P19931 | FALSE | FALSE | FALSE |
| 2443 | cydB | P0ABK2 | FALSE | FALSE | FALSE |
| 2444 | yfcD | P65556 | FALSE | FALSE | FALSE |
| 2445 | yfbO | P76485 | FALSE | FALSE | FALSE |
| 2446 | gppA | P25552 | FALSE | FALSE | FALSE |
| 2447 | allE | P75713 | FALSE | TRUE | FALSE |
| 2448 | rffH | P61887 | FALSE | FALSE | FALSE |
| 2449 | tusB | P45530 | FALSE | FALSE | FALSE |
| 2450 | rplD | P60723 | FALSE | FALSE | FALSE |
| 2451 | ispD | Q46893 | FALSE | FALSE | FALSE |
| 2452 | dusB | P0ABT5 | FALSE | FALSE | FALSE |
| 2453 | plsC | P26647 | FALSE | FALSE | FALSE |
| 2454 | mdtP | P32714 | FALSE | TRUE | FALSE |
| 2455 | rimI | P0A944 | FALSE | FALSE | FALSE |
| 2456 | yggP | P52048 | FALSE | TRUE | TRUE |
| 2457 | waaA | P0AC75 | FALSE | FALSE | FALSE |
| 2458 | araF | P02924 | FALSE | FALSE | FALSE |
| 2459 | clcA | P37019 | FALSE | FALSE | FALSE |
| 2460 | yfaE | P0ABW3 | FALSE | FALSE | FALSE |
| 2461 | yddM | P67699 | FALSE | FALSE | FALSE |
| 2462 | yphE | P77509 | FALSE | FALSE | FALSE |
| 2463 | yejE | P33915 | FALSE | MISSING | FALSE |
| 2464 | wcaF | P0ACD2 | FALSE | FALSE | FALSE |

Supplementary Table 3: E. coli predicted metal-binding proteome *(continued)*

| # | Gene Name | UniProt ID | MoM Prediction | M3D Prediction | GASS Prediction |
|---|---|---|---|---|---|
| 2465 | ilvC | P05793 | TRUE | FALSE | FALSE |
| 2466 | rpsH | P0A7W7 | FALSE | MISSING | FALSE |
| 2467 | mutT | P08337 | FALSE | FALSE | FALSE |
| 2468 | apbE | P0AB85 | FALSE | FALSE | FALSE |
| 2469 | aroA | P0A6D3 | FALSE | FALSE | FALSE |
| 2470 | menE | P37353 | FALSE | FALSE | FALSE |
| 2471 | uppP | P60932 | FALSE | FALSE | FALSE |
| 2472 | galE | P09147 | FALSE | FALSE | TRUE |
| 2473 | rlhA | P76104 | FALSE | FALSE | FALSE |
| 2474 | inaA | P27294 | FALSE | TRUE | TRUE |
| 2475 | kbp | P0ADE6 | FALSE | FALSE | FALSE |
| 2476 | cheR | P07364 | FALSE | FALSE | FALSE |
| 2477 | yfeX | P76536 | FALSE | FALSE | FALSE |
| 2478 | gltD | P09832 | FALSE | FALSE | FALSE |
| 2479 | nepI | P0ADL1 | TRUE | MISSING | FALSE |
| 2480 | rnb | P30850 | FALSE | FALSE | FALSE |
| 2481 | bglA | Q46829 | FALSE | FALSE | FALSE |
| 2482 | trmO | P28634 | FALSE | FALSE | FALSE |
| 2483 | gatD | P0A9S3 | TRUE | TRUE | FALSE |
| 2484 | iap | P10423 | FALSE | TRUE | FALSE |
| 2485 | fsaA | P78055 | FALSE | FALSE | FALSE |
| 2486 | bcsZ | P37651 | FALSE | FALSE | FALSE |
| 2487 | yafE | P30866 | TRUE | TRUE | FALSE |
| 2488 | hyaB | P0ACD8 | TRUE | TRUE | FALSE |
| 2489 | sbp | P0AG78 | FALSE | FALSE | FALSE |
| 2490 | dgoK | P31459 | FALSE | TRUE | FALSE |
| 2491 | ddpD | P77268 | TRUE | FALSE | FALSE |
| 2492 | sapA | Q47622 | FALSE | FALSE | FALSE |
| 2493 | nanE | P0A761 | FALSE | FALSE | TRUE |
| 2494 | malZ | P21517 | FALSE | FALSE | FALSE |
| 2495 | dam | P0AEE8 | TRUE | FALSE | FALSE |
| 2496 | envY | P10805 | FALSE | FALSE | FALSE |
| 2497 | yjbR | P0AF50 | FALSE | FALSE | FALSE |
| 2498 | ynjB | P76223 | FALSE | TRUE | FALSE |
| 2499 | croE | P75975 | FALSE | MISSING | FALSE |
| 2500 | usg | P08390 | FALSE | FALSE | FALSE |
| 2501 | tehB | P25397 | FALSE | FALSE | FALSE |
| 2502 | thiD | P76422 | FALSE | FALSE | FALSE |
| 2503 | nlpC | P23898 | FALSE | FALSE | FALSE |
| 2504 | garK | P23524 | FALSE | FALSE | FALSE |
| 2505 | yegD | P36928 | FALSE | FALSE | FALSE |
| 2506 | yoaJ | C1P603 | FALSE | MISSING | FALSE |
| 2507 | ddlA | P0A6J8 | FALSE | FALSE | FALSE |
| 2508 | dmlR | P76250 | FALSE | FALSE | FALSE |
| 2509 | nrfA | P0ABK9 | FALSE | TRUE | TRUE |
| 2510 | dapA | P0A6L2 | FALSE | FALSE | FALSE |
| 2511 | yfdI | P76507 | TRUE | FALSE | FALSE |
| 2512 | nuoN | P0AFF0 | FALSE | FALSE | FALSE |
| 2513 | mqsR | Q46865 | FALSE | FALSE | FALSE |
| 2514 | ycjG | P51981 | FALSE | FALSE | FALSE |
| 2515 | fhuE | P16869 | FALSE | FALSE | FALSE |
| 2516 | pppA | Q46836 | TRUE | TRUE | FALSE |
| 2517 | csdE | P0AGF2 | FALSE | FALSE | FALSE |
| 2518 | ykgO | Q2EEQ2 | TRUE | MISSING | FALSE |
| 2519 | xylA | P00944 | TRUE | TRUE | FALSE |
| 2520 | ycfT | P75955 | FALSE | FALSE | FALSE |

Supplementary Table 3: E. coli predicted metal-binding proteome *(continued)*

| # | Gene Name | UniProt ID | MoM Prediction | M3D Prediction | GASS Prediction |
|---|---|---|---|---|---|
| 2521 | dgoR | P31460 | TRUE | TRUE | TRUE |
| 2522 | eutM | P0ABF4 | FALSE | MISSING | FALSE |
| 2523 | insA1 | P0CF07 | TRUE | TRUE | FALSE |
| 2524 | rplY | P68919 | FALSE | FALSE | FALSE |
| 2525 | ompT | P09169 | FALSE | FALSE | FALSE |
| 2526 | gnd | P00350 | FALSE | FALSE | FALSE |
| 2527 | ptsN | P69829 | FALSE | FALSE | FALSE |
| 2528 | syd | P0A8U0 | FALSE | FALSE | FALSE |
| 2529 | rng | P0A9J0 | TRUE | FALSE | FALSE |
| 2530 | hypA | P0A700 | TRUE | TRUE | TRUE |
| 2531 | citF | P75726 | TRUE | FALSE | FALSE |
| 2532 | maeB | P76558 | FALSE | FALSE | FALSE |
| 2533 | yohJ | P60632 | FALSE | MISSING | FALSE |
| 2534 | galR | P03024 | FALSE | TRUE | FALSE |
| 2535 | tsf | P0A6P1 | FALSE | FALSE | FALSE |
| 2536 | ygiD | P24197 | TRUE | TRUE | FALSE |
| 2537 | ydfC | P21418 | FALSE | MISSING | FALSE |
| 2538 | fucO | P0A9S1 | TRUE | TRUE | TRUE |
| 2539 | yjjW | P39409 | TRUE | FALSE | FALSE |
| 2540 | casB | P76632 | FALSE | FALSE | FALSE |
| 2541 | kptA | P39380 | FALSE | FALSE | FALSE |
| 2542 | nanX | P39352 | FALSE | MISSING | FALSE |
| 2543 | mazE | P0AE72 | FALSE | FALSE | FALSE |
| 2544 | yggF | P21437 | FALSE | FALSE | FALSE |
| 2545 | ydjG | P77256 | FALSE | FALSE | FALSE |
| 2546 | dmlA | P76251 | FALSE | FALSE | FALSE |
| 2547 | folE | P0A6T5 | TRUE | TRUE | TRUE |
| 2548 | iscA | P0AAC8 | FALSE | FALSE | FALSE |
| 2549 | bcp | P0AE52 | FALSE | FALSE | FALSE |
| 2550 | yjfP | P39298 | FALSE | FALSE | FALSE |
| 2551 | yggR | P52052 | FALSE | FALSE | FALSE |
| 2552 | frvB | P32154 | FALSE | FALSE | FALSE |
| 2553 | accC | P24182 | FALSE | FALSE | TRUE |
| 2554 | hflD | P25746 | FALSE | FALSE | TRUE |
| 2555 | deoA | P07650 | FALSE | FALSE | FALSE |
| 2556 | ynfE | P77374 | FALSE | FALSE | FALSE |
| 2557 | fadD | P69451 | FALSE | TRUE | FALSE |
| 2558 | ccp | P37197 | FALSE | FALSE | FALSE |
| 2559 | dnaN | P0A988 | FALSE | FALSE | FALSE |
| 2560 | lsrB | P76142 | FALSE | FALSE | FALSE |
| 2561 | codB | P0AA82 | FALSE | FALSE | FALSE |
| 2562 | gntK | P46859 | FALSE | FALSE | TRUE |
| 2563 | rplV | P61175 | FALSE | FALSE | FALSE |
| 2564 | gpmA | P62707 | FALSE | TRUE | TRUE |
| 2565 | cmtB | P69824 | FALSE | FALSE | FALSE |
| 2566 | yhcA | P28722 | FALSE | FALSE | TRUE |
| 2567 | rnk | P0AFW4 | FALSE | FALSE | FALSE |
| 2568 | fumC | P05042 | FALSE | FALSE | TRUE |
| 2569 | yecM | P52007 | FALSE | TRUE | FALSE |
| 2570 | thpR | P37025 | FALSE | FALSE | FALSE |
| 2571 | ybcL | P77368 | FALSE | FALSE | FALSE |
| 2572 | lptC | P0ADV9 | FALSE | FALSE | FALSE |
| 2573 | sseA | P31142 | FALSE | FALSE | FALSE |
| 2574 | ytfK | P0ADE2 | FALSE | MISSING | FALSE |

Supplementary Table 4: TPP Hits

| # | Gene Name | UniProt ID | Bound Metal | Effect | Adjusted p-value | dTm |
|---|---|---|---|---|---|---|
| 1 | cysS | P21888 | zinc | Destabilised | 0.0000023 | -4.4388 |
| 2 | tilS | P52097 | - | Destabilised | 0.0000076 | -6.1481 |
| 3 | fdhE | P13024 | iron | Destabilised | 0.0000096 | -7.3603 |
| 4 | hisI | P06989 | - | Destabilised | 0.0000098 | -13.0904 |
| 5 | idi | Q46822 | manganese, magnesium, zinc | Destabilised | 0.0000098 | -10.1409 |
| 6 | moeB | P12282 | zinc | Destabilised | 0.0000099 | -9.6083 |
| 7 | nagC | P0AF20 | - | Destabilised | 0.0000175 | -6.0215 |
| 8 | fbp | P0A993 | magnesium | Destabilised | 0.0000192 | -1.6098 |
| 9 | gloB | P0AC84 | zinc | Destabilised | 0.0000271 | -14.2231 |
| 10 | lpxC | P0A725 | zinc, iron | Destabilised | 0.0000278 | -5.6840 |
| 11 | rsgA | P39286 | zinc | Destabilised | 0.0000278 | -3.8464 |
| 12 | purA | P0A7D4 | magnesium | Destabilised | 0.0000369 | -2.7300 |
| 13 | corC | P0AE78 | cobalt, magnesium | Destabilised | 0.0000380 | -6.3482 |
| 14 | ycfH | P0AFQ7 | cobalt, manganese, nickel | Destabilised | 0.0000454 | -9.9221 |
| 15 | frmA | P25437 | zinc | Destabilised | 0.0000521 | -4.2136 |
| 16 | tdh | P07913 | zinc, cobalt, cadmium, iron, manganese | Destabilised | 0.0000690 | -6.0410 |
| 17 | dcp | P24171 | zinc, calcium | Stabilised | 0.0000762 | 2.8428 |
| 18 | yjbQ | P0AF48 | - | Destabilised | 0.0000779 | -9.3933 |
| 19 | hisB | P06987 | magnesium, zinc | Destabilised | 0.0000962 | -6.7220 |
| 20 | pepP | P15034 | manganese | Destabilised | 0.0000989 | -8.4662 |
| 21 | cobB | P75960 | zinc | Destabilised | 0.0001216 | -3.9384 |
| 22 | orn | P0A784 | zinc | Destabilised | 0.0001519 | -3.7389 |
| 23 | selB | P14081 | - | Stabilised | 0.0001519 | 2.7691 |
| 24 | mtfA | P76346 | zinc | Destabilised | 0.0001519 | -6.7808 |
| 25 | yajD | P0AAQ2 | zinc | Destabilised | 0.0001691 | NA |
| 26 | argE | P23908 | zinc, cobalt | Destabilised | 0.0002841 | -5.9284 |
| 27 | pckA | P22259 | calcium, manganese, magnesium | Destabilised | 0.0003536 | -2.2213 |
| 28 | cca | P06961 | magnesium, nickel | Destabilised | 0.0004232 | -6.9505 |
| 29 | mmuM | Q47690 | zinc | Destabilised | 0.0004232 | -5.2781 |
| 30 | pepN | P04825 | zinc | Destabilised | 0.0004949 | -2.2427 |
| 31 | map | P0AE18 | cobalt, zinc, manganese, iron, sodium | Destabilised | 0.0004949 | -2.3987 |
| 32 | dapE | P0AED7 | zinc, cobalt | Destabilised | 0.0004949 | -9.6163 |
| 33 | glmM | P31120 | magnesium | Destabilised | 0.0005462 | -5.8614 |
| 34 | mqsA | Q46864 | zinc | Destabilised | 0.0005462 | -8.1971 |
| 35 | cpdA | P0AEW4 | iron | Destabilised | 0.0007248 | -10.0286 |
| 36 | eptB | P37661 | calcium | Destabilised | 0.0010161 | -3.7897 |
| 37 | tadA | P68398 | zinc | Destabilised | 0.0010161 | -6.6022 |
| 38 | pepT | P29745 | zinc | Destabilised | 0.0012244 | -9.7083 |
| 39 | phnP | P16692 | zinc, manganese | Destabilised | 0.0012517 | -9.1559 |
| 40 | tgt | P0A847 | zinc | Stabilised | 0.0013195 | 3.9224 |
| 41 | gatZ | P0C8J8 | - | Destabilised | 0.0014025 | -4.6571 |
| 42 | ybeL | P0AAT9 | - | Destabilised | 0.0017002 | NA |
| 43 | pepQ | P21165 | manganese | Destabilised | 0.0019051 | -8.9666 |
| 44 | prlC | P27298 | zinc | Destabilised | 0.0020501 | -2.1297 |
| 45 | aroG | P0AB91 | - | Destabilised | 0.0020936 | -1.7984 |
| 46 | bfr | P0ABD3 | iron | Destabilised | 0.0020936 | NA |
| 47 | mpaA | P0ACV6 | zinc | Destabilised | 0.0020936 | -9.3585 |
| 48 | pgm | P36938 | magnesium | Destabilised | 0.0020936 | -3.6970 |
| 49 | thrS | P0A8M3 | zinc | Destabilised | 0.0028703 | -2.4605 |
| 50 | rihA | P41409 | calcium | Destabilised | 0.0028945 | -22.1970 |
| 51 | mazG | P0AEY3 | magnesium | Destabilised | 0.0030821 | -3.9861 |
| 52 | metQ | P28635 | - | Destabilised | 0.0033889 | NA |
| 53 | zur | P0AC51 | zinc | Destabilised | 0.0036578 | -10.6161 |
| 54 | dgoR | P31460 | zinc | Destabilised | 0.0046165 | -4.3864 |
| 55 | yfcE | P67095 | manganese | Destabilised | 0.0046943 | -11.1178 |
| 56 | uxuR | P39161 | - | Destabilised | 0.0047685 | -4.7263 |

Supplementary Table 4: TPP Hits *(continued)*

| # | Gene Name | UniProt ID | Bound Metal | Effect | Adjusted p-value | dTm |
|---|---|---|---|---|---|---|
| 57 | obgE | P42641 | magnesium | Stabilised | 0.0048925 | 1.3382 |
| 58 | gmhA | P63224 | zinc | Stabilised | 0.0050454 | 2.1733 |
| 59 | yfeX | P76536 | iron | Stabilised | 0.0051422 | 0.5795 |
| 60 | glnD | P27249 | magnesium | Destabilised | 0.0051746 | -3.5438 |
| 61 | bglX | P33363 | - | Destabilised | 0.0051746 | -1.6013 |
| 62 | rbn | P0A8V0 | zinc | Destabilised | 0.0051747 | -9.2487 |
| 63 | glgA | P0A6U8 | - | Destabilised | 0.0055544 | -1.6838 |
| 64 | phnO | P16691 | divalent metal cation | Destabilised | 0.0057081 | -2.5284 |
| 65 | nrdR | P0A8D0 | zinc | Destabilised | 0.0058776 | -6.1495 |
| 66 | yegU | P76418 | - | Destabilised | 0.0064935 | -2.9541 |
| 67 | wrbA | P0A8G6 | - | Destabilised | 0.0068248 | -3.7145 |
| 68 | exuR | P0ACL2 | - | Destabilised | 0.0070431 | -4.3283 |
| 69 | ftnA | P0A998 | iron | Destabilised | 0.0083474 | NA |
| 70 | ndk | P0A763 | magnesium | Stabilised | 0.0084548 | 2.6841 |
| 71 | ligA | P15042 | zinc, magnesium | Destabilised | 0.0106686 | -3.6707 |
| 72 | ycdX | P75914 | zinc | Destabilised | 0.0109670 | NA |
| 73 | rng | P0A9J0 | magnesium | Destabilised | 0.0110868 | -0.5236 |
| 74 | tyrB | P04693 | - | Destabilised | 0.0120402 | -2.0430 |
| 75 | pepD | P15288 | zinc, cobalt | Destabilised | 0.0120402 | -2.8866 |
| 76 | galT | P09148 | zinc, iron | Destabilised | 0.0124756 | -3.5575 |
| 77 | metE | P25665 | zinc | Destabilised | 0.0151077 | -0.9735 |
| 78 | yhbS | P63417 | - | Stabilised | 0.0166405 | 1.6912 |
| 79 | leuA | P09151 | manganese | Destabilised | 0.0202881 | -1.3717 |
| 80 | ycaR | P0AAZ7 | - | Destabilised | 0.0214624 | NA |
| 81 | ycaO | P75838 | magnesium | Stabilised | 0.0214624 | 0.9458 |
| 82 | dnaX | P06710 | zinc | Destabilised | 0.0250662 | -2.8011 |
| 83 | nadR | P27278 | magnesium | Destabilised | 0.0268028 | -2.0829 |
| 84 | prfB | P07012 | - | Stabilised | 0.0272865 | 2.8088 |
| 85 | cytR | P0ACN7 | - | Destabilised | 0.0274708 | -7.9955 |
| 86 | gatY | P0C8J6 | zinc | Destabilised | 0.0274708 | -2.4402 |
| 87 | polA | P00582 | - | Destabilised | 0.0286285 | -1.1874 |
| 88 | pyrG | P0A7E5 | magnesium | Destabilised | 0.0293541 | -1.1339 |
| 89 | add | P22333 | zinc | Destabilised | 0.0305079 | -4.1848 |
| 90 | rnb | P30850 | magnesium | Stabilised | 0.0305079 | 1.6595 |
| 91 | glk | P0A6V8 | - | Stabilised | 0.0306836 | 0.9756 |
| 92 | hcxA | P45579 | zinc | Destabilised | 0.0339901 | -2.6646 |
| 93 | leuS | P07813 | - | Destabilised | 0.0389376 | -1.0344 |
| 94 | yrdA | P0A9W9 | zinc | Stabilised | 0.0448373 | 3.0755 |
| 95 | rlmG | P42596 | - | Stabilised | 0.0452008 | 0.6569 |

Supplementary Table 5: Hit Overlaps

| # | Gene Name | UniProt ID | Bound Metal | LiP | TPP | SEC | <10 Å to active site |
|---|---|---|---|---|---|---|---|
| 1 | ycfH | P0AFQ7 | cobalt, manganese, nickel | TRUE | TRUE | TRUE | FALSE |
| 2 | add | P22333 | zinc | TRUE | TRUE | FALSE | TRUE |
| 3 | lpxC | P0A725 | zinc, iron | TRUE | TRUE | FALSE | TRUE |
| 4 | metE | P25665 | zinc | TRUE | TRUE | FALSE | TRUE |
| 5 | ndk | P0A763 | magnesium | TRUE | TRUE | FALSE | TRUE |
| 6 | pepD | P15288 | zinc, cobalt | TRUE | TRUE | FALSE | TRUE |
| 7 | pepN | P04825 | zinc | TRUE | TRUE | FALSE | TRUE |
| 8 | pepT | P29745 | zinc | TRUE | TRUE | FALSE | TRUE |
| 9 | pgm | P36938 | magnesium | TRUE | TRUE | FALSE | TRUE |
| 10 | yfeX | P76536 | iron | TRUE | TRUE | FALSE | TRUE |
| 11 | argE | P23908 | zinc, cobalt | TRUE | TRUE | FALSE | FALSE |
| 12 | aroG | P0AB91 | - | TRUE | TRUE | FALSE | FALSE |
| 13 | bfr | P0ABD3 | iron | TRUE | TRUE | FALSE | FALSE |
| 14 | cysS | P21888 | zinc | TRUE | TRUE | FALSE | FALSE |
| 15 | dnaX | P06710 | zinc | TRUE | TRUE | FALSE | FALSE |
| 16 | gatZ | P0C8J8 | - | TRUE | TRUE | FALSE | FALSE |
| 17 | gloB | P0AC84 | zinc | TRUE | TRUE | FALSE | FALSE |
| 18 | gmhA | P63224 | zinc | TRUE | TRUE | FALSE | FALSE |
| 19 | hcxA | P45579 | zinc | TRUE | TRUE | FALSE | FALSE |
| 20 | hisB | P06987 | magnesium, zinc | TRUE | TRUE | FALSE | FALSE |
| 21 | hisI | P06989 | - | TRUE | TRUE | FALSE | FALSE |
| 22 | leuA | P09151 | manganese | TRUE | TRUE | FALSE | FALSE |
| 23 | ligA | P15042 | zinc, magnesium | TRUE | TRUE | FALSE | FALSE |
| 24 | map | P0AE18 | cobalt, zinc, manganese, iron, sodium | TRUE | TRUE | FALSE | FALSE |
| 25 | mmuM | Q47690 | zinc | TRUE | TRUE | FALSE | FALSE |
| 26 | nrdR | P0A8D0 | zinc | TRUE | TRUE | FALSE | FALSE |
| 27 | pepP | P15034 | manganese | TRUE | TRUE | FALSE | FALSE |
| 28 | pepQ | P21165 | manganese | TRUE | TRUE | FALSE | FALSE |
| 29 | purA | P0A7D4 | magnesium | TRUE | TRUE | FALSE | FALSE |
| 30 | pyrG | P0A7E5 | magnesium | TRUE | TRUE | FALSE | FALSE |
| 31 | tyrB | P04693 | - | TRUE | TRUE | FALSE | FALSE |
| 32 | wrbA | P0A8G6 | - | TRUE | TRUE | FALSE | FALSE |
| 33 | yajD | P0AAQ2 | zinc | TRUE | TRUE | FALSE | FALSE |
| 34 | ybeL | P0AAT9 | - | TRUE | TRUE | FALSE | FALSE |
| 35 | ycdX | P75914 | zinc | TRUE | TRUE | FALSE | FALSE |
| 36 | yrdA | P0A9W9 | zinc | TRUE | TRUE | FALSE | FALSE |
| 37 | zur | P0AC51 | zinc | TRUE | TRUE | FALSE | FALSE |
| 38 | bolA | P0ABE2 | - | TRUE | FALSE | TRUE | FALSE |
| 39 | cmoA | P76290 | - | TRUE | FALSE | TRUE | FALSE |
| 40 | metK | P0A817 | magnesium, potassium, manganese, cobalt | TRUE | FALSE | TRUE | FALSE |
| 41 | rlmI | P75876 | - | TRUE | FALSE | TRUE | FALSE |
| 42 | rplB | P60422 | zinc | TRUE | FALSE | TRUE | FALSE |
| 43 | rplC | P60438 | - | TRUE | FALSE | TRUE | FALSE |
| 44 | rplD | P60723 | - | TRUE | FALSE | TRUE | FALSE |
| 45 | rplE | P62399 | - | TRUE | FALSE | TRUE | FALSE |
| 46 | rplF | P0AG55 | - | TRUE | FALSE | TRUE | FALSE |
| 47 | rplJ | P0A7J3 | - | TRUE | FALSE | TRUE | FALSE |
| 48 | rplM | P0AA10 | zinc | TRUE | FALSE | TRUE | FALSE |
| 49 | rplN | P0ADY3 | - | TRUE | FALSE | TRUE | FALSE |
| 50 | rplO | P02413 | - | TRUE | FALSE | TRUE | FALSE |
| 51 | rplQ | P0AG44 | - | TRUE | FALSE | TRUE | FALSE |
| 52 | rplR | P0C018 | - | TRUE | FALSE | TRUE | FALSE |
| 53 | rplV | P61175 | - | TRUE | FALSE | TRUE | FALSE |
| 54 | rplW | P0ADZ0 | - | TRUE | FALSE | TRUE | FALSE |

Supplementary Table 5: Hit Overlaps *(continued)*

| # | Gene Name | UniProt ID | Bound Metal | LiP | TPP | SEC | <10 Å to active site |
|---|---|---|---|---|---|---|---|
| 55 | rplX | P60624 | - | TRUE | FALSE | TRUE | FALSE |
| 56 | rplY | P68919 | - | TRUE | FALSE | TRUE | FALSE |
| 57 | rpmA | P0A7L8 | - | TRUE | FALSE | TRUE | FALSE |
| 58 | rpmB | P0A7M2 | - | TRUE | FALSE | TRUE | FALSE |
| 59 | rpmD | P0AG51 | - | TRUE | FALSE | TRUE | FALSE |
| 60 | rpmF | P0A7N4 | - | TRUE | FALSE | TRUE | FALSE |
| 61 | rpmG | P0A7N9 | - | TRUE | FALSE | TRUE | FALSE |
| 62 | rpsC | P0A7V3 | - | TRUE | FALSE | TRUE | FALSE |
| 63 | rpsH | P0A7W7 | - | TRUE | FALSE | TRUE | FALSE |
| 64 | rpsJ | P0A7R5 | - | TRUE | FALSE | TRUE | FALSE |
| 65 | rpsL | P0A7S3 | - | TRUE | FALSE | TRUE | FALSE |
| 66 | rpsP | P0A7T3 | - | TRUE | FALSE | TRUE | FALSE |
| 67 | rpsU | P68679 | - | TRUE | FALSE | TRUE | FALSE |
| 68 | rsfS | P0AAT6 | - | TRUE | FALSE | TRUE | FALSE |
| 69 | thiI | P77718 | iron | TRUE | FALSE | TRUE | FALSE |
| 70 | aceB | P08997 | - | TRUE | FALSE | FALSE | TRUE |
| 71 | amyA | P26612 | calcium, sodium | TRUE | FALSE | FALSE | TRUE |
| 72 | asnB | P22106 | - | TRUE | FALSE | FALSE | TRUE |
| 73 | bcp | P0AE52 | - | TRUE | FALSE | FALSE | TRUE |
| 74 | deoC | P0A6L0 | - | TRUE | FALSE | FALSE | TRUE |
| 75 | eno | P0A6P9 | magnesium | TRUE | FALSE | FALSE | TRUE |
| 76 | fbaA | P0AB71 | zinc | TRUE | FALSE | FALSE | TRUE |
| 77 | fruB | P69811 | - | TRUE | FALSE | FALSE | TRUE |
| 78 | fruK | P0AEW9 | magnesium | TRUE | FALSE | FALSE | TRUE |
| 79 | gapA | P0A9B2 | - | TRUE | FALSE | FALSE | TRUE |
| 80 | gatB | P37188 | - | TRUE | FALSE | FALSE | TRUE |
| 81 | glmS | P17169 | - | TRUE | FALSE | FALSE | TRUE |
| 82 | gloA | P0AC81 | nickel | TRUE | FALSE | FALSE | TRUE |
| 83 | gltB | P09831 | iron | TRUE | FALSE | FALSE | TRUE |
| 84 | gpmI | P37689 | manganese | TRUE | FALSE | FALSE | TRUE |
| 85 | guaA | P04079 | - | TRUE | FALSE | FALSE | TRUE |
| 86 | guaC | P60560 | potassium | TRUE | FALSE | FALSE | TRUE |
| 87 | hemB | P0ACB2 | zinc, magnesium | TRUE | FALSE | FALSE | TRUE |
| 88 | hisD | P06988 | zinc, manganese | TRUE | FALSE | FALSE | TRUE |
| 89 | lysA | P00861 | - | TRUE | FALSE | FALSE | TRUE |
| 90 | mdh | P61889 | - | TRUE | FALSE | FALSE | TRUE |
| 91 | metAS | P07623 | - | TRUE | FALSE | FALSE | TRUE |
| 92 | msrB | P0A746 | zinc, iron | TRUE | FALSE | FALSE | TRUE |
| 93 | pepA | P68767 | manganese | TRUE | FALSE | FALSE | TRUE |
| 94 | ptsI | P08839 | magnesium | TRUE | FALSE | FALSE | TRUE |
| 95 | speD | P0A7F6 | magnesium | TRUE | FALSE | FALSE | TRUE |
| 96 | tpx | P0A862 | - | TRUE | FALSE | FALSE | TRUE |
| 97 | trpGD | P00904 | magnesium | TRUE | FALSE | FALSE | TRUE |
| 98 | yccX | P0AB65 | - | TRUE | FALSE | FALSE | TRUE |
| 99 | aas | P31119 | - | TRUE | FALSE | FALSE | FALSE |
| 100 | accD | P0A9Q5 | zinc | TRUE | FALSE | FALSE | FALSE |
| 101 | aceA | P0A9G6 | magnesium, manganese | TRUE | FALSE | FALSE | FALSE |
| 102 | aceE | P0AFG8 | magnesium | TRUE | FALSE | FALSE | FALSE |
| 103 | acnB | P36683 | iron | TRUE | FALSE | FALSE | FALSE |
| 104 | acuI | P26646 | - | TRUE | FALSE | FALSE | FALSE |
| 105 | ade | P31441 | manganese, iron | TRUE | FALSE | FALSE | FALSE |
| 106 | adhE | P0A9Q7 | iron | TRUE | FALSE | FALSE | FALSE |
| 107 | ahpC | P0AE08 | - | TRUE | FALSE | FALSE | FALSE |
| 108 | ahpF | P35340 | - | TRUE | FALSE | FALSE | FALSE |
| 109 | alaS | P00957 | zinc | TRUE | FALSE | FALSE | FALSE |
| 110 | argB | P0A6C8 | - | TRUE | FALSE | FALSE | FALSE |

Supplementary Table 5: Hit Overlaps *(continued)*

| # | Gene Name | UniProt ID | Bound Metal | LiP | TPP | SEC | <10 Å to active site |
|---|---|---|---|---|---|---|---|
| 111 | argG | P0A6E4 | - | TRUE | FALSE | FALSE | FALSE |
| 112 | aroB | P07639 | zinc, cobalt | TRUE | FALSE | FALSE | FALSE |
| 113 | aroH | P00887 | - | TRUE | FALSE | FALSE | FALSE |
| 114 | asnS | P0A8M0 | - | TRUE | FALSE | FALSE | FALSE |
| 115 | aspS | P21889 | - | TRUE | FALSE | FALSE | FALSE |
| 116 | atpC | P0A6E6 | - | TRUE | FALSE | FALSE | FALSE |
| 117 | atpD | P0ABB4 | - | TRUE | FALSE | FALSE | FALSE |
| 118 | avtA | P09053 | - | TRUE | FALSE | FALSE | FALSE |
| 119 | bipA | P0DTT0 | - | TRUE | FALSE | FALSE | FALSE |
| 120 | btuE | P06610 | - | TRUE | FALSE | FALSE | FALSE |
| 121 | can | P61517 | zinc | TRUE | FALSE | FALSE | FALSE |
| 122 | carA | P0A6F1 | - | TRUE | FALSE | FALSE | FALSE |
| 123 | carB | P00968 | magnesium, manganese | TRUE | FALSE | FALSE | FALSE |
| 124 | cbpM | P63264 | - | TRUE | FALSE | FALSE | FALSE |
| 125 | cfa | P0A9H7 | - | TRUE | FALSE | FALSE | FALSE |
| 126 | clpB | P63284 | - | TRUE | FALSE | FALSE | FALSE |
| 127 | copA | Q59385 | copper, magnesium | TRUE | FALSE | FALSE | FALSE |
| 128 | cpdB | P08331 | divalent metal cation | TRUE | FALSE | FALSE | FALSE |
| 129 | cysK | P0ABK5 | - | TRUE | FALSE | FALSE | FALSE |
| 130 | degP | P0C0V0 | - | TRUE | FALSE | FALSE | FALSE |
| 131 | degQ | P39099 | - | TRUE | FALSE | FALSE | FALSE |
| 132 | deoB | P0A6K6 | manganese, cobalt, magnesium | TRUE | FALSE | FALSE | FALSE |
| 133 | dhaL | P76014 | magnesium | TRUE | FALSE | FALSE | FALSE |
| 134 | dksA | P0ABS1 | zinc | TRUE | FALSE | FALSE | FALSE |
| 135 | dmsA | P18775 | iron, molybdenum | TRUE | FALSE | FALSE | FALSE |
| 136 | dnaJ | P08622 | zinc | TRUE | FALSE | FALSE | FALSE |
| 137 | dosC | P0AA89 | iron, magnesium | TRUE | FALSE | FALSE | FALSE |
| 138 | dsbC | P0AEG6 | - | TRUE | FALSE | FALSE | FALSE |
| 139 | efeO | P0AB24 | - | TRUE | FALSE | FALSE | FALSE |
| 140 | erpA | P0ACC3 | iron | TRUE | FALSE | FALSE | FALSE |
| 141 | fabI | P0AEK4 | - | TRUE | FALSE | FALSE | FALSE |
| 142 | feoC | P64638 | iron | TRUE | FALSE | FALSE | FALSE |
| 143 | folP | P0AC13 | magnesium | TRUE | FALSE | FALSE | FALSE |
| 144 | fre | P0AEN1 | iron | TRUE | FALSE | FALSE | FALSE |
| 145 | frsA | P04335 | - | TRUE | FALSE | FALSE | FALSE |
| 146 | ftsH | P0AAI3 | zinc | TRUE | FALSE | FALSE | FALSE |
| 147 | fur | P0A9A9 | zinc, iron | TRUE | FALSE | FALSE | FALSE |
| 148 | fusA | P0A6M8 | - | TRUE | FALSE | FALSE | FALSE |
| 149 | gcvP | P33195 | - | TRUE | FALSE | FALSE | FALSE |
| 150 | gcvR | P0A9I3 | - | TRUE | FALSE | FALSE | FALSE |
| 151 | gdhA | P00370 | - | TRUE | FALSE | FALSE | FALSE |
| 152 | ghrB | P37666 | - | TRUE | FALSE | FALSE | FALSE |
| 153 | glf | P37747 | - | TRUE | FALSE | FALSE | FALSE |
| 154 | glgP | P0AC86 | - | TRUE | FALSE | FALSE | FALSE |
| 155 | glnA | P0A9C5 | magnesium | TRUE | FALSE | FALSE | FALSE |
| 156 | gloC | P75849 | zinc | TRUE | FALSE | FALSE | FALSE |
| 157 | glpX | P0A9C9 | manganese | TRUE | FALSE | FALSE | FALSE |
| 158 | glsA1 | P77454 | - | TRUE | FALSE | FALSE | FALSE |
| 159 | gltX | P04805 | zinc | TRUE | FALSE | FALSE | FALSE |
| 160 | glyA | P0A825 | zinc | TRUE | FALSE | FALSE | FALSE |
| 161 | gpt | P0A9M5 | magnesium | TRUE | FALSE | FALSE | FALSE |
| 162 | groEL | P0A6F5 | magnesium | TRUE | FALSE | FALSE | FALSE |
| 163 | grpE | P09372 | - | TRUE | FALSE | FALSE | FALSE |
| 164 | grxB | P0AC59 | - | TRUE | FALSE | FALSE | FALSE |
| 165 | grxD | P0AC69 | iron | TRUE | FALSE | FALSE | FALSE |
| 166 | gstA | P0A9D2 | - | TRUE | FALSE | FALSE | FALSE |

Supplementary Table 5: Hit Overlaps *(continued)*

| # | Gene Name | UniProt ID | Bound Metal | LiP | TPP | SEC | <10 Å to active site |
|---|---|---|---|---|---|---|---|
| 167 | gyrA | P0AES4 | - | TRUE | FALSE | FALSE | FALSE |
| 168 | hemL | P23893 | - | TRUE | FALSE | FALSE | FALSE |
| 169 | hisC | P06986 | - | TRUE | FALSE | FALSE | FALSE |
| 170 | hrpA | P43329 | - | TRUE | FALSE | FALSE | FALSE |
| 171 | icd | P08200 | magnesium, manganese | TRUE | FALSE | FALSE | FALSE |
| 172 | ileS | P00956 | zinc | TRUE | FALSE | FALSE | FALSE |
| 173 | ilvB | P08142 | magnesium | TRUE | FALSE | FALSE | FALSE |
| 174 | iscA | P0AAC8 | iron | TRUE | FALSE | FALSE | FALSE |
| 175 | iscR | P0AGK8 | iron | TRUE | FALSE | FALSE | FALSE |
| 176 | iscU | P0ACD4 | copper, iron, zinc | TRUE | FALSE | FALSE | FALSE |
| 177 | ispB | P0AD57 | magnesium | TRUE | FALSE | FALSE | FALSE |
| 178 | ispF | P62617 | zinc, manganese | TRUE | FALSE | FALSE | FALSE |
| 179 | ispG | P62620 | iron | TRUE | FALSE | FALSE | FALSE |
| 180 | katE | P21179 | iron | TRUE | FALSE | FALSE | FALSE |
| 181 | katG | P13029 | iron | TRUE | FALSE | FALSE | FALSE |
| 182 | kgtP | P0AEX3 | - | TRUE | FALSE | FALSE | FALSE |
| 183 | letB | P76272 | - | TRUE | FALSE | FALSE | FALSE |
| 184 | lon | P0A9M0 | - | TRUE | FALSE | FALSE | FALSE |
| 185 | lpdA | P0A9P0 | zinc | TRUE | FALSE | FALSE | FALSE |
| 186 | luxS | P45578 | iron | TRUE | FALSE | FALSE | FALSE |
| 187 | lysC | P08660 | - | TRUE | FALSE | FALSE | FALSE |
| 188 | maeB | P76558 | magnesium, manganese | TRUE | FALSE | FALSE | FALSE |
| 189 | manA | P00946 | zinc | TRUE | FALSE | FALSE | FALSE |
| 190 | maoP | P0ADN2 | - | TRUE | FALSE | FALSE | FALSE |
| 191 | metF | P0AEZ1 | - | TRUE | FALSE | FALSE | FALSE |
| 192 | metG | P00959 | zinc | TRUE | FALSE | FALSE | FALSE |
| 193 | miaB | P0AEI1 | iron | TRUE | FALSE | FALSE | FALSE |
| 194 | minD | P0AEZ3 | - | TRUE | FALSE | FALSE | FALSE |
| 195 | mreB | P0A9X4 | - | TRUE | FALSE | FALSE | FALSE |
| 196 | mrp | P0AF08 | iron | TRUE | FALSE | FALSE | FALSE |
| 197 | msrC | P76270 | - | TRUE | FALSE | FALSE | FALSE |
| 198 | mukF | P60293 | calcium | TRUE | FALSE | FALSE | FALSE |
| 199 | murA | P0A749 | - | TRUE | FALSE | FALSE | FALSE |
| 200 | nfo | P0A6C1 | zinc, manganese | TRUE | FALSE | FALSE | FALSE |
| 201 | nikR | P0A6Z6 | nickel | TRUE | FALSE | FALSE | FALSE |
| 202 | nusA | P0AFF6 | - | TRUE | FALSE | FALSE | FALSE |
| 203 | pdeK | P37649 | - | TRUE | FALSE | FALSE | FALSE |
| 204 | pfkA | P0A796 | magnesium | TRUE | FALSE | FALSE | FALSE |
| 205 | pgk | P0A799 | - | TRUE | FALSE | FALSE | FALSE |
| 206 | pgl | P52697 | - | TRUE | FALSE | FALSE | FALSE |
| 207 | pheS | P08312 | magnesium | TRUE | FALSE | FALSE | FALSE |
| 208 | phoA | P00634 | magnesium, zinc | TRUE | FALSE | FALSE | FALSE |
| 209 | pldB | P07000 | - | TRUE | FALSE | FALSE | FALSE |
| 210 | pnp | P05055 | magnesium, manganese | TRUE | FALSE | FALSE | FALSE |
| 211 | ppa | P0A7A9 | magnesium, zinc | TRUE | FALSE | FALSE | FALSE |
| 212 | ppiB | P23869 | - | TRUE | FALSE | FALSE | FALSE |
| 213 | proS | P16659 | - | TRUE | FALSE | FALSE | FALSE |
| 214 | pta | P0A9M8 | zinc | TRUE | FALSE | FALSE | FALSE |
| 215 | purC | P0A7D7 | - | TRUE | FALSE | FALSE | FALSE |
| 216 | purD | P15640 | magnesium, manganese | TRUE | FALSE | FALSE | FALSE |
| 217 | purH | P15639 | - | TRUE | FALSE | FALSE | FALSE |
| 218 | puuE | P50457 | - | TRUE | FALSE | FALSE | FALSE |
| 219 | pyrB | P0A786 | - | TRUE | FALSE | FALSE | FALSE |
| 220 | pyrI | P0A7F3 | zinc | TRUE | FALSE | FALSE | FALSE |
| 221 | queC | P77756 | zinc | TRUE | FALSE | FALSE | FALSE |
| 222 | radD | P33919 | zinc | TRUE | FALSE | FALSE | FALSE |

Supplementary Table 5: Hit Overlaps *(continued)*

| # | Gene Name | UniProt ID | Bound Metal | LiP | TPP | SEC | <10 Å to active site |
|---|---|---|---|---|---|---|---|
| 223 | rapZ | P0A894 | - | TRUE | FALSE | FALSE | FALSE |
| 224 | rbsK | P0A9J6 | potassium, magnesium, manganese | TRUE | FALSE | FALSE | FALSE |
| 225 | recR | P0A7H6 | zinc | TRUE | FALSE | FALSE | FALSE |
| 226 | rffH | P61887 | magnesium | TRUE | FALSE | FALSE | FALSE |
| 227 | rnc | P0A7Y0 | magnesium | TRUE | FALSE | FALSE | FALSE |
| 228 | rne | P21513 | magnesium, zinc | TRUE | FALSE | FALSE | FALSE |
| 229 | rplA | P0A7L0 | - | TRUE | FALSE | FALSE | FALSE |
| 230 | rplI | P0A7R1 | - | TRUE | FALSE | FALSE | FALSE |
| 231 | rplK | P0A7J7 | - | TRUE | FALSE | FALSE | FALSE |
| 232 | rpmE | P0A7M9 | zinc | TRUE | FALSE | FALSE | FALSE |
| 233 | rpmH | P0A7P5 | - | TRUE | FALSE | FALSE | FALSE |
| 234 | rpmI | P0A7Q1 | - | TRUE | FALSE | FALSE | FALSE |
| 235 | rpmJ | P0A7Q6 | - | TRUE | FALSE | FALSE | FALSE |
| 236 | rpoA | P0A7Z4 | - | TRUE | FALSE | FALSE | FALSE |
| 237 | rpoC | P0A8T7 | zinc, magnesium | TRUE | FALSE | FALSE | FALSE |
| 238 | rpsA | P0AG67 | - | TRUE | FALSE | FALSE | FALSE |
| 239 | rpsD | P0A7V8 | - | TRUE | FALSE | FALSE | FALSE |
| 240 | rpsE | P0A7W1 | - | TRUE | FALSE | FALSE | FALSE |
| 241 | rpsG | P02359 | - | TRUE | FALSE | FALSE | FALSE |
| 242 | rpsI | P0A7X3 | - | TRUE | FALSE | FALSE | FALSE |
| 243 | rpsK | P0A7R9 | - | TRUE | FALSE | FALSE | FALSE |
| 244 | rpsM | P0A7S9 | - | TRUE | FALSE | FALSE | FALSE |
| 245 | rpsN | P0AG59 | - | TRUE | FALSE | FALSE | FALSE |
| 246 | rpsO | P0ADZ4 | - | TRUE | FALSE | FALSE | FALSE |
| 247 | rpsR | P0A7T7 | - | TRUE | FALSE | FALSE | FALSE |
| 248 | rpsS | P0A7U3 | - | TRUE | FALSE | FALSE | FALSE |
| 249 | rpsT | P0A7U7 | - | TRUE | FALSE | FALSE | FALSE |
| 250 | selO | P77649 | magnesium | TRUE | FALSE | FALSE | FALSE |
| 251 | slyD | P0A9K9 | nickel, copper, zinc, cobalt | TRUE | FALSE | FALSE | FALSE |
| 252 | sodA | P00448 | manganese | TRUE | FALSE | FALSE | FALSE |
| 253 | speA | P21170 | magnesium | TRUE | FALSE | FALSE | FALSE |
| 254 | speE | P09158 | - | TRUE | FALSE | FALSE | FALSE |
| 255 | sra | P68191 | - | TRUE | FALSE | FALSE | FALSE |
| 256 | suhB | P0ADG4 | magnesium, lithium | TRUE | FALSE | FALSE | FALSE |
| 257 | tabA | P0AF96 | - | TRUE | FALSE | FALSE | FALSE |
| 258 | tas | P0A9T4 | - | TRUE | FALSE | FALSE | FALSE |
| 259 | thrA | P00561 | sodium | TRUE | FALSE | FALSE | FALSE |
| 260 | thrB | P00547 | - | TRUE | FALSE | FALSE | FALSE |
| 261 | topA | P06612 | magnesium, zinc, manganese, calcium | TRUE | FALSE | FALSE | FALSE |
| 262 | trhO | P24188 | - | TRUE | FALSE | FALSE | FALSE |
| 263 | trmJ | P0AE01 | - | TRUE | FALSE | FALSE | FALSE |
| 264 | trpE | P00895 | magnesium | TRUE | FALSE | FALSE | FALSE |
| 265 | trxB | P0A9P4 | - | TRUE | FALSE | FALSE | FALSE |
| 266 | tsaD | P05852 | iron, magnesium | TRUE | FALSE | FALSE | FALSE |
| 267 | tsf | P0A6P1 | zinc | TRUE | FALSE | FALSE | FALSE |
| 268 | ttcA | P76055 | iron, magnesium | TRUE | FALSE | FALSE | FALSE |
| 269 | ubiD | P0AAB4 | manganese | TRUE | FALSE | FALSE | FALSE |
| 270 | uspD | P0AAB8 | - | TRUE | FALSE | FALSE | FALSE |
| 271 | uspG | P39177 | - | TRUE | FALSE | FALSE | FALSE |
| 272 | xthA | P09030 | magnesium, manganese | TRUE | FALSE | FALSE | FALSE |
| 273 | yaeH | P62768 | - | TRUE | FALSE | FALSE | FALSE |
| 274 | yahJ | P77554 | - | TRUE | FALSE | FALSE | FALSE |
| 275 | yahK | P75691 | zinc | TRUE | FALSE | FALSE | FALSE |
| 276 | ybcJ | P0AAS7 | - | TRUE | FALSE | FALSE | FALSE |

Supplementary Table 5: Hit Overlaps *(continued)*

| # | Gene Name | UniProt ID | Bound Metal | LiP | TPP | SEC | <10 Å to active site |
|---|---|---|---|---|---|---|---|
| 277 | ybeY | P0A898 | zinc, nickel | TRUE | FALSE | FALSE | FALSE |
| 278 | ybgI | P0AFP6 | divalent metal cation | TRUE | FALSE | FALSE | FALSE |
| 279 | ybiI | P41039 | zinc | TRUE | FALSE | FALSE | FALSE |
| 280 | ybjQ | P0A8C1 | - | TRUE | FALSE | FALSE | FALSE |
| 281 | yceD | P0AB28 | - | TRUE | FALSE | FALSE | FALSE |
| 282 | ycfP | P0A8E1 | - | TRUE | FALSE | FALSE | FALSE |
| 283 | ycgL | P0AB43 | - | TRUE | FALSE | FALSE | FALSE |
| 284 | ychJ | P37052 | - | TRUE | FALSE | FALSE | FALSE |
| 285 | yciF | P21362 | - | TRUE | FALSE | FALSE | FALSE |
| 286 | yciK | P31808 | - | TRUE | FALSE | FALSE | FALSE |
| 287 | ycjY | P76049 | - | TRUE | FALSE | FALSE | FALSE |
| 288 | yeaD | P39173 | - | TRUE | FALSE | FALSE | FALSE |
| 289 | yejK | P33920 | - | TRUE | FALSE | FALSE | FALSE |
| 290 | yfgJ | P76575 | - | TRUE | FALSE | FALSE | FALSE |
| 291 | yfhM | P76578 | - | TRUE | FALSE | FALSE | FALSE |
| 292 | yghU | Q46845 | - | TRUE | FALSE | FALSE | FALSE |
| 293 | ygiW | P0ADU5 | - | TRUE | FALSE | FALSE | FALSE |
| 294 | yhhW | P46852 | copper, iron, zinc, cobalt | TRUE | FALSE | FALSE | FALSE |
| 295 | yhhX | P46853 | - | TRUE | FALSE | FALSE | FALSE |
| 296 | yjdM | P0AFJ1 | - | TRUE | FALSE | FALSE | FALSE |
| 297 | yobA | P0AA57 | copper | TRUE | FALSE | FALSE | FALSE |
| 298 | glk | P0A6V8 | - | FALSE | TRUE | TRUE | - |
| 299 | nadR | P27278 | magnesium | FALSE | TRUE | TRUE | - |
| 300 | obgE | P42641 | magnesium | FALSE | TRUE | TRUE | - |
| 301 | phnO | P16691 | divalent metal cation | FALSE | TRUE | TRUE | - |
| 302 | rlmG | P42596 | - | FALSE | TRUE | TRUE | - |
| 303 | selB | P14081 | - | FALSE | TRUE | TRUE | - |
| 304 | bglX | P33363 | - | FALSE | TRUE | FALSE | - |
| 305 | cca | P06961 | magnesium, nickel | FALSE | TRUE | FALSE | - |
| 306 | cobB | P75960 | zinc | FALSE | TRUE | FALSE | - |
| 307 | corC | P0AE78 | cobalt, magnesium | FALSE | TRUE | FALSE | - |
| 308 | cpdA | P0AEW4 | iron | FALSE | TRUE | FALSE | - |
| 309 | cytR | P0ACN7 | - | FALSE | TRUE | FALSE | - |
| 310 | dapE | P0AED7 | zinc, cobalt | FALSE | TRUE | FALSE | - |
| 311 | dcp | P24171 | zinc, calcium | FALSE | TRUE | FALSE | - |
| 312 | dgoR | P31460 | zinc | FALSE | TRUE | FALSE | - |
| 313 | eptB | P37661 | calcium | FALSE | TRUE | FALSE | - |
| 314 | exuR | P0ACL2 | - | FALSE | TRUE | FALSE | - |
| 315 | fbp | P0A993 | magnesium | FALSE | TRUE | FALSE | - |
| 316 | fdhE | P13024 | iron | FALSE | TRUE | FALSE | - |
| 317 | frmA | P25437 | zinc | FALSE | TRUE | FALSE | - |
| 318 | ftnA | P0A998 | iron | FALSE | TRUE | FALSE | - |
| 319 | galT | P09148 | zinc, iron | FALSE | TRUE | FALSE | - |
| 320 | gatY | P0C8J6 | zinc | FALSE | TRUE | FALSE | - |
| 321 | glgA | P0A6U8 | - | FALSE | TRUE | FALSE | - |
| 322 | glmM | P31120 | magnesium | FALSE | TRUE | FALSE | - |
| 323 | glnD | P27249 | magnesium | FALSE | TRUE | FALSE | - |
| 324 | idi | Q46822 | manganese, magnesium, zinc | FALSE | TRUE | FALSE | - |
| 325 | leuS | P07813 | - | FALSE | TRUE | FALSE | - |
| 326 | mazG | P0AEY3 | magnesium | FALSE | TRUE | FALSE | - |
| 327 | metQ | P28635 | - | FALSE | TRUE | FALSE | - |
| 328 | moeB | P12282 | zinc | FALSE | TRUE | FALSE | - |
| 329 | mpaA | P0ACV6 | zinc | FALSE | TRUE | FALSE | - |
| 330 | mqsA | Q46864 | zinc | FALSE | TRUE | FALSE | - |
| 331 | mtfA | P76346 | zinc | FALSE | TRUE | FALSE | - |
| 332 | nagC | P0AF20 | - | FALSE | TRUE | FALSE | - |

Supplementary Table 5: Hit Overlaps *(continued)*

| # | Gene Name | UniProt ID | Bound Metal | LiP | TPP | SEC | <10 Å to active site |
|---|---|---|---|---|---|---|---|
| 333 | orn | P0A784 | zinc | FALSE | TRUE | FALSE | - |
| 334 | pckA | P22259 | calcium, manganese, magnesium | FALSE | TRUE | FALSE | - |
| 335 | phnP | P16692 | zinc, manganese | FALSE | TRUE | FALSE | - |
| 336 | polA | P00582 | - | FALSE | TRUE | FALSE | - |
| 337 | prfB | P07012 | - | FALSE | TRUE | FALSE | - |
| 338 | prlC | P27298 | zinc | FALSE | TRUE | FALSE | - |
| 339 | rbn | P0A8V0 | zinc | FALSE | TRUE | FALSE | - |
| 340 | rihA | P41409 | calcium | FALSE | TRUE | FALSE | - |
| 341 | rnb | P30850 | magnesium | FALSE | TRUE | FALSE | - |
| 342 | rng | P0A9J0 | magnesium | FALSE | TRUE | FALSE | - |
| 343 | rsgA | P39286 | zinc | FALSE | TRUE | FALSE | - |
| 344 | tadA | P68398 | zinc | FALSE | TRUE | FALSE | - |
| 345 | tdh | P07913 | zinc, cobalt, cadmium, iron, manganese | FALSE | TRUE | FALSE | - |
| 346 | tgt | P0A847 | zinc | FALSE | TRUE | FALSE | - |
| 347 | thrS | P0A8M3 | zinc | FALSE | TRUE | FALSE | - |
| 348 | tilS | P52097 | - | FALSE | TRUE | FALSE | - |
| 349 | uxuR | P39161 | - | FALSE | TRUE | FALSE | - |
| 350 | ycaO | P75838 | magnesium | FALSE | TRUE | FALSE | - |
| 351 | ycaR | P0AAZ7 | - | FALSE | TRUE | FALSE | - |
| 352 | yegU | P76418 | - | FALSE | TRUE | FALSE | - |
| 353 | yfcE | P67095 | manganese | FALSE | TRUE | FALSE | - |
| 354 | yhbS | P63417 | - | FALSE | TRUE | FALSE | - |
| 355 | yjbQ | P0AF48 | - | FALSE | TRUE | FALSE | - |
| 356 | aidB | P33224 | - | FALSE | FALSE | TRUE | - |
| 357 | cspE | P0A972 | - | FALSE | FALSE | TRUE | - |
| 358 | dam | P0AEE8 | - | FALSE | FALSE | TRUE | - |
| 359 | darP | P0A8X0 | - | FALSE | FALSE | TRUE | - |
| 360 | dcm | P0AED9 | - | FALSE | FALSE | TRUE | - |
| 361 | eutC | P19636 | cobalt | FALSE | FALSE | TRUE | - |
| 362 | fbaB | P0A991 | - | FALSE | FALSE | TRUE | - |
| 363 | fecA | P13036 | iron | FALSE | FALSE | TRUE | - |
| 364 | ffh | P0AGD7 | - | FALSE | FALSE | TRUE | - |
| 365 | fnr | P0A9E5 | iron | FALSE | FALSE | TRUE | - |
| 366 | folE | P0A6T5 | zinc | FALSE | FALSE | TRUE | - |
| 367 | glgB | P07762 | - | FALSE | FALSE | TRUE | - |
| 368 | groES | P0A6F9 | metal cation | FALSE | FALSE | TRUE | - |
| 369 | hflX | P25519 | magnesium | FALSE | FALSE | TRUE | - |
| 370 | ibaG | P0A9W6 | iron | FALSE | FALSE | TRUE | - |
| 371 | ihfB | P0A6Y1 | - | FALSE | FALSE | TRUE | - |
| 372 | lipA | P60716 | iron | FALSE | FALSE | TRUE | - |
| 373 | livF | P22731 | - | FALSE | FALSE | TRUE | - |
| 374 | mlaF | P63386 | - | FALSE | FALSE | TRUE | - |
| 375 | nagD | P0AF24 | magnesium, manganese, cobalt, zinc | FALSE | FALSE | TRUE | - |
| 376 | nagZ | P75949 | - | FALSE | FALSE | TRUE | - |
| 377 | nudI | P52006 | magnesium | FALSE | FALSE | TRUE | - |
| 378 | proQ | P45577 | - | FALSE | FALSE | TRUE | - |
| 379 | pssA | P23830 | - | FALSE | FALSE | TRUE | - |
| 380 | pxpA | P75746 | - | FALSE | FALSE | TRUE | - |
| 381 | rbfA | P0A7G2 | - | FALSE | FALSE | TRUE | - |
| 382 | rho | P0AG30 | - | FALSE | FALSE | TRUE | - |
| 383 | ribE | P61714 | - | FALSE | FALSE | TRUE | - |
| 384 | rimM | P0A7X6 | - | FALSE | FALSE | TRUE | - |
| 385 | rimP | P0A8A8 | - | FALSE | FALSE | TRUE | - |
| 386 | rlmE | P0C0R7 | magnesium | FALSE | FALSE | TRUE | - |

Supplementary Table 5: Hit Overlaps *(continued)*

| # | Gene Name | UniProt ID | Bound Metal | LiP | TPP | SEC | <10 Å to active site |
|---|---|---|---|---|---|---|---|
| 387 | rlmJ | P37634 | - | FALSE | FALSE | TRUE | - |
| 388 | rlmL | P75864 | - | FALSE | FALSE | TRUE | - |
| 389 | rluC | P0AA39 | - | FALSE | FALSE | TRUE | - |
| 390 | rnm | P77766 | manganese | FALSE | FALSE | TRUE | - |
| 391 | rnpA | P0A7Y8 | - | FALSE | FALSE | TRUE | - |
| 392 | rplT | P0A7L3 | - | FALSE | FALSE | TRUE | - |
| 393 | rplU | P0AG48 | - | FALSE | FALSE | TRUE | - |
| 394 | rpmC | P0A7M6 | - | FALSE | FALSE | TRUE | - |
| 395 | rppH | P0A776 | magnesium, zinc, manganese | FALSE | FALSE | TRUE | - |
| 396 | rpsB | P0A7V0 | zinc | FALSE | FALSE | TRUE | - |
| 397 | rsmB | P36929 | - | FALSE | FALSE | TRUE | - |
| 398 | selD | P16456 | magnesium | FALSE | FALSE | TRUE | - |
| 399 | smpB | P0A832 | - | FALSE | FALSE | TRUE | - |
| 400 | sucA | P0AFG3 | magnesium | FALSE | FALSE | TRUE | - |
| 401 | tdk | P23331 | zinc | FALSE | FALSE | TRUE | - |
| 402 | tmaR | P0A8M6 | - | FALSE | FALSE | TRUE | - |
| 403 | trmA | P23003 | - | FALSE | FALSE | TRUE | - |
| 404 | udk | P0A8F4 | - | FALSE | FALSE | TRUE | - |
| 405 | uvrA | P0A698 | zinc | FALSE | FALSE | TRUE | - |
| 406 | waaB | P27127 | - | FALSE | FALSE | TRUE | - |
| 407 | xynR | P77300 | - | FALSE | FALSE | TRUE | - |
| 408 | yadG | P36879 | - | FALSE | FALSE | TRUE | - |
| 409 | ybeD | P0A8J4 | - | FALSE | FALSE | TRUE | - |
| 410 | ybhA | P21829 | magnesium, manganese, cobalt, zinc | FALSE | FALSE | TRUE | - |
| 411 | ybiB | P30177 | - | FALSE | FALSE | TRUE | - |
| 412 | ybjD | P75828 | - | FALSE | FALSE | TRUE | - |
| 413 | ycbB | P22525 | - | FALSE | FALSE | TRUE | - |
| 414 | ycdY | P75915 | - | FALSE | FALSE | TRUE | - |
| 415 | yciH | P08245 | - | FALSE | FALSE | TRUE | - |
| 416 | yegP | P76402 | - | FALSE | FALSE | TRUE | - |
| 417 | yehS | P33355 | - | FALSE | FALSE | TRUE | - |
| 418 | ygiF | P30871 | metal cation | FALSE | FALSE | TRUE | - |
| 419 | yheV | P0ADW8 | - | FALSE | FALSE | TRUE | - |
| 420 | yibL | P0ADK8 | - | FALSE | FALSE | TRUE | - |
| 421 | yjaG | P32680 | - | FALSE | FALSE | TRUE | - |
| 422 | yjgM | P39337 | - | FALSE | FALSE | TRUE | - |
| 423 | yjhU | P39356 | - | FALSE | FALSE | TRUE | - |
| 424 | yncE | P76116 | - | FALSE | FALSE | TRUE | - |

Supplementary Table 6: SEC-MS Hits

| # | Gene Name | UniProt ID | Bound Metal | Ribosome | Binds Nucleotides | Binds DNA/RNA |
|---|---|---|---|---|---|---|
| 1 | rplB | P60422 | zinc | + | - | + |
| 2 | rplC | P60438 | - | + | - | + |
| 3 | rplD | P60723 | - | + | - | + |
| 4 | rplE | P62399 | - | + | - | + |
| 5 | rplF | P0AG55 | - | + | - | + |
| 6 | rplJ | P0A7J3 | - | + | - | + |
| 7 | rplM | P0AA10 | zinc | + | - | + |
| 8 | rplN | P0ADY3 | - | + | - | + |
| 9 | rplO | P02413 | - | + | - | + |
| 10 | rplR | P0C018 | - | + | - | + |
| 11 | rplT | P0A7L3 | - | + | - | + |
| 12 | rplU | P0AG48 | - | + | - | + |
| 13 | rplV | P61175 | - | + | - | + |
| 14 | rplW | P0ADZ0 | - | + | - | + |
| 15 | rplX | P60624 | - | + | - | + |
| 16 | rplY | P68919 | - | + | - | + |
| 17 | rpmA | P0A7L8 | - | + | - | + |
| 18 | rpmB | P0A7M2 | - | + | - | + |
| 19 | rpmC | P0A7M6 | - | + | - | + |
| 20 | rpmG | P0A7N9 | - | + | - | + |
| 21 | rpsC | P0A7V3 | - | + | - | + |
| 22 | rpsH | P0A7W7 | - | + | - | + |
| 23 | rpsJ | P0A7R5 | - | + | - | + |
| 24 | rpsL | P0A7S3 | - | + | - | + |
| 25 | rpsP | P0A7T3 | - | + | - | + |
| 26 | rpsU | P68679 | - | + | - | + |
| 27 | rplQ | P0AG44 | - | + | - | - |
| 28 | rpmD | P0AG51 | - | + | - | - |
| 29 | rpmF | P0A7N4 | - | + | - | - |
| 30 | rpsB | P0A7V0 | zinc | + | - | - |
| 31 | ffh | P0AGD7 | - | - | + | + |
| 32 | hflX | P25519 | magnesium | - | + | + |
| 33 | nadR | P27278 | magnesium | - | + | + |
| 34 | obgE | P42641 | magnesium | - | + | + |
| 35 | rho | P0AG30 | - | - | + | + |
| 36 | selB | P14081 | - | - | + | + |
| 37 | thiI | P77718 | iron | - | + | + |
| 38 | uvrA | P0A698 | zinc | - | + | + |
| 39 | folE | P0A6T5 | zinc | - | + | - |
| 40 | glk | P0A6V8 | - | - | + | - |
| 41 | groES | P0A6F9 | metal cation | - | + | - |
| 42 | livF | P22731 | - | - | + | - |
| 43 | metK | P0A817 | magnesium, potassium, manganese, cobalt | - | + | - |
| 44 | mlaF | P63386 | - | - | + | - |
| 45 | pxpA | P75746 | - | - | + | - |
| 46 | selD | P16456 | magnesium | - | + | - |
| 47 | tdk | P23331 | zinc | - | + | - |
| 48 | udk | P0A8F4 | - | - | + | - |
| 49 | yadG | P36879 | - | - | + | - |
| 50 | ybjD | P75828 | - | - | + | - |
| 51 | aidB | P33224 | - | - | - | + |
| 52 | bolA | P0ABE2 | - | - | - | + |
| 53 | cspE | P0A972 | - | - | - | + |
| 54 | dam | P0AEE8 | - | - | - | + |
| 55 | darP | P0A8X0 | - | - | - | + |
| 56 | dcm | P0AED9 | - | - | - | + |

Supplementary Table 6: SEC-MS Hits *(continued)*

| # | Gene Name | UniProt ID | Bound Metal | Ribosome | Binds Nucleotides | Binds DNA/RNA |
|---|---|---|---|---|---|---|
| 57 | fnr | P0A9E5 | iron | - | - | + |
| 58 | ihfB | P0A6Y1 | - | - | - | + |
| 59 | proQ | P45577 | - | - | - | + |
| 60 | rimM | P0A7X6 | - | - | - | + |
| 61 | rlmI | P75876 | - | - | - | + |
| 62 | rlmJ | P37634 | - | - | - | + |
| 63 | rlmL | P75864 | - | - | - | + |
| 64 | rluC | P0AA39 | - | - | - | + |
| 65 | rnpA | P0A7Y8 | - | - | - | + |
| 66 | rsmB | P36929 | - | - | - | + |
| 67 | smpB | P0A832 | - | - | - | + |
| 68 | trmA | P23003 | - | - | - | + |
| 69 | xynR | P77300 | - | - | - | + |
| 70 | ybiB | P30177 | - | - | - | + |
| 71 | yjhU | P39356 | - | - | - | + |
| 72 | yncE | P76116 | - | - | - | + |
| 73 | cmoA | P76290 | - | - | - | - |
| 74 | eutC | P19636 | cobalt | - | - | - |
| 75 | fbaB | P0A991 | - | - | - | - |
| 76 | fecA | P13036 | iron | - | - | - |
| 77 | glgB | P07762 | - | - | - | - |
| 78 | ibaG | P0A9W6 | iron | - | - | - |
| 79 | lipA | P60716 | iron | - | - | - |
| 80 | nagD | P0AF24 | magnesium, manganese, cobalt, zinc | - | - | - |
| 81 | nagZ | P75949 | - | - | - | - |
| 82 | nudI | P52006 | magnesium | - | - | - |
| 83 | phnO | P16691 | divalent metal cation | - | - | - |
| 84 | pssA | P23830 | - | - | - | - |
| 85 | rbfA | P0A7G2 | - | - | - | - |
| 86 | ribE | P61714 | - | - | - | - |
| 87 | rimP | P0A8A8 | - | - | - | - |
| 88 | rlmE | P0C0R7 | magnesium | - | - | - |
| 89 | rlmG | P42596 | - | - | - | - |
| 90 | rnm | P77766 | manganese | - | - | - |
| 91 | rppH | P0A776 | magnesium, zinc, manganese | - | - | - |
| 92 | rsfS | P0AAT6 | - | - | - | - |
| 93 | sucA | P0AFG3 | magnesium | - | - | - |
| 94 | tmaR | P0A8M6 | - | - | - | - |
| 95 | waaB | P27127 | - | - | - | - |
| 96 | ybeD | P0A8J4 | - | - | - | - |
| 97 | ybhA | P21829 | magnesium, manganese, cobalt, zinc | - | - | - |
| 98 | ycbB | P22525 | - | - | - | - |
| 99 | ycdY | P75915 | - | - | - | - |
| 100 | ycfH | P0AFQ7 | cobalt, manganese, nickel | - | - | - |
| 101 | yciH | P08245 | - | - | - | - |
| 102 | yegP | P76402 | - | - | - | - |
| 103 | yehS | P33355 | - | - | - | - |
| 104 | ygiF | P30871 | metal cation | - | - | - |
| 105 | yheV | P0ADW8 | - | - | - | - |
| 106 | yibL | P0ADK8 | - | - | - | - |
| 107 | yjaG | P32680 | - | - | - | - |
| 108 | yjgM | P39337 | - | - | - | - |
